# Abnormal features of human self-domestication in bipolar disorder

**DOI:** 10.1101/2020.04.28.065581

**Authors:** Antonio Benítez-Burraco, Ethan Hansen

## Abstract

Bipolar disorder (BD) is a severe mental condition characterized by episodes of elevated mood and depression. Being a heritable condition, it features a complex genetic architecture, and it is not still clear how genes contribute to the onset and course of the disease. In this paper we adopted an evolutionary-genomic approach to this condition, focusing on changes occurring during human evolution as a source of our distinctive cognitive and behavioral phenotype. We show clinical evidence that the BD phenotype can be construed as an abnormal presentation of the human self-domestication phenotype. We further demonstrate that candidate genes for BD significantly overlap with candidates for mammal domestication, and that this common set of genes is enriched in functions that are important for the BD phenotype, especially neurotransmitter homeostasis. Finally, we show that candidates for domestication are differentially expressed in brain regions involved in BD pathology, particularly, the hippocampus and the prefrontal cortex. Overall, this link between domestication and BD should facilitate a better understanding of the BD etiopathology.

## Introduction

Bipolar disorder (BD) is a severe psychiatric condition characterized by recurrent episodes of elevated mood (mania) and depression, together with changes in activity levels, with depression being usually more common and longer lasting than mania (Judd et al., 2008; Anderson et al., 2012). BD is one of the most heritable psychiatric conditions, but like similar diseases, it exhibits a complex genetic architecture, with dozens of candidate genes and risk factors that overlap with the genetic determinants of other high-prevalence cognitive conditions, especially schizophrenia (SZ) and major depression (see Gordovez and McMahon, 2020 for a recent review). These findings are in line with the so-called ‘omnigenic’ theories of complex diseases, according to which they mostly result from changes in the expression pattern of most of the genes expressed in the affected tissues, but not from the dysfunction of single genes, with a strong effect on selected pathways that drive disease etiology (Boyle et al., 2017; Peedicayil and Grayson, 2018a,b). Still, one can expect that certain regulatory pathways or biological mechanisms are more affected than others and ultimately that intermediate, disease-specific phenotypes can be identified with finite genetic determinants (Jakobson and Jarosz, 2019). As noted by Gordovez and McMahon (2020), this would be also the case with BD, as some specific biological pathways are expected to be differentially impaired in this condition.

One promising approach to make more tractable the genetic study of complex diseases like BD is adopting an evolutionary perspective. A robust link exists between evolution and abnormal development, with genes positively selected in our species being enriched in candidates for high-prevalence, complex cognitive conditions like SZ (Srinivasan et al., 2016) or autism spectrum disorders (ASD) (Polimanti and Gelernter, 2017). Even in conditions with a neat genetic origin, like Williams syndrome (WS), abnormally expressed genes outside the WS region are enriched in protein-coding genes that have been positively selected in modern humans compared to extinct hominins (Benítez-Burraco, 2020a). This overlap between genes selected in humans (and seemingly accounting for the evolution of the human brain, human cognition, and human behavior) and genes causing complex cognitive conditions seems to be explained by the fact that recently evolved components of the human phenotype are less resistant to gene mutations (and more generally, to developmental perturbations) because they lack the robust compensatory mechanisms to damage commonly found in biological functions shaped by selective pressures that have acted over longer periods of time (see Toro et al., 2010 for a useful discussion on ASD). Ultimately, recent changes in our genome might have uncovered (or ‘decanalized’) widespread cryptic variation, ultimately resulting in complex disease (see Gibson 2009 for details). As a consequence, genomic changes resulting in evolutionary advantages persist even if they also cause diseases.

Overall, this is the evolutionary-genomic approach we have adopted in this paper to gain some insight into the molecular etiopathogenesis of BD. Specifically, considering that BD impacts key aspects of our species-specific distinctive cognitive and behavioral phenotype, we have built on a particular hypothesis about the origins of our cognition and behavior, namely, the self-domestication account of human evolution. Briefly stated, humans are hypothesized to have experienced a domestication process similar to that underlying the evolution of domesticated strains of mammals, except that it was driven by intra-species factors such as co-parenting, changes in human foraging ecology, or the rise of community living (Hare et al., 2012; Pisor and Surbeck, 2019). This process of self-domestication is thought to have facilitated the emergence of many of our species-specific biological, cognitive, and behavioral features, including our sophisticated technology and culture (see Hare, 2017 for review). This hypothesis is supported by findings in humans, compared to extant primates and extinct hominins, of many features commonly exhibited by domesticates, including smaller skulls/brains, juvenile-like faces, hairlessness, paedomorphic features, a prolonged juvenile period, less marked sexual dimorphism, reduced reactive aggression, increased prosocial behavior, and increased play behavior, among others (Shea, 1989; Leach, 2003; Somel et al., 2009; Zollikofer and Ponce de León, 2010; Herrmann et al., 2011; Plavcan, 2012; Stringer, 2016; see Hare, 2017 for review). In further support of this view, regions positively selected in modern humans compared to extinct hominins are enriched in candidate genes for domestication in mammals (Theofanopoulou et al., 2017). Interestingly, complex genetic conditions involving cognitive deficits, behavioral anomalies, and abnormal socialization patterns, like SZ and ASD, are characterized by abnormal presentations of traits associated with (self-)domestication; moreover, candidate genes for domestication are overrepresented among the candidates for these conditions and/or exhibit altered expression profiles in the brains of affected people (Benítez-Burraco et al., 2016; Benítez-Burraco et al., 2017; Benítez-Burraco, 2020b). But the same pattern holds in conditions with a neat molecular etiology, like WS, as genes outside the affected genomic region that are differentially-expressed in patients are likewise enriched in candidates for domestication (Niego and Benítez-Burraco, 2019). Overall, this supports the view that a deep link exists between cognitive disease and self-domestication (as a crucial aspect of human evolution) and specifically, that examining the presentation of (self-)domestication features in people with BD can help unravel the etiology of this condition.

The paper is structured as follows. First, we provide a detailed characterization of features of self-domestication in subjects with BD, with a focus on their distinctive physical, cognitive, and behavioral clinical profile. Second, we examine the overlap between candidate genes for BD and domestication. We provide a functional characterization of the common genes as well as some hints about their expression pattern in the brains of affected people. We conclude with a discussion about the suitability of construing BD as an abnormal ontogenetic itinerary for human cognition and behavior, resulting in part from changes in genes involved in domestication.

## 2. Domestication features in BD

As noted in the previous section, domestication results in a set of morphological, neuroendocrine, and behavioral traits that usually co-occur. This is the ‘domestication syndrome’, hypothesized as stemming from a selection for reduced stress/reactivity (‘tameness’) leading to, in effect, mild loss-of-function of the neural crest (Wilkins et al., 2014; but see Sánchez-Villagra et al., 2016 for a critical view). Features include reduction in brain/cranial capacity, floppy ears, smaller teeth, docile/juvenile behavior, shorter muzzles, and altered reproductive cycles, as well as underlying neural crest and HPA axis changes, when comparing domesticated animals to their wild counterparts (Wilkins et al., 2014). As we discussed in the previous section, many of these traits are also found in modern humans compared to extinct hominins and most extant primates. As we show below, most of these features are also found altered in patients with BD, with some of them being attenuated, while others appear augmented (Figure 1).

**Figure 1.**
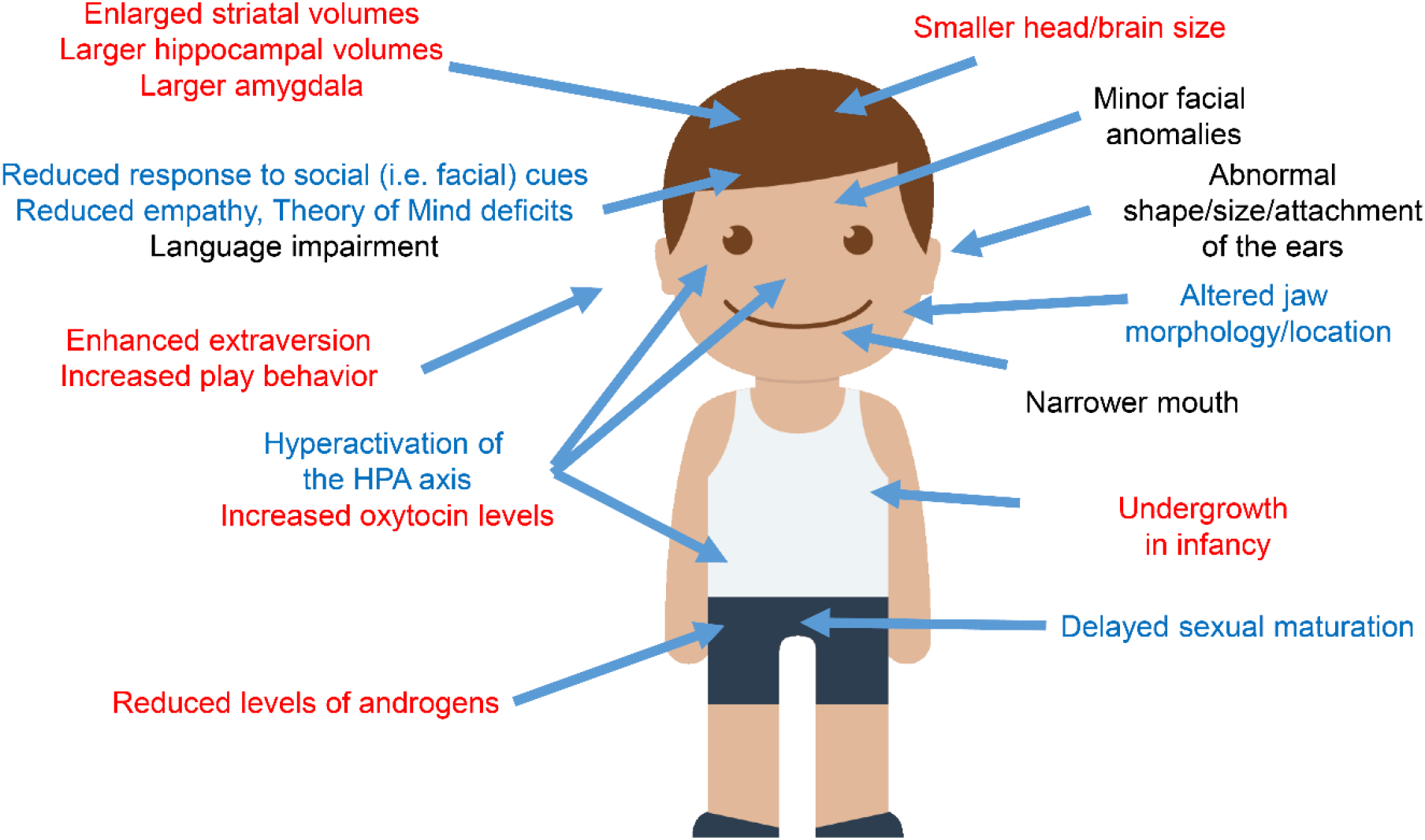
Domestication features in subjects with BD. Features found exacerbated are colored in red, whereas traits found attenuated are highlighted in blue. The picture of the child was gathered from Iconfinder output (available at http://www.iconfinder.com/icons/525448/boy_child_kid_male_man_person_white_icon).

### Minor physical anomalies

Minor physical anomalies (MPAs) are more common in those with neurodevelopmental disorders and are considered to reflect altered brain development given the common neuroectodermal origin of the brain and skin, with shared sensitivity to genetic and/or environmental perturbation during embryogenesis (Myers et al., 2017). Evidence surrounding MPAs in BD is mixed (Tényi et al., 2009), with some studies showing no significant difference in total MPA rate compared to SZ, others showing no significant difference compared to healthy controls (Sanches et al., 2008), and some indicating increased MPA rate within specific body regions (Akabaliev et al., 2011; Berecz et al., 2017). When comparing patients with affective psychosis to those with non-affective psychosis and healthy controls, no significant differences in total MPA scores were found by McGrath et al. (2002), but those with affective psychosis (including BD) were found to have wider skulls, shorter lower thirds of the face, and more brachycephalic skulls, reminiscent of the domestication syndrome. Significant increases in dermatoglyphic malformations such as ridge dissociation have also been reported (Gutiérrez et al., 1998) (though the evidence in this area is also mixed; see Vonk et al., 2014) and were associated with earlier age of onset of BD. Increased dermatoglyphic malformations and genetic correlations led Vonk et al. (2014) to conclude that smaller brain volumes in those at risk for BD are related to abnormal development of fetal ectoderm, from which both brain and skin tissue are derived, between the tenth and fifteenth weeks of gestation. Specific examples or areas of statistically significantly increased MPA in BD include furrowed tongue, enlarged gap between toes I and II, high or steepled palate, and lowered eye fissures, with abnormal ear size/shape/attachment, flat occiput, and other MPAs shared with SZ patients (Trixler et al., 2001; Lloyd et al., 2008; Akabaliev et al., 2011; Berecz et al., 2017). Surface imaging of BD patients compared with SZ patients and controls by 3D laser describes anomalies found in male and female patients with BD including widening of the nose and face overall, narrowing of the mouth, and chin displacement upward, as well as sex-specific changes in nose length (shorter in females, longer in males) (Hennessy et al., 2010). Compared to controls and patients with SZ, the jaws of patients with BD were wider (in males) and upwardly displaced (in females). Additionally, patients with BD and SZ were found to share common, especially frontonasal dysmorphology, such as mouth narrowing. In a study demonstrating significant differences between the three groups, a pattern emerges with decreasing levels of (especially cephalofacial) MPAs from SZ, to BD, to normal controls, suggesting ‘a continuum of neurodevelopmental adversity’ to match the clinical spectrum of psychotic disorders (Akabaliev et al., 2014).

### Brain and cognition

In BD and SZ, total cerebral volume is decreased, suggesting a shared neurodevelopmental etiology (Nasrallah, 1991); this finding is significantly associated with BD duration (Hallahan et al., 2011). The brain in patients with BD is also notable for increased amygdala volume (Altshuler et al., 1998, 2000; Strakowski et al., 1999; Hallahan et al., 2011), decreased hippocampus volume (Hajek et al., 2012; Han et al., 2019) (though increased with lithium treatment according to Hallahan et al., 2011), increased ventricle:brain ratio (Post et al., 2003), enlarged caudate nuclei (Noga et al., 2001), and increased right putamen volumes (Hallahan et al., 2011). As noted, the brain of domesticated animals is smaller than the brain of their wild conspecific. Contrary to people with BD, some domesticated animals like domesticated rats exhibit reduced volumes of the striatal regions (Kruska and Schott, 1977). By contrast, and like subjects with BD, many domesticated mammals (laboratory rats, pigs, sheep, poodles and llamas) exhibit reduced hippocampal volumes (and seemingly, hippocampal function) as well as increased amygdala volumes (Kruska, 1988, 2005). Additionally, bonobos—another species thought to have self-domesticated—have a larger amygdala compared to chimpanzees (Rilling et al., 2012). The amygdala and the hippocampus are part of the limbic system, which regulates important behavioral and cognitive functions such as emotion, motivation, and long-term memory (see Rolls, 2015 for review).

Evidence as to impairment of social cognition in BD is mixed: on one measure of social cognition (MSCEIT), patients with SZ evidenced social cognition deficits compared to controls, while those with BD did not—even separately assessing BD with and without psychotic features (Nitzburg et al., 2015). On the other hand, theory of mind deficits as assessed with the MASC tool have been found in patients with BD and their first-degree relatives (Santos et al., 2017). Interestingly, theory of mind deficits in BD are most severe during the manic phase, especially if psychotic symptoms are present (Hawken et al., 2016). Furthermore, one aspect of social cognition—facial emotion recognition (FER)—has been found to be a moderate, stable deficit in BD in a quantitative review by Kohler et al. (2011). This deficit can be seen across mood states in both adults and children with BD as well as children with first-degree relatives with BD (Brotman et al., 2008; Van Rheenen and Rossell, 2014). As assessed with fMRI, patients with BD displayed greater amygdalar activation, compared to healthy controls, when processing fearful facial expressions (Yurgelun-Todd et al., 2000). Empathic responses by self-report in BD have been shown to be decreased relative to controls (and higher relative to patients with SZ), though actual performance on empathy behavioral tasks was similar to that of controls (Derntl et al., 2012; Seidel et al., 2012), consistent with previous findings of reduced cognitive empathy (perspective taking) and increased personal distress at others’ negative experiences in BD (Cusi et al., 2010). Another aspect of social cognition, theory of mind, has been reported to be impaired in BD—in symptomatic (both depressed and manic) patients (Kerr et al., 2003), as well as in euthymic and remitted patients (Bora et al., 2005), with greater impairment seen in cognitive than emotional theory of mind (Barrera et al., 2013). Contrary to people with BD, domesticated animals (dogs vs. wolves, domestic vs. wild foxes, and bonobos vs. chimpanzees) show more sensitivity and attentiveness to human social cues, like eye or facial movements (Hare, 2017).

Finally, regarding language (dis)abilities, it should be noted that human self-domestication has been proposed as a key facilitator of language evolution (Benítez-Burraco and Kempe, 2018; Thomas and Kirby, 2018). Language impairment is a central if overlooked component of BD, with meta-analysis indicating verbal fluency as moderately impaired, with significantly greater impairment in euthymic than manic patients (Raucher-Chéné et al., 2017). Language differences in BD also include impaired accuracy when emotionally labeling ‘happy intonations’ in males (Van Rheenen and Rossell, 2013) and abnormal prefrontal activation during language tasks (Curtis et al., 2007).

### Behavioral traits and neuroendocrine impairments

The hypothalamic–pituitary–adrenal (HPA) axis, which regulates a great number of body functions and is thought to underlie stress reactivity changes in domestication, has been shown to be hyperactive in patients with BD, with hypersecretion of cortisol compared to controls (Cervantes et al., 2001; Girshkin et al., 2014). HPA axis dysfunction has been suggested as a trait abnormality in BD given enhanced cortisol response in the dexamethasone/corticotropin-releasing hormone test relative to healthy controls, without significant difference between remitted and non-remitted patients (Watson et al., 2004). Basal plasma adrenocorticotropic hormone (ACTH) levels are also elevated in BD, according to a 2016 meta-analysis (Belvederi Murri et al.), and glucocorticoid receptor responsiveness is reduced (Fries et al., 2014). This HPA axis dysfunction has been hypothesized to underlie the cyclicity of BD mood phases by Daban et al. (2005), who suggest that increases in ACTH and cortisol precede manic states, while depressive states may be caused by chronically elevated cortisol levels. In support of this, HPA axis hyperactivity in BD appears to be most pronounced during mania (Belvederi Murri et al., 2016). HPA axis dysfunction also appears to factor into the illness progression of BD, as post-dexamethasone cortisol levels—elevated in patients with BD compared to controls overall—positively correlate with total number of mood episodes experienced (Fries et al., 2014). By contrast, in domesticated animals the function of the HPA axis is significantly reduced compared to wild conspecifics, resulting in decreased levels of glutocorticoids, basal plasma ACTH levels, and adrenal response to stress, as well as less pronounced cortisol response to novelties (Naumenko and Belyaev, 1980; Kruska 1988; Kunzl and Sachser, 1999; Trut et al., 2009; Zipser et al., 2014; Kaiser et al., 2015).

Relatedly, oxytocin (OT) and vasopressin (AVP; also known as antidiuretic hormone, ADH) have been proposed as factors in the etiology of BD as well as, in different ways, SZ, WS, and ASD (Dai et al., 2012; Perez-Rodriguez et al., 2015; Iovino et al., 2018). OT has been described as a driver of social cognition and mentalization (Striepens et al., 2011; Wojciak et al., 2012; Crespi, 2016), including the multimodality that characterizes higher-order linguistic abilities (Theofanopoulou 2016). Specifically, OT inhibits the HPA axis’ stress triggered activity (Neumann, 2002). While Rutigliano et al. (2016) found insufficient evidence for use of peripheral OT/AVP levels as biomarkers of psychiatric disorders, OT has been found to be elevated in BD II compared to MDD and controls, pre- and post-treatment (Lien et al., 2017). This is consistent with the findings of Turan et al. (2013) in a study that also found serum OT to be highest in currently manic BD patients, compared to depressed and remitted BD patients (as well as healthy controls). Domesticated animals exhibit higher densities of both OT and AVP cells (Ruan and Zhang, 2016). Oxytocin has been associated to human-animal interactions, especially when eye contact is involved in communicative settings (see Beetz et al., 2012 for review).

Inasmuch as domestication entails a selection for hypersociality and low reactivity—in effect, tameness, the basis of selection in the famous fox farm experiment (Trut, 1999)—personality traits such extraversion and openness are worth noting. For instance, WS has been proposed as a condition of hyperdomestication and is characterized by hypersociability (Niego and Benítez-Burraco, 2019), with the genetic locus involved being correlated with hypersociability in dogs compared to wolves (vonHoldt et al., 2017). Perhaps not surprisingly, in subjects with WS, basal OT and AVP levels are increased, and the increase correlates with the tendency to approach strangers and emotionality (Dai et al., 2012). Reflecting the gregariousness associated with the manic state, trait extraversion and openness have been associated with hypomanic personality traits (Meyer, 2002; Durbin et al., 2009). Mania has also been correlated with trait extraversion, as well as neuroticism and negatively, agreeableness (Quilty et al., 2009). Finally, playful thinking has been associated with mania, compared to SZ, as measured on the Thought Disorder Index (Daniels et al., 1988). According to Kisko et al. (2018), CACNA1C-haploinsufficient rats exhibit deficits in prosocial 50-kHz ultrasonic vocalizations famously thought to reflect laughter due to association with play (Panksepp, 2005). This deficit was thought to reflect the derivation of less reward from social play from the haploinsufficient animals. Domestication is known to increase play behavior in animals (Hart, 1985; Himmler et al., 2013; Kaiser et al., 2015).

### Other features

With neoteny a classic feature in the domestication syndrome, it is notable that small head circumference (less than 32 cm at birth) has been found to correlate with later diagnosis of BD by Pugliese et al. (2019); inadequate maternal weight gain was inversely associated with BD (while positively associated with SZ). Considering another aspect of neoteny, evidence is mixed as to whether age at menarche is delayed (Bisaga et al., 2002) or not significantly different (Dunjic-Kostic et al., 2016) in women with BD compared to healthy controls (Williams et al., 2007; Tondo et al., 2017). (Further complicating the picture, girls with early menarche experience higher rates of BD and other psychiatric conditions (Platt et al., 2017).) Age of menarche in women with BD appears to indicate the course of the disorder, with being strongly correlated with irritable temperament scores and inversely correlated with depressive and cyclothymic scores as well as duration of depressive episodes (Kesebir et al., 2013). Age at menarche has also been negatively associated with number of depressive episodes (Dunjic-Kostic et al., 2016)

Looking to the reproductive cycle in BD more broadly, the impact of menstruation on mood in BD is instructive. A systematic review by Teatero et al. (2014) found that 64 to 68 percent of women with BD in retrospective studies reported mood changes with menses, as well as considerable diagnostic overlap with (DSM-4) premenstrual syndrome and premenstrual dysphoric disorder. In a study by Perich et al. (2017), 77 percent of women with BD reported increased mood symptoms during peri/menstrual and postnatal periods which—relative to women who did not report reproductive cycle-related changes in symptoms—was correlated with more severe BD course, including earlier age of onset with mood episodes and increased likelihood of rapid cycling and mixed mood states. Menorrhagia, polymenorrhea, oligomenorrhea, and amenorrhea are among the many reproductive cycle abnormalities reported among women with BD (Rasgon et al., 2003, 2005b; Williams et al., 2007; Barron et al., 2008). Indeed, women with BD experience menstrual dysfunction (irregularity or abnormally increased/decreased length of menstrual cycles) prior to the onset of psychiatric illness at a significantly higher rate relative to women with unipolar depression and healthy controls (Joffe et al., 2006), and women with unipolar depression were not significantly likelier to experience such dysfunction than healthy controls. This serves to highlight the centrality of mania in BD and the potentially explanatory role of oxytocin in the disorder, as discussed by Crespi’s (2016) examination of the ‘symmetrical’ comorbidities of ASD and polycystic ovarian syndrome (PCOS) on the one hand and BD and endometriosis on the other, proposed to be the results of decreased and increased oxytocin levels, respectively (though PCOS comorbidity with BD has also been reported by Qadri et al., 2018). Menstrual psychosis is also thought to be a manifestation of BD pathology (Abe and Ohta, 1995; Brockington, 2005, 2011), and menses can trigger exacerbation of psychosis in BD and other psychotic disorders (Reilly et al., 2019).

In line with this and with the reduction of androgen levels seen in domestication, BD is associated with decreased testosterone levels among untreated first-episode patients (Feng et al., 2019) and remitted male patients (Keshri et al., 2018). Interestingly, in BD patients in a current depressive episode, testosterone levels were significantly decreased among male patients compared to healthy male controls and significantly increased among female patients compared to healthy female controls (Wooderson et al., 2015), consistent with findings of hyperandrogenemia in women with BD (Rasgon et al., 2005a; Zerouni et al., 2013). In both males and females with BD (and controlling for sex), testosterone levels positively correlated with number of manic episodes as well as number of suicide attempts (with no correlation found between testosterone levels and aggression) (Sher et al., 2012). Among females with BD, a positive correlation was found between testosterone levels and number of past major depressive episodes and number of suicide attempts (Sher et al., 2014). At follow-up, elevated baseline testosterone levels predicted suicide attempts such that a 10 ng/dl increase in testosterone translated to a 16.9-times increased probability of suicide attempt. While it should be noted that one study found no significant differences in salivary testosterone levels either between BD-I patients and healthy controls or between euthymic, depressed, and manic BD patients (Mousavizadegan and Maroufi, 2018), there are reports of hypomania (Freinhar and Alvarez, 1985) and mania (Elboga and Sayiner, 2018) induced by androgen administration, as well as homicide after a man with BD was administered testosterone for hypogonadism (Sher and Landers, 2014). In a case with comorbid Klinefelter syndrome, however, testosterone therapy has been reported to effectively treat a pattern of frequent manic episodes refractory to treatment with mood stabilizers and second-generation antipsychotics (Kawahara et al., 2015). (Of note, testosterone therapy in early childhood has been shown to improve neurodevelopmental outcomes including language and intellectual ability in Klinefelter syndrome (47, XXY) patients, compared to untreated controls (Samango‐Sprouse et al., 2013).)

## 3. Genetic signatures of domestication and the genetics of BD

As noted in the introduction, BD has a strong genetic background. With the aim of testing whether our hypothesis is supported by available molecular evidence, we assessed whether genes that have been related to BD are overrepresented among candidates for domestication and neural crest (NC) development and function. To achieve this, we compiled an extended, updated list of BD-associated genes from two different curated databases: the Genetic Database for Bipolar Disorder (BDgene) (http://bdgene.psych.ac.cn/) and the Psychiatric disorders Gene association NETwork (PsyGeNet) (www.psygenet.org/). The BDgene database contains candidates resulting from different methodological approaches: genome-wide association analyses (GWAs), candidate-gene association studies, linkage studies, mutational studies, copy number variation (CNV) analyses, and meta-analyses for association or linkage studies. Candidates result from literature review via PubMed followed by manual curation. Candidate genes may bear pathogenic SNPs, be associated to pathological haplotypes or CNVs, be found mutated in familial forms of the disease, result from candidate gene approaches and functional studies, result from linkage studies, association studies, or GWAs, be part of pathways associated to the disease, or interact with known candidates for BD (see http://bdgene.psych.ac.cn/data.do for details). Of the 1192 genes discussed in the BD database, we considered only 599 genes categorized as positive, for which at least one significant study has been found under the significance criteria used by the database’s authors (see http://bdgene.psych.ac.cn/data.do#summary, table 3). Regarding the PsyGeNet database, it has been developed by automatic extraction of information from PubMed abstracts using the text mining tool BeFree (http://ibi.imim.es/befree/), followed by curation by experts in the domain (see Gutiérrez-Sacristán et al., 2015 for details). The number of genes associated to BD and related disorders in the PsyGeNet database is 514. However, we considered only the genes for which the evidence index was 1 (n = 374), implying that all of the evidence reviewed by the experts supports the existence of an association between the gene and the disease (for each gene, experts validate a maximum of five publications, selected among the most recent ones). The entire list of candidate genes for BD considered in our study resulted from merging these two lists (599 plus 374 genes) and removing the overlapping genes (see Supplemental file 1).

Regarding candidates for domestication, we compiled a comprehensive, updated list of candidates by merging the list compiled by Benítez-Burraco and collaborators (2017) in their paper on domestication features in schizophrenia with the list compiled by Theofanopoulou and collaborators (2017) in their work on signs of positive selection of candidates for domestication in humans. The merged list encompasses 764 genes (see Supplemental file 1). Because of the purported involvement, as noted, of the NC in the domestication syndrome (Wilkins et al., 2014), we have also considered candidate genes for NC development and function in our study. For this, we have used the list compiled by Benítez-Burraco and collaegues (2017) in their work on domestication features in schizophrenia, which comprises 89 genes gathered following pathogenic and functional criteria: neurocristopathy-associated genes annotated in the OMIM database (http://omim.org/), NC markers, genes that are functionally involved in NC induction and specification, genes involved in NC signaling (within NC-derived structures), and genes involved in cranial NC differentiation (see Supplemental file 1).

We then conducted a Fisher’s exact test to determine the significance of the overlapping between these lists, using protein-coding genes obtained from the Ensembl BioMart (GRCh38.p12) as background. We found a significant overlap (*p* = 3.37e-07) between candidates for BD and candidates for domestication. By contrast, the overlapping between candidates for BD and candidates for NC development and function was not significant (*p* = 0.168). Figure 2 displays the list of 62 common candidates for BD and domestication.

**Figure 2.**
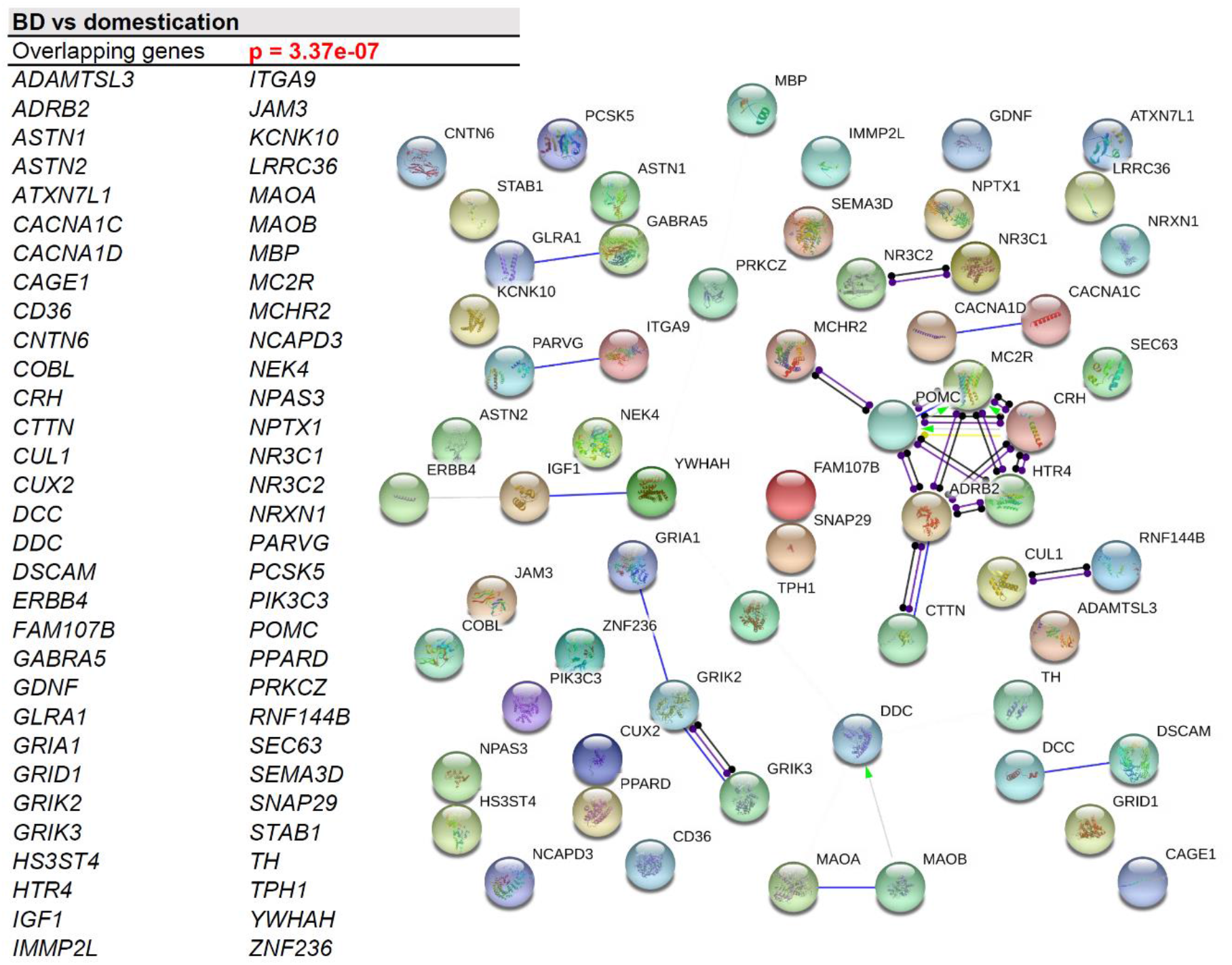
Candidates for BD that are also candidates for animal domestication. A. List of genes that are cited as candidate for BD and that show signals of positive selection in domesticated animals. B. Functional interactions among the genes that are candidates for BD and for domestication. The diagram shows the network of known functional interactions among the proteins encoded by the genes. The networks were drawn with String (version 11.0; Szklarczyk et al. 2015) license-free software (http://string-db.org/), using the molecular action visualization. Colored nodes symbolize the proteins. The color of the edges represents different kinds of known protein-protein associations. Green: activation, red: inhibition, dark blue: binding, light blue: phenotype, dark purple: catalysis, light purple: post-translational modification, black: reaction, yellow: transcriptional regulation. Edges ending in an arrow symbolize positive effects, edges ending in a bar symbolize negative effects, whereas edges ending in a circle symbolize unspecified effects. The medium confidence value was .0400 (a 40% probability that a predicted link exists between two enzymes in the same metabolic map in the KEGG database: http://www.genome.jp/kegg/pathway.html). The diagram only represents the attested connectivity between the involved proteins, derived from curated databases or experimentally determined, but it must be mapped onto particular biochemical networks, signaling pathways, cellular properties, aspects of neuronal function, or cell-types of interest to gain a more accurate view of its relevance for the presentation of domesticated features in BD (see the text and the Supplemental file 2 for further details).

### Functional characterization of genes of interest

#### GO analyses

We expected that the 62 genes we highlight here as part of the shared signature of domestication and BD play roles at the molecular and cellular levels, and/or map to biological processes, regulatory pathways, cell types, or aspects of brain development and function, of interest not only for BD etiopathogenesis, but also for human cognitive evolution. For this reason, we conducted gene ontology analyses (GO) of the set of common candidates for BD and domestication. Functional enrichment analyses were performed with Enrichr (amp.pharm.mssm.edu/Enrichr; Chen et al., 2013; Kuleshov et al., 2016). We considered biological processes, molecular functions, cellular components, or human pathological phenotypes with associated *p*-values < 0.05 as enriched. We found that these genes are significantly related to processes, functions, cellular components, and pathological phenotypes of interest for the BD phenotype (Figure 3). Specifically, we found that this set of genes is significantly enriched in aspects of neurotransmitter homeostasis and function, mostly dopaminergic (GO:0042420; GO:0042417; GO:0001963) and glutamatergic transmission (GO:0035249; GO:0008066; GO:0004970; GO:0008328), but also indolamine (i.e. serotonin) (GO:0006586) and catecholamine transmission (GO:0042424), and mostly impacting on postsynaptic potential (GO:2000463; GO:0099529; GO:1904315). These genes are also predicted to be significantly involved in dendrite development (GO:2000171) and axonal growth (GO:0044295). Not surprisingly, these genes are enriched in proteins related to ion channel activity (GO:0022824, GO:0008331, GO:0005237, GO:1990454, GO:0034706, GO:0034705). Also of interest is their association to insulin receptor signaling pathway (GO:0046628), insulin-like growth factor receptor binding (GO:0005159), and low-density lipoprotein receptor activity (GO:0005041).

**Figure 3.**
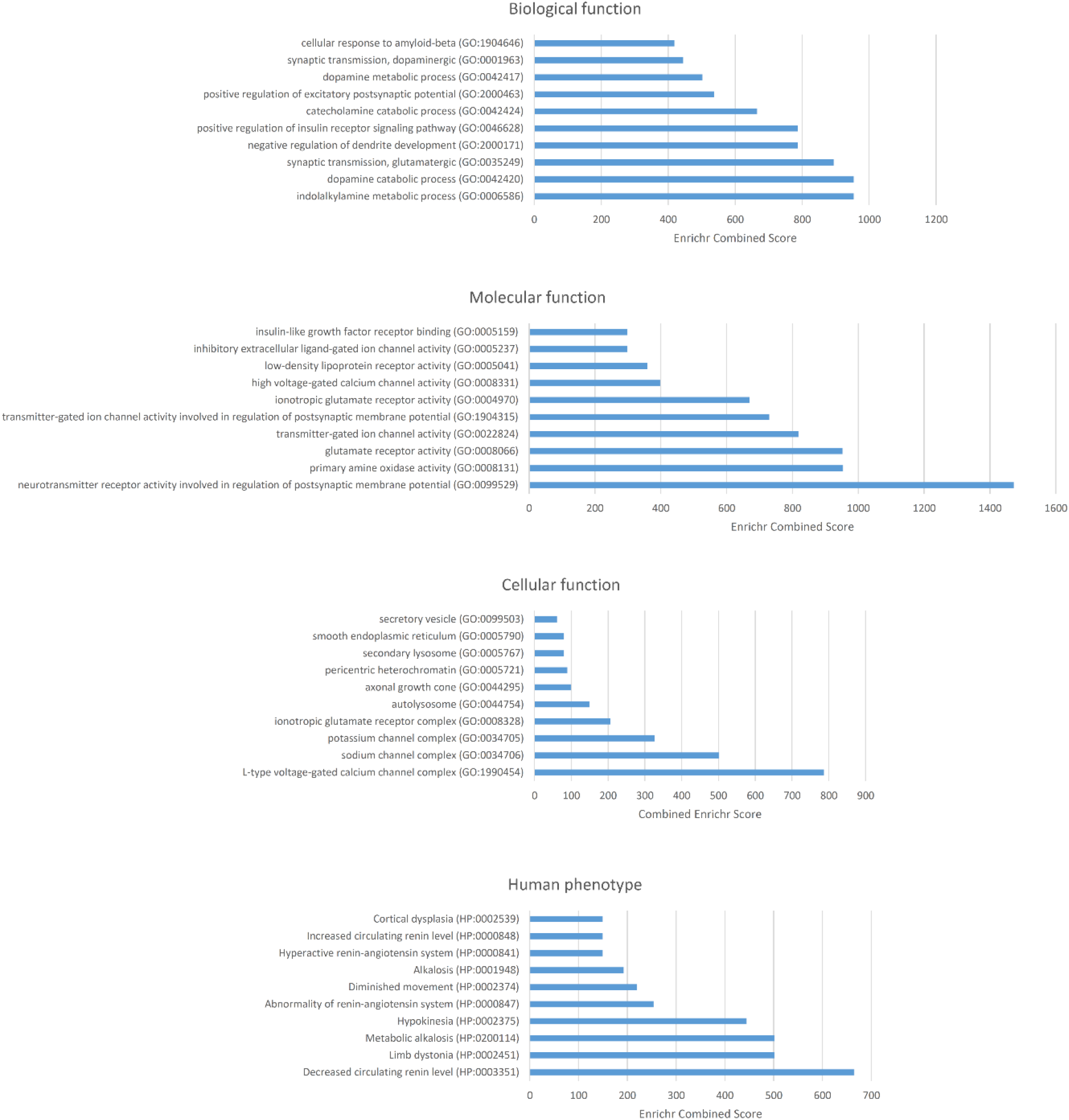
Functional enrichment analysis according to Enrichr of the set of genes that are candidates for BD and for domestication. The graphs show the enrichment in biological processes, molecular function, cellular components, and human pathological phenotypes (from top to bottom). Only the top 10 functions have been included, and only then if their *p* < 0.05. The *p*-value was computed using Fisher’s exact test. Enriched categories are ordered according to their Enrichr Combined Scores. This is a combination of the *p*-value and the *z*-score calculated by multiplying the two scores (Combined Score = ln(*p*-value) * *z*-score). The *z*-score is computed using a modification of Fisher’s exact test and assesses the deviation from the expected rank. The Combined Score provides a compromise between both methods and is claimed to report the best rankings when compared with the other scoring schemes. See http://amp.pharm.mssm.edu/Enrichr/help#background&q=5 for details.

Alterations in dopamine homeostasis have been claimed to contribute significantly to the BD phenotype, with high levels of dopamine underlying the manic phase of the disease and reduced dopamine levels accounting for depressive symptoms (Berk et al., 2007). Recent neurobiological and pharmacological findings (reviewed by Ashok et al., 2017) support the view that increased dopamine receptor availability underlies mania, whereas increased dopamine transporter levels may account for depression. Evolutionary changes in dopaminergic innervation have been proposed to account for human-specific cognitive abilities, like language. Specifically, changes in dopaminergic afferents of the striatum have been argued to underlie aspects of speech and language evolution (Raghanti et al., 2016). Interestingly, dopamine is involved in social behavior in animal species, with the level of dopaminergic system activity influencing social support between individual via diverse cognitive functions (Lin et al., 2011). According to Yamaguchi and colleagues (2015), evolutionary changes of dopamine signaling and associated genetic variants might yield advantages at the group level even if they result in disadvantages at the individual level. Domesticated foxes exhibit increased dopamine levels in the left half of the striatum compared to aggressive foxes, which may be related to reduced function of the pituitary-adrenal system (Trut et al., 2000). Interestingly, changes in dopaminergic and other neuromodulatory systems have been suggested to be related to the emergence of human-distinctive traits via self-domestication (Calvey, 2019). Likewise, glutamatergic neurotransmission is involved in excitatory activity of the brain, and patients with BD exhibit abnormally increased glutamate levels in the frontal areas, compared to neurotypical controls, as well as evidence of abnormal glutamatergic neurotransmission (Hashimoto et al. 2007; Eastwood and Harrison, 2010; Gigante et al., 2012; Gottschalk et al., 2015). Genes involved in glutamate metabolism have been subject to strong selection in many domesticated strains of mammals, including dog, pig, and fox, seemingly contributing to their increased tameness and distinctive social cognitive capabilities (Li et al., 2014; Moon et al., 2015; Wang et al., 2018; O’Rourke and Boeckx, 2020). Interestingly, modern humans also show changes in glutamate (most notably, kainate and metabotropic) receptor genes compared to extinct hominins, reinforcing the view that human self-domestication resulted in changes in hypothalamic-pituitary-adrenal axis excitation—and ultimately, the attenuation of the stress response—via changes in glutamatergic innervation (O’Rourke and Boeckx, 2020).

Enrichment in genes related to the insulin-like growth factor 1 is also of interest, given the diverse physiological activities played by this factor, from growth hormone regulation to heart development, genitourinary development, and musculoskeletal development (see Puche and Castilla-Cortázar, 2012 for review). Within the brain, the insulin-like growth factor 1 contributes to brain development and plasticity as well as myelinization and synapse formation (see Puche and Castilla-Cortázar, 2012 for review). This circumstance might account for some of the developmental features found in people with BD, including altered growth and reproductive cycle, as reviewed in the previous section. Abnormally high levels of insulin-like growth factor 1 have been found in the blood of patients with BD (Da Silva et al. 2017). It has been suggested that these increased levels correlate with manic episodes and might constitute a compensatory mechanism against excitotoxicity (Ferensztajn-Rochowiak et al., 2019). Finally, the enrichment in genes related to lipoproteins is also of interest in view of claims that mood disorders and cardiometabolic disease might share biological mechanisms, including candidate genes (Amare et al., 2017). Specifically, patients with BD suffer higher mortality due to circulatory-related problems such as heart attacks (Hayes et al., 2015). Pharmacotherapy for BD has been claimed to potentially increase susceptibility for type 2 diabetes mellitus and dyslipidemia, which are known cardiovascular risk factors (Correll et al., 2015), and insulin resistance is more prevalent relative to controls even among psychotropic-naïve BD patients (Guha et al., 2014). As noted, our set of genes is also enriched in genes involved in insulin signaling.

Finally, genes that are candidates for both domestication and BD are significantly associated to human pathological phenotypes impacting the renin-angiotensin system (HP:0003351, HP:0000847, HP:0000841, HP:0000848), which is mostly involved in the regulation of blood pressure and electrolyte balance, as well as vascular resistance, but which has been also linked to mood disorders (Mohite et al., 2019). Patients with BD show higher plasma renin activity compared to controls (Barbosa et al., 2020). In animals, the inhibition of selected components of the renin-angiotensin system reverses the hypertension and cognitive deficits associated with mood disorders (Mohite et al. 2019). This circumstance is seemingly related to the cardiovascular problems observed in people with BD noted above. Interestingly, our set of genes are also significantly associated to motor problems, including diminished movement (HP:0002374), dystonia (HP:0002451), and hypokinesia (HP:0002375). Patients with BD have been shown to exhibit impaired motor sequencing (Negash et al., 2004), as well as impaired fine motor skills associated to depressive phases (Malhi et al., 2007). Abnormal rhythm is more prevalent in mood disorders than in schizophrenia (Boks et al. 2000). Psychomotor disturbances commonly found in major depressive disorders can result not only in reduced mobility, but also in in decreased speech production. Interestingly, this can result from reduced coupling between cortical and striatal regions (Liberg et al., 2014). Finally, it is also of interest that these genes are significantly related to cortical dysplasia (HP:0002539) considering that BD entails decreases in anterior cortical volumes over time, including gray matter contraction and reduced white matter expansion (Najt et al., 2016), and that epilepsy (a common outcome of cortical dysplasia) and BD frequently co-occur and share a similar pathophysiology (Knott et al., 2015).

A detailed functional characterization of these 62 common candidates for BD and domestication is provided in Supplemental file 2, including connections with the etiopathogenesis of BD and related conditions.

#### Network analyses

Additionally, we expected that some of these shared genes between the BD and domestication signatures are functionally interconnected. In order to check these possibilities, we used String 11.0 (www.string-db.org), a license-free software that predicts direct/physical and indirect/functional associations between proteins derived from several sources (high-throughput experiments, conserved coexpression, genomic context, and knowledge previously gained from text mining) (Szklarczyk et al. 2015). As shown in Figure 2, String 11.0 points to quite robust functional links among several of these genes, particularly, *ADRB2*, *CRH*, *CTTN*, *HTR4*, *MC2R*, and *MCHR2*, which are mostly involved in the interactions between the noradrenergic system and the hormonal system responsible for the response to stress, which, as discussed in sections 1 and 2 above, are altered in both BD and domestication. The core component of this network is *ADRB2*, which encodes the beta-2-adrenergic receptor, which has a high binding affinity for epinephrine, but which also activates the receptors of the corticotropin releasing hormone (CRH) (Vranjkovic et al., 2014), important for stress response and adrenocorticotropic hormone (ACTH) release and function (Sapolsky et al., 2000; Keller-Wood, 2015). One of the proteins involved in the recycling of the beta-2-adrenergic receptor is cortactin, a cell-adhesion protein encoded by *CTTN* (Vistein and Puthenveedu, 2014). CRH is secreted by the paraventricular nucleus of the hypothalamus in response to stress and stimulates the production and secretion of ACTH by the anterior pituitary, which then acts on the adrenal cortex to stimulate the synthesis and release of glucocorticoids, which are crucial for a normal response to stressful perturbations (Sapolsky et al., 2000; Keller-Wood, 2015). Interestingly, glutamate receptors, which are subject to selection in domesticates, mediate CRH release and thereby modulate the HPA axis response to stress (see O’Rourke and Boeckx, 2020 for details). Glucocorticoid receptors have a negative feedback effect on both CRH neurons in the hypothalamus and *CRH* expression in the hypothalamus, but also downregulate *POMC* in the pituitary (Keller-Wood, 2015). *POMC* contributes to the homeostasis of up to ten biologically active peptides, including ACTH, melanocyte-stimulating hormone (MSH), and the opioid-receptor ligand beta-endorphin (Smith and Funder, 1988). As noted in the previous section, subjects with BD exhibit increased levels of cortisol and ACTH, but not of CRH, with cortisol levels positively correlating with the manic phase (Belvederi Murri et al. 2015). Interestingly, variants of most of the genes related to the HPA axis seem to be associated not with a direct risk of suffering BD, but with different clinical presentations. Specifically, two *CRH* polymorphisms have been associated with the severity of psychotic symptoms in BD (Leszczynska-Rodziewicz et al., 2012; Leszczynska-Rodziewicz et al., 2013). Likewise, although first genetic analyses did not find molecular variation in *POMC* in bipolar illness (e.g. Feder et al., 1985), increased levels of pro-opiomelanocortin have been found in BD patients (Stelzhammer et al., 2015), and subtle alterations in pro-opiomelanocortin processing have been suggested to occur in BD (Berrettini et al., 1985). In mammals both *POMC* and *CRH* are upregulated after stress in the hypothalamus (Lightman and Young, 1988, Givalois et al., 2000, Lee et al., 2005). Two other important components of the network are *MC2R* and *HTR4*, encoding, respectively, one of the receptors for ACTH (Detera-Wadleigh et al., 1995; Bickeböller et al., 1997) and a serotonin receptor (Ohtsuki et al., 2002). See Supplemental file 2 for a detailed characterization of these genes.

The remaining candidate genes appear isolated and are not suggested to be functionally interconnected with the genes we have previously highlighted under the stringent conditions we have employed (we have only relied on curated databases of molecular interactions and on experimentally determined interactions between proteins, and used a confidence value of 0.04, indicating a 40 percent probability that a predicted link exists between two proteins in the same map in the KEGG database: http://www.genome.jp/kegg/pathway.html). Nonetheless, most of these genes play relevant roles in cognitive functions affected in patients with BD, as we show in Supplemental file 2.

### Gene expression patterns

Finally, we investigated the expression profiles of the genes that are candidates for both BD and domestication. A heat map of the expression levels of these genes in the samples of the Human tissue compendium (Novartis) (Su et al., 2004), as generated by Gene Set Enrichment Analysis (GSEA) software (http://software.broadinstitute.org/gsea/index.jsp), is shown in Figure 4. According to the Human Gene Atlas (Su et al., 2004), genes that are candidates for BD and domestication are predicted to be preferentially expressed in several areas of the brain, particularly, the amygdala (*p* = 0.0006878; Enrichr combined score = 40.85), parietal lobe (*p* = 0.07760; Enrichr combined score = 31.71), and prefrontal cortex (*p* = 0.02257; Enrichr combined score = 11.74). Outside the brain, they are preferentially expressed in body regions involved in domestication, particularly, the adrenal cortex (*p* = 0.04283; Enrichr combined score = 19.18). As noted in section 2 above, the amygdala is found enlarged in subjects with BD, as well as in domesticated strains of animals. Regarding the temporal lobe, meta-analyses of the available data suggest that BD entails significant dysfunction of the temporal lobe impacting executive function, verbal memory, and emotion regulation (Whalley et al., 2012; Bostock et al., 2017), as well as structural anomalies in this region, mostly decreased cortical thickness and reduced grey matter (Selvaraj et al., 2012; Hanford et al., 2016). Likewise, abnormal activation patterns in neural networks including the prefrontal cortex have been observed in people with BD compared to other mood and cognitive disorders, like unipolar depression (Han et al., 2019). In a similar vein, morphometric studies have found volume reduction as well as decreased grey matter in the prefrontal cortex (Selvaraj et al., 2012; Roda et al., 2015; Hanford et al., 2016).

**Figure 4.**
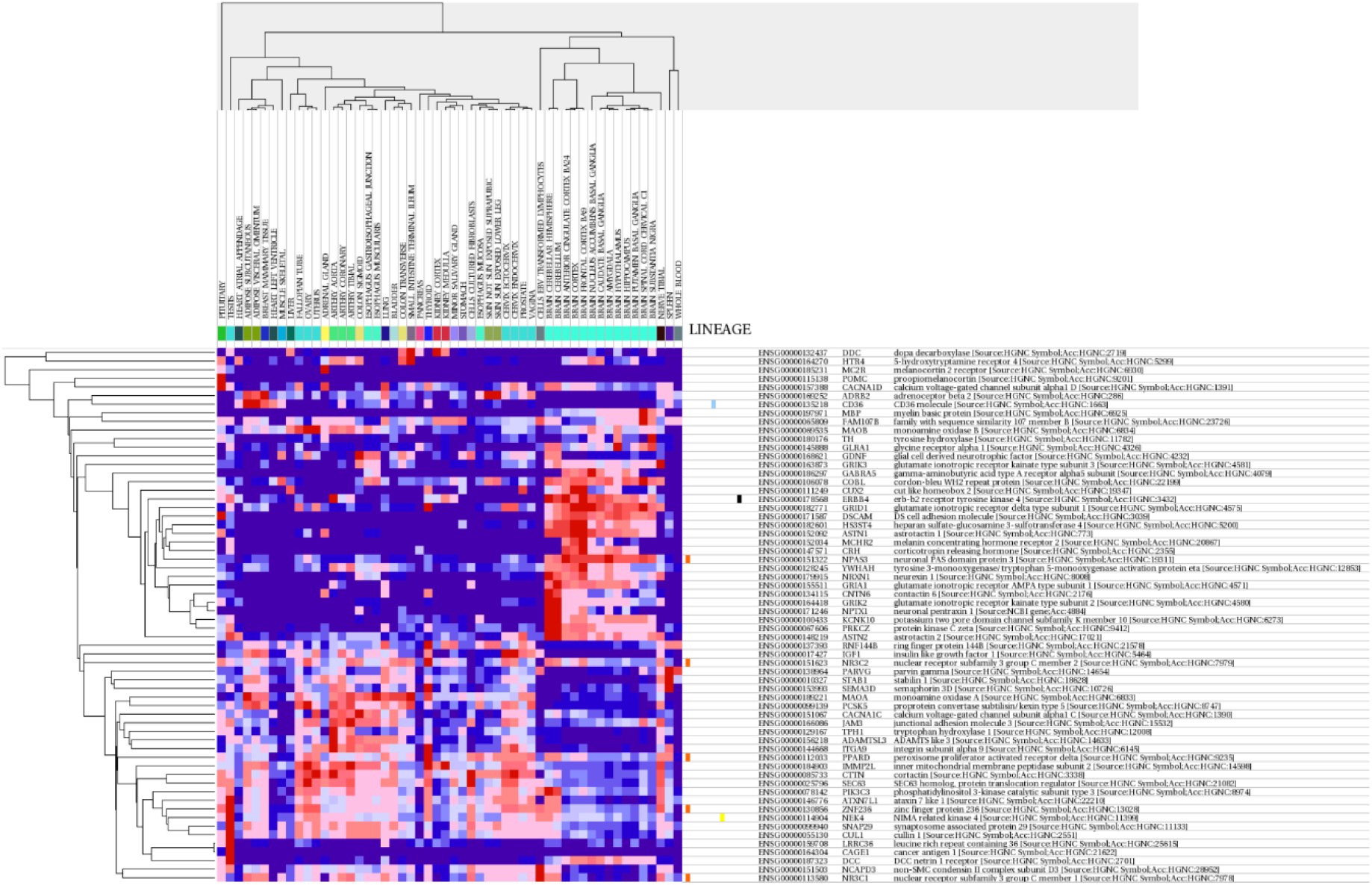
Heat maps of the expression levels of the set of genes that are candidates for both BD and domestication. The heat map was generated by the Gene Set Enrichment Analysis (GSEA) software using the samples of the Human tissue compendium (Novartis) (Su et al., 2004). GSEA is a computational method that determines whether an a priori defined set of genes shows statistically significant, concordant differences between two biological states. The heat maps include dendrograms clustering gene expression by gene and samples. Genes are identified by probe identifier, gene symbol, description, and gene family. See http://software.broadinstitute.org/gsea/index.jsp for details.

Given that BD is primarily a cognitive condition and that our set of genes is predicted to be significantly expressed in brain areas of interest for BD pathogenesis, we interrogated whether candidates for domestication are differentially expressed in vivo in selected areas of the brain of patients. For this, we relied on available datasets of differentially expressed genes (DEGs) as found in the GEO repository (https://www.ncbi.nlm.nih.gov/gds). Specifically, we made use of recent data produced by Lanz and colleagues (2019) (GSE53987; supplemental table 5), who analyzed DEGs in postmortem samples of the prefrontal cortex (PFC), the hippocampus (HIP), and the striatum (STR) of people with BD. We considered only DEGs with expression *p*-values < 0.01 (see Supplemental file 3). We then conducted a Fisher’s Exact Test (95 percent confidence interval), to check whether these three sets of genes overlap significantly with candidates for domestication. To calculate this, we considered (i) the number of overlapping genes, (ii) the number of DEGs in the brain of patients, (iii) the number of domestication genes, and (iv) the total number of genes present in the microarray (Supplemental file 3). We found that candidates for domestication are significantly dysregulated in the PFC (*p* = 0.00249) and HIP (*p* = 0.001806), but not in the STR (*p* = 0.2729); these brain regions are not only involved in the pathophysiology of BD, as noted, but are also affected by domestication processes. Figure 5 shows the domestication candidates that are significantly downregulated and upregulated in the prefrontal cortex and hippocampus of patients.

**Figure 5.**
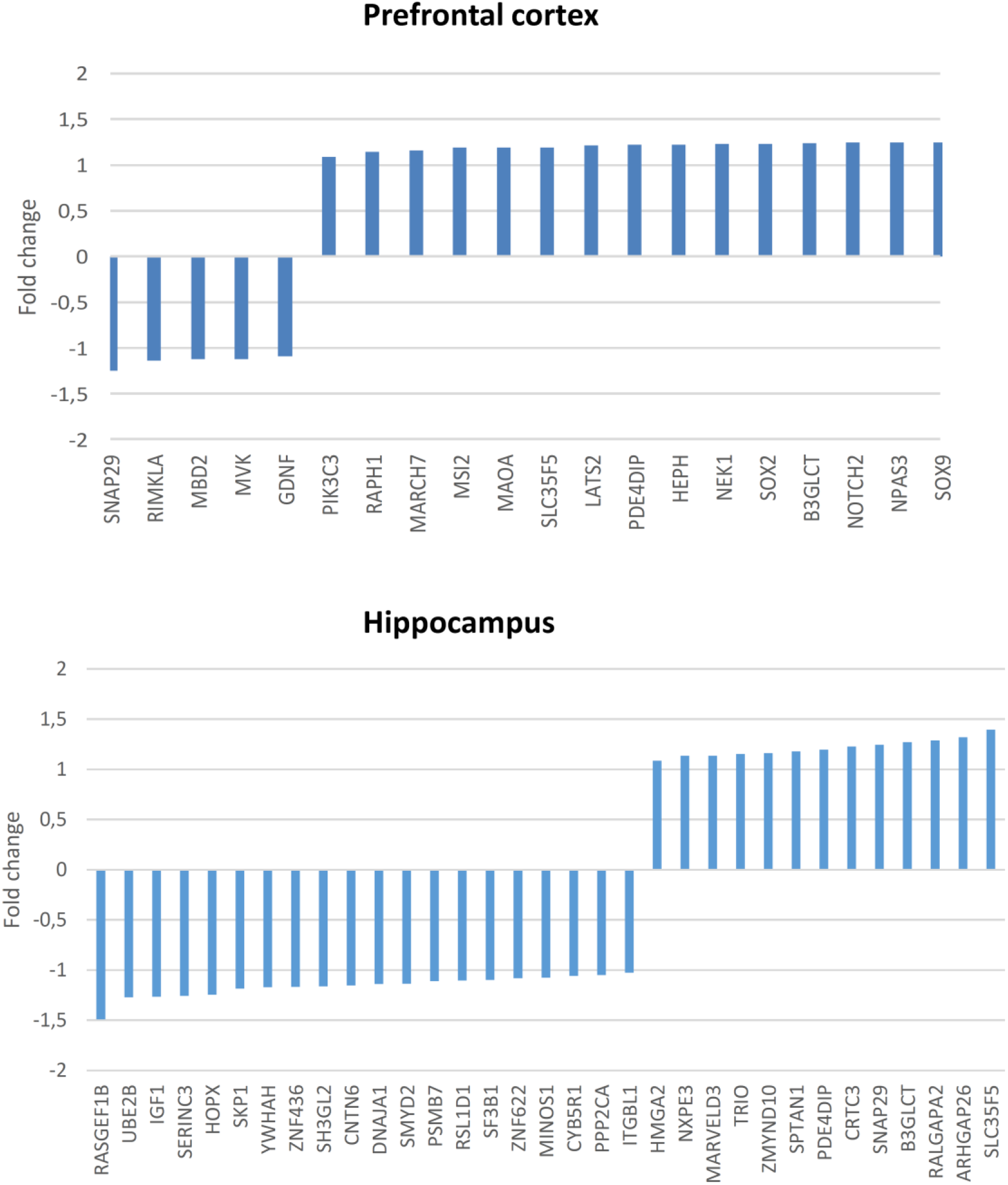
Changes in the expression levels of candidates for domestication in the prefrontal cortex (top) and the hippocampus (bottom) of subjects with BD compared to neurotypical controls. Only DEGs with expression *p*-values < 0.01 have been considered (see Supplemental files 3 and 4 for details).

We conducted GO analyses of these two set of genes with Enrichr. As previously, we considered biological processes, molecular functions, cellular components, or human pathological phenotypes as enriched if their *p* < 0.05. We found that candidates for domestication that are differentially expressed in the prefrontal cortex of people with BD are significantly involved in neurodevelopment (GO:0048484) and the regulation of catecholamine secretion (GO:0033605), as well as in the regulation of gene expression, via satellite DNA binding (GO:0003696), enhancer sequence-specific DNA binding (GO:0001158), methyl-CpG binding (GO:0008327), and miRNA binding (GO:0035198). Likewise, they are significantly associated to human pathological phenotypes impacting brain development and function, particularly, cortical dysplasia (HP:0002539) (see Supplemental file 4 for further details). Regarding candidates for domestication that are differentially expressed in the hippocampus of subjects with BD, they are significantly involved in gene expression regulation (including histone H2A ubiquitination (GO:0033522), DNA binding (GO:2000679), histone-serine phosphorylation (GO:0035404), histone methyltransferase activity (GO:0046975), and AT DNA binding (GO:0003680)), as well as dendrite extension (GO:0097484), and regulation of the activity of the insulin-like growth factor (GO:0005159) and glucocorticoids (GO:0035259). Similar to genes that are differentially expressed in the prefrontal cortex, these genes are significantly associated to human pathological phenotypes impacting on brain development and function, like progressive microcephaly (HP:0000253) and cortical dysplasia (HP:0002539) (see Supplemental file 4 for further details).

## Discussion and conclusions

Overall, our findings suggest that the complex phenotype exhibited by people with BD can be construed as a combination of hypo- and hyperdomesticated features. This is in line with the mixed nature of this disorder, entailing both schizophrenia-like features (with SZ being in turn a hyperdomesticated human phenotype; see Benítez-Burraco et al., 2017 for discussion) and ASD-like features (with ASD being in turn a hypodomesticated human phenotype; see Benítez-Burraco et al., 2016 for discussion). This is also in line with previous accounts of BD as an intermediate phenotype within the continuum of human social cognition (Crespi and Badcock, 2008). Such a view emphasizes the role of genomic imprinting in the etiology of SZ and ASD—with relative biases toward maternal expression in SZ and paternal expression in ASD—and interprets features in SZ and ASD in light of the interaction of sex differences and imprinting effects. Like SZ, BD is associated with DNA methylation changes, including of glutamate receptor and neurodevelopment genes (Mill et al., 2008). Accordingly, future work could investigate sex differences in BD—especially those relating to domestication-relevant characteristics, as described in section 2—in connection with imprinting effects. At present, available data are very scarce. Still, we found 3 of our common candidates for domestication and BD (*DDC*, *GABRA5*, and *TH*) in the Geneimprint database (http://www.geneimprint.com/), with at least one of them (*GABRA5*) showing robust evidence of a paternal expression effect (data not shown).

Further study of the role of imprinting effects, sex differences, and their interaction may also illuminate a conundrum that unfolds in the application of the self-domestication hypothesis to human psychiatric/neurodevelopmental disorders: why is it that human disorders able to be construed as hyperdomesticated phenotypes are characterized by hypersociability (as in the case of WS; see Niego and Benítez-Burraco, 2019) or hypermentalization (as in the case of SZ; see Benítez-Burraco et al., 2017), but rarely both? Psychosis, for example, is rarely seen in WS (though has been reported in a female patient by Salgado and Martins-Correia, 2014), and SZ is characterized by persistent and severe social deficits despite (or perhaps in part because of) hypermentalization (Savla et al, 2013). BD, especially considering the manic phase, might have been thought to be a hyperdomesticated state with both hypersociable and hypermentalizing (e.g., psychotic) features, but as we have seen, the condition is characterized by deficits in social cognition and other hypo- and hyperdomesticated features.

A key pair of imprinting disorders that Crespi and Badcock (2008) cite in support of their hypothesis is Angelman syndrome—the paternally-biased condition characterized by sociability and ASD-like features—and Prader-Willi syndrome—the maternally-biased condition characterized by infantile undergrowth and, often, psychosis, commonly as part of atypical bipolar or schizoaffective disorder (Singh et al., 2019). Somewhat similarly, mutations in *MECP2* lead to Rett syndrome in females and are typically lethal in males, but surviving males exhibit psychosis, usually as part of atypical BD, as well as impaired language development and pyramidal/parkinsonian signs (termed PPMX syndrome, OMIM 300055) (Lambert et al., 2016).

Among neurodevelopmental disorders and neurocristopathies characterized by a happy/sociable demeanor and other traits that could suggest an altered (possibly hyper-)domestication phenotype— including Pitt-Hopkins syndrome (OMIM 610954), Mowat-Wilson syndrome (OMIM 235730), Cornelia de Lange syndrome 5 (OMIM 300882), Glass syndrome (OMIM 612313), Christianson syndrome (OMIM 300243), Skraban-Deardorff syndrome (OMIM 617616), Nablus mask-like facial syndrome (608156), NDAGSCW (OMIM 617807), NEDMAGA (OMIM 617865), and pontocerebellar hypoplasia type 11 (OMIM 617695)—none are characterized by psychosis. An important exception to this rule is Down syndrome (DS), commonly characterized by a happy, sociable demeanor as well as comorbidities—e.g., Hirschsprung disease—and other features possibly suggestive of an underlying neurocristopathy (see *DSCAM* in Supplemental file 2). DS has been found by Dykens and colleagues (2015) to be unique among intellectual disabilities for higher rates of psychosis (43 percent compared to 13 percent in a pooled group of those with other IDs), and 81 percent of DS patients with psychosis were female (*p* < .001, with no significant sex difference among those with psychosis in the pooled group). Further study of the interaction of sex differences and any imprinting effects, then, might be crucial to understanding the varied manifestations of hyperdomestication in humans.

Overall, because of the relevance of the self-domestication hypothesis for explaining human evolution, and because of the strong link that exists between evolutionary novelties and complex disease, the genes highlighted in this paper should be considered as potentially relevant etiological factors of the bipolar condition.

## Acknowledgements

The authors thank Dr. Dafni Anastasiadi, from the Department of Breeding and Genomics, Plant and Food Research, University of Auckland, for helping with the analysis of gene expression datasets.

## Statement of Ethics

The research conducted for the paper relied on previously published data by others and available datasets, hence no ethics approval was required

## Author Contributions

ABB and EH conceived the paper, performed the data analyses and wrote and approved the final manuscript

## Supplemental data

Supplemental file 1. List of candidates for BD and for domestication.

Supplemental file 2. Functional characterization of genes that are candidates for both domestication and BD.

Supplemental file 3. DEGs in the prefrontal cortex, the hippocampus and the striatum of patients with BD and overlapping with candidate genes for domestication. DEGs have been gathered from Lanz and colleagues’ 2019 supplemental table 5. Only DEGs with expression *p*-values < 0.01 have been considered. The list of candidates for domestication is the one included in Supplemental file 1. A Venn diagram (right, top) shows in each case the relation between the set of DEGs and the set of candidates for domestication. The diagram was drawn with the webtool designed by the bioinformatics evolutionary genomics group at the University of Gent, Belgium (http://bioinformatics.psb.ugent.be/webtools/Venn/). The list of overlapping genes is displayed below, together with the data used for the Fisher’s exact test (at bottom). DEGs in the prefrontal cortex are highlighted in green, DEGs in the hippocampus are highlighted in blue, and DEGs in the striatum are highlighted in brown.

Supplemental file 4. Functional enrichment analysis according to Enrichr of the set of genes that are differentially expressed in the prefrontal cortex (left) and the hippocampus (right) of patients with BD and that are also candidates for BD. The graphs show the enrichment in biological processes, molecular function, cellular components, and human pathological phenotypes (from top to bottom). Only the top 10 functions have been included and only if their *p* < 0.05. The *p*-value was computed using Fisher’s exact test. Enriched categories are ordered according to their Enrichr Combined Scores. This is a combination of the *p*-value and the *z*-score calculated by multiplying the two scores (Combined Score = ln(*p*-value) * *z*-score). The *z*-score is computed using a modification of Fisher’s exact test and assesses the deviation from the expected rank. The Combined Score provides a compromise between both methods and is claimed to report the best rankings when compared with the other scoring schemes. See http://amp.pharm.mssm.edu/Enrichr/help#background&q=5 for details.

## References

1. Abe, K., and Ohta, M. (1995). Recurrent brief episodes with psychotic features in adolescence: periodic psychosis of puberty revisited. Br J Psychiatry 167, 507–513. doi:10.1192/bjp.167.4.507.

2. Akabaliev, V. H., Sivkov, S. T., and Mantarkov, M. Y. (2014). Minor physical anomalies in schizophrenia and bipolar I disorder and the neurodevelopmental continuum of psychosis. Bipolar Disorders 16, 633–641. doi:10.1111/bdi.12211.

3. Akabaliev, V., Sivkov, S., Mantarkov, M., and Ahmed-Popova, F. (2011). Minor physical anomalies in patients with bipolar I disorder and normal controls. Journal of Affective Disorders 135, 193–200. doi:10.1016/j.jad.2011.07.019.

4. Alsabban S, Rivera M, McGuffin P (2011) Genome-wide searches for bipolar disorder genes. Curr Psychiatry Rep. 13(6):522–527

5. Altshuler, L. L., Bartzokis, G., Grieder, T., Curran, J., and Mintz, J. (1998). Amygdala enlargement in bipolar disorder and hippocampal reduction in schizophrenia: an MRI study demonstrating neuroanatomic specificity. Arch. Gen. Psychiatry 55, 663–664.

6. Altshuler, L. L., Bartzokis, G., Grieder, T., Curran, J., Jimenez, T., Leight, K., et al. (2000). An MRI study of temporal lobe structures in men with bipolar disorder or schizophrenia. Biological Psychiatry 48, 147–162. doi:10.1016/S0006-3223(00)00836-2.

7. Amare AT, Schubert KO, Klingler-Hoffmann M, Cohen-Woods S, Baune BT (2017) The genetic overlap between mood disorders and cardiometabolic diseases: a systematic review of genome wide and candidate gene studies. Transl Psychiatry 7(1):e1007. doi: 10.1038/tp.2016.261.

8. Anderson IM, Haddad PM, Scott J (2012) Bipolar disorder. BMJ 345:e8508. doi: 10.1136/bmj.e8508.

9. Ashok AH, Marques TR, Jauhar S, Nour MM, Goodwin GM, Young AH, Howes OD (2017) The dopamine hypothesis of bipolar affective disorder: the state of the art and implications for treatment. Mol Psychiatry 22(5):666–679. doi: 10.1038/mp.2017.16.

10. Barbosa IG, Ferreira GC, Andrade Júnior DF, Januário CR, Belisário AR, Bauer ME, Simões E Silva AC (2020) The renin angiotensin system and bipolar disorder: a systematic review. Protein Pept Lett. doi: 10.2174/0929866527666200127115059

11. Barrera, Á., Vázquez, G., Tannenhaus, L., Lolich, M., and Herbst, L. (2013). Theory of mind and functionality in bipolar patients with symptomatic remission. Revista de Psiquiatría y Salud Mental (English Edition) 6, 67–74. doi:10.1016/j.rpsmen.2012.07.003.

12. Barron, M. L., Flick, L. H., Cook, C. A., Homan, S. M., and Campbell, C. (2008). Associations between Psychiatric Disorders and Menstrual Cycle Characteristics. Arch Psychiatr Nurs 22, 254–265. doi:10.1016/j.apnu.2007.11.001.

13. Beetz A, Uvnäs-Moberg K, Julius H, Kotrschal K (2012) Psychosocial and psychophysiological effects of human-animal interactions: the possible role of oxytocin. Front. Psychol. 3:234. doi:10.3389/fpsyg.2012.00234.

14. Belvederi Murri M, Prestia D, Mondelli V, Pariante C, Patti S, Olivieri B, Arzani C, Masotti M, Respino M, Antonioli M, Vassallo L, Serafini G, Perna G, Pompili M, Amore M. (2016) The HPA axis in bipolar disorder: Systematic review and meta-analysis. Psychoneuroendocrinology. 63:327–42. doi: 10.1016/j.psyneuen.2015.10.014.

15. Belvederi Murri, M., Prestia, D., Mondelli, V., Pariante, C., Patti, S., Olivieri, B., et al. (2016). The HPA axis in bipolar disorder: Systematic review and meta-analysis. Psychoneuroendocrinology 63, 327–342. doi:10.1016/j.psyneuen.2015.10.014.

16. Benítez-Burraco A (2020a). Genes dysregulated in the blood of people with Williams Syndrome are enriched in protein-coding genes positively selected in humans. Eur J Med Genet 30: 103828. doi: 10.1016/j.ejmg.2019.103828

17. Benítez-Burraco A (2020b) Genes positively selected in domesticated mammals are significantly dysregulated in the blood of people with Autism Spectrum Disorders. Mol Syndromol 10(6): 306–312 doi: 10.1159/000505116

18. Benítez-Burraco A, Di Pietro L, Barba M, Lattanzi W (2017) Schizophrenia and human self-domestication: an evolutionary linguistics approach. Brain Behav Evol 89(3): 162–184

19. Benítez-Burraco A, Lattanzi W, Murphy E (2016) Language impairments in ASD resulting from a failed domestication of the human brain. Front Neurosci 10: 373. doi: 10.3389/fnins.2016.00373.

20. Benítez-Burraco, A., and Kempe, V. (2018). The Emergence of Modern Languages: Has Human Self-Domestication Optimized Language Transmission? Front. Psychol. 9. doi:10.3389/fpsyg.2018.00551.

21. Benítez-Burraco, A., Pietro, L. D., Barba, M., and Lattanzi, W. (2017). Schizophrenia and Human Self-Domestication: An Evolutionary Linguistics Approach. BBE 89, 162–184. doi:10.1159/000468506.

22. Berecz, H., Csábi, G., Jeges, S., Herold, R., Simon, M., Halmai, T., et al. (2017). Minor physical anomalies in bipolar I and bipolar II disorders – Results with the Méhes Scale. Psychiatry Research 249, 120–124. doi:10.1016/j.psychres.2017.01.014.

23. Berrettini WH, Nurnberger JI Jr, Chan JS, Chrousos GP, Gaspar L, Gold PW, Seidah NG, Simmons-Alling S, Goldin LR, Chrétien M, Gershon ES (1985) Pro-opiomelanocortin-related peptides in cerebrospinal fluid: a study of manic-depressive disorder. Psychiatry Res. 16(4):287–302.

24. Bhambhvani HP, Simmons M, Haroutunian V, Meador-Woodruff JH (2016) Decreased expression of cortactin in the schizophrenia brain. Neuroreport 27(3):145–50. doi: 10.1097/WNR.0000000000000514.

25. Bhat S, Dao DT, Terrillion CE, Arad M, Smith RJ, Soldatov NM, Gould TD (2012) CACNA1C (Cav1.2) in the pathophysiology of psychiatric disease. Prog Neurobiol 99(1):1–14.

26. Bickeböller H, Kistler M, Scholz M (1997) Investigation of the candidate genes ACTHR and golf for bipolar illness by the transmission/disequilibrium test. Genet Epidemiol. 14(6):575–80.

27. Bisaga, K., Petkova, E., Cheng, J., Davies, M., Feldman, J. F., and Whitaker, A. H. (2002). Menstrual Functioning and Psychopathology in a County-Wide Population of High School Girls. Journal of the American Academy of Child & Adolescent Psychiatry 41, 1197–1204. doi:10.1097/00004583-200210000-00009.

28. Boks MP, Russo S, Knegtering R, Bosch van den RJ (2000) The specificity of neurological signs in schizophrenia: a review. Schizophr Res. 43: 109–116. 10.1016/S0920-9964(99)00145-0.

29. Bora, E., Vahip, S., Gonul, A. S., Akdeniz, F., Alkan, M., Ogut, M., et al. (2005). Evidence for theory of mind deficits in euthymic patients with bipolar disorder. Acta Psychiatrica Scandinavica 112, 110–116. doi:10.1111/j.1600-0447.2005.00570.x.

30. Bostock ECS, Kirkby KC, Garry MI, Taylor BVM (2017) Systematic review of cognitive function in euthymic bipolar disorder and pre-surgical temporal lobe epilepsy. Front Psychiatry 8: 133. doi: 10.3389/fpsyt.2017.00133.

31. Boyle EA, Li YI, Pritchard JK (2017) An expanded view of complex traits: from polygenic to omnigenic. Cell 169(7):1177–1186. doi: 10.1016/j.cell.2017.05.038. Review.

32. Brockington, I. (2005). Menstrual psychosis. World Psychiatry 4, 9–17.

33. Brockington, I. F. (2011). Menstrual Psychosis: A Bipolar Disorder with a Link to the Hypothalamus. Curr Psychiatry Rep 13, 193. doi:10.1007/s11920-011-0191-5.

34. Brotman, M. A., Guyer, A. E., Lawson, E. S., Horsey, S. E., Rich, B. A., Dickstein, D. P., et al. (2008). Facial Emotion Labeling Deficits in Children and Adolescents at Risk for Bipolar Disorder. AJP 165, 385–389. doi:10.1176/appi.ajp.2007.06122050.

35. Calvey T (2019) Human self-domestication and the extended evolutionary synthesis of addiction: how humans evolved a unique vulnerability. Neuroscience 419:100–107. doi: 10.1016/j.neuroscience.2019.09.013.

36. Cervantes, P., Gelber, S., Kin, F., Nair, V. N. P., and Schwartz, G. (2001). Circadian secretion of cortisol in bipolar disorder. J Psychiatry Neurosci 26, 411–416.

37. Chang, S. H., Gao, L., Li, Z., Zhang, W. N., Du, Y. And Wang, J. (2013) BDgene: A genetic database for bipolar disorder and its overlap with schizophrenia and major depressive disorder. Biol Psychiatry 74(10):727–33. doi: 10.1016/j.biopsych.2013.04.016

38. Correll CU, Detraux J, De Lepeleire J, De Hert M (2015) Effects of antipsychotics, antidepressants and mood stabilizers on risk for physical diseases in people with schizophrenia, depression and bipolar disorder. World Psychiatry 14: 119–136

39. Crespi B, Badcock C (2008) Psychosis and autism as diametrical disorders of the social brain. Behav Brain Sci. 31(3):241–61; discussion 261-320. doi: 10.1017/S0140525X08004214.

40. Crespi, B. J. (2016). Oxytocin, testosterone, and human social cognition: Oxytocin and social behavior. Biological Reviews 91, 390–408. doi:10.1111/brv.12175.

41. Curtis, V. A., Thompson, J. M., Seal, M. L., Monks, P. J., Lloyd, A. J., Harrison, L., et al. (2007). The nature of abnormal language processing in euthymic bipolar I disorder: evidence for a relationship between task demand and prefrontal function. Bipolar Disorders 9, 358–369. doi:10.1111/j.1399-5618.2007.00422.x.

42. Cusi, A., MacQueen, G. M., and McKinnon, M. C. (2010). Altered self-report of empathic responding in patients with bipolar disorder. Psychiatry Research 178, 354–358. doi:10.1016/j.psychres.2009.07.009.

43. da Silva EG, Pfaffenseller B, Walz J, Stertz L, Fries G, Rosa AR, Magalhães PV (2017) Peripheral insulin-like growth factor 1 in bipolar disorder. Psychiatry Res. 250:30–34. doi: 10.1016/j.psychres.2017.01.061.

44. Daban, C., Vieta, E., Mackin, P., and Young, A. H. (2005). Hypothalamic-pituitary-adrenal Axis and Bipolar Disorder. Psychiatric Clinics of North America 28, 469–480. doi:10.1016/j.psc.2005.01.005.

45. Dai L, Carter CS, Ying J, Bellugi U, Pournajafi-Nazarloo H, Korenberg JR (2012). Oxytocin and vasopressin are dysregulated in Williams syndrome, a genetic disorder affecting social behavior. PLoS ONE 7(6):e38513. Doi:10.1371/journal.pone.0038513

46. Dai, L., Carter, C. S., Ying, J., Bellugi, U., Pournajafi-Nazarloo, H., and Korenberg, J. R. (2012). Oxytocin and Vasopressin Are Dysregulated in Williams Syndrome, a Genetic Disorder Affecting Social Behavior. PLoS One 7. doi:10.1371/journal.pone.0038513.

47. Daly CJ, McGrath JC (2011) Previously unsuspected widespread cellular and tissue distribution of β-adrenoceptors and its relevance to drug action. Trends Pharmacol Sci. 32(4):219–26.

48. Daly RJ (2004) Cortactin signalling and dynamic actin networks. Biochem J. 382(Pt 1):13–25.

49. Daniels, E. K., Shenton, M. E., Holzman, P. S., Benowitz, L. I., Coleman, M., Levin, S., et al. (1988). Patterns of thought disorder associated with right cortical damage, schizophrenia, and mania. Am J Psychiatry 145, 944–949. doi:10.1176/ajp.145.8.944.

50. Derntl, B., Seidel, E.-M., Schneider, F., and Habel, U. (2012). How specific are emotional deficits? A comparison of empathic abilities in schizophrenia, bipolar and depressed patients. Schizophrenia Research 142, 58–64. doi:10.1016/j.schres.2012.09.020.

51. Detera-Wadleigh SD, Yoon SW, Berrettini WH, Goldin LR, Turner G, Yoshikawa T, Rollins DY, Muniec D, Nurnberger JI Jr, Gershon ES (1995) Adrenocorticotropin receptor/melanocortin receptor-2 maps within a reported susceptibility region for bipolar illness on chromosome 18. Am J Med Genet. 60(4):317–21.

52. Doyle, A. E., Biederman, J., Ferreira, M. A., Wong, P., Smoller, J. W., and Faraone, S. V. (2010). Suggestive linkage of the CBCL juvenile bipolar disorder phenotype to 1p21, 6p21 and 8q21. J Am Acad Child Adolesc Psychiatry 49, 378–387.

53. Dunjic-Kostic, B., Pantovic Stefanovic, M., Lackovic, M., Damjanovic, A., Jovanovic, A., Totic-Poznanovic, S., et al. (2016). Age at menarche is related to number of previous depressive episodes in patients with bipolar disorder. European Psychiatry 33, S338. doi:10.1016/j.eurpsy.2016.01.1180.

54. Durbin, C. E., Schalet, B. D., Hayden, E. P., Simpson, J., and Jordan, P. L. (2009). Hypomanic personality traits: A multi-method exploration of their association with normal and abnormal dimensions of personality. Journal of Research in Personality 43, 898–905. doi:10.1016/j.jrp.2009.04.010.

55. Dykens, E. M., Shah, B., Davis, B., Baker, C., Fife, T., and Fitzpatrick, J. (2015). Psychiatric disorders in adolescents and young adults with Down syndrome and other intellectual disabilities. J Neurodev Disord 7. doi:10.1186/s11689-015-9101-1.

56. Eastwood SL, Harrison PJ (2010) Markers of glutamate synaptic transmission and plasticity are increased in the anterior cingulate cortex in bipolar disorder. Biol. Psychiatry 67: 1010–1016. doi: 10.1016/j.biopsych.2009.12.004

57. Elboga, G., and Sayiner, Z. A. (2018). Rare cause of manic period trigger in bipolar mood disorder: testosterone replacement. Case Reports 2018, bcr-2018–225108. doi:10.1136/bcr-2018-225108.

58. Feng, G., Kang, C., Yuan, J., Zhang, Y., Wei, Y., Xu, L., et al. (2019). Neuroendocrine abnormalities associated with untreated first episode patients with major depressive disorder and bipolar disorder. Psychoneuroendocrinology 107, 119–123. doi:10.1016/j.psyneuen.2019.05.013.

59. Ferensztajn-Rochowiak E, Kaczmarek M, Wójcicka M, Kaufman-Szukalska E, Dziuda S,, Remlinger-Molenda A, Szeliga-Neymann A, Losy J, Rybakowski JK (2019) Glutamate-related antibodies and peripheral insulin-like growth factor in bipolar disorder and lithium prophylaxis. Neuropsychobiology 77(1):49–56. doi: 10.1159/000493740.

60. Freinhar, J. P., and Alvarez, W. (1985). Androgen-induced hypomania. J Clin Psychiatry 46, 354–355.

61. Fries, G. R., Vasconcelos-Moreno, M. P., Gubert, C., dos Santos, B. T. M. Q., Sartori, J., Eisele, B., et al. (2014). Hypothalamic-Pituitary-Adrenal Axis Dysfunction and Illness Progression in Bipolar Disorder. Int J Neuropsychopharmacol 18. doi:10.1093/ijnp/pyu043.

62. Gibson G (2009) Decanalization and the origin of complex disease. Nature Rev. Genet. 10: 134–140.

63. Gigante AD, Bond DJ, Lafer B, Lam RW, Young LT, Yatham LN (2012) Brain glutamate levels measured by magnetic resonance spectroscopy in patients with bipolar disorder: a meta-analysis. Bipolar Disord. 14: 478–487. doi: 10.1111/j.1399-5618.2012.01033.x

64. Girshkin, L., Matheson, S. L., Shepherd, A. M., and Green, M. J. (2014). Morning cortisol levels in schizophrenia and bipolar disorder: A meta-analysis. Psychoneuroendocrinology 49, 187–206. doi:10.1016/j.psyneuen.2014.07.013.

65. Givalois L, Arancibia S, Tapia-Arancibia L (2000) Concomitant changes in CRH mRNA levels in rat hippocampus and hypothalamus following immobilization stress. Mol. Brain Res. 75: 166–171

66. Gordovez FJA, McMahon FJ (2020) The genetics of bipolar disorder. Mol Psychiatry 25(3):544–559. doi: 10.1038/s41380-019-0634-7.

67. Gottschalk MG, Wesseling H, Guest PC, Bahn S (2015) Proteomic enrichment analysis of psychotic and affective disorders reveals common signatures in presynaptic glutamatergic signaling and energy metabolism. Int. J. Neuropsychopharmacol. 18: pyu019. doi: 10.1093/ijnp/pyu019

68. Gray NW, Kruchten AE, Chen J, McNiven MA (2005) A dynamin-3 spliced variant modulates the actin/cortactin-dependent morphogenesis of dendritic spines. J Cell Sci. 118(Pt 6):1279–90.

69. Green, E., Elvidge, G., Jacobsen, N., Glaser, B., Jones, I., O’Donovan, M. C., et al. (2005). Localization of Bipolar Susceptibility Locus by Molecular Genetic Analysis of the Chromosome 12q23-q24 Region in Two Pedigrees With Bipolar Disorder and Darier’s Disease. AJP 162, 35–42. doi:10.1176/appi.ajp.162.1.35.

70. Greenwood, T. A., Akiskal, H. S., Akiskal, K. K., and Kelsoe, J. R. (2012). Genome-Wide Association Study of Temperament in Bipolar Disorder Reveals Significant Associations with Three Novel Loci. Biological Psychiatry 72, 303–310. doi:10.1016/j.biopsych.2012.01.018.

71. Guha, P., Bhowmick, K., Mazumder, P., Ghosal, M., Chakraborty, I., and Burman, P. (2014). Assessment of Insulin Resistance and Metabolic Syndrome in Drug Naive Patients of Bipolar Disorder. Indian J Clin Biochem 29, 51–56. doi:10.1007/s12291-012-0292-x.

72. Gutiérrez, B., Van Os, J., Vallès, V., Guillamat, R., Campillo, M., and Fañanás, L. (1998). Congenital dermatoglyphic malformations in severe bipolar disorder. Psychiatry Research 78, 133–140. doi:10.1016/S0165-1781(98)00016-X.

73. Gutiérrez-Sacristán A, Grosdidier S, Valverde O, Torrens M, Bravo À, Piñero J, Sanz F, Furlong LI (2015) PsyGeNET: a knowledge platform on psychiatric disorders and their genes. Bioinformatics 31(18):3075–7. doi: 10.1093/bioinformatics/btv301.

74. Hajek, T., Kopecek, M., Höschl, C., and Alda, M. (2012). Smaller hippocampal volumes in patients with bipolar disorder are masked by exposure to lithium: a meta-analysis. J Psychiatry Neurosci 37, 333–343. doi:10.1503/jpn.110143.

75. Hallahan, B., Newell, J., Soares, J. C., Brambilla, P., Strakowski, S. M., Fleck, D. E., et al. (2011). Structural Magnetic Resonance Imaging in Bipolar Disorder: An International Collaborative Mega-Analysis of Individual Adult Patient Data. Biological Psychiatry 69, 326–335. doi:10.1016/j.biopsych.2010.08.029.

76. Han, K.-M., Kim, A., Kang, W., Kang, Y., Kang, J., Won, E., et al. (2019). Hippocampal subfield volumes in major depressive disorder and bipolar disorder. European Psychiatry 57, 70–77. doi:10.1016/j.eurpsy.2019.01.016.

77. Hanford LC, Nazarov A, Hall GB, Sassi RB (2016) Cortical thickness in bipolar disorder: a systematic review. Bipolar Disord. 18(1):4–18. doi: 10.1111/bdi.12362.

78. Hare B (2017) Survival of the friendliest: Homo sapiens evolved via selection for prosociality. Ann Rev Psychol 68: 24.1–24.32 DOI: 10.1146/annurev-psych-010416-044201

79. Hare B, Wobber V, Wrangham R (2012) The self-domestication hypothesis: Evolution of bonobo psychology is due to selection against aggression. Animal Behav 83(3), 573–585.

80. Hart BL (1985) The behaviour of domestic animals. New York: W.H. Freeman and Company.

81. Hashimoto K, Sawa A, Iyo M (2007) Increased levels of glutamate in brains from patients with mood disorders. Biol. Psychiatry 62: 1310–1316. doi: 10.1016/j.biopsych.2007.03.017

82. Hawken, E. R., Harkness, K. L., Lazowski, L. K., Summers, D., Khoja, N., Gregory, J. G., et al. (2016). The manic phase of Bipolar disorder significantly impairs theory of mind decoding. Psychiatry Research 239, 275–280. doi:10.1016/j.psychres.2016.03.043.

83. Hayes JF, Miles J, Walters K, King M, Osborn DPJ (2015) A systematic review and meta-analysis of premature mortality in bipolar affective disorder. Acta Psychiatr. Scand. 131: 417–425, 10.1111/acps.12408

84. Hennessy, R. J., Baldwin, P. A., Browne, D. J., Kinsella, A., and Waddington, J. L. (2010). Frontonasal dysmorphology in bipolar disorder by 3D laser surface imaging and geometric morphometrics: Comparisons with schizophrenia. Schizophr Res 122, 63–71. doi:10.1016/j.schres.2010.05.001.

85. Hering H, Sheng M (2003) Activity-dependent redistribution and essential role of cortactin in dendritic spine morphogenesis. J Neurosci. 23(37):11759–69.

86. Herrmann E, Hare B, Cissewski J, Tomasello MA (2011) Comparison of temperament in nonhuman apes and human infants. Dev Sci 14: 1393–1405.

87. Himmler BT, Stryjek R, Modlinska K, Derksen SM, Pisula W, Pellis SM. (2013). How domestication modulates play behavior: a comparative analysis between wild ratsand a laboratory strain of Rattus norvegicus. J Comp Psychol. 127(4):453–64. doi: 10.1037/a0032187.

88. Insel PA (2011) β(2)-Adrenergic receptor polymorphisms and signaling: Do variants influence the “memory” of receptor activation? Sci Signal 4(185):pe37.

89. Iovino, M., Messana, T., De Pergola, G., Iovino, E., Dicuonzo, F., Guastamacchia, E., et al. (2018). The Role of Neurohypophyseal Hormones Vasopressin and Oxytocin in Neuropsychiatric Disorders. Endocrine, Metabolic & Immune Disorders - Drug Targets 18, 341–347. doi:10.2174/1871530318666180220104900.

90. Jakobson CM, Jarosz DF (2019) Molecular origins of complex heritability in natural genotype-to-phenotype relationships. Cell Syst. 8(5):363–379.e3. doi: 10.1016/j.cels.2019.04.002.

91. Joffe, H., Kim, D. R., Foris, J. M., Baldassano, C. F., Gyulai, L., Hwang, C. H., et al. (2006). Menstrual Dysfunction Prior to Onset of Psychiatric Illness Is Reported More Commonly by Women With Bipolar Disorder Than by Women With Unipolar Depression and Healthy Controls. J Clin Psychiatry 67, 297–304.

92. Joiner ML, Lisé MF, Yuen EY, Kam AY, Zhang M, Hall DD, Malik ZA, Qian H, Chen Y, Ulrich JD, Burette AC, Weinberg RJ, Law PY, El-Husseini A, Yan Z, Hell JW (2010) Assembly of a beta2-adrenergic receptor--GluR1 signalling complex for localized cAMP signalling. EMBO J. 29(2):482–95.

93. Judd LL, Schettler PJ, Solomon DA, Maser JD, Coryell W, Endicott J et al. (2008) Psychosocial disability and work role function compared across the long-term course of bipolar I, bipolar II and unipolar major depressive disorders. J Affect Disord 108: 49–58.

94. Kaiser S, Hennessy MB, Sachser N (2015) Domestication affects the structure, development and stability of biobehavioural profiles. Front Zool 12(Suppl 1):S19. doi:10.1186/1742-9994-12-S1-S19.

95. Kawahara, K., Jono, T., Nishi, Y., Ushijima, H., and Ikeda, M. (2015). Effects of testosterone therapy on bipolar disorder with Klinefelter syndrome. General Hospital Psychiatry 37, 192.e1–192.e2. doi:10.1016/j.genhosppsych.2014.12.003.

96. Keller-Wood M (2015) Hypothalamic-pituitary--adrenal axis-feedback control. Compr Physiol. 5(3):1161–82. doi: 10.1002/cphy.c140065.

97. Kerr, N., Dunbar, R. I. M., and Bentall, R. P. (2003). Theory of mind deficits in bipolar affective disorder. Journal of Affective Disorders 73, 253–259. doi:10.1016/S0165-0327(02)00008-3.

98. Kesebir, S., Yaşan Şair, B., Unübol, B., and Tatlıdil Yaylacı, E. (2013). Is there a relationship between age at menarche and clinical and temperamental characteristics in bipolar disorder? Ann Clin Psychiatry 25, 121–124.

99. Keshri, N., Nandeesha, H., and Kattimani, S. (2018). Elevated interleukin-17 and reduced testosterone in bipolar disorder. Relation with suicidal behaviour. Asian Journal of Psychiatry 36, 66–68. doi:10.1016/j.ajp.2018.06.011.

100. Kisko, T. M., Braun, M. D., Michels, S., Witt, S. H., Rietschel, M., Culmsee, C., et al. (2018). Cacna1c haploinsufficiency leads to pro-social 50-kHz ultrasonic communication deficits in rats. Dis Model Mech 11. doi:10.1242/dmm.034116.

101. Kohler, C. G., Hoffman, L. J., Eastman, L. B., Healey, K., and Moberg, P. J. (2011). Facial emotion perception in depression and bipolar disorder: A quantitative review. Psychiatry Research 188, 303–309. doi:10.1016/j.psychres.2011.04.019.

102. Kruska D (1988) Mammalian domestication and its effect on brain structure and behavior. In: Jerison HJ, Jerison I. (eds) Intelligence and Evolutionary Biology. NATO ASI Series (Series G: Ecological Sciences), vol 17. Springer, Berlin, Heidelberg

103. Kruska D, Schott U (1977) Comparative-quantitative investigations of brains of wild and laboratory rats. J Hirnforsch. 18(1):59–67.

104. Kruska DC (2005). On the evolutionary significance of encephalization in some eutherian mammals: effects of adaptive radiation, domestication, and feralization. Brain Behav Evol. 65(2):73–108.

105. Kubo Y, Baba K, Toriyama M, Minegishi T, Sugiura T, Kozawa S, Ikeda K, Inagaki N. (2015) Shootin1-cortactin interaction mediates signal-force transduction for axon outgrowth. J Cell Biol. 210(4):663–76. doi: 10.1083/jcb.201505011.

106. Lambert, S., Maystadt, I., Boulanger, S., Vrielynck, P., Destrée, A., Lederer, D., et al. (2016). Expanding phenotype of p.Ala140Val mutation in MECP2 in a 4 generation family with X-linked intellectual disability and spasticity. Eur J Med Genet 59, 522–525. doi:10.1016/j.ejmg.2016.07.003.

107. Lanz TA, Reinhart V, Sheehan MJ, Rizzo SJS, Bove SE, James LC, Volfson D, Lewis DA, Kleiman RJ (2019) Postmortem transcriptional profiling reveals widespread increase in inflammation in schizophrenia: a comparison of prefrontal cortex, striatum, and hippocampus among matched tetrads of controls with subjects diagnosed with schizophrenia, bipolar or major depressive disorder. Transl Psychiatry 9(1):151. doi: 10.1038/s41398-019-0492-8.

108. Leach HM (2003) Human domestication reconsidered. Curr Anthropol 44: 349–368

109. Lee H.-C., Chang D.-E., Yeom M., Kim G.-H., Choi K.-D., Shim I., Lee H.-J., Hahm D.-H (2005) Gene expression profiling in hypothalamus of immobilization-stressed mouse using cDNA microarray. Mol. Brain Res. 135: 293–300

110. Leszczynska-Rodziewicz, A., Maciukiewicz, M., Szczepankiewicz, A., Poglodzinski, A., Hauser, J. (2013) Association between OPCRIT dimensions and polymorphisms of HPA axis genes in bipolar disorder. J. Affect. Disord. 151: 744–747

111. Leszczynska-Rodziewicz, A., Szczepankiewicz, A., Dmitrzak-Weglarz, M., Skibinska, M., Hauser, J. (2012) Association between functional polymorphism of the AVPR1b gene and polymorphism rs1293651 of the CRHR1 gene and bipolar disorder with psychotic features. J. Affect. Disord., 138 (3): 490–493

112. Lett TA, Zai CC, Tiwari AK, Shaikh SA, Likhodi O, Kennedy JL, Müller DJ (2011) ANK3, CACNA1C and ZNF804A gene variants in bipolar disorders and psychosis subphenotype. World J Biol Psychiatry. 2011;12(5):392–397.

113. Li Y, Wang GD, Wang MS, Irwin DM, Wu DD, Zhang YP (2014) Domestication of the dog from the wolf was promoted by enhanced excitatory synaptic plasticity: a hypothesis. Genome Biol Evol. 6(11):3115–21. doi: 10.1093/gbe/evu245.

114. Liberg B, Klauser P, Harding IH, Adler M, Rahm C, Lundberg J, Masterman T, Wachtler C, Jonsson T, Kristoffersen-Wiberg M, Pantelis C, Wahlund B (2014) Functional and structural alterations in the cingulate motor area relate to decreased fronto-striatal coupling in major depressive disorder with psychomotor disturbances. Front Psychiatry 5:176. doi: 10.3389/fpsyt.2014.00176.

115. Lien, Y.-J., Chang, H. H., Tsai, H.-C., Kuang Yang, Y., Lu, R.-B., and See Chen, P. (2017). Plasma oxytocin levels in major depressive and bipolar II disorders. Psychiatry Research 258, 402–406. doi:10.1016/j.psychres.2017.08.080.

116. Lightman S, Young 3rd W (1988) Corticotrophin-releasing factor, vasopressin and pro-opiomelanocortin mRNA responses to stress and opiates in the rat. J. Physiol. 403: 511

117. Lin SH, Chen PS, Yeh TL, Yang YK (2011) The dopamine hypothesis of social support. Med Hypotheses 77(5):753–5. doi: 10.1016/j.mehy.2011.07.030.

118. Litonjua AA, Gong L, Duan QL, Shin J, Moore MJ, Weiss ST, Johnson JA, Klein TE, Altman RB (2010) Very important pharmacogene summary ADRB2. Pharmacogenet Genomics. 20(1):64–9. doi: 10.1097/FPC.0b013e3

119. Lloyd, T., Dazzan, P., Dean, K., Park, S. B. G., Fearon, P., Doody, G. A., et al. (2008). Minor physical anomalies in patients with first-episode psychosis: their frequency and diagnostic specificity. Psychological Medicine 38, 71–77. doi:10.1017/S0033291707001158.

120. Lo, M.-T., Hinds, D. A., Tung, J. Y., Franz, C., Fan, C.-C., Wang, Y., et al. (2017). Genome-wide analyses for personality traits identify six genomic loci and show correlations with psychiatric disorders. Nature Genetics 49, 152–156. doi:10.1038/ng.3736.

121. Løtvedt P, Fallahshahroudi A, Bektic L, Altimiras J, Jensen P (2017) Chicken domestication changes expression of stress-related genes in brain, pituitary and adrenals. Neurobiol Stress. 7:113–121. doi: 10.1016/j.ynstr.2017.08.002.

122. MacGillavry HD, Kerr JM, Kassner J, Frost NA, Blanpied TA (2016) Shank-cortactin interactions control actin dynamics to maintain flexibility of neuronal spines and synapses. Eur J Neurosci. 43(2):179–93. doi: 10.1111/ejn.13129. Epub 2015 Dec 22.

123. Malhi GS, Ivanovski B, Hadzi-Pavlovic D, Mitchell PB, Vieta E, Sachdev P (2007) Neuropsychological deficits and functional impairment in bipolar depression, hypomania and euthymia. Bipolar Disord 9: 114–125. 10.1111/j.1399-5618.2007.00324.x.

124. McGrath, J., El-Saadi, O., Grim, V., Cardy, S., Chapple, B., Chant, D., et al. (2002). Minor Physical Anomalies and Quantitative Measures of the Head and Face in Patients With Psychosis. Arch Gen Psychiatry 59, 458–464. doi:10.1001/archpsyc.59.5.458.

125. Meyer, T. D. (2002). The Hypomanic Personality Scale, the Big Five, and their relationship to depression and mania. Personality and Individual Differences 32, 649–660. doi:10.1016/S0191-8869(01)00067-8.

126. Mohite S, Sanches M, Teixeira AL (2019) Exploring the evidence implicating the renin-angiotensin system (RAS) in the physiopathology of mood disorders. Protein Pept Lett. doi: 10.2174/0929866527666191223144000.

127. Moon S, Kim TH, Lee KT, Kwak W, Lee T, Lee SW, Kim MJ, Cho K, Kim N, Chung WH, Sung S, Park T, Cho S, Groenen MA, Nielsen R, Kim Y, Kim H (2015) A genome-wide scan for signatures of directional selection in domesticated pigs. BMC Genomics. 16:130. doi: 10.1186/s12864-015-1330-x.

128. Mousavizadegan, S., and Maroufi, M. (2018). Comparison of salivary testosterone levels in different phases of bipolar I disorder and control group. J Res Med Sci 23. doi:10.4103/jrms.JRMS_1009_17.

129. Myers, L., Anderlid, B.-M., Nordgren, A., Willfors, C., Kuja-Halkola, R., Tammimies, K., et al. (2017). Minor physical anomalies in neurodevelopmental disorders: a twin study. Child Adolesc Psychiatry Ment Health 11. doi:10.1186/s13034-017-0195-y.

130. Najt P, Wang F, Spencer L, Johnston JA, Cox Lippard ET, Pittman BP, Lacadie C, Staib LH, Papademetris X, Blumberg HP (2016) Anterior cortical development during adolescence in bipolar disorder. Biol Psychiatry 79(4):303–10. doi: 10.1016/j.biopsych.2015.03.026.

131. Nasrallah, H. A. (1991). Neurodevelopmental aspects of bipolar affective disorder. Biological Psychiatry 29, 1–2. doi:10.1016/0006-3223(91)90205-Z.

132. Naumenko EV, Belyaev DK (1980) Neuroendocrine mechanisms in animal domestication. In: Belyaev DK, editor. Problems in general genetics. Proceedings XIY Int Cong Genet. 2, 2. Moscow: Mir; pp. 12–24.

133. Negash A, Kebede D, Alem A, Melaku Z, Deyessa N, Shibire T, Fekadu A, Fekadu D, Jacobsson L, Kullgren G: Neurological soft signs in bipolar I disorder patients (2004) J Affect Disord. 80: 221–230. 10.1016/S0165-0327(03)00116-2.

134. Niego A, Benítez-Burraco A (2019) Williams syndrome, human self-domestication, and language evolution. Front Psychol 10: 521.

135. Niego, A., and Benítez-Burraco, A. (2019). Williams Syndrome, Human Self-Domestication, and Language Evolution. Front. Psychol. 10. doi:10.3389/fpsyg.2019.00521.

136. Nitzburg, G. C., Burdick, K. E., Malhotra, A. K., and DeRosse, P. (2015). Social cognition in patients with schizophrenia spectrum and bipolar disorders with and without psychotic features. Schizophrenia Research: Cognition 2, 2–7. doi:10.1016/j.scog.2014.12.003.

137. Noga, J. T., Vladar, K., and Torrey, E. F. (2001). A volumetric magnetic resonance imaging study of monozygotic twins discordant for bipolar disorder. Psychiatry Research: Neuroimaging 106, 25–34. doi:10.1016/S0925-4927(00)00084-6.

138. O’Rourke T, Boeckx C (2020) Glutamate receptors in domestication and modern human evolution. Neurosci Biobehav Rev. 108:341–357. doi: 10.1016/j.neubiorev.2019.10.004.

139. Panksepp, J. (2005). Beyond a Joke: From Animal Laughter to Human Joy? Science 308, 62–63. doi:10.1126/science.1112066.

140. Peedicayil J, Grayson DR (2018a) An epigenetic basis for an omnigenic model of psychiatric disorders. J Theor Biol. 2018 443:52–55. doi: 10.1016/j.jtbi.2018.01.027.

141. Peedicayil J, Grayson DR (2018b) Some implications of an epigenetic-based omnigenic model of psychiatric disorders. J Theor Biol. 452:81–84. doi: 10.1016/j.jtbi.2018.05.014.

142. Perez-Rodriguez, M. M., Mahon, K., Russo, M., Ungar, A. K., and Burdick, K. E. (2015). Oxytocin and Social Cognition in Affective and Psychotic Disorders. Eur Neuropsychopharmacol 25, 265–282. doi:10.1016/j.euroneuro.2014.07.012.

143. Perich, T. A., Roberts, G., Frankland, A., Sinbandhit, C., Meade, T., Austin, M.-P., et al. (2017). Clinical characteristics of women with reproductive cycle–associated bipolar disorder symptoms. Australian & New Zealand Journal of Psychiatry 51, 161–167. doi:10.1177/0004867416670015.

144. Pisor AC, Surbeck M (2019) The evolution of intergroup tolerance in nonhuman primates and humans. Evol Anthropol 28(4):210–223.

145. Platt, J., Colich, N. L., McLaughlin, K. A., Gary, D., and Keyes, K. M. (2017). Transdiagnostic psychiatric disorder risk associated with early age of menarche: a latent modeling approach. Compr Psychiatry 79, 70–79. doi:10.1016/j.comppsych.2017.06.010.

146. Plavcan JM (2012) Sexual size dimorphism, canine dimorphism, and male-male competition in primates. Hum Nat. 23:45–67.

147. Polimanti R, Gelernter J (2017) Widespread signatures of positive selection in common risk alleles associated to autism spectrum disorder. PLoS Genet. 13(2):e1006618. doi: 10.1371/journal.pgen.1006618

148. Post, R. M., Speer, A. M., Hough, C. J., and Xing, G. (2003). Neurobiology of bipolar illness: implications for future study and therapeutics. Ann Clin Psychiatry 15, 85–94.

149. Psychiatric GWAS Consortium Bipolar Disorder Working Group (2011) Large-scale genome-wide association analysis of bipolar disorder identifies a new susceptibility locus near ODZ4. Nat Genet. 2011;43(10):977–983.

150. Pugliese, V., Bruni, A., Carbone, E. A., Calabrò, G., Cerminara, G., Sampogna, G., et al. (2019). Maternal stress, prenatal medical illnesses and obstetric complications: Risk factors for schizophrenia spectrum disorder, bipolar disorder and major depressive disorder. Psychiatry Research 271, 23–30. doi:10.1016/j.psychres.2018.11.023.

151. Qadri, S., Hussain, A., Bhat, M. H., and Baba, A. A. (2018). Polycystic Ovary Syndrome in Bipolar Affective Disorder: A Hospital-based Study. Indian J Psychol Med 40, 121–128. doi:10.4103/IJPSYM.IJPSYM_284_17.

152. Quilty, L. C., Sellbom, M., Tackett, J. L., and Bagby, R. M. (2009). Personality trait predictors of bipolar disorder symptoms. Psychiatry Research 169, 159–163. doi:10.1016/j.psychres.2008.07.004.

153. Racz B, Weinberg RJ (2004) The subcellular organization of cortactin in hippocampus. J Neurosci. 24(46):10310–7.

154. Raghanti MA, Edler MK, Stephenson AR, Wilson LJ, Hopkins WD, Ely JJ, Erwin JM, Jacobs B, Hof PR, Sherwood CC (2016) Human-specific increase of dopaminergic innervation in a striatal region associated with speech and language: A comparative analysis of the primate basal ganglia. J Comp Neurol 524(10):2117–29. doi: 10.1002/cne.23937.

155. Rasgon, N. L., Altshuler, L. L., Fairbanks, L., Elman, S., Bitran, J., Labarca, R., et al. (2005a). Reproductive function and risk for PCOS in women treated for bipolar disorder. Bipolar Disord 7, 246–259. doi:10.1111/j.1399-5618.2005.00201.x.

156. Rasgon, N. L., Reynolds, M. F., Elman, S., Saad, M., Frye, M. A., Bauer, M., et al. (2005b). Longitudinal evaluation of reproductive function in women treated for bipolar disorder. Journal of Affective Disorders 89, 217–225. doi:10.1016/j.jad.2005.08.002.

157. Rasgon, N., Bauer, M., Glenn, T., Elman, S., and Whybrow, P. C. (2003). Menstrual cycle related mood changes in women with bipolar disorder. Bipolar Disord 5, 48–52.

158. Raucher-Chéné, D., Achim, A. M., Kaladjian, A., and Besche-Richard, C. (2017). Verbal fluency in bipolar disorders: A systematic review and meta-analysis. Journal of Affective Disorders 207, 359–366. doi:10.1016/j.jad.2016.09.039.

159. Reilly, T. J., Sagnay de la Bastida, V. C., Joyce, D. W., Cullen, A. E., and McGuire, P. (2019). Exacerbation of Psychosis During the Perimenstrual Phase of the Menstrual Cycle: Systematic Review and Meta-analysis. Schizophr Bull. doi:10.1093/schbul/sbz030.

160. Rilling JK, Scholz J, Preuss TM, Glasser MF, Errangi BK, Behrens TE (2012). Differences between chimpanzees and bonobos in neural systems supporting socialcognition. Soc Cogn Affect Neurosci. 7(4):369–79. doi: 10.1093/scan/nsr017.

161. Rolls ET (2015) Limbic systems for emotion and for memory, but no single limbic system. Cortex 62:119–57. doi: 10.1016/j.cortex.2013.12.005.

162. Ruan C, Zhang Z (2016) Laboratory domestication changed the expression patterns of oxytocin and vasopressin in brains of rats and mice. Anat Sci Int. 91(4):358–70. doi: 10.1007/s12565-015-0311-0.

163. Rutigliano, G., Rocchetti, M., Paloyelis, Y., Gilleen, J., Sardella, A., Cappucciati, M., et al. (2016). Peripheral oxytocin and vasopressin: Biomarkers of psychiatric disorders? A comprehensive systematic review and preliminary meta-analysis. Psychiatry Research 241, 207–220. doi:10.1016/j.psychres.2016.04.117.

164. Salgado, H., and Martins-Correia, L. (2014). Williams syndrome and psychosis: a case report. J Med Case Rep 8, 49. doi:10.1186/1752-1947-8-49.

165. Samango‐Sprouse, C. A., Sadeghin, T., Mitchell, F. L., Dixon, T., Stapleton, E., Kingery, M., et al. (2013). Positive effects of short course androgen therapy on the neurodevelopmental outcome in boys with 47,XXY syndrome at 36 and 72 months of age. American Journal of Medical Genetics Part A 161, 501–508. doi:10.1002/ajmg.a.35769.

166. Sanches, M., Keshavan, M. S., Brambilla, P., and Soares, J. C. (2008). Neurodevelopmental basis of bipolar disorder: A critical appraisal. Progress in Neuro-Psychopharmacology and Biological Psychiatry 32, 1617–1627. doi:10.1016/j.pnpbp.2008.04.017.

167. Sánchez-Villagra, M. R., Geiger, M., and Schneider, R. A. (2016). The taming of the neural crest: a developmental perspective on the origins of morphological covariation in domesticated mammals. R Soc Open Sci 3. doi:10.1098/rsos.160107.

168. Santos, J. M., Pousa, E., Soto, E., Comes, A., Roura, P., Arrufat, F. X., et al. (2017). Theory of Mind in Euthymic Bipolar Patients and First-Degree Relatives. J. Nerv. Ment. Dis. 205, 207–212. doi:10.1097/NMD.0000000000000595.

169. Sapolsky RM, Romero LM, Munck AU (2000) How do glucocorticoids influence stress responses? Integrating permissive, suppressive, stimulatory, and preparative actions. Endocr Rev. 21(1):55–89.

170. Savla, G. N., Vella, L., Armstrong, C. C., Penn, D. L., and Twamley, E. W. (2013). Deficits in domains of social cognition in schizophrenia: a meta-analysis of the empirical evidence. Schizophr Bull 39, 979–992. doi:10.1093/schbul/sbs080.

171. Seidel, E.-M., Habel, U., Finkelmeyer, A., Hasmann, A., Dobmeier, M., and Derntl, B. (2012). Risk or resilience? Empathic abilities in patients with bipolar disorders and their first-degree relatives. Journal of Psychiatric Research 46, 382–388. doi:10.1016/j.jpsychires.2011.11.006.

172. Selvaraj S, Arnone D, Job D, Stanfield A, Farrow TF, Nugent AC, Scherk H, Gruber O, Chen X, Sachdev PS, Dickstein DP, Malhi GS, Ha TH, Ha K, Phillips ML, McIntosh AM (2012) Grey matter differences in bipolar disorder: a meta-analysis of voxel-based morphometry studies. Bipolar Disord. 14(2):135–45. doi: 10.1111/j.1399-5618.2012.01000.x.

173. Shea B (1989) Heterochrony in human evolution: the case for neoteny reconsidered. Am J Phys Anthropol 32: 69e101.

174. Sher, L., and Landers, S. (2014). Bipolar disorder, testosterone administration, and homicide: A case report. International Journal of Psychiatry in Clinical Practice 18, 215–216. doi:10.3109/13651501.2014.894075.

175. Sher, L., Grunebaum, M. F., Sullivan, G. M., Burke, A. K., Cooper, T. B., Mann, J. J., et al. (2012). Testosterone levels in suicide attempters with bipolar disorder. J Psychiatr Res 46. doi:10.1016/j.jpsychires.2012.06.016.

176. Sher, L., Grunebaum, M. F., Sullivan, G. M., Burke, A. K., Cooper, T. B., Mann, J. J., et al. (2014). Association of testosterone levels and future suicide attempts in females with bipolar disorder. J Affect Disord 166, 98–102. doi:10.1016/j.jad.2014.04.068.

177. Singh, D., Sasson, A., Rusciano, V., Wakimoto, Y., Pinkhasov, A., and Angulo, M. (2019). Cycloid Psychosis Comorbid with Prader-Willi Syndrome: A Case Series. Am. J. Med. Genet. A 179, 1241–1245. doi:10.1002/ajmg.a.61181.

178. Smith AK, Dimulescu I, Falkenberg VR, Narasimhan S, Heim C, Vernon SD, Rajeevan MS (2008) Genetic evaluation of the serotonergic system in chronic fatigue syndrome. Psychoneuroendocrinology 33(2): 188–197. 10.1016/j.psyneuen.2007.11.001.

179. Smith, A.I. Funder, J.W. (1988) Proopiomelanocortin processing in the pituitary, central nervous system and peripheral tissues. Endocr. Rev. 9, 159–179.

180. Soeiro-de-Souza, M. G., Otaduy, M. C. G., Dias, C. Z., Bio, D. S., Machado-Vieira, R., and Moreno, R. A. (2012). The impact of the CACNA1C risk allele on limbic structures and facial emotions recognition in bipolar disorder subjects and healthy controls. Journal of Affective Disorders 141, 94–101. doi:10.1016/j.jad.2012.03.014.

182. Somel M, Franz H, Yan Z, Lorenc A, Guo S, Giger T, Kelso J, Nickel B, Dannemann M, Bahn S, Webster MJ, Weickertm CS, Lachmann M, Pääbo S, Khaitovich P (2009) Transcriptional neoteny in the human brain. Proc Natl Acad Sci U S A. 106(14):5743–8. doi: 10.1073/pnas.0900544106.

183. Srinivasan S, Bettella F, Mattingsdal M, Wang Y, Witoelar A, Schork AJ, Thompson WK, Zuber V; Schizophrenia Working Group of the Psychiatric Genomics Consortium, The International Headache Genetics Consortium, Winsvold BS, Zwart JA, Collier DA, Desikan RS, Melle I, Werge T, Dale AM, Djurovic S, Andreassen OA. (2016) Genetic markers of human evolution are enriched in schizophrenia. Biol Psychiatry 80(4):284–292. doi: 10.1016/j.biopsych.2015.10.009.

184. Stelzhammer V, Alsaif M, Chan MK, Rahmoune H, Steeb H, Guest PC, Bahn S (2015) Distinct proteomic profiles in post-mortem pituitary glands from bipolar disorder and major depressive disorder patients. Psychiatr Res. 60:40–8. doi: 10.1016/j.jpsychires.2014.09.022.

185. Strakowski, S. M., DelBello, M. P., Sax, K. W., Zimmerman, M. E., Shear, P. K., Hawkins, J. M., et al. (1999). Brain magnetic resonance imaging of structural abnormalities in bipolar disorder. Arch. Gen. Psychiatry 56, 254–260.

186. Striepens, N., Kendrick, K. M., Maier, W., and Hurlemann, R. (2011). Prosocial effects of oxytocin and clinical evidence for its therapeutic potential. Frontiers in Neuroendocrinology 32, 426–450. doi:10.1016/j.yfrne.2011.07.001.

187. Stringer C (2016) The origin and evolution of Homo sapiens. Philos Trans R Soc Lond B Biol Sci 371: 20150237.

188. Teatero, M. L., Mazmanian, D., and Sharma, V. (2014). Effects of the menstrual cycle on bipolar disorder. Bipolar Disorders 16, 22–36. doi:10.1111/bdi.12138.

189. Tényi, T., Trixler, M., and Csábi, G. (2009). Minor physical anomalies in affective disorders. A review of the literature. Journal of Affective Disorders 112, 11–18. doi:10.1016/j.jad.2008.04.025.

190. Theofanopoulou C (2016) Implications of oxytocin in human linguistic cognition: from genome to phenome. Front Neurosci. 10:271. doi: 10.3389/fnins.2016.00271.

191. Theofanopoulou C, Boeckx C, Jarvis ED (2017) A hypothesis on a role of oxytocin in the social mechanisms of speech and vocal learning. Proc Biol Sci. 284(1861), pii: 20170988. doi: 10.1098/rspb.2017.0988.

192. Thomas, J., and Kirby, S. (2018). Self domestication and the evolution of language. Biol Philos 33, 9. doi:10.1007/s10539-018-9612-8.

193. Tondo, L., Pinna, M., Serra, G., Chiara, L. D., and Baldessarini, R. J. (2017). Age at menarche predicts age at onset of major affective and anxiety disorders. European Psychiatry 39, 80–85. doi:10.1016/j.eurpsy.2016.08.001.

194. Toro R, Konyukh M, Delorme R, Leblond C, Chaste P, Fauchereau F, Coleman M, Leboyer M, Gillberg C, Bourgeron T (2010) Key role for gene dosage and synaptic homeostasis in autism spectrum disorders. Trends Genet. 26: 363–72.

195. Trixler, M., Tényi, T., Csábi, G., and Szabó, R. (2001). Minor physical anomalies in schizophrenia and bipolar affective disorder. Schizophrenia Research 52, 195–201. doi:10.1016/S0920-9964(00)00182-1.

196. Trut L (1999) Early canid domestication: the farm-fox experiment: foxes bred for tamability in a 40-year experiment exhibit remarkable transformations that suggest an interplay between behavioral genetics and development. Am Sci 87(2):160–169.

197. Trut L, Oskina I, Kharlamova A (2009) Animal evolution during domestication: the domesticated fox as a model. BioEssays 31(3):349–360. doi:10.1002/bies.200800070.

198. Trut LN1, Pliusnina IZ, Kolesnikova LA, Kozlova ON (2000) Interhemisphere neurochemical differences in the brain of silver foxes selected for behavior and the problem of directed asymmetry. Genetika 36(7):942–6.

199. Trut, L. N. (1999). Early Canid Domestication: The Farm-Fox Experiment. American Scientist 87. Available at: http://adsabs.harvard.edu/abs/1999AmSci..87.T [Accessed March 12, 2020].

200. Turan, T., Uysal, C., Asdemir, A., and Kılıç, E. (2013). May oxytocin be a trait marker for bipolar disorder? Psychoneuroendocrinology 38, 2890–2896. doi:10.1016/j.psyneuen.2013.07.017.

201. Ueda S, Negishi M, Katoh H (2013) Rac GEF Dock4 interacts with cortactin to regulate dendritic spine formation. Mol Biol Cell. 24(10):1602–13. doi: 10.1091/mbc.E12-11-0782.

202. Van Rheenen, T. E., and Rossell, S. L. (2013). Auditory-prosodic processing in bipolar disorder; from sensory perception to emotion. Journal of Affective Disorders 151, 1102–1107. doi:10.1016/j.jad.2013.08.039.

203. Van Rheenen, T. E., and Rossell, S. L. (2014). Phenomenological predictors of psychosocial function in bipolar disorder: Is there evidence that social cognitive and emotion regulation abnormalities contribute? Aust N Z J Psychiatry 48, 26–35. doi:10.1177/0004867413508452.

204. Vistein R, Puthenveedu MA (2014) Src regulates sequence-dependent beta-2 adrenergic receptor recycling via cortactin phosphorylation. Traffic 15(11):1195–205. doi: 10.1111/tra.12202.

205. vonHoldt, B. M., Shuldiner, E., Koch, I. J., Kartzinel, R. Y., Hogan, A., Brubaker, L., et al. (2017). Structural variants in genes associated with human Williams-Beuren syndrome underlie stereotypical hypersociability in domestic dogs. Science Advances 3, e1700398. doi:10.1126/sciadv.1700398.

206. Vonk, R., van der Schot, A. C., van Baal, G. C. M., van Oel, C. J., Nolen, W. A., and Kahn, R. S. (2014). Dermatoglyphics in relation to brain volumes in twins concordant and discordant for bipolar disorder. European Neuropsychopharmacology 24, 1885–1895. doi:10.1016/j.euroneuro.2014.09.010.

207. Vranjkovic O, Gasser PJ, Gerndt CH, Baker DA, Mantsch JR (2014) Stress-induced cocaine seeking requires a beta-2 adrenergic receptor-regulated pathway from the ventral bed nucleus of the stria terminalis that regulates CRF actions in the ventral tegmental area. J Neurosci. 34(37):12504–14. doi: 10.1523/JNEUROSCI.0680-14.2014.

208. Wang X, Pipes L, Trut LN, Herbeck Y, Vladimirova AV, Gulevich RG, Kharlamova AV, Johnson JL, Acland GM, Kukekova AV, Clark AG (2018) Genomic responses to selection for tame/aggressive behaviors in the silver fox (Vulpes vulpes). Proc Natl Acad Sci U S A. 115(41):10398–10403. doi: 10.1073/pnas.1800889115.

209. Watson, S., Gallagher, P., Ritchie, J. C., Ferrier, I. N., and Young, A. H. (2004). Hypothalamic-pituitary-adrenal axis function in patients with bipolar disorder. Br J Psychiatry 184, 496–502. doi:10.1192/bjp.184.6.496.

210. Weaver AM (2008) Cortactin in tumor invasiveness. Cancer Lett 265(2):157–66. doi: 10.1016/j.canlet.2008.02.066.

211. Whalley HC, Papmeyer M, Sprooten E, Lawrie SM, Sussmann JE, McIntosh AM (2012) Review of functional magnetic resonance imaging studies comparing bipolar disorder and schizophrenia. Bipolar Disord. 14(4):411–31. doi: 10.1111/j.1399-5618.2012.01016.x.

212. Wilkins, A. S., Wrangham, R. W., and Fitch, W. T. (2014). The “Domestication Syndrome” in Mammals: A Unified Explanation Based on Neural Crest Cell Behavior and Genetics. Genetics 197, 795–808. doi:10.1534/genetics.114.165423.

213. Williams, K. E., Marsh, W. K., and Rasgon, N. L. (2007). Mood disorders and fertility in women: a critical review of the literature and implications for future research. Hum Reprod Update 13, 607–616. doi:10.1093/humupd/dmm019.

214. Wójciak P, Remlinger-Molenda A, Rybakowski J (2012) The role of oxytocin and vasopressin in central nervous system activity and mental disorders. Psychiatr Pol. 46(6):1043–52.

215. Woodbury-Smith, M., Bilder, D. A., Morgan, J., Jerominski, L., Darlington, T., Dyer, T., et al. (2017). Combined genome-wide linkage and targeted association analysis of head circumference in autism spectrum disorder families. J Neurodev Disord 9. doi:10.1186/s11689-017-9187-8.

216. Wooderson, S. C., Gallagher, P., Watson, S., and Young, A. H. (2015). An exploration of testosterone levels in patients with bipolar disorder. BJPsych Open 1, 136–138. doi:10.1192/bjpo.bp.115.001008.

217. Yamaguchi Y, Lee YA, Goto Y (2015) Dopamine in socioecological and evolutionary perspectives: implications for psychiatric disorders. Front Neurosci. 9:219. doi: 10.3389/fnins.2015.00219.

218. Yurgelun-Todd, D. A., Gruber, S. A., Kanayama, G., Killgore, W. D., Baird, A. A., and Young, A. D. (2000). fMRI during affect discrimination in bipolar affective disorder. Bipolar Disord 2, 237–248.

219. Zerouni, C., Kummerow, E., Martinez, M., Diaz, A., Ezequiel, U., and Wix-Ramos, R. (2013). Affective disorder and hyperandrogenism. Recent Pat Endocr Metab Immune Drug Discov 7, 77–79.

220. Zipser B, Schleking A, Kaiser S, Sachser N (2014) Effects of domestication on biobehavioural profiles: a comparison of domestic guinea pigs and wild cavies from early to late adolescence. Front Zool. 11(1):30. doi: 10.1186/1742-9994-11-30.

221. Zollikofer CPE, Ponce de León MS (2010) The evolution of hominin ontogenies. Semin. Cell. Dev. Biol. 21, 441–452.

