## Supplemental file 2 for "Abnormal features of human self-domestication in bipolar disorder"

### Functional characterization of common candidates for BD and domestication

#### ADAMTSL3

Calling to mind the changes in body size and vertebral number in domestication (Teletchea, 2019), the extracellular matrix metalloprotease-encoding gene *ADAMTSL3* is associated with human height (Weedon et al., 2008) and is a noted selection signature involving body size in domestic pigs (Wilkinson et al., 2013). In partial tetrasomy 15q—a rare condition entailing autism-like developmental delay and learning/language deficits as well as retrognathia, flattened nasal bridge, and hyperpigmentation (Akahoshi et al., 2004; Tatton-Brown et al., 2009)—overexpression of *ADAMTSL3* is thought to be central to associated cardiovascular anomalies (George-Abraham et al., 2012), especially as subsequent patients with duplications that did not include the gene were found not to have cardiac defects (Levy et al., 2012; Xu et al., 2014). *ADAMTSL3* has been proposed to be a schizophrenia candidate gene after sequencing of patients with schizophrenia (SZ) and ultra-high density association analysis (Dow et al., 2011), noting that the gene is widely expressed in the brain throughout the lifespan. Additionally, genome-wide association study (GWAS) analyses by Pardiñas et al. (2018) and by the Schizophrenia Working Group of the Psychiatric Genomics Consortium (Ripke et al., 2014) have found *ADAMTSL3* to be associated with SZ. Single-nucleotide polymorphisms (SNPs) in the gene were found most strongly associated with SZ by Need et al. (2009).

#### ADRB2

Adrenoceptor Beta 2 (*ADRB2*) encodes the β-2 adrenergic receptor, which has a high binding affinity for epinephrine and which plays a pivotal role in the regulation of the cardiac, pulmonary, vascular, endocrine, and central nervous systems: not surprisingly, genetic polymorphisms in the gene have been related to several clinical conditions, including asthma, diabetes, obesity, and hypertension (Litonjua et al., 2010). *ADRB2* is highly expressed in the hippocampus and the cortex (Daly and McGrath, 2011), and specifically, in dendritic spines, which suggests that the gene might be involved in memory formation (Joiner, et al., 2010; Insel, 2011). A notable ultimate effector of this gene is the calcium channel $Ca_V1.2$, encoded in part by key bipolar disorder (BD) and domestication candidate gene *CACNA1C* (see below) (PubChem). It has also been shown to interact with the oxytocin receptor (encoded by *OTR*, a candidate gene of both autism spectrum disorder (ASD) and SZ) (Wrzal et al., 2012). There is moderate evidence for a genetic association between *ADRB2* and ASD in the AGRE cohort (Cheslack-Postava et al., 2007) and a study of dizygotic twins by Connors et al. (2005). A translocation with breakpoint *ADRB2* has also been reported with ASD (Vincent et al., 2009). There is also evidence that it modulates memory, particularly fear memories (e.g., in the context of posttraumatic stress disorder (PTSD)) (Johansen et al., 2011; Shi et al., 2018). Other conditions associated with mutations in this gene include nocturnal asthma and obesity (PubChem). Indeed, in dogs, *ADRB2* is hypothesized to have driven shifts of lipid metabolism during dietary changes co-occurring with domestication (Freedman et al., 2016).

#### ASTN1, ASTN2

*ASTN1* encodes astrotactin, an integral membrane protein expressed on neurons and facilitating their glial-guided migration (required for laminar assembly in the development of cortical regions), while ASTN2 complexes with ASTN1 during this process and regulates surface expression of ASTN1 (Wilson et al., 2010). In mammals, ASTN2 regulates the orientation of hair follicles, a feature altered in domesticates (Chang et al., 2015). *ASTN1* is associated with BD (Gratacòs et al., 2009), SZ (Kähler et al., 2008), and alcohol dependence (Hill et al., 2012). *ASTN2* has been implicated in ASD, speech delay, attention deficit hyperactivity disorder (ADHD), intellectual disability (ID), and other neurodevelopmental disorders (Lionel et al., 2014; Anazi et al., 2017; Autism Spectrum Disorders Working Group of The Psychiatric Genomics Consortium, 2017), with mutations also associated with decreased hippocampal volume and Alzheimer disease (AD) (Wang et al., 2015a; Hibar et al., 2017). Disruption of *ASTN2* is also linked to SZ (Vrijenhoek et al., 2008), with an adjacent SNP associated with both BD and SZ (Wang et al., 2010). *ASTN1* is positively selected in the domestic horse, while *ASTN2* is positively selected in domestic cattle (Theofanopoulou et al., 2017).

**ATXN7L1**

Ataxin 7-like 1, encoded by *ATXN7L1*, is related to Ataxin 7—associated with spinocerebellar ataxia type 7—though it is poorly characterized itself (Gamage et al., 2013). A microdeletion including *ATXN7L1* was reported in a case of a child with language delay, macrocephaly and overgrowth, prognathism, large ears, and ID (Uliana et al., 2010). A de novo inversion disrupting *ATXN7L1* in addition to *PDE1C* was also reported in a child with language and motor delay, ID, and near-macrocephaly (Gamage et al., 2013). The gene has also been linked to late-onset Alzheimer disease (Cummings et al., 2012). Positively selected in both cat and dog, *ATXN7L1* is one of fifteen genes selected in two or more domesticated species (Montague et al., 2014; Theofanopoulou et al., 2017) and one of four candidate genes in which SNPs distinguish free-breeding dogs and pure-breed dogs (Pilot et al., 2016).

**CACNA1C**

A key candidate gene for BD—as well as SZ, ASD, and domestication—is *CACNA1C*, which encodes the alpha 1C subunit of the L-type calcium channel, $Ca_V1.2$ (Ferreira et al., 2008; Green et al., 2010; Curtis et al., 2011; Niego and Benítez-Burraco, 2019). CACNA1C is required for hippocampal neurogenesis (Völkening et al., 2017), which is increased in foxes selected for tameness (Huang et al., 2015). Variants of the gene associate to deficits in reversal learning, as seen in BD and SZ (Sykes et al., 2019). *CACNA1C* expression is also modulated during associative learning (Sykes et al., 2018). Additionally, embryonic deletion of the gene in mouse glutamatergic neurons causes defects in synaptic plasticity, sociability, and cognition (Dedic et al., 2018).

*CACNA1C* gain-of-function mutation in humans causes Timothy syndrome (TS), which is characterized by QT prolongation and syndactyly as well as common findings of small and widely spaced teeth, autism, language and social delays, hypertrophic cardiomyopathy, baldness for the first two years of life, and craniofacial anomalies including low-set ears, small upper jaw, and flat nasal bridge (Splawski et al., 2004; Napolitano et al., 2015). A TS patient with a rare *CACNA1C* mutation who survived beyond childhood was found to have BD with psychotic features, with ten manic episodes by the age of 30 (Gershon et al., 2014). Meanwhile, haploinsufficiency in *CACNA1C* has been reported in a case of expressive language impairment/delay with anteverted nares and motor-skills delay; speech delay, language impairment, learning difficulties, microcephaly, facial dysmorphisms, and ID characterize related deletions (Mio et al., 2020). Haploinsufficiency in the related *CACNA1A* causes ASD and/or ADHD as well as executive dysfunction, hyperactivity-impulsivity, ID, and childhood epilepsy (Damaj et al., 2015).

In the five-disorder GWAS analysis, *CACNA1C* and another L-type calcium channel, *CACNB2*, were found to be associated to BD, ASD, SZ, ADHD, and SZ, suggesting a pleiotropic effect (Cross-Disorder Group of the Psychiatric Genomics Consortium, 2013). In BD and SZ, *CACNA1C* brain expression has shown to be decreased in postmortem studies (Gershon et al., 2014; Kabir et al., 2016).

*CACNA1C*—especially the intronic SNP within it, rs1006737—has been associated to BD by GWAS analysis (Ferreira et al., 2008; Liu et al., 2011; Sklar et al., 2011), association (Zhang et al., 2013), and case-control (Bigos et al., 2010) studies, as has hypermethylation of the gene (Starnawska et al., 2016). The risk SNP predicted increased prefrontal cortex (PFC) activity in executive cognition and increased hippocampus activity in emotional processing (including an emotional faces task) in healthy participants and was associated with increased *CACNA1C* expression in the human brain (Bigos et al., 2010). It is also associated with decreased semantic verbal fluency in healthy subjects (Krug et al., 2010) and decreased executive function in BD (Soeiro-de-Souza et al., 2013). Variants in *CACNA1C* also correlate with BDNF levels in BD (Smedler et al., 2019).

*CACNA1C* has been associated with SZ in GWAS analyses by Pardiñas et al. (2018), the Schizophrenia Working Group of the Psychiatric Genomics Consortium (Ripke et al., 2014), and the Schizophrenia Psychiatric GWAS Consortium (2011). The *CACNA1C* risk allele has been associated with white matter abnormalities (Mallas et al., 2017) and amygdala volume differences (Wolf et al., 2014) in SZ, as well as amygdala activation in BD (Tesli et al., 2013), calling to mind fear-related changes of domestication (see for example Brusini et al., 2018).

*CACNA1C* is considered a strong candidate for ASD based upon syndromic autism in Timothy syndrome as well as whole-genome sequencing (Jiang et al., 2013), whole-exome sequencing (Iossifov et al., 2014), targeted sequence enrichment (Brett et al., 2014), and targeted deep sequencing of postmortem ASD brains (D'Gama et al., 2015). In rodents, haploinsufficiency in the gene results in deficits in prosocial ultrasonic vocalization (Kisko et al., 2018; Redecker et al., 2019) and other social deficits that were rescuable with eIF2α inhibition, suggesting excitation/inhibition imbalance in the etiology of BD and SZ (Kabir et al., 2017).

This domestication candidate gene has been under positive selection in cattle (Qanbari et al., 2014; Theofanopoulou et al., 2017) and is hypermethylated in anatomically modern humans (AMH) relative to Neanderthals (Murphy and Benítez-Burraco, 2018). Work by Ramachandran et al. (2013) has demonstrated that $Ca_V1.2$ channels, while not directly affecting cranial neural crest migration, are expressed in neural crest-derived chondrocytes of the mandible, regulating mandibular development. (A case report of a patient with BD and periodic paralysis describes micrognathia in addition to flat nasal bridge, low-set ears, and malformed auricular cartilage (Raveendranathan et al., 2012).)

### CACNA1D

The 1D subunit of the L-type calcium channel, $Ca_V1.3$, is encoded by *CACNA1D*, which is widely expressed in the neural crest-derived chromaffin cells of the adrenal gland (Comunanza et al., 2010; Marcantoni et al., 2010; Vandael et al., 2015), the hypothesized indirect target of selection in domestication (Wilkins et al., 2014). *CACNA1D* is also expressed in the central nervous system (CNS)—especially the hippocampus, where it is required for proper neurogenesis (Marschallinger et al., 2015)—with evidence for involvement in memory and learning (Berger and Bartsch, 2014; Liu et al., 2014b), including context-conditioned fear memory in mice (McKinney and Murphy, 2006). Loss of function of *CACNA1D* leads to the SANDD (sinoatrial node dysfunction and deafness) syndrome (Baig et al., 2011); the gene has also been associated with Parkinson disease (Berger and Bartsch, 2014), while a de novo *CACNA1D* mutation was identified in case of twins with language delay and self-injurious behavior (Hofer et al., 2020).

The gene plays an important role in neuropsychiatric pathology, with one SNP in *CACNA1D* linked to BD, SZ, ASD, major depressive disorder (MDD), and ADHD in the five-disorder GWAS (Cross-Disorder Group of the Psychiatric Genomics Consortium, 2013). *CACNA1D* is a strong candidate gene for ASD, with de novo gain-of-function mutations linked to sporadic ASD (Iossifov et al., 2012; O'Roak et al., 2012; Pinggera et al., 2015, 2017; Kabir et al., 2016; Hofer et al., 2020). Self-biting behavior, as seen in ASD and neurogenetic disorders such as Lesch-Nyhan syndrome, has been shown to be induced in mice by agonism of L-type calcium channels (Jinnah et al., 1999; Kasim and Jinnah, 2003), while activation of $Ca_V1.3$ channels in the ventral tegmental area (VTA) induced deficits in social behavior (Martínez-Rivera et al., 2017). SNPs have been associated with BD (Ament et al., 2015; Kabir et al., 2016; Ross et al., 2016), and L-type calcium channel antagonists have been used in the treatment of BD (Cipriani et al., 2016). *CACNA1D* has also been associated with SZ in the GWAS analysis by Pardiñas et al. (2018).

Compared to prehistoric ancestors, *CACNA1D* has been positively selected in domestic horses (Schubert et al., 2014) and AMH (Racimo, 2016; Theofanopoulou et al., 2017). As noted by Theofanopoulou et al. (2017), it is also selected in vocal learning avians (Nam et al., 2010; Friedrich et al., 2019).

**CAGE1**
Cancer-associated gene-1 (*CAGE1*) is expressed in testis normally, but also in a variety of cancer tissues (at higher levels) (Park et al., 2003). It was one of six genes deleted in two patients who shared facial dysmorphisms including low-set and deformed ears, short philtrum, and prominent forehead in addition to early feeding difficulty and ID; short nose, micrognathia/retrognathia, heart defects, and hair abnormalities were also reported (Kuipers et al., 2013). *CAGE1* is positively selected in the domestic cat (Montague et al., 2014) and is differentially expressed in the prefrontal cortices of tame versus aggressive silver foxes, subjects of the renowned fox farm domestication experiment (Wang et al., 2018).

**CD36**
*CD36* encodes a protein alternately known as glycoprotein IV (GPIV) that functions as a scavenger receptor for fatty acid translocation and lipid metabolism regulation in the brain (Abumrad et al., 2005). In the brain, it is expressed at especially high levels in the amygdala and hypothalamus (Zapata et al., 2016). *CD36* is also reported as having a role in learning ability (Abumrad et al., 2005; Moullé et al., 2012; Niego and Benítez-Burraco, 2019). It has been proposed as a therapeutic target in the treatment of BD, inasmuch as valproate for BD decreases production of arachidonoyl-CoA, and given polyunsaturated fatty acid deficiencies in BD, SZ, and other neuropsychiatric conditions (Iwayama et al., 2010; Moullé et al., 2012). *CD36* was one of several domestication candidate genes found to be upregulated in the blood of patients with Williams syndrome, another condition hypothesized to be related to the domestication phenotype (Niego and Benítez-Burraco, 2019). The gene is also thought to have a role in changes in canine fear and aggression during domestication (Zapata et al., 2016), consistent with increased anxiety and aggression in *CD36*-knockout mice (Zhang et al., 2016).

**CNTN6**
*CNTN6* encodes Contactin 6, a neural cell adhesion molecule with roles in synaptogenesis (especially in the cerebellum) and dendrite orientation, that also acts as a NOTCH1 ligand (Mercati et al., 2017). It is associated with BD (Kerner et al., 2011), ASD as a candidate with strong evidence (Firth et al., 2009; Roohi et al., 2009; van Daalen et al., 2011; Iossifov et al., 2014; Poot, 2014; Juan-Perez et al., 2018, 6) including auditory perception abnormalities in ASD (Mercati et al., 2017), SZ (Juan-Perez et al., 2018), ID (Kashevarova et al., 2014), and other neurodevelopmental disorders (Oguro-Ando et al., 2017, though see Repnikova et al., 2020). Additionally, *CNTN6* duplications substantially increase risk of Tourette syndrome (Huang et al., 2017).

Findings reported with single-gene *CNTN6* copy number variations (CNVs) include ID with speech/language delay, microcephaly/dolichocephaly, small ears with malformations, flat nasal bridge, small nose, anteverted nostrils, short philtrum, teeth abnormalities, and micrognathia (Kashevarova et al., 2014). Macrocephaly, brachycephaly, short stature, feeding difficulties—with personal and family histories of ASD, BD, SZ, ADHD, and learning disability—have also been reported (Hu et al., 2015).

*CNTN6* is positively selected in the domestic horse (Schubert et al., 2014; Theofanopoulou et al., 2017) and is a candidate gene for the Persian cat breed phenotype, the extreme brachycephaly and small ears of which may represent an ‘exaggeration’ of the domestication syndrome (Bertolini et al., 2016).

**COBL**
COBL is an actin nucleator, preferentially expressed in the brain, that is involved in inducing neurites and neurite branching (Ahuja et al., 2007). It is a candidate gene for ASD (Griswold et al., 2012) and is among the genes duplicated in the imprinting disorder Silver-Russell syndrome, characterized by a small, triangular face, macrocephaly relative to a growth-restricted body, and ear anomalies (Hitchins et al., 2002; Spiteri et al., 2017). *COBL* is also enriched in neural tube defects (Ishida et al., 2018). The gene is positively selected in cattle (Theofanopoulou et al., 2017).

**CRH**

*CRH* encodes corticotropin-releasing hormone, which stimulates adrenocorticotropic hormone (ACTH) production by the pituitary gland. Aside from its role in the hypothalamic-pituitary-adrenal (HPA) axis, it also acts as a neurotransmitter with roles in, for instance, the storage of fear memories in the hippocampus (Chen et al., 2012c). CRH is also thought to affect dopamine release, suggesting a role in stress-induced substance relapse (Payer et al., 2017). BD is marked by HPA axis dysfunction; e.g., the cortisol response to the dexamethasone-suppressed/CRH test in patients with BD is enhanced compared to controls (Watson et al., 2004), and CRH levels are decreased in patients with BD relative to controls (Monfrim et al., 2014). CRH levels are also decreased in children with ASD and in Bull Terrier dogs with an ASD-like phenotype (Tsilioni et al., 2014). CRH release, in conjunction with high circulating glucocorticoid levels, is implicated in stress-induced regression of dendritic spines, potentially precipitating depression or psychosis, especially in adolescence (Bennett, 2008). Epigenetic modification of *CRH* correlates with severity of suicide behavior and general psychiatric risk (Jokinen et al., 2017). Indeed, CRH abnormalities are reported in PTSD (Heim et al., 1997; Baker et al., 1999), suicide (Fawcett et al., 1997), and both postpartum and endogenous depression (Young et al., 1990; Bloch et al., 2005). In domestication, which is hypothesized as a selection for decreased HPA axis reactivity via mild neurocristopathy (Wilkins et al., 2014), CRH is decreased in the hypothalamus of foxes selected for domestication (Trut et al., 2009). In addition, glutamate receptors, which are highly selected in domestication, can mediate CRH release and thereby modulate the HPA axis stress response (O'Rourke and Boeckx, 2020).

**CTTN**

*CTTN* encodes cortactin, an inducer of actin polymerization and activator of the actin-related protein (Arp)2/3 complex, which is important for formation and stabilization of branched actin networks (Daly, 2004). A cell-adhesion protein, it is involved in the regulation of adherens-type junctions and cytoskeletal organization, and ultimately in controlling cell shape and cell migration (Daly, 2004; see Weaver, 2008 for review). It is highly expressed in dendritic spines, especially in the hippocampus, and interacts with Shank proteins, thought to be important in ASD (Hering and Sheng, 2003; Ueda et al., 2013), as well as the ASD candidate gene *DIP2A* (Ma et al., 2019b). In collaboration with components of the postsynaptic density, like Shank proteins, which stabilize it at the synapse, cortactin contributes to regulate synapse morphology and function (MacGillavry et al., 2016). Likewise, cortactin is involved as well in dendritic spine formation (Hering and Sheng, 2003; Gray et al., 2005; Ueda et al., 2013) and axon outgrowth (Kubo et al. 2015), with an impact on morphological changes in the neuron associated to brain activity, like long-term potentiation (Racz and Weinberg, 2004). Cortactin also regulates dendritic spine formation via interaction with DOCK4, encoded by *DOCK4*, a candidate gene in ASD and SZ, consistent with dendritic spine pathology characterizing these and other neuropsychiatric disorders (Penzes et al., 2011; Ueda et al., 2013). *CTTN* is downregulated in the brain of people with schizophrenia (Bhambhvani et al., 2016). Cortactin is also involved in the recycling of the beta-2-adrenergic receptor, encoded by *ADRB2* (Vistein and Puthenveedu, 2014). It is also involved in the proliferation and invasiveness of many cancers, including glioma (Weaver, 2008; Su et al., 2018; Ramos-García et al., 2019). *CTTN* is positively selected in cats (Montague et al., 2014; Theofanopoulou et al., 2017).

**CUL1**

*CUL1* encodes a scaffolding protein that interacts with SKP1 and F-box protein to form the SCF ubiquitin ligase complex, which targets cell-cycle proteins and is involved in the regulation of the Hedgehog, Wnt, and NF-κB pathways (Maniatis, 1999). *CUL1* expression is increased in glioma cells and other cancers and plays a role in their migration and spread (Fan et al., 2014; Chen et al., 2016; Ren et al., 2019). In a study of patients with Weaver syndrome, a patient with deletion of *CUL1* in addition to *EZH2* (usually causative in the syndrome) exhibited severe ID, overgrowth, macrocephaly, retrognathia, large ears, low nasal bridge, long philtrum, and cardiac anomalies (Imagawa et al., 2017). *CUL1* is positively selected in domestic cattle.

**CUX2**

The cut homeodomain transcription factor CUX2 has been proposed to regulate expression of neural cell adhesion molecules and is involved in dendritic spine formation and synaptogenesis in the upper cortical layers of the brain (Jacobsen et al., 2001; Cubelos et al., 2010). Indeed, failed interaction of *CUX2* with *LHX2* is hypothesized to be associated with ASD and SZ (Yang et al., 2020a). *CUX2* is also involved in hippocampal neurogenesis (Yamada et al., 2015). It has been considered as a candidate gene in BD (Jacobsen et al., 2001; Glaser et al., 2005; Kato, 2007), and there is suggestive evidence for *CUX2* as a candidate in ASD from whole-exome sequencing (De Rubeis et al., 2014) and cases of patients with ASD, ID, and epilepsy in which *CUX2* variants are identified (Barington et al., 2018; Chatron et al., 2018). Autistic features, inappropriate laughter, and non-verbal status have also been reported in these cases (Chatron et al., 2018). It is positively selected in dogs (Freedman et al., 2016; Theofanopoulou et al., 2017).

**DCC**

Deleted in colorectal cancer (DCC) is a receptor for netrin-1 and plays a key role in axon guidance during brain development (Finci et al., 2015; Kang et al., 2018), specifically projection of thalamocortical axons from the dorsal thalamus to the cortex (Braisted et al., 2000). It interacts with the SLIT-ROBO pathway, which has been implicated in vocal learning, as well as with the domestication candidate and neural crest cell gene, *DSCAM* (Wang et al., 2015b; Theofanopoulou et al., 2017). *DCC*-deficient mice have increased basal dopamine levels and altered dendritic spine density in neurons of the medial prefrontal cortex (mPFC), which were associated with altered reward responses (Grant et al., 2007). DCC has been proposed to regulate mPFC connectivity during adolescence by giving rise to cognitive flexibility through anatomical and electrophysiological changes, to affect dopaminergic innervation in the mPFC (Manitt et al., 2013), and to be associated with adolescent emergence of SZ and other psychiatric conditions (Vosberg et al., 2020). *DCC* has been associated with mood instability (Ward et al., 2017), SZ (Grant et al., 2012), behaviors seen in depression (Torres-Berrío et al., 2017), and ASD (Nisar et al., 2019), and is involved in mirror movement disorder (Vosberg et al., 2019). *DCC* is a neural crest-related gene positively selected in both cat and horse (Montague et al., 2014; Schubert et al., 2014), and a high sequence conservation region deleted in AMH (hCONDEL) has been identified upstream of *DCC*, though it is common to Neanderthals (McLean et al., 2011; Theofanopoulou et al., 2017).

**DDC**

*DDC* encodes DOPA decarboxylase, which converts L-DOPA to dopamine (itself the precursor to adrenaline and noradrenaline) and 5-hydroxytryptophan to serotonin (Lauritsen et al., 2002). *DDC* is thought to be a minor susceptibility gene for BD (Børglum et al., 1999; Jahnes et al., 2002), especially when paternally transmitted (Børglum et al., 2003). Indeed, mania is thought to arise from a hyperdopaminergic state (Ashok et al., 2017). There is suggestive evidence for *DDC* as a candidate for ASD (Toma et al., 2013 (though see Lauritsen et al., 2002)). It is also associated with ADHD (Ribasés et al., 2009). While *DDC* polymorphism is reported not to be associated with SZ (Zhang et al., 2004), age of onset of SZ appears to be correlated with *DDC* genotype (Børglum et al., 2001). It is positively selected in the domestic dog (Cagan and Blass, 2016) and upregulated in the striatum of AMH compared to apes (Calvey, 2019).

**DSCAM**

*DSCAM* encodes the Down syndrome (DS) cell adhesion molecule, a protein with diverse functions in neurodevelopment including dendrite arborization (Maynard and Stein, 2012), inhibition of synaptic plasticity (Simmons et al., 2017), and axon fasciculation (Bruce et al., 2017). It is expressed in the developing cortex, hippocampus, and medulla as well as the neural crest and its derivatives (Yamakawa et al., 1998; Barlow et al., 2002).

Given expression of *DSCAM* in the neural crest and its status as a candidate gene for defects in Down syndrome (Barlow et al., 2002), it is worth noting that trisomy 21 causes a neurocristopathy in mice, with deficits in neural crest cell migration, mitosis, and delamination (Roper et al., 2009; Deitz, 2013), potentially suggesting that the craniofacial, dental, cardiac, and gastrointestinal abnormalities seen in DS —including an association with Hirschsprung disease (Friedmacher and Puri, 2013)—might be explained by underlying neurocristopathy.

Located near the BD linkage locus chromosome 21q22, *DSCAM* is linked to BD by association and postmortem brain expression studies with allele- and genotype-specific differences in expression levels (though overall expression was not significantly different between BD patients and controls) (Amano et al., 2008). In addition, there is strong evidence for *DSCAM* as a candidate gene for ASD, with several de novo loss-of-function variants identified in patients with ASD (Iossifov et al., 2014; Wang et al., 2016; Guo et al., 2018) and evidence from whole genome sequencing (Turner et al., 2016; C Yuen et al., 2017), GWAS analysis (Autism Spectrum Disorders Working Group of The Psychiatric Genomics Consortium, 2017), and targeted sequencing (Stessman et al., 2017). The gene is positively selected in domestic cattle (Qanbari et al., 2014; Theofanopoulou et al., 2017)

**ERBB4**

*ERBB4* encodes the receptor for neuregulins—especially the SZ candidate neuregulin1 (NRG1) (Yang et al., 2003; Munafò et al., 2006)—important for neural (particularly, neural crest-derived cranial ganglia) and cardiac development (Lee et al., 1995). ERBB4/NRG1 signaling has roles in myelination, synaptogenesis, and synaptic plasticity (Mei and Nave, 2014), regulating, for example, migration of GABAergic interneurons from ganglionic eminences to the cortex (Li et al., 2012). ERBB4/NRG1 signaling is important for oligodendrocyte development, with loss of ERBB4 signaling leading to myelin abnormalities similar to those seen in SZ and BD and analogous behaviors in the deficient mice (Roy et al., 2007) (see *MBP* below). *ERBB4* mutation is associated with amyotrophic lateral sclerosis (ALS) (Takahashi et al., 2013) as well as ID and speech delay (Kasnauskiene et al., 2013). Downregulation of hippocampal ERBB4/NRG1 signaling is important in learning and memory (Tian et al., 2017), with ERBB4 inhibition rescuing hippocampal long-term potentiation impairments and fear memory deficits in a mouse model of Angelman syndrome (AS) (Kaphzan et al., 2012). *ERBB4* and *NRG1* are also associated with social behavior, with mice deficient in these genes spending more time with unfamiliar mice, calling to mind the hypersociability seen in AS and Williams syndrome (WS) (Moy et al., 2009; Toth, 2019). On the other hand, *NRG1*-heterozygous mice exhibit decreased social approach, as in ASD (Ehrlichman et al., 2009). Glutamatergic impairment via *NRG1* overexpression in a mouse model required *LIMK1*—a gene deleted in WS and duplicated in some cases of ASD (Yin et al., 2013; Mei and Nave, 2014).

*ERBB4* is associated with BD (Goes et al., 2011; Chen et al., 2012b), and deletion of *ERBB4* from noradrenergic neurons in the mouse locus coeruleus results in catecholamine elevation, giving rise to mania-like behaviors, rescuable with the mood stabilizer lithium, dopamine antagonism, or norepinephrine antagonism (Cao et al., 2018). Polymorphisms in *ERBB4* are associated with enhanced linguistic abilities, IQ scores, and memory in BD (Rolstad et al., 2015). *ERBB4* has been identified as a susceptibility gene for SZ by association (Sun et al., 2008a), linkage (Lewis et al., 2003; Ng et al., 2009), and expression (Silberberg et al., 2006) studies, and its role in SZ is thought to be related to its interactions in glutamatergic signaling (Banerjee et al., 2010) (see *GRIK3* below). *NRG1*, meanwhile, is associated with verbal fluency in SZ (Kircher et al., 2009), and ERBB4/NRG1 signaling enhances synchronized PFC oscillations, which are reduced in SZ (Hou et al., 2014; Murphy and Benítez-Burraco, 2017). Finally, *ERBB4* also interacts with *PTEN*, a candidate for a subtype of ASD characterized by macrocephaly and language delay with reduced processing speed and working memory (Naqvi et al., 2000; Buxbaum et al., 2007; Tilot et al., 2015).

*ERBB4* is positively selected in both AMH (Pickrell et al., 2009; Theofanopoulou et al., 2017) and domestic cattle, in which it is a candidate gene for coat color variation (Qanbari et al., 2014). Its ligand, *NRG2*, is selected in dog, cat, and cattle, the only gene under positive selection in three domesticates in the study by Theofanopoulou et al. (2017). *ERBB4*-deficient mice exhibit faulty pathfinding of hindbrain-derived cranial neural crest cells (Golding et al., 2000), suggesting a role in skull globularization (Murphy and Benítez-Burraco, 2017) and, relatedly, brain language-readiness in AMH (Boeckx and Benítez-Burraco, 2014; Benítez-Burraco et al., 2016).

**FAM107B**

*FAM107B* encodes the heat shock-inducible tumor small (HITS) protein and is widely expressed, especially in breast, thyroid, gonadal, and brain tissue (Nakajima et al., 2012). In the mouse brain, *FAM107B* is expressed in the hippocampus during embryogenesis, suggesting roles in neuronal migration and corticogenesis (Masana et al., 2015), though its function is yet to be fully understood. Expression is also decreased in several, especially gastrointestinal, cancers (Nakajima et al., 2012; Guo et al., 2017) and altered in spinocerebellar ataxia type 2 (Scoles et al., 2017). *FAM107B* has been linked to BD by GWAS analysis (Johnson et al., 2009; Nurnberger et al., 2014) and interacts with *FAM107A*, which is increased in the prefrontal cortex of patients with BD and SZ (Shao and Vawter, 2008) and located at a susceptibility locus of BD, SZ, and ASD (Nakajima and Koizumi, 2014). It has also been implicated in animal models of puerperal psychosis and (transgenerational) depression (Quilter et al., 2012; Zhang et al., 2017a). The gene is positively selected in the dog (Freedman et al., 2016; Theofanopoulou et al., 2017).

**GABRA5**

*GABRA5* encodes the α5 subunit of the type A receptor for the major inhibitory neurotransmitter in the brain, γ-aminobutyric acid (GABA), which is thought to be of prime importance in BD (Cotter et al., 2002; Brambilla et al., 2003) and ASD (Blatt et al., 2001; Hussman, 2001). It is primarily expressed in hippocampus (Sieghart and Sperk, 2002), and its expression is modulated by *ULK4*, CNVs of which are associated with ASD, SZ, and language delay, among other disorders (Liu et al., 2018). *GABRA5* reduction is associated with increased fear, and inhibition of the receptors reverses memory deficits in mice (Ponder et al., 2007; Wang et al., 2012; Li et al., 2014). Interestingly, *GABRA5* is among the genes deleted in the Prader-Willi/Angelman syndromes (PWS, AS), and it flanks a gene expressed in neural crest cells and associated with ASD and epilepsy, *GABRB3*, deletion of which causes ocular hypopigmentation as seen in PWS and AS, with *GABRA5* also downregulated with ocular hypopigmentation (Delahanty et al., 2016). Mutations in *GABRA5* are associated with early-onset epileptic encephalopathies (Hernandez et al., 2019), in which microcephaly, autistic behaviors, nonverbal status, and developmental delay have also been reported (Butler et al., 2018). *GABRA5* has been linked to BD by case-control and genetic association studies (Papadimitriou et al., 2001b; Otani et al., 2005; Kato, 2007), particularly with predominance of manic over depressive symptoms in BD (Papadimitriou et al., 1998). It is also associated with unipolar depression (Oruc et al., 1997) and with later age of onset of SZ (Papadimitriou et al., 2001a), with *GABRA5* modulation modifying dopaminergic activity in a mouse model of SZ (Gill et al., 2011). In mice, *GABRA5* deletion causes ASD-like behaviors including reductions in sociality and vocalization (Zurek et al., 2016) and ASD-like EEG and sleep-wake patterns (Mesbah-Oskui et al., 2017), consistent with associations to ASD in humans (Menold et al., 2001), particularly verbal communication deficits (Yang et al., 2017) (though see Mahdavi et al., 2018).

Through analyzing QTLs thought to underlie tameness in rats in a domestication experiment parallel to the farm fox study, *GABRA5* has been identified as a candidate gene affecting tameness (particularly fear) (Albert et al., 2011), consistent with prior findings of lower GABA levels in the brains of tame compared to aggressive rats (Albert et al., 2008). *GABRA5* and *TPH1* (see below) were the two candidate genes near regions of positive selection. The gene is positively selected in dogs (Li et al., 2014).

**GDNF**

Glial cell-derived neurotrophic factor, encoded by *GDNF*, facilitates differentiation and synaptogenesis of dopaminergic neurons (Christophersen et al., 2007; Ledda et al., 2007), and appears to have neuroprotective effects (Tajdaran et al., 2016; Al-Ayadhi et al., 2019) and promote neuronal survival (Coulpier et al., 2002).

In BD, SNPs in *GDNF* associate differently with patients' severity and functionality scores (Safari et al., 2017). Circulating GDNF levels in patients with BD have been found to vary with mood state and differ from those in healthy controls (Rosa et al., 2006; Barbosa et al., 2011). GDNF levels are higher in manic patients and lower in SZ patients relative to healthy controls (Tunca et al., 2015). While peripheral GDNF levels in ASD patients have not been seen to differ from controls (Rodrigues et al., 2014), they did rise with auditory integration therapy, with autistic behavior also declining in scale-rated severity (Al-Ayadhi et al., 2019). Meanwhile, high GDNF levels relative to controls are reported in untreated children with ADHD (Bilgiç et al., 2017).

Mutations in *GDNF* and *RET*—which encodes the receptor of GDNF—are the primary causes of Hirschsprung disease, in which neural crest cells of the enteric nervous system (ENS) fail to colonize the distal colon; normally, GDNF induces enteric neural crest cells to mature and acts as a chemoattractant during their migration (Schriemer et al., 2016). *GDNF* also has a minor role in pheochromocytoma, a tumor of neural crest-derived chromaffin cells (Dahia et al., 1997; Woodward et al., 1997; Powers et al., 2009). *GDNF* is under selection in domesticated Atlantic salmon (López et al., 2018) and is a key candidate gene proposed by Wilkins et al. (2014) as part of the neural crest-mediated hypothesis of domestication.

**GLRA1**
*GLRA1* encodes the glycine receptor, subunit α-1, a ligand-gated chloride channel expressed in the CNS (primarily spinal cord) that mediates postsynaptic inhibition (Mowrey et al., 2013). *GLRA1* mutation causes the rare familial disorder, hyperekplexia, characterized by an exaggerated startle response (Paucar et al., 2018, 1) and sometimes accompanied by speech delay, developmental delay, and learning difficulties (Thomas et al., 2013). *GLRA1* is downregulated in a mouse model of Angelman syndrome (Low and Chen, 2010), and polymorphism in the gene is associated with trait schizotypy (Vora et al., 2018). A related gene, *GLRA2*, is associated with ASD (Pilorge et al., 2016). *GLRA1* is positively selected in the domestic dog (Freedman et al., 2016; Theofanopoulou et al., 2017)

**GRIA1**
*GRIA1* encodes the glutamate receptor-1. Glutamate receptors are the primary excitatory neurotransmitter in mammals and are involved in long-term potentiation, with learning impaired in deficient mice (Mead and Stephens, 2003; Montague et al., 2014). *GRIA1* is associated with BD (Kerner et al., 2009), with *GRIA1* expression in neurons in the nucleus accumbens affecting mania-like behaviors in a mouse model (Parekh et al., 2018). It has been identified as a candidate gene for SZ in GWAS analyses (Ripke et al., 2014) and linkage studies (Lewis et al., 2003; Ng et al., 2009), with *GRIA1* found upregulated in the dorsolateral prefrontal cortices of patients with SZ (O'Connor and Hemby, 2007) though downregulated in hippocampus (Eastwood et al., 1995). It is also a strong candidate for ASD, with missense variants reported with ASD (De Rubeis et al., 2014; Iossifov et al., 2014) as well as in specific learning disabilities (Geisheker et al., 2017) and ID (de Ligt et al., 2012). *GRIA1* is positively selected in both cat and AMH and interacts with the domestication candidate *GRIK3* (Montague et al., 2014; Theofanopoulou et al., 2017).

**GRID1**
*GRID1* encodes the glutamate receptor δ-1 subunit that is important in synaptic plasticity in addition to excitatory synaptic transmission and presynaptic terminal differentiation (Kuroyanagi et al., 2009; Zhang et al., 2018a). It is also required for burst firing of dopaminergic neurons (Benamer et al., 2018). A 10-Hz parieto-occipital α rhythm, associated near the locus of *GRID1* (Salmela et al., 2016), has roles in lexical decision making and embedding of γ rhythms generated during language processing (Murphy, 2015), which is perturbed during lexical and sentence processing in SZ (Murphy and Benítez-Burraco, 2016).

*GRID1* has been associated with BD by case-parent trios and exome sequencing (Fallin et al., 2005; Zhang et al., 2018a). *GRID1*-knockout mice exhibit hyperactivity, aggression, social interaction deficits, depression-like behavior rescuable with lithium, and decreased anxiety-like behavior (Yadav et al., 2012).

Additionally, *GRID1* has been associated to SZ in linkage studies (Lewis et al., 2003; Ng et al., 2009) and association studies (Guo et al., 2007). It is also a strong candidate for ASD, with deletions encompassing the gene reported in ASD (Griswold et al., 2012) and overrepresented in ASD cases (Glessner et al., 2009). It is positively selected in domestic horse (Schubert et al., 2014; Theofanopoulou et al., 2017).

**GRIK2**

*GRIK2* encodes the ionotropic glutamate receptor kainite 2. Polymorphisms in *GRIK2* are associated with aggression during mania in patients with BD (Ma et al., 2019a), with *GRIK2*-knockout mice exhibiting aggression and decreased anxiety (Shaltiel et al., 2008). Strong evidence links *GRIK2* to ASD, within the Han Chinese (Shuang et al., 2004) and European (Holt et al., 2010) populations, by affected sib-pair study (Jamain et al., 2002) and with ASD-associated CNVs (Griswold et al., 2012). In SZ, maternal transmission disequilibrium of *GRIK2* has been shown (Bah et al., 2004), though no association was found in a case-control study (Shibata et al., 2002). However, *GRIK2* expression in the brains of patients with SZ is decreased (Meador-Woodruff et al., 2001; Bah et al., 2004). *GRIK2* has also been associated with obsessive-compulsive disorder (OCD) (Mattheisen et al., 2015) and nonsyndromic autosomal recessive mental retardation (Motazacker et al., 2007), while a gain-of-function mutation is reported in a case with language delay and ID (Guzmán et al., 2017). Selective sweep in *GRIK2* has been identified in rabbit domestication (Carneiro et al., 2014). Furthermore, *GRIK2* is expressed in higher levels in frontal cortices of dog compared to wolf and is upregulated in multiple domesticates relative to wild counterparts (Li et al., 2014). It is positively selected in dog, rabbit, and duck (O'Rourke and Boeckx, 2020)

**GRIK3**

*GRIK3* encodes inotropic glutamate receptor kainate 3. Maternal linkage and association of *GRIK3* has been associated with BD-II (but not BD-I) and MDD (Schiffer and Heinemann, 2007; Cherlyn et al., 2010). *GRIK3* has been determined a susceptibility gene for SZ in association studies (Begni et al., 2002; Allen et al., 2008; Sun et al., 2008a; Ahmad et al., 2009) and linkage studies (Lewis et al., 2003; Ng et al., 2009). Interestingly, serum levels of glutamate-related antibodies to the NMDA receptor and GAD are elevated in patients with acute mania relative to lithium-treated patients (Ferensztajn-Rochowiak et al., 2019), calling to mind the psychosis and mania that can be seen in anti-NMDA receptor encephalitis (Barry et al., 2015). A key domestication candidate, it has been positively selected in AMH, dog, cattle, and sheep, one of only five genes selected in both AMH and multiple domesticated species (Freedman et al., 2016; Li et al., 2014; Theofanopoulou et al., 2017; O'Rourke and Boeckx, 2020). As detailed by O'Rourke and Boeckx (2020), glutamate receptors modulate the stress response via the HPA axis, and various features of the domestication phenotype could be attributable to changes in glutamatergic receptors during domestication, noting that glutamate receptors are expressed on melanocytes (Hoogduijn et al., 2006) and that glutamatergic signaling is important in the regulation of oligodendrocytes and (especially learning-dependent) myelination (Gautier et al., 2015; Spitzer et al., 2016).

**HS3ST4**

*HS3ST4* encodes heparan sulfate-glucosamine 3-sulfotransferase 4, which is enriched in the hippocampus, thought to be involved in neurotransmitter receptor activity and synaptic transmission, and involved in the pathogenesis of Herpes simplex virus-1 (Tiwari et al., 2005; Debette et al., 2015). It is involved in the production of heparan sulfate, which has a role in blood coagulation (Reynolds et al., 2019). It has been associated to BD by CNV and GWA studies (Johnson et al., 2009; Grozeva et al., 2010). *HS3ST4* was one of two genome-wide associations in a study of verbal declarative memory in nondemented older adults (Debette et al., 2015). It is positively selected in cattle (Qanbari et al., 2014; Theofanopoulou et al., 2017).

**HTR4**

*HTR4* encodes 5-hydroxytryptamine receptor 4, part of the serotonin receptor family, expressed among other places in the CNS (especially basal ganglia and hippocampus), heart, and adrenal glands (Hegde and Eglen, 1996; Varnäs et al., 2003), in which location overexpression can be associated with ACTHindependent bilateral macronodular adrenal hyperplasias (AIMAH) (Cartier et al., 2003), a cause of psychosis (Shah et al., 2019). Polymorphisms in *HTR4* are associated with BD (Ohtsuki et al., 2002). The gene has also been identified as a susceptibility gene for SZ in linkage studies (Lewis et al., 2003; Ng et al., 2009), with a particular SZ-susceptibility haplotype (Suzuki et al., 2003). Hypomethylation of the *HTR4* promoter is associated with ASD in males (Hu et al., 2020), and a translocation with breakpoint near *HTR4* and *ADRB2* is reported in a case of ASD (Vincent et al., 2009). *HTR4* is selected in pig and dog domestication (Li et al., 2013; Leno-Colorado et al., 2017; Theofanopoulou et al., 2017). Mutations in *HTR2A* have been found to result in a dysregulation of HPA axis and serotonergic system (Smith et al., 2008).

**IGF1**

*IGF1* encodes insulin-like growth factor 1, a protein that regulates growth and development and is important for brain growth and regulation of myelination (Carson et al., 1993). Within the adult brain, IGF-1 is expressed in the hippocampus (Kar et al., 1993) and involved in synaptic plasticity, neuronal excitability, and gap junctional communication (Fernandez et al., 2007). It alleviates neuroinflammation and NMDA-induced neurotoxicity (Riikonen, 2016).

*IGF1* has been identified as a candidate gene for BD by GWAS (Pereira et al., 2011). Peripheral IGF-1 levels are increased in patients with BD (Kim et al., 2013; da Silva et al., 2017)—including during and after acute mania, in what is hypothesized to be a compensation to protect against excitotoxicity (Liu et al., 2014a; Ferensztajn-Rochowiak et al., 2019)—and correlate with lithium response (Squassina et al., 2013).

IGF-1 levels are also increased in patients in the initial stages of SZ (Palomino et al., 2013, in a study that did not find statistically signficant elevation in patients with BD), though IGF-1 levels are found to be decreased in antipsychotic-naïve patients several years after the onset of SZ (Venkatasubramanian et al., 2007), which may be implicated in the increased insulin resistance of even antipsychotic-naïve, first-episode patients with SZ (Ryan et al., 2003) and perhaps the body size differences in SZ hypothesized to be mediated by genomic imprinting effects (Crespi and Badcock, 2008; Benítez-Burraco et al., 2017).

Given decreased IGF-1 levels in cerebrospinal fluid (CSF) of children with ASD compared to controls, as well as thinner myelin layers in the brains of those with ASD, Steinman and Mankuta (2019) have suggested that IGF-1 may ameliorate symptoms of ASD; indeed, it has been used to treat Rett syndrome (Pini et al., 2014). IGF-1 levels are also low in Williams syndrome, consistent with growth restriction across all ages (Levy-Shraga et al., 2018).

The gene is positively selected in AMH (Racimo, 2016; Theofanopoulou et al., 2017) and domestic cat (Montague et al., 2014). *IGF1* is a major determinant of body size in dogs, with a single SNP haplotype common to all small breeds (Sutter et al., 2007; Wayne and vonHoldt, 2012); while the mutation appears to post-date domestication of the dog, it is thought to have arisen early in dog domestication (Gray et al., 2010). *IGF1* expression is increased or has been under selection in several domesticates, including chicken (Beccavin et al., 2001), trout (Devlin et al., 2009), and salmon (Tymchuk et al., 2009).

**IMMP2L**

*IMMP2L* encodes subunit 2 of the peptidase required for transport across the inner mitochondrial membrane (Burri et al., 2005). It is a candidate for Gilles de la Tourette syndrome (GTS) (Bertelsen et al., 2014; Gimelli et al., 2014), a disorder of multiple vocal and motor tics that also affects language processing (Frank, 1978)—even speeding some aspects of grammar usage (Walenski et al., 2007). A region in *IMMP2L* with a breakpoint linked to GTS and speech delay/reduced speech development (in addition to microgenia and an ear malformation) has also been associated with ASD and speech-language disorder (Petek et al., 2001). Indeed, there is suggestive evidence linking *IMMP2L* to ASD (Maestrini et al., 2010; Zhang et al., 2018b) (though see Petek et al., 2007), and patients with CNV duplications/deletions of

*IMMP2L* have been reported to exhibit ASD symptoms, language delay, and numerous minor physical anomalies including microcephaly, micrognathia, large central incisors, and brachydactyly (Gimelli et al., 2014). It has also been associated with SZ in GWAS analyses by Ripke et al. (2014) and Pardiñas et al. (2018). An insertion/deletion polymorphism in a sweep region in *IMMP2L* distinguishes domestic and wild rabbits (Carneiro et al., 2014).

**ITGA9**

Integrin alpha 9, encoded by *ITGA9*, is the α9 subunit of the α9β1 integrin, which serves as a receptor for brain-derived neurotrophic factor (BDNF) and other neurotrophins (Secolin et al., 2013). As a cell adhesion molecule, it binds extracellular matrix (ECM) ligands and is expressed, among other places, on the surface of human and mice egg cells, where it facilitates binding and fusion with sperm (Vjugina et al., 2009).

An *ITGA9* SNP is associated with BD, and *ITGA9* transcripts were increased in peripheral blood of patients with BD relative to controls, suggesting that more abundant α9β1 integrin bind and therefore decrease the availability of BDNF (Secolin et al., 2013), decreased levels of which are characteristic of BD (Fernandes et al., 2011). A GWAS investigating broad psychosis (SZ, schizoaffective disorder, and BD with psychotic features) identified a locus near *ITGA9* as one of three loci meriting further investigation, though no clear association was determined (Psychosis Endophenotypes International Consortium and Wellcome Trust Case-Control Consortium, 2014).

*ITGA9* is positively selected in AMH (Racimo, 2016; Theofanopoulou et al., 2017) and domestic cats, with ECM receptor interactions enriched (Montague et al., 2014). Interestingly, given the similarities of phenotypic changes in domestication and island evolution (Sánchez-Villagra et al., 2016), *ITGA9* contains a fixed mutation in the Korean Jeju horse, enriched in ECM receptor interactions and dilated cardiomyopathy (Srikanth et al., 2019). *ITGA9* also underlies a sweep region detected in domestic cattle, hypothesized to be part of the reproductive changes targeted during domestication (Ramey et al., 2013).

**JAM3**

*JAM3* encodes a protein that forms part of tight junctions. Mutations in *JAM3* cause a syndrome similar to pseudo-TORCH syndrome, entailing hemorrhagic brain destruction, subependymal calcification, and congenital cataracts, suggesting roles for *JAM3* in lens development and CNS endothelium integrity (Mochida et al., 2010; Akawi et al., 2013). SNPs in *JAM3* have been associated with BD in a GWAS analysis by Baum et al. (2008). The gene is selected in domestic horses (Theofanopoulou et al., 2017).

**KCNK10**

*KCNK10* encodes the potassium channel $K_{2P}10.1$, also known as TREK-2, a leak channel that contributes importantly to the background potassium current in dorsal root ganglion neurons and plays a significant role in neuron excitability (Talley et al., 2001; Kang and Kim, 2006; Deng et al., 2009). It is expressed in the CNS, primarily cerebellum and spinal cord (Talley et al., 2001), as well as in the developing mouse cerebral cortex, with a role in neuronal migration (Hur et al., 2012; Bando et al., 2014). The TREK-2 channel is involved in migraine induction via its impact on neuron excitability, (Royal et al., 2019), calling to mind sensory sensitivity as seen in migraines and ASD. In a study of DNA methylation in patients with SZ, *KCNK10* was one of 13 genes associated to cerebellar volume compared to healthy controls (Liu et al., 2015). The channel has a role in mood disorders, demonstrating inhibition by antipsychotics and antidepressants, though not by mood stabilizers (Kennard et al., 2005; Thümmler et al., 2007; Kim et al., 2017). It is in the same family as *KCNK9*, mutations in which underlie the Birk-Barel imprinting syndrome, characterized by ID with limited speech, feeding difficulty, micro/retrognathia, dolichocephaly, narrow forehead, ear abnormalities, high nasal bridge, broadened tip of nares, short and broad philtrum, and large incisors (Barel et al., 2008; Zadeh and Graham, 2017). *KNCK10* has been selected in domestic horses (Schubert et al., 2014; Theofanopoulou et al., 2017).

**LRRC36**

*LRRC36* encodes leucine-rich repeat-containing protein 36. While little is known about it, the gene has been associated to BD by GWAS (Moskvina et al., 2009) and is selected in domestic cat (Montague et al., 2014).

**MAOA**

*MAOA* encodes the monoamine oxidase protein A, located on the outer mitochondrial membrane, which metabolizes the neurotransmitters serotonin, norepinephrine, and dopamine; a loss-of-function mutation leading to elevations in these neurotransmitters gives rise to Brunner syndrome, characterized by impulsive aggression, hypersexuality, and moderate mental retardation (Brunner et al., 1993). A gene variant with low-activity MAOA (dubbed 'the warrior gene') has been associated with 'extremely violent' criminal history in a study of Finnish prisoners (Tiihonen et al., 2015) and with later antisocial behavior in maltreated children (Caspi et al., 2002), consistent with findings of aggressive behavior in *MAOA*-knockout mice (Cases et al., 1995). *MAOA* has been identified in linkage studies as a susceptibility gene for SZ (Ng et al., 2009), and mutations in the gene have also been associated with BD (Lim et al., 1995; Shih and Thompson, 1999; Preisig et al., 2000; Craddock et al., 2005; Fan et al., 2010; Eslami Amirabadi et al., 2015), unipolar depression/MDD (as treated with MAOA-selective inhibitors such as Moclobemide) (Fan et al., 2010), social phobia (as also sometimes treated with MAOA-inhibitors) (Liebowitz et al., 1993; Montgomery, 1999; Voltas et al., 2015), panic disorder (Gupta et al., 2015), ADHD (Das et al., 2006; Gupta et al., 2015), and conduct disorder (Prom-Wormley et al., 2009), as well as variability in aggressiveness and impulsivity among adult males of the general (i.e., non-patient) population (Manuck et al., 2000). In males, *MAOA* promoter methylation is associated with schizophrenia (Chen et al., 2012d) and several other psychiatric disorders (Ziegler and Domschke, 2018), with findings of increased *MAOA* expression in brains of patients with SZ postmortem (Purves-Tyson et al., 2017).

With regard to domestication, different polymorphisms of the gene have been demonstrated in wild animals compared to domestic species (Saif et al., 2018). In dogs, a study of *MAOA* polymorphism across breed groups found increased diversity in ancient breeds compared to modern breeds (Sacco et al., 2017). One polymorphism, c.-212 A > G, was notable as it suggests replacement of the G allele, the major allele in wolves, with the A allele, the major allele in dogs and other domesticates, as part of the domestication process; it also indicates the potential loss of a heat shock factor binding site, leading the authors to consider whether it increases docility by functionally increasing *MAOA* expression, though this remains to be demonstrated. Polymorphism in the *MAOA* promoter has also been associated with aggression in cattle (Eusebi et al., 2020). Epigenetic regulation also appears to modify *MAOA* expression in dogs, as different DNA methylation of the promoter among dog breeds is associated with expression in the brain (Eo et al., 2016). DNA methylation of *MAOA* (in addition to *TPH1* (see below), *OXTR*, *COMT*, *HTR1A*, *WFS1*, and *SLC6A4*) has also been shown to distinguish domestic dogs from gray wolves (Banlaki et al., 2017).

**MAOB**

MAOB functions like its MAOA counterpart but preferentially metabolizes benzylamine and phenylethylamine in addition to dopamine (Shih and Thompson, 1999). It is expressed in many neural crest derivatives, including cranial and dental mesenchyme, aorta, etc. (Vitalis et al., 2003). It is also associated with negative emotionality (Dlugos et al., 2009).

*MAOB* has been linked to BD by association analysis (Lin et al., 2000), and the selective MAOB inhibitor pargyline has been reported to induce manic psychosis (Folks and Arnold, 1983). Furthermore, platelet MAOB—a proposed peripheral marker of brain MAOB activity—decreases with lamotrigine treatment in bipolar depression (Muck-Seler et al., 2008). In rats, lamotrigine inhibits serotonin reuptake (Southam et al., 1998; Consoni et al., 2006), and in mice, *MAOB* overexpression is associated with increased turnover of serotonin in the nucleus accumbens (Kato et al., 2018) (see *TPH1* below). Treatment with selective

MAOB inhibition has been shown to be beneficial in both unipolar and bipolar depression (Mendlewicz and Youdim, 1983).

A locus near *MAOB* is associated with SZ by linkage study (Dann et al., 1997), and recently selected haploptypes of *MAOB* are associated with SZ susceptibility (Carrera et al., 2009). Overall, evidence of *MAOB* association with SZ is mixed, with stronger support for an association with psychosis more generally (see Bergen et al., 2009). There is also suggestive evidence of a role for *MAOB* in ASD, from whole-exome sequencing (Al-Mubarak et al., 2017) and microdeletion (e.g., Whibley et al., 2010) studies, with *MAOB* affecting serotonin metabolism and specific behaviors in ASD (Chakraborti et al., 2016).

*MAOB* is also associated with PTSD with psychosis (Pivac et al., 2007), Alzheimer disease (Saura et al., 1994) including AD with psychotic features (Mimica et al., 2008), ADHD (Ribasés et al., 2009), and suicidality (Niculescu et al., 2015). It is positively selected in the domestic dog (Cagan and Blass, 2016; Theofanopoulou et al., 2017) and associated with differences in aggression between dog breeds (Hashizume et al., 2005).

Regarding both genes, *MAOA* and *MAOB* have been associated with risk for Parkinson disease (Hotamisligil et al., 1994; Liu et al., 2014c). An atypical form of the retinopathy Norrie disease, entailing contiguous deletion of *MAOA* and *MAOB* in addition to the disease-defining *NDP* gene, has been reported with microcephaly, psychotic features, autism-like behavior, lack of verbal language, small stature, hypogonadism, epilepsy, and developmental delay (Berger et al., 1992; Collins et al., 1992; Chen et al., 1995; Saito et al., 2001; Suárez-Merino et al., 2001; Rodriguez-Revenga et al., 2007; Whibley et al., 2010; Smith et al., 2012; Jia et al., 2017). Notably, a patient with duplication of *MAOA*, *MAOB*, and *NDP* has been described in a case report as having Hirschsprung disease, friendly temperament, osteoporosis, intractable epilepsy, and developmental delay (Klitten et al., 2011).

In *MAOA/B* knockout mice, high brain serotonin levels are observed in addition to ASD-like behaviors, including social and communication deficits (Bortolato et al., 2013). In addition, the intron 2 dinucleotide repeats in *MAOA* and *MAOB*, *MAin2* and *MBin2* (respectively), exhibit increased polymorphism in bonobos compared to chimpanzees (Garai et al., 2014), reflecting differences in aggression between these species, for which self-domestication has been proposed an explanation (Hare et al., 2012).

**MBP**

*MBP* encodes myelin basic protein, a key component of the myelin sheath of oligodendrocytes and (neural crest-derived) Schwann cells in the central and peripheral nervous systems, respectively, which influences velocity of axonal impulse conduction (Stassart et al., 2018). It is primarily known for its roles in multiple sclerosis and other myelin disorders (Banik, 1992; Shah and Tobias, 2006). Regarding behavior, socially isolated mice exhibit deficits in PFC myelination, with behavioral and transcriptional changes that were reversed with social reintegration, suggesting social experience modifies myelination in a form of adult plasticity (Liu et al., 2012a).

*MBP* expression is decreased in the brains of patients with BD and patients with SZ (Tkachev et al., 2003), with oligodendrocyte dysfunction thought to be important in the etiology of both conditions (Yu et al., 2014a). *MBP* has been associated to BD by GWAS (Baum et al., 2008a), and in patients with BD, expression of *MBP* in peripheral blood is increased during mania: of ten candidate genes for mood state biomarkers, *MBP* scored highest, and five were involved in myelination (Le-Niculescu et al., 2009). Abnormal myelination has been reported in BD and SZ (Chambers and Perrone-Bizzozero, 2004) and even correlated to verbal memory in BD (Sehmbi et al., 2018). In addition, *MBP* has been associated with SZ by convergent functional genomics (Ayalew et al., 2012) and case-control analysis (Baruch et al., 2009). *MBP* expression in the blood of patients with SZ is decreased (Glatt et al., 2011), consistent with findings of myelin and oligodendrocyte abnormalities in SZ (Davis et al., 2003; Haroutunian and Davis, 2007;

Takahashi et al., 2011). Myelin abnormalities also characterize ASD (Gozzi et al., 2012; Steinman and Mankuta, 2019), with *MBP* upregulated in the cerebellum of those with ASD (Zeidán-Chuliá et al., 2016) and elevated levels of autoantibodies against MBP found in the peripheral blood of children with ASD and their mothers (Singh et al., 1993; Abou-Donia et al., 2019). Myelin abnormalities also characterize Williams syndrome (Marenco et al., 2007; Arlinghaus et al., 2011; Avery et al., 2012; Barak et al., 2019), and interestingly, deletion of *GTF2I*—a gene deleted in WS that underlies hypersociability in that condition as well as in dog domestication (vonHoldt et al., 2017)—causes myelin abnormalities and WS-like hypersociability in mice that were both rescuable with a remyelinating drug (Barak et al., 2019).

*MBP* is positively selected in the domestic dog (Freedman et al., 2016; Theofanopoulou et al., 2017; Pendleton et al., 2018), and reduced white matter anisotropy—in addition to reduced amygdala and enlarged mPFC volumes—in domestic compared to wild rabbits is thought to have been part of reducing the fear response in the domestication of this species (Brusini et al., 2018).

**MC2R**

*MC2R* encodes the melanocortin receptor-2, the ACTH receptor, expressed in the brain and adrenal gland (Gragnoli, 2014). Because the neural crest hypothesis of domestication centers around HPA axis-mediated stress (Wilkins et al., 2014), HPA axis genes are particularly of note; indeed, mutations in *MC2R* are found in familial glucocorticoid deficiency, resulting in a low-cortisol state (Clark et al., 1993). The gene has been implicated in BD by linkage studies, with evidence for a parent-of-origin effect (Detera-Wadleigh et al., 1995; Bickeböller et al., 1997; Segurado et al., 2003). *MC2R* is preferentially expressed in the adrenal cortex, and in domestic chickens, it is upregulated in adrenal glands relative to wild counterparts (Fallahsharoudi et al., 2015; Løtvedt et al., 2017). It is positively selected in domestic dogs (Theofanopoulou et al., 2017) and has been proposed as a candidate gene for herding behavior in canids (Hall and Wynne, 2012). A related gene, *MC1R*, is associated with pigmentation/coat-color variation in humans and several domesticates (Switonski et al., 2013); it is also linked to BD, as are *MC3R*, *MC4R*, and *MC5R* (Cheng et al., 2006; Gragnoli, 2014). Notably, foxes selected for tameness have lower ACTH levels both at baseline (Gulevich et al., 2004; Trut et al., 2004) and in response to stress (Hekman et al., 2018) compared to those selected for aggression, a finding paralleled in rats (Shikhevich et al., 2002).

**MCHR2**

*MCHR2* encodes melanin-concentrating hormone (MCH) receptor-2, a G protein-coupled receptor (GPCR) for MCH, which is important in feeding and energy metabolism as well as mood, with MCH promoting depression- and anxiety-like behaviors (Barson et al., 2013). *MCHR2* is expressed in the CNS (especially in the amygdala, temporal cortex, and hippocampus) and weakly in the periphery (pituitary, pancreas, and adipose tissue) (Presse et al., 2014). It is not functionally expressed in rodents, but in transgenic mice expressing human *MCHR2*, it was found to protect against diet-induced obesity (Chee et al., 2014). *MCHR2* has been associated to BD by association mapping, and to BD and SZ by genotyping (Miller et al., 2009; Jamra et al., 2010), consistent with GWAS results indicating risk loci for both conditions near *MCHR2* (Levinson et al., 2000; Dick et al., 2003; Lambert et al., 2005). Notably for the domestication phenotype, MCH leads to pale skin in fish (Presse et al., 2014). *MCHR2* is positively selected in domestic dog (Freedman et al., 2016; Theofanopoulou et al., 2017).

**NCAPD3**

*NCAPD3* encodes a subunit of the condensin-2 complex, which is involved in chromosome condensation in mitosis (Martin et al., 2016). Mutations in *NCAPD3* cause primary autosomal recessive microcephaly 22, with autistic behavior, sloping forehead, ID, and low birthweight also reported (Martin et al., 2016). Alternate splicing of *NCAPD3* distinguishes those with history of psychosis from those without (Glatt et al., 2009). *NCAPD3* was enriched for CNV case hits in a combined ASD and ADHD sample, with CNV regions common to both disorders excluded (Martin et al., 2014); CNVs in *NCAPD3* have also been associated with SZ (Magri et al., 2010).

In two Jacobsen syndrome patients with a distal 11q microdeletion, *NCAPD3* was one of five candidate genes for developmental delay/ID; microcephaly, severe ID, cardiac defects, low birth weight, and facial dysmorphism were reported (Ji et al., 2010). Phenotypes commonly seen in Jacobsen syndrome include growth restriction, ID, cardiac defects, macrocrania, trigonocephaly, short nose with anteverted nares, nasal bridge that is flattened or prominent, small ears that may be low-set and posteriorly rotated (plus other ear malformations), retrognathia, and philtrum abnormalities (Mattina et al., 2009), as well as an increased risk of ASD (Maruani et al., 2015). *NCAPD3* has been positively selected in domestic horse (Schubert et al., 2014; Theofanopoulou et al., 2017).

**NEK4**

*NEK4* encodes a never-in-mitosis A kinase that regulates replicative senescence and cell cycle arrest in response to double-stranded DNA damage (Nguyen et al., 2012). It is highly expressed in the brain and is a ciliopathy candidate gene (Coene et al., 2011), notable in light of the role of cilia in neurodevelopment (Takata et al., 2017). Indeed, *NEK4* is one of twenty genes associated with neuropsychiatric conditions including SZ, BD, and ASD shown to be important for cilia stability in a loss-of-function study by Marley and von Zastrow (2012).

*NEK4* is linked to SZ by GWAS analyses by the Schizophrenia Working Group of the Psychiatric Genomics Consortium (Ripke et al., 2014). Dysregulated alternate splicing of regions encompassing *NEK4* has also been implicated in SZ (Takata et al., 2017). The Psychiatric GWAS Consortium Bipolar Disorder Working Group has found, in combined analysis of non-overlapping SZ and BD GWAS samples, an association of SNPs in a four-gene region including *NEK4* with both conditions (Sklar et al., 2011). In the study proposing *STAB1* as a candidate gene for BD, Witt et al. (2014) note its proximity to the *NEK4-ITIH1-ITIH3-ITIH4* region. Furthermore, BD and SZ risk alleles at the chromosome 3p21.1 region correlate with *NEK4* expression as well as dendritic spine pathology (Yang et al., 2020b), which has been implicated in the pathophysiology of ASD, BD, and SZ (Penzes et al., 2011; Konopaske et al., 2014).

*NEK4* has been selected in both domestic cats (Montague et al., 2014) and AMH (Racimo, 2016; Theofanopoulou et al., 2017).

**NPAS3**

*NPAS3* is a transcription factor primarily expressed in the brain, thought to have played a significant role in human brain evolution given that it contains the largest number of accelerated regulatory sequences in the human genome, with other genes with high numbers of human-accelerated elements also associated with ASD and SZ (Kamm et al., 2013). It is known for its role in neurodevelopment, being expressed in the human fetal brain—especially the hippocampus, where it supports neurogenesis (Michaelson et al., 2017), which is altered in BD, SZ, and other psychiatric disorders (Kempermann et al., 2008; Nurnberger et al., 2014; Michaelson et al., 2017)—throughout gestation (Gould and Kamnasaran, 2011) and is a candidate gene for holoprosencephaly, with which comorbid schizophrenia has been reported (Hercig et al., 1994; Kamnasaran et al., 2005).

Interestingly, *NPAS3* is also a candidate gene for Sotos syndrome (Visser et al., 2010), which is characterized by overgrowth in childhood, delays in cognitive and motor development including speech delay and learning difficulty, cardiac anomalies, and reports of social inhibition and psychosis (Ruggieri and Arberas, 2003; Compton et al., 2004; Baujat and Cormier-Daire, 2007). The overgrowth is often preceded by neonatal feeding difficulties, as in Prader-Willi syndrome, with which it also shares behavioral characteristics (Sheth et al., 2015), calling to mind the effects of imprinting which Badcock and Crespi (2006) have hypothesized to be part of the etiology of ASD and SZ. Sotos syndrome also entails several domestication-relevant minor physical anomalies including macrocephaly, large ears, pointed chin, micrognathia as well as prominent jaw, and anteverted nares (Baujat and Cormier-Daire, 2007).

*NPAS3* has been determined a master regulator of risk genes of neuropsychiatric illness, including SZ, BD, MDD, ADHD, and ID, as well as regulating *UBE3A*, mutations in which can underlie Angelman syndrome and Prader-Willi syndrome (Pickard et al., 2006; Michaelson et al., 2017). *NPAS3* has been identified as a susceptibility gene for SZ in linkage studies by Lewis et al. (2003) and Ng et al. (2009), with several variants associated with SZ (Macintyre et al., 2010; González-Peñas et al., 2015) and comorbid learning disability (Pickard et al., 2005). A translocation in the gene was found to segregate in a family affected by SZ (Kamnasaran et al., 2003; Pickard et al., 2005), and other mutations in *NPAS3* have similarly segregated in families with SZ (Yu et al., 2014b), with resultant protein aggregation in one case (Nucifora et al., 2016). It has also been associated with BD (Pickard et al., 2009; Nurnberger et al., 2014), including by GWAS analysis (Huang et al., 2010).

In domestication, *NPAS3* has been selected in domestic cattle (Qanbari et al., 2014), and decreased copy number variation of *NPAS3* was found in domestic compared to wild goats (Dong et al., 2015).

**NPTX1**
Neuronal pentraxin-1 (NP1), encoded by the *NPTX1* gene, is expressed in the CNS where it is involved with neuronal apoptosis (Hooper et al., 2017). *NPTX1* has been found to be a candidate gene for ASD by in silico analysis (Iurov et al., 2010) and is a candidate for BD based on breakpoint analysis (Rajkumar et al., 2015). In mice, NP1 has been shown to be involved in neurodegeneration (Hooper et al., 2017). *NPTX1* has been positively selected in domestic cattle (Theofanopoulou et al., 2017).

**NR3C1**
*NR3C1* encodes the glucocorticoid receptor (GR) and is ubiquitously expressed, with GR expression and responsivity implicated in depression and PTSD (Yehuda et al., 2002; Ruiz et al., 2007; Chourbaji et al., 2008). Glucocorticoid levels affect stability of dendritic spines, with regression potentially precipitating depression or psychosis (Bennett Ao, 2008). *NR3C1* undergoes epigenetic modification in response to in utero stress, which has been hypothesized to play a role in SZ, ASD, and depression (Kundakovic and Jaric, 2017; Watkeys et al., 2018). It has been associated with BD (Ceulemans et al., 2011), and in adult offspring of people with BD, *NR3C1* had higher methylation rates than controls (Duffy et al., 2019). In women with PTSD, *NR3C1* methylation negatively correlated with severity of illness (Schechter et al., 2015). Notably, epigenetic modification in the context of stress and fear is hypothesized to be an important aspect of domestication (Jensen, 2015). In MDD, *NR3C1* polymorphism predict attention and working memory (Keller et al., 2017). *NR3C1* expression is increased in domesticated chicken compared to wild counterparts (Fallahsharoudi et al., 2015); it is also a candidate gene for reproductive seasonality in domestic versus wild rabbits (Carneiro et al., 2015).

**NR3C2**
*NR3C2* encodes the aldosterone or mineralocorticoid receptor (MR), which also binds glucocorticoids. SNPs in *NR3C2* are moderately associated with BD but are distributed widely over the gene (Ceulemans et al., 2011). *NR3C2* is considered a strong candidate for ASD with whole-exome sequencing (De Rubeis et al., 2014), whole-genome sequencing (Turner et al., 2016), and analyses (Krumm et al., 2014; Ruzzo et al., 2019; Zhou et al., 2019). Interestingly, *NR3C2* polymorphisms predict memory performance in MDD (Keller et al., 2017) and correlate with intelligence (Pan et al., 2011). The gene is positively selected in domestic horse (Schubert et al., 2014; Theofanopoulou et al., 2017)

**NRXN1**
*NRXN1* encodes Neurexin-1, a presynaptic cell adhesion molecule that critically influences neuritogenesis (Gjørlund et al., 2012) and synaptogenesis, complexing with postsynaptic neuroligin to facilitate development of glutamatergic and GABAergic synapses (Jenkins et al., 2016). Deletion in *NRXN1* is characterized by severe language delay, ASD, seizures, and ID (Ching et al., 2010; Béna et al., 2013). In a study of three unrelated patients with *NRXN1* deletions, diagnoses included ID, autism, and BD; shared

dysmorphisms included a long face and prominent premaxilla; tooth, ear, and philtrum abnormalities were also reported (Viñas-Jornet et al., 2014), with increased head size reported in other cases (Schaaf et al., 2012). In another study of patients with deletions in *NRXN1*, co-occurring CNVs including Williams syndrome deletion and 22q11.2 (DiGeorge region) duplications were reported, with microcephaly affecting the patient with comorbid WS and several others, and autism, speech delay, social communication difficulties, developmental delay, and learning difficulties also reported (Curran et al., 2013). *NRXN1* is mutated in Pitt-Hopkins-like syndrome, characterized by severe ID, lack of speech, and macrostomia (Zweier et al., 2009). Reduced social approach—and increased aggression in males—is observed in *NRXN1*-knockout mice (Grayton et al., 2013), while heterozygotes exhibit loss of social memory (Dachtler et al., 2015). Intriguingly, ASD-like behaviors in mice with mutated *NRXN1* can be reversed with later inhibition of the protein (Rabaneda et al., 2014).

*NRXN1* is a strong ASD candidate gene supported by linkage and CNV analyses (Autism Genome Project Consortium et al., 2007; Sanders et al., 2011; Prasad et al., 2012), mutation analysis (Liu et al., 2012c), GWAS (Walker and Scherer, 2013), and studies of rare and ultra-rare sequence variants (Feng et al., 2006; Kim et al., 2008; Yan et al., 2008). In postmortem brain samples, *NRXN1-β* expression was increased in patients with SZ, while *NRXN1-α* was increased in those with BD, relative to healthy controls (Jenkins et al., 2016). CNVs in *NRXN1* are also associated with BD (Noor et al., 2014), and the gene has been linked to BD by GWAS analysis (O'Dushlaine et al., 2011). *NRXN1* is found within CNVs conferring increased risk of SZ (Rujescu et al., 2009; Magri et al., 2010; Marshall et al., 2017), and *NRXN1* polymorphism associates with antipsychotic treatment response in SZ (Jenkins et al., 2014). *NRXN1* deletions have also been reported in families with SZ (Todarello et al., 2014). The role of *NRXN1* in overlapping ASD and SZ susceptibility has been proposed to be related to white matter abnormalities associated with *NRXN1* mutation (Voineskos et al., 2011). CNVs in *NRXN1* also increase risk of Tourette syndrome (Huang et al., 2017) and correlate with antidepressant response in MDD (Tansey et al., 2014).

*NRXN1* is a domestication candidate positively selected in cattle (Qanbari et al., 2014; Theofanopoulou et al., 2017).

**PARVG**

Part of the parvin family, *PARVG* binds actin as a focal adhesion protein expressed preferentially in lymphoid tissue (Korenbaum et al., 2001). It has been identified as a potential driver gene of Alzheimer disease (Mukherjee et al., 2019), and reduced parvin expression in cancer, relative to normal tissue, has been reported (Korenbaum et al., 2001). In 22q13.31 CNV deletions associated with schizophrenia risk, *PARVG* is one of several genes deleted (Liu et al., 2012b). It is positively selected in domestic cats (Montague et al., 2014).

**PCSK5**

Proprotein convertase subtilisin/kexin type 5, encoded by *PCSK5*, cleaves proteins such as growth/differentiation factor 11 (GDF11), which sets anterior/posterior patterning by regulation of *Hox* gene expression (Essalmani et al., 2008; Tsuda et al., 2011). It is also involved in lipoprotein and insulin metabolism and has been linked with cognitive impairment via its effects on blood pressure (Chen et al., 2019). It is expressed in the CNS, especially the spinal cord and the pineal gland, with studies in zebrafish suggesting a role in spatial awareness and sensation of the environment (Chitramuthu et al., 2010; Ivanov et al., 2018).

Nonsynonymous mutations in *PCSK5* have been shown in patients with VACTERL (vertebral, anorectal, cardiac, tracheoesophageal, renal, limb malformation) and caudal regression syndrome, and mice with mutated *PCSK5* display similar malformations (Szumska et al., 2008).

Microdeletions including *PCSK5* have been reported in cases of patients exhibiting autistic behavior, speech delay, microcephaly, ready laughter, global developmental delay, and minor physical anomalies including teeth abnormalities, short nose, micrognathia, flat nasal bridge, prominent chin, philtrum abnormalities, hypertelorism, macrostomia, and hypertrichosis (Boudry-Labis et al., 2013; Ivanov et al., 2018). SNPs near *PCSK5* were found to be associated with SZ by Need and colleagues (2009).

**PIK3C3**

The class III phosphoinositide 3-kinase, encoded by *PIK3C3*, is a regulator of vesicular trafficking and autophagy that also plays a role in axon extension and migration of cortical neurons during embryonic neurodevelopment (Inaguma et al., 2016). Widely expressed in the brain, *PIK3C3*'s catalytic product PI3P is expressed in dendritic spines, with deletion of the gene resulting in loss of synapses, gliosis, and neurodegeneration (Zhou et al., 2010; Wang et al., 2011). *PIK3C3* has been identified as a susceptibility gene for SZ in linkage studies (Lewis et al., 2003), with variants in its promoter associated with SZ (Saito et al., 2005; Tang et al., 2008) and BD (Stopkova et al., 2004; Carrard et al., 2011). *PIK3C3* CNVs are associated with ASD (Sbacchi et al., 2010; Ji et al., 2016), as are CNVs in the *PIK3R4* gene which encodes a protein that interacts with *PIK3C3* to regulate the PI3K pathway (Cuscó et al., 2008). Indeed, the PI3K pathway is oppositely altered in SZ and ASD (Crespi et al., 2010), and deletion of *PIK3C3* has been detected in a patient with specific learning disorders (Inaguma et al., 2016). The gene has been selected during horse (Schubert et al., 2014) and duck (Zhang et al., 2018c) domestication.

**POMC**

Pro-opiomelanocortin, encoded by the *POMC* gene, is cleaved to produce ACTH, α-MSH (melanocyte-stimulating hormone), and endogenous opioids such as β-endorphin. Through these peptides, *POMC* is thought to contribute to pigmentation / coat color changes seen in domestication (Cieslak et al., 2011). POMC is part of the central melanocortin system which regulates appetite and is a drug target in obesity treatment in humans (Zhan, 2018). Indeed, a deletion in *POMC* is thought to drive food interest and obesity in Labrador retriever dogs (Raffan et al., 2016), and loss-of-function mutations in humans cause early obesity with adrenal insufficiency and red hair (Krude et al., 1998, 2003). Despite mutations in *POMC* not being associated with BD or SZ in initial analyses (Feder et al., 1985), increased levels of pro-opiomelanocortin have been found in the pituitary glands of BD patients postmortem (Stelzhammer et al, 2015). ACTH levels are also increased in BD (Belvederi Murri et al., 2016), and hyposecretion of POMC products has been proposed as a mechanism in the pathophysiology of ASD (Chamberlain and Herman, 1990), with POMC products' release profile correlating with self-injurious behavior (Sandman et al., 2000). Pituitary *POMC* expression is lower in domesticated chicken than in wild counterparts, hypothesized to factor into the reduced HPA axis reactivity that is exhibited by domesticated chickens and the proposed target of domestication selection (Wilkins et al., 2014; Løtvedt et al., 2017). As part of this general reduction of HPA axis activity in domestication, *POMC* expression reduction is also seen in foxes selected for tameness in the fox farm domestication experiment (Trut et al., 2009).

**PPARD**

*PPARD* encodes peroxisome proliferator-activated receptor-δ, a transcription factor and nuclear hormone receptor with diverse roles including fat metabolism, the Wnt pathway, keratinocyte differentiation, and chondrocyte proliferation and differentiation (Di-Poï et al., 2004; Zandi et al., 2008; Ren et al., 2011). Its highest expression is in the embryonic brain, suggesting a role in neurodevelopment (Zandi et al., 2008). *PPARD* is necessary for normal neuron function and is repressed in Huntington disease (Dickey et al., 2016). It is associated with BD, consistent with Wnt dysfunction in the condition (Zandi et al., 2008b), and with SZ (Sun et al., 2008b). Among domesticates, *PPARD* inhibits cartilage growth in the external ear of pigs, while a missense mutation in the gene (which likely occurred after domestication) results in large, floppy ears in the Chinese Erhualian breed (Ren et al., 2011; Zhang et al., 2017b).

**PRKCZ**
*PRKCZ* encodes protein kinase C, zeta (PKCζ), which is involved in maintenance of long-term memory (Migues et al., 2010; Li et al., 2011; though see Lee et al., 2013). The gene is deleted in most cases of 1p36 deletion syndrome, which is characterized by ID, speech/language delay, ASD/autistic behavior, restricted growth, microcephaly, brachycephaly, wide nasal bridge, flat nose, and pointed chin (Gajecka et al., 2007; Cunha et al., 2014). The condition often also entails cardiac malformations or cardiomyopathy, for which *PRKCZ* is a candidate gene (Zaveri et al., 2014). *PRKCZ* has been linked to BD (Kandaswamy et al., 2012), SZ with comorbid tobacco use (Ma et al., 2020), ASD (Chow et al., 2012), including anxiety in ASD (Gao et al., 2019), and ethanol consumption in mice (Lee et al., 2014). It is positively selected in domestic horse (Schubert et al., 2014; Theofanopoulou et al., 2017).

**RNF144B**
*RNF144B* encodes a ubiquitin ligase and regulates cell proliferation and apoptosis as well as NF-κB in macrophages (Raphaka et al., 2017). Accordingly, it is associated with cancer (e.g., Zhou et al., 2018) and may have a role in dilated cardiomyopathy (Peche et al., 2013). *RNF144B* has been linked to ASD in a twin study (Hu et al., 2019) and is differentially expressed in human neurons after androgen exposure, suggesting a potential role in the male excess hypothesis of ASD (Quartier et al., 2018). It is upregulated in patients with SZ, persisting even weeks into antipsychotic treatment (Kumarasinghe et al., 2013). *RNF144B* is positively selected in domestic cattle (Theofanopoulou et al., 2017).

**SEC63**
*SEC63* encodes part of the protein translocon complex of the endoplasmic reticulum (ER), with mutations causing autosomal dominant polycystic liver disease (Davila et al., 2004). This is thought to be due to ER stress, which is also associated with disorders of myelin, with zebrafish models exhibiting both liver pathology and reduced myelination (Monk et al., 2013). Another study in zebrafish, focused on craniofacial development, found *SEC63* to be involved in regulating neural crest cell development and migration (Wang et al., 2019). SNPs in *SEC63* are among those that differentiate wild and domesticated Eurasian perch (Chen et al., 2017). *SEC63* has also been shown to dysregulated in autism with a fragile X mutation versus autism with a 15q11–q13 duplication (Nishimura et al., 2007).

**SEMA3D**
*SEMA3D* encodes the ligand of plexin A2, which is associated with SZ, and is involved in axon guidance and neuroplasticity (Kruger et al., 2005; Fujii et al., 2011). It is expressed in dorsal root ganglia (Takahashi et al., 2009) and in enteric neural crest cells, as a candidate in Hirschsprung disease (Jiang et al., 2015). It has been associated with BD by GWAS (Kuo et al., 2014) and with ASD (Sbacchi et al., 2010). Additionally, it is a candidate gene for SZ based on a genotyping of a 500-patient sample (Fujii et al., 2011), having also been shown to be downregulated in the prefrontal cortices of patients with SZ (Gilabert-Juan et al., 2015). It is positively selected in the domestic dog (Theofanopoulou et al., 2017).

**SNAP29**
*SNAP29* encodes a SNARE protein involved in synaptic transmission (Su et al., 2001). It is encompassed by the 22q11 region associated with SZ, BD, and ASD, and is deleted in the neurocristopathy DiGeorge syndrome, which is associated with SZ and BD (Saito et al., 2001; Clements et al., 2017). *SNAP29* is a candidate driver of 22q11.2 CNV-associated ASD, and polymorphisms in its promoter are associated with SZ (Saito et al., 2001; Wonodi et al., 2005); linkage and association studies have also connected the gene to SZ (Lewis et al., 2003; Sun et al., 2008a; Marshall et al., 2017). Loss-of-function *SNAP29* mutations cause CEDNIK (cerebral dysgenesis, neuropathy, ichthyosis and keratoderma) syndrome (Fuchs-Telem et al., 2011). It is positively selected in the domestic dog (Cagan and Blass, 2016; Theofanopoulou et al., 2017).

**STAB1**

*STAB1* encodes the Stabilin-1 (also known as FEEL-1) scavenger receptor that binds low-density lipoprotein and is linked with sexual dimorphism in fat distribution (Heid et al., 2010). It is expressed on sinusoidal endothelial cells and tissue macrophages and upregulated during inflammation and tumor progression (Kzhyshkowska, 2010), binding bacteria and modulating angiogenesis (Adachi and Tsujimoto, 2002). *STAB1* is located near the *NEK4-ITIH1-ITIH3-ITIH4* region associated with SZ and BD and was established as a candidate gene for BD by Witt et al. (2014), who demonstrated that it is upregulated during both mania and euthymia. It has also been linked to SZ by GWAS (Lee et al., 2013b), whole exome sequencing (Li et al., 2020), and GWAS analysis by the Schizophrenia Working Group of the Psychiatric Genomics Consortium (Ripke et al., 2014). *STAB1* is also upregulated in the peripheral blood of patients with Williams syndrome (Niego and Benítez-Burraco, 2019). Additionally, it has been associated with Parkinson disease (Germer et al., 2019), pediatric multiple sclerosis (Liguori et al., 2019) and Alzheimer disease (Giil et al., 2017). It is positively selected in both AMH and horse (Schubert et al., 2014; Racimo, 2016), one of 41 genes selected in both AMH and one or more domesticates (Theofanopoulou et al., 2017).

**TH**

*TH* encodes tyrosine hydroxylase, which catalyzes the conversion of tyrosine into L-dopa, the rate-limiting step in the synthesis of the catecholamines—dopamine, epinephrine, and norepinephrine (Nagatsu, 1995). Melanin is also synthesized from L-dopa, with tyrosinase the rate-limiting enzyme in this pathway in mammals—though expressing TH in the retinal pigment epithelium cells of albino mice rescues visual function that is commonly compromised by tyrosinase mutation in albinism (Lavado et al., 2006). *TH* is expressed in the substantia nigra and locus ceruleus of the CNS as well as the adrenal medulla (Nagatsu and Ichinose, 1991). Mutations in *TH* are associated with autosomal recessive dopa-responsive dystonia, which is characterized by Parkinsonism in children (Liu et al., 2004).

*TH* has been connected to BD susceptibility by association (Bellivier et al., 1998) and linkage analysis studies (Lim et al., 1993; Smyth et al., 1996; Lewis et al., 2003; Ng et al., 2009), though some studies have not supported this (e.g., Furlong et al., 1999). It has also been identified as a susceptibility gene for SZ by linkage studies (Lewis et al., 2003; Ng et al., 2009), with TH expression associated with decreased full-scale and verbal IQ (Horiguchi et al., 2014) and suicide risk (Hu et al., 2014) in SZ. TH was also downregulated in rats chronically treated with the atypical antipsychotic clozapine (Tejedor-Real et al., 2003), and epigenetic modulation leading to increased TH protein levels has been found in neurons isolated from patients with BD and SZ (Pai et al., 2019). TH is also increased in the locus coeruleus of patients with MDD (Guo et al., 2019), and TH activity is a marker of chronic stress in laboratory and domestic animals (Chobotská et al., 1998).

In the silver fox domestication experiment, TH enzymatic activity was increased in tame compared to aggressive foxes in the locus coeruleus, hypothalamus and cortex (Dygalo et al., 1988). *TH* was positively selected in dog domestication (Theofanopoulou et al., 2017), with polymorphisms in the gene associated with differences in aggressive (Takeuchi et al., 2005; Cagan and Blass, 2016) and active-impulsive (Kubinyi et al., 2012; Rigterink and Houpt, 2014) behavior between breeds. Interestingly, *TH* was a locus of domestication in the silkworm, with lower expression levels corresponding to lighter body color in domesticated versus wild silkworms (in insects, TH rather than tyrosinase is the rate-limiting enzyme in melanin synthesis) (Yu et al., 2011). With regard to AMH, TH-expressing interneurons are found more abundantly in the human striatum and neocortex than in those of apes, specifically in the dopaminergic innervation associated with speech and language (Raghanti et al., 2016; Calvey, 2019).

**TPH1**

*TPH1* encodes tryptophan hydroxylase, which catalyzes the rate-limiting step in the production of serotonin; *TPH2* is the predominant isoform expressed in the brain, while *TPH1* is expressed in peripheral tissues including the gut, skin, pineal gland, spleen, and thymus (Walther and Bader, 2003; Albert et al., 2011). While CNS expression of *TPH1* has been reported in a human postmortem study—in amygdala, hypothalamus, hippocampus, and raphe nucleus, though at lower levels than *TPH2* (Zill et al., 2007)—there is some uncertainty whether this is distinct from pineal projection (Zhang et al., 2006). In the mouse brain, meanwhile, *TPH1* is expressed in late development, potentially with a role in the development of serotonergic neurons (Nakamura et al., 2006). In any case, *TPH1* polymorphism (in particular, the A218C allele) is strongly associated with suicidal behavior (Rujescu et al., 2003; Bellivier et al., 2004; Li and He, 2006; Beden et al., 2016), especially as part of SZ (Saetre et al., 2010) and MDD (Galfalvy et al., 2009). Polymorphisms in the gene are also associated with decreased PFC activation in response inhibition tasks (Ruocco et al., 2016), impulsive-aggressive behavior (New et al., 1998), reduced anger control (Baud et al., 2009), and increased likelihood of developing borderline personality disorder following childhood abuse (Wilson et al., 2012).

*TPH1* has been identified as a susceptibility gene for SZ by association studies (Allen et al., 2008; Sun et al., 2008a), linkage studies (Lewis et al., 2003; Ng et al., 2009), case-control studies (Watanabe et al., 2007), and others. It is also associated with risk of MDD as well as treatment response (Peters et al., 2004; Viikki et al., 2010), temperament (Andre et al., 2013), and amygdala activation in response to facial stimuli (Lee et al., 2009) in MDD. *TPH1* has been linked to risk of BD (and alcohol dependence) in Caucasian but not Asian populations (Lai et al., 2005; Choi et al., 2010; Chen et al., 2012a). Serotonin deficit is thought to play a key role in the pathophysiology of mania (Shiah and Yatham, 2000), though serotonin reuptake inhibition is known to induce mania in BD (Howland, 1996).

DNA methylation of *TPH1* has been shown to distinguish domestic dogs from gray wolves (Banlaki et al., 2017), and *TPH1* was determined a candidate gene affecting aggression, one of two candidate genes near regions of positive selection in the rat tameness experiment by Albert et al. (2011), in line with prior findings of decreased serotonin levels in the brains of tame relative to aggressive rats (Albert et al., 2008). TPH activity is also increased in domesticated foxes compared to aggressive animals (Popova et al., 1991).

**YWHAH**

Preferentially expressed in the brain, *YWHAH* encodes the η-subtype of the 14-3-3 protein and prevents degradation of the glucocorticoid receptor, positively regulating its transcriptional activation (Kim et al., 2005; Grover et al., 2009). It has been identified as a SZ susceptibility gene by linkage studies (Lewis et al., 2003; Ng et al., 2009) and is also a positional (due to its 22q12 location) and functional candidate gene for BD (Kelsoe et al., 2001; Grover et al., 2009). *YWHAH* is among several genes thought to mediate environmental stress in the etiology of both SZ and BD via the eIF2B complex (Carter, 2007). *YWHAH* is under positive selection in domestic dog and interacts with the domestication candidate *BRAF*, which is positively selected in cat, horse, and AMH (Theofanopoulou et al., 2017).

**ZNF236**

The Kruppel-like zinc finger 236, encoded by *ZNF236*, is expressed ubiquitously in humans but at highest levels in the brain and skeletal muscle, and is regulated by glucose levels (Holmes et al., 1999). It has been considered as a candidate gene for childhood disintegrative disorder—a pre-DSM-5 subtype of ASD characterized by severe regression in two or more areas of development including language, social skill, play, and others—and is thought to play a role in regulating transcription (Gupta et al., 2017). Additionally, *ZNF236* is linked with chromosome 18q deletion syndrome as one of five genes deleted in one of the smallest microdeletions reported (Tassano et al., 2016). This syndrome is characterized by microcephaly, aural atresia and other ear abnormalities, flat midface/nasal bridge, short stature, cardiac anomalies, cerebral demyelination, and ID, with speech delay, smooth philtrum, short upturned nose, and other facial dysmorphisms also reported (Mark et al., 2013; Tassano et al., 2016). Patients with 18q deletion have been reported to have high rates of psychotic, manic, and depressive symptoms (Zavala et al., 2010) as well as autism (O'Donnell et al., 2010) (though neither of these studies identified *ZNF236* as a candidate gene).

References

Abou-Donia, M. B., Suliman, H. B., Siniscalco, D., Antonucci, N., and ElKafrawy, P. (2019). De novo Blood Biomarkers in Autism: Autoantibodies against Neuronal and Glial Proteins. *Behav Sci (Basel)* 9. doi:10.3390/bs9050047.

Abumrad, N. A., Ajmal, M., Pothakos, K., and Robinson, J. K. (2005). CD36 expression and brain function: does CD36 deficiency impact learning ability? *Prostaglandins Other Lipid Mediat.* 77, 77–83. doi:10.1016/j.prostaglandins.2004.09.012.

Adachi, H., and Tsujimoto, M. (2002). FEEL-1, a novel scavenger receptor with in vitro bacteria-binding and angiogenesis-modulating activities. *J. Biol. Chem.* 277, 34264–34270. doi:10.1074/jbc.M204277200.

Ahmad, Y., Bhatia, M. S., Mediratta, P. K., Sharma, K. K., Negi, H., Chosdol, K., et al. (2009). Association between the ionotropic glutamate receptor kainate3 (GRIK3) Ser310Ala polymorphism and schizophrenia in the Indian population. *World J. Biol. Psychiatry* 10, 330–333. doi:10.3109/15622970802688044.

Ahuja, R., Pinyol, R., Reichenbach, N., Custer, L., Klingensmith, J., Kessels, M. M., et al. (2007). Cordon-bleu is an actin nucleation factor and controls neuronal morphology. *Cell* 131, 337–350. doi:10.1016/j.cell.2007.08.030.

Akahoshi, K., Spritz, R. A., Fukai, K., Mitsui, N., Matsushima, K., and Ohashi, H. (2004). Mosaic supernumerary inv dup(15) chromosome with four copies of the P gene in a boy with pigmentary dysplasia. *American Journal of Medical Genetics Part A* 126A, 290–292. doi:10.1002/ajmg.a.20580.

Akawi, N. A., Canpolat, F. E., White, S. M., Quilis-Esquerra, J., Sanchez, M. M., Gamundi, M. J., et al. (2013). Delineation of the Clinical, Molecular and Cellular Aspects of Novel JAM3 Mutations Underlying the Autosomal Recessive Hemorrhagic Destruction of the Brain, Subependymal Calcification and Congenital Cataracts. *Hum Mutat* 34, 498–505. doi:10.1002/humu.22263.

Al-Ayadhi, L., El-Ansary, A., Bjørklund, G., Chirumbolo, S., and Mostafa, G. A. (2019). Impact of Auditory Integration Therapy (AIT) on the Plasma Levels of Human Glial Cell Line–Derived Neurotrophic Factor (GDNF) in Autism Spectrum Disorder. *J Mol Neurosci* 68, 688–695. doi:10.1007/s12031-019-01332-w.

Albert, F. W., Hodges, E., Jensen, J. D., Besnier, F., Xuan, Z., Rooks, M., et al. (2011). Targeted resequencing of a genomic region influencing tameness and aggression reveals multiple signals of positive selection. *Heredity* 107, 205–214. doi:10.1038/hdy.2011.4.

Albert, F. W., Shchepina, O., Winter, C., Römpler, H., Teupser, D., Palme, R., et al. (2008). Phenotypic differences in behavior, physiology and neurochemistry between rats selected for tameness and for defensive aggression towards humans. *Hormones and Behavior* 53, 413–421. doi:10.1016/j.yhbeh.2007.11.010.

Allen, N. C., Bagade, S., McQueen, M. B., Ioannidis, J. P. A., Kavvoura, F. K., Khoury, M. J., et al. (2008). Systematic meta-analyses and field synopsis of genetic association studies in schizophrenia: the SzGene database. *Nat. Genet.* 40, 827–834. doi:10.1038/ng.171.

Al-Mubarak, B., Abouelhoda, M., Omar, A., AlDhalaan, H., Aldosari, M., Nester, M., et al. (2017). Whole exome sequencing reveals inherited and de novo variants in autism spectrum disorder: a trio study from Saudi families. *Sci Rep* 7, 5679. doi:10.1038/s41598-017-06033-1.

Amano, K., Yamada, K., Iwayama, Y., Detera-Wadleigh, S. D., Hattori, E., Toyota, T., et al. (2008). Association study between the Down syndrome cell adhesion molecule (DSCAM) gene and bipolar disorder. *Psychiatric Genetics* 18, 1–10. doi:10.1097/YPG.0b013e3281ac238e.

Ament, S. A., Szelinger, S., Glusman, G., Ashworth, J., Hou, L., Akula, N., et al. (2015). Rare variants in neuronal excitability genes influence risk for bipolar disorder. *Proc Natl Acad Sci U S A* 112, 3576–3581. doi:10.1073/pnas.1424958112.

Anazi, S., Maddirevula, S., Salpietro, V., Asi, Y. T., Alsahli, S., Alhashem, A., et al. (2017). Expanding the genetic heterogeneity of intellectual disability. *Hum. Genet.* 136, 1419–1429. doi:10.1007/s00439-017-1843-2.

Andre, K., Kampman, O., Viikki, M., Illi, A., Setälä-Soikkeli, E., Poutanen, O., et al. (2013). TPH1 A218C polymorphism and temperament in major depression. *BMC Psychiatry* 13, 118. doi:10.1186/1471-244X-13-118.

Arlinghaus, L. R., Thornton-Wells, T. A., Dykens, E. M., and Anderson, A. W. (2011). Alterations in diffusion properties of white matter in Williams syndrome. *Magnetic Resonance Imaging* 29, 1165–1174. doi:10.1016/j.mri.2011.07.012.

Ashok, A. H., Marques, T. R., Jauhar, S., Nour, M. M., Goodwin, G. M., Young, A. H., et al. (2017). The dopamine hypothesis of bipolar affective disorder: the state of the art and implications for treatment. *Mol Psychiatry* 22, 666–679. doi:10.1038/mp.2017.16.

Autism Genome Project Consortium, Szatmari, P., Paterson, A. D., Zwaigenbaum, L., Roberts, W., Brian, J., et al. (2007). Mapping autism risk loci using genetic linkage and chromosomal rearrangements. *Nat. Genet.* 39, 319–328. doi:10.1038/ng1985.

Autism Spectrum Disorders Working Group of The Psychiatric Genomics Consortium (2017). Meta-analysis of GWAS of over 16,000 individuals with autism spectrum disorder highlights a novel locus at 10q24.32 and a significant overlap with schizophrenia. *Mol Autism* 8, 21. doi:10.1186/s13229-017-0137-9.

Avery, S. N., Thornton-Wells, T. A., Anderson, A. W., and Blackford, J. U. (2012). White matter integrity deficits in prefrontal–amygdala pathways in Williams syndrome. *NeuroImage* 59, 887–894. doi:10.1016/j.neuroimage.2011.09.065.

Ayalew, M., Le-Niculescu, H., Levey, D. F., Jain, N., Changala, B., Patel, S. D., et al. (2012). Convergent functional genomics of schizophrenia: from comprehensive understanding to genetic risk prediction. *Molecular Psychiatry* 17, 887–905. doi:10.1038/mp.2012.37.

Badcock, C., and Crespi, B. (2006). Imbalanced genomic imprinting in brain development: an evolutionary basis for the aetiology of autism. *Journal of Evolutionary Biology* 19, 1007–1032. doi:10.1111/j.1420-9101.2006.01091.x.

Bah, J., Quach, H., Ebstein, R. P., Segman, R. H., Melke, J., Jamain, S., et al. (2004). Maternal transmission disequilibrium of the glutamate receptor GRIK2 in schizophrenia. *Neuroreport* 15, 1987–1991. doi:10.1097/00001756-200408260-00031.

Baig, S. M., Koschak, A., Lieb, A., Gebhart, M., Dafinger, C., Nürnberg, G., et al. (2011). Loss of Ca v 1.3 ( CACNA1D ) function in a human channelopathy with bradycardia and congenital deafness. *Nature Neuroscience* 14, 77–84. doi:10.1038/nn.2694.

Baker, D. G., West, S. A., Nicholson, W. E., Ekhator, N. N., Kasckow, J. W., Hill, K. K., et al. (1999). Serial CSF corticotropin-releasing hormone levels and adrenocortical activity in combat veterans with posttraumatic stress disorder. *Am J Psychiatry* 156, 585–588. doi:10.1176/ajp.156.4.585.

Bando, Y., Hirano, T., and Tagawa, Y. (2014). Dysfunction of KCNK Potassium Channels Impairs Neuronal Migration in the Developing Mouse Cerebral Cortex. *Cereb Cortex* 24, 1017–1029. doi:10.1093/cercor/bhs387.

Banerjee, A., Macdonald, M. L., Borgmann-Winter, K. E., and Hahn, C.-G. (2010). Neuregulin 1-erbB4 pathway in schizophrenia: From genes to an interactome. *Brain Res. Bull.* 83, 132–139. doi:10.1016/j.brainresbull.2010.04.011.

Banik, N. L. (1992). Pathogenesis of myelin breakdown in demyelinating diseases: role of proteolytic enzymes. *Crit Rev Neurobiol* 6, 257–271.

Banlaki, Z., Cimarelli, G., Viranyi, Z., Kubinyi, E., Sasvari-Szekely, M., and Ronai, Z. (2017). DNA methylation patterns of behavior-related gene promoter regions dissect the gray wolf from domestic dog breeds. *Mol Genet Genomics* 292, 685–697. doi:10.1007/s00438-017-1305-5.

Barak, B., Zhang, Z., Liu, Y., Nir, A., Trangle, S. S., Ennis, M., et al. (2019). Neuronal deletion of Gtf2i , associated with Williams syndrome, causes behavioral and myelin alterations rescuable by a remyelinating drug. *Nature Neuroscience* 22, 700–708. doi:10.1038/s41593-019-0380-9.

Barbosa, I. G., Huguet, R. B., Sousa, L. P., Abreu, M. N. S., Rocha, N. P., Bauer, M. E., et al. (2011). Circulating levels of GDNF in bipolar disorder. *Neuroscience Letters* 502, 103–106. doi:10.1016/j.neulet.2011.07.031.

Barel, O., Shalev, S. A., Ofir, R., Cohen, A., Zlotogora, J., Shorer, Z., et al. (2008). Maternally Inherited Birk Barel Mental Retardation Dysmorphism Syndrome Caused by a Mutation in the Genomically Imprinted Potassium Channel KCNK9. *Am J Hum Genet* 83, 193–199. doi:10.1016/j.ajhg.2008.07.010.

Barington, M., Risom, L., Ek, J., Uldall, P., and Ostergaard, E. (2018). A recurrent de novo CUX2 missense variant associated with intellectual disability, seizures, and autism spectrum disorder. *Eur J Hum Genet* 26, 1388–1391. doi:10.1038/s41431-018-0184-5.

Barlow, G. M., Lyons, G. E., Richardson, J. A., Sarnat, H. B., and Korenberg, J. R. (2002). DSCAM: an endogenous promoter drives expression in the developing CNS and neural crest. *Biochemical and Biophysical Research Communications* 299, 1–6. doi:10.1016/S0006-291X(02)02548-2.

Barry, H., Byrne, S., Barrett, E., Murphy, K. C., and Cotter, D. R. (2015). Anti-N-methyl-d-aspartate receptor encephalitis: review of clinical presentation, diagnosis and treatment. *BJPsych Bull* 39, 19–23. doi:10.1192/pb.bp.113.045518.

Barson, J. R., Morganstern, I., and Leibowitz, S. F. (2013). Complementary Roles of Orexin and Melanin-Concentrating Hormone in Feeding Behavior. *Int J Endocrinol* 2013. doi:10.1155/2013/983964.

Baruch, K., Silberberg, G., Aviv, A., Shamir, E., Bening-Abu-Shach, U., Baruch, Y., et al. (2009). Association between golli-MBP and schizophrenia in the Jewish Ashkenazi population: are regulatory regions involved? *Int J Neuropsychopharmacol* 12, 885–894. doi:10.1017/S1461145708009887.

Baud, P., Perroud, N., Courtet, P., Jaussent, I., Relecom, C., Jollant, F., et al. (2009). Modulation of anger control in suicide attempters by TPH-1. *Genes Brain Behav.* 8, 97–100. doi:10.1111/j.1601-183X.2008.00451.x.

Baujat, G., and Cormier-Daire, V. (2007). Sotos syndrome. *Orphanet J Rare Dis* 2, 36. doi:10.1186/1750-1172-2-36.

Baum, A. E., Akula, N., Cabanero, M., Cardona, I., Corona, W., Klemens, B., et al. (2008a). A genome-wide association study implicates diacylglycerol kinase eta (DGKH) and several other genes in the etiology of bipolar disorder. *Molecular Psychiatry* 13, 197–207. doi:10.1038/sj.mp.4002012.

Baum, A., Hamshere, M., Green, E., Cichon, S., Rietschel, M., Noethen, M., et al. (2008b). Meta-analysis of two genome-wide association studies of bipolar disorder reveals important points of agreement. *Mol Psychiatry* 13, 466–467. doi:10.1038/mp.2008.16.

Beccavin, C., Chevalier, B., Cogburn, L. A., Simon, J., and Duclos, M. J. (2001). Insulin-like growth factors and body growth in chickens divergently selected for high or low growth rate. *J. Endocrinol.* 168, 297–306. doi:10.1677/joe.0.1680297.

Beden, O., Senol, E., Atay, S., Ak, H., Altintoprak, A. E., Kiyan, G. S., et al. (2016). TPH1 A218 allele is associated with suicidal behavior in Turkish population. *Leg Med (Tokyo)* 21, 15–18. doi:10.1016/j.legalmed.2016.05.005.

Begni, S., Popoli, M., Moraschi, S., Bignotti, S., Tura, G. B., and Gennarelli, M. (2002). Association between the ionotropic glutamate receptor kainate 3 ( GRIK3 ) ser310ala polymorphism and schizophrenia. *Molecular Psychiatry* 7, 416–418. doi:10.1038/sj.mp.4000987.

Bellivier, F., Chaste, P., and Malafosse, A. (2004). Association between the TPH gene A218C polymorphism and suicidal behavior: a meta-analysis. *Am. J. Med. Genet. B Neuropsychiatr. Genet.* 124B, 87–91. doi:10.1002/ajmg.b.20015.

Bellivier, F., Leboyer, M., Courtet, P., Buresi, C., Beaufils, B., Samolyk, D., et al. (1998). Association Between the Tryptophan Hydroxylase Gene and Manic-depressive Illness. *Arch Gen Psychiatry* 55, 33–37. doi:10.1001/archpsyc.55.1.33.

Belvederi Murri, M., Prestia, D., Mondelli, V., Pariante, C., Patti, S., Olivieri, B., et al. (2016). The HPA axis in bipolar disorder: Systematic review and meta-analysis. *Psychoneuroendocrinology* 63, 327–342. doi:10.1016/j.psyneuen.2015.10.014.

Béna, F., Bruno, D. L., Eriksson, M., van Ravenswaaij-Arts, C., Stark, Z., Dijkhuizen, T., et al. (2013). Molecular and clinical characterization of 25 individuals with exonic deletions of NRXN1 and comprehensive review of the literature. *Am. J. Med. Genet. B Neuropsychiatr. Genet.* 162B, 388–403. doi:10.1002/ajmg.b.32148.

Benamer, N., Marti, F., Lujan, R., Hepp, R., Aubier, T. G., Dupin, A. a. M., et al. (2018). GluD1, linked to schizophrenia, controls the burst firing of dopamine neurons. *Mol. Psychiatry* 23, 691–700. doi:10.1038/mp.2017.137.

Benítez-Burraco, A., Pietro, L. D., Barba, M., and Lattanzi, W. (2017). Schizophrenia and Human Self-Domestication: An Evolutionary Linguistics Approach. *BBE* 89, 162–184. doi:10.1159/000468506.

Benítez-Burraco, A., Theofanopoulou, C., and Boeckx, C. (2016). Globularization and Domestication. *Topoi*, 1–14. doi:10.1007/s11245-016-9399-7.

Bennett, M. R., AO (2008). Stress and Anxiety in Schizophrenia and Depression: Glucocorticoids, Corticotropin-Releasing Hormone and Synapse Regression. *Aust N Z J Psychiatry* 42, 995–1002. doi:10.1080/00048670802512073.

Bergen, S. E., Fanous, A. H., Walsh, D., O'Neill, F. A., and Kendler, K. S. (2009). Polymorphisms in SLC6A4, PAH, GABRB3, and MAOB and Modification of Psychotic Disorder Features. *Schizophr Res* 109, 94–97. doi:10.1016/j.schres.2009.02.009.

Berger, S. M., and Bartsch, D. (2014). The role of L-type voltage-gated calcium channels Cav1.2 and Cav1.3 in normal and pathological brain function. *Cell Tissue Res* 357, 463–476. doi:10.1007/s00441-014-1936-3.

Berger, W., Meindl, A., van de Pol, T. J. R., Cremers, F. P. M., Ropers, H. H., Döerner, C., et al. (1992). Isolation of a candidate gene for Norrie disease by positional cloning. *Nature Genetics* 1, 199–203. doi:10.1038/ng0692-199.

Bertelsen, B., Melchior, L., Jensen, L. R., Groth, C., Glenthøj, B., Rizzo, R., et al. (2014). Intragenic deletions affecting two alternative transcripts of the IMMP2L gene in patients with Tourette syndrome. *Eur J Hum Genet* 22, 1283–1289. doi:10.1038/ejhg.2014.24.

Bertolini, F., Gandolfi, B., Kim, E. S., Haase, B., Lyons, L. A., and Rothschild, M. F. (2016). Evidence of selection signatures that shape the Persian cat breed. *Mamm Genome* 27, 144–155. doi:10.1007/s00335-016-9623-1.

Bhambhvani, H. P., Simmons, M., Haroutunian, V., and Meador-Woodruff, J. H. (2016). Decreased Expression of Cortactin in Schizophrenia Brain. *Neuroreport* 27, 145–150. doi:10.1097/WNR.0000000000000514.

Bickeböller, H., Kistler, M., and Scholz, M. (1997). Investigation of the candidate genes ACTHR and golf for bipolar illness by the transmission/disequilibrium test. *Genet. Epidemiol.* 14, 575–580. doi:10.1002/(SICI)1098-2272(1997)14:6<575::AID-GEPI4>3.0.CO;2-Z.

Bigos, K. L., Mattay, V. S., Callicott, J. H., Straub, R. E., Vakkalanka, R., Kolachana, B., et al. (2010). Genetic variation in CACNA1C affects brain circuitries related to mental illness. *Arch Gen Psychiatry* 67, 939–945. doi:10.1001/archgenpsychiatry.2010.96.

Bilgiç, A., Toker, A., Işık, Ü., and Kılınç, İ. (2017). Serum brain-derived neurotrophic factor, glial-derived neurotrophic factor, nerve growth factor, and neurotrophin-3 levels in children with attention-deficit/hyperactivity disorder. *Eur Child Adolesc Psychiatry* 26, 355–363. doi:10.1007/s00787-016-0898-2.

Blatt, G. J., Fitzgerald, C. M., Guptill, J. T., Booker, A. B., Kemper, T. L., and Bauman, M. L. (2001). Density and Distribution of Hippocampal Neurotransmitter Receptors in Autism: An Autoradiographic Study. *J Autism Dev Disord* 31, 537–543. doi:10.1023/A:1013238809666.

Bloch, M., Rubinow, D. R., Schmidt, P. J., Lotsikas, A., Chrousos, G. P., and Cizza, G. (2005). Cortisol response to ovine corticotropin-releasing hormone in a model of pregnancy and parturition in euthymic women with and without a history of postpartum depression. *J. Clin. Endocrinol. Metab.* 90, 695–699. doi:10.1210/jc.2004-1388.

Boeckx, C., and Benítez-Burraco, A. (2014). Globularity and language-readiness: generating new predictions by expanding the set of genes of interest. *Front Psychol* 5. doi:10.3389/fpsyg.2014.01324.

Børglum, A. D., Bruun, T. G., Kjeldsen, T. E., Ewald, H., Mors, O., Kirov, G., et al. (1999). Two novel variants in the DOPA decarboxylase gene: association with bipolar affective disorder. *Mol. Psychiatry* 4, 545–551. doi:10.1038/sj.mp.4000559.

Børglum, A. D., Hampson, M., Kjeldsen, T. E., Muir, W., Murray, V., Ewald, H., et al. (2001). Dopa decarboxylase genotypes may influence age at onset of schizophrenia. *Mol. Psychiatry* 6, 712–717. doi:10.1038/sj.mp.4000902.

Børglum, A. D., Kirov, G., Craddock, N., Mors, O., Muir, W., Murray, V., et al. (2003). Possible parent-of-origin effect of Dopa decarboxylase in susceptibility to bipolar affective disorder. *Am. J. Med. Genet. B Neuropsychiatr. Genet.* 117B, 18–22. doi:10.1002/ajmg.b.10030.

Bortolato, M., Godar, S. C., Alzghoul, L., Zhang, J., Darling, R. D., Simpson, K. L., et al. (2013). Monoamine oxidase A and A/B knockout mice display autistic-like features. *Int. J. Neuropsychopharmacol.* 16, 869–888. doi:10.1017/S1461145712000715.

Boudry-Labis, E., Demeer, B., Le Caignec, C., Isidor, B., Mathieu-Dramard, M., Plessis, G., et al. (2013). A novel microdeletion syndrome at 9q21.13 characterised by mental retardation, speech delay, epilepsy and characteristic facial features. *European Journal of Medical Genetics* 56, 163–170. doi:10.1016/j.ejmg.2012.12.006.

Braisted, J. E., Catalano, S. M., Stimac, R., Kennedy, T. E., Tessier-Lavigne, M., Shatz, C. J., et al. (2000). Netrin-1 Promotes Thalamic Axon Growth and Is Required for Proper Development of the Thalamocortical Projection. *J Neurosci* 20, 5792–5801. doi:10.1523/JNEUROSCI.20-15-05792.2000.

Brambilla, P., Perez, J., Barale, F., Schettini, G., and Soares, J. C. (2003). GABAergic dysfunction in mood disorders. *Mol. Psychiatry* 8, 721–737, 715. doi:10.1038/sj.mp.4001362.

Brett, M., McPherson, J., Zang, Z. J., Lai, A., Tan, E.-S., Ng, I., et al. (2014). Massively Parallel Sequencing of Patients with Intellectual Disability, Congenital Anomalies and/or Autism Spectrum Disorders with a Targeted Gene Panel. *PLoS One* 9. doi:10.1371/journal.pone.0093409.

Bruce, F. M., Brown, S., Smith, J. N., Fuerst, P. G., and Erskine, L. (2017). DSCAM promotes axon fasciculation and growth in the developing optic pathway. *Proc. Natl. Acad. Sci. U.S.A.* 114, 1702–1707. doi:10.1073/pnas.1618606114.

Brunner, H. G., Nelen, M., Breakefield, X. O., Ropers, H. H., and Oost, B. A. (1993). Abnormal Behavior Associated with a Point Mutation in the Structural Gene for Monoamine Oxidase A. *Science (New York, N.Y.)* 262, 578–80. doi:10.1126/science.8211186.

Brusini, I., Carneiro, M., Wang, C., Rubin, C.-J., Ring, H., Afonso, S., et al. (2018). Changes in brain architecture are consistent with altered fear processing in domestic rabbits. *Proc Natl Acad Sci U S A* 115, 7380–7385. doi:10.1073/pnas.1801024115.

Burri, L., Strahm, Y., Hawkins, C. J., Gentle, I. E., Puryer, M. A., Verhagen, A., et al. (2005). Mature DIABLO/Smac Is Produced by the IMP Protease Complex on the Mitochondrial Inner Membrane. *MBoC* 16, 2926–2933. doi:10.1091/mbc.e04-12-1086.

Butler, K. M., Moody, O. A., Schuler, E., Coryell, J., Alexander, J. J., Jenkins, A., et al. (2018). De novo variants in GABRA2 and GABRA5 alter receptor function and contribute to early-onset epilepsy. *Brain* 141, 2392–2405. doi:10.1093/brain/awy171.

Buxbaum, J. D., Cai, G., Chaste, P., Nygren, G., Goldsmith, J., Reichert, J., et al. (2007). Mutation Screening of the PTEN Gene in Patients With Autism Spectrum Disorders and Macrocephaly. *Am J Med Genet B Neuropsychiatr Genet* 144B, 484–491. doi:10.1002/ajmg.b.30493.

C Yuen, R. K., Merico, D., Bookman, M., L Howe, J., Thiruvahindrapuram, B., Patel, R. V., et al. (2017). Whole genome sequencing resource identifies 18 new candidate genes for autism spectrum disorder. *Nat. Neurosci.* 20, 602–611. doi:10.1038/nn.4524.

Cagan, A., and Blass, T. (2016). Identification of genomic variants putatively targeted by selection during dog domestication. *BMC Evol Biol* 16. doi:10.1186/s12862-015-0579-7.

Calvey, T. (2019). Human Self-Domestication and the Extended Evolutionary Synthesis of Addiction: How Humans Evolved a Unique Vulnerability. *Neuroscience* 419, 100–107. doi:10.1016/j.neuroscience.2019.09.013.

Cao, S.-X., Zhang, Y., Hu, X.-Y., Hong, B., Sun, P., He, H.-Y., et al. (2018). ErbB4 deletion in noradrenergic neurons in the locus coeruleus induces mania-like behavior via elevated catecholamines. *eLife* 7. doi:10.7554/eLife.39907.

Carneiro, M., Piorno, V., Rubin, C.-J., Alves, J. M., Ferrand, N., Alves, P. C., et al. (2015). Candidate genes underlying heritable differences in reproductive seasonality between wild and domestic rabbits. *Animal Genetics* 46, 418–425. doi:10.1111/age.12299.

Carneiro, M., Rubin, C.-J., Palma, F. D., Albert, F. W., Alföldi, J., Barrio, A. M., et al. (2014). Rabbit genome analysis reveals a polygenic basis for phenotypic change during domestication. *Science* 345, 1074–1079. doi:10.1126/science.1253714.

Carrard, A., Salzmann, A., Perroud, N., Gafner, J., Malafosse, A., and Karege, F. (2011). Genetic association of the Phosphoinositide-3 kinase in schizophrenia and bipolar disorder and interaction with a BDNF gene polymorphism. *Brain Behav* 1, 119–124. doi:10.1002/brb3.23.

Carrera, N., Sanjuán, J., Moltó, M. D., Carracedo, A., and Costas, J. (2009). Recent adaptive selection at MAOB and ancestral susceptibility to schizophrenia. *Am. J. Med. Genet. B Neuropsychiatr. Genet.* 150B, 369–374. doi:10.1002/ajmg.b.30823.

Carson, M. J., Behringer, R. R., Brinster, R. L., and McMorris, F. A. (1993). Insulin-like growth factor I increases brain growth and central nervous system myelination in tTransgenic mice. *Neuron* 10, 729–740. doi:10.1016/0896-6273(93)90173-O.

Carter, C. J. (2007). EIF2B and Oligodendrocyte Survival: Where Nature and Nurture Meet in Bipolar Disorder and Schizophrenia? *Schizophr Bull* 33, 1343–1353. doi:10.1093/schbul/sbm007.

Cartier, D., Lihrmann, I., Parmentier, F., Bastard, C., Bertherat, J., Caron, P., et al. (2003). Overexpression of serotonin4 receptors in cisapride-responsive adrenocorticotropin-independent bilateral macronodular adrenal hyperplasia causing Cushing's syndrome. *J. Clin. Endocrinol. Metab.* 88, 248–254. doi:10.1210/jc.2002-021107.

Cases, O., Seif, I., Grimsby, J., Gaspar, P., Chen, K., Pournin, S., et al. (1995). Aggressive Behavior and Altered Amounts of Brain Serotonin and Norepinephrine in Mice Lacking MAOA. *Science* 268, 1763–1766.

Caspi, A., McClay, J., Moffitt, T. E., Mill, J., Martin, J., Craig, I. W., et al. (2002). Role of Genotype in the Cycle of Violence in Maltreated Children. *Science* 297, 851–854. doi:10.1126/science.1072290.

Ceulemans, S., Zutter, S. D., Heyrman, L., Norrback, K.-F., Nordin, A., Nilsson, L.-G., et al. (2011). Evidence for the involvement of the glucocorticoid receptor gene in bipolar disorder in an isolated northern Swedish population. *Bipolar Disorders* 13, 614–623. doi:10.1111/j.1399-5618.2011.00960.x.

Chakraborti, B., Verma, D., Karmakar, A., Jaiswal, P., Sanyal, A., Paul, D., et al. (2016). Genetic variants of MAOB affect serotonin level and specific behavioral attributes to increase autism spectrum disorder (ASD) susceptibility in males. *Prog. Neuropsychopharmacol. Biol. Psychiatry* 71, 123–136. doi:10.1016/j.pnpbp.2016.07.001.

Chamberlain, R. S., and Herman, B. H. (1990). A novel biochemical model linking dysfunctions in brain melatonin, proopiomelanocortin peptides, and serotonin in autism. *Biol. Psychiatry* 28, 773–793. doi:10.1016/0006-3223(90)90513-2.

Chambers, J. S., and Perrone-Bizzozero, N. I. (2004). Altered myelination of the hippocampal formation in subjects with schizophrenia and bipolar disorder. *Neurochem. Res.* 29, 2293–2302. doi:10.1007/s11064-004-7039-x.

Chang, H., Cahill, H., Smallwood, P. M., Wang, Y., and Nathans, J. (2015). Identification of Astrotactin2 as a Genetic Modifier That Regulates the Global Orientation of Mammalian Hair Follicles. *PLoS Genet* 11. doi:10.1371/journal.pgen.1005532.

Chatron, N., Møller, R. S., Champaigne, N. L., Schneider MGenCouns, A. L., Kuechler, A., Labalme, A., et al. (2018). The epilepsy phenotypic spectrum associated with a recurrent CUX2 variant. *Ann Neurol* 83, 926–934. doi:10.1002/ana.25222.

Chee, M. J. S., Pissios, P., Prasad, D., and Maratos-Flier, E. (2014). Expression of Melanin-Concentrating Hormone Receptor 2 Protects Against Diet-Induced Obesity in Male Mice. *Endocrinology* 155, 81–88. doi:10.1210/en.2013-1738.

Chen, D., Liu, F., Yang, C., Liang, X., Shang, Q., He, W., et al. (2012a). Association between the TPH1 A218C polymorphism and risk of mood disorders and alcohol dependence: evidence from the current studies. *J Affect Disord* 138, 27–33. doi:10.1016/j.jad.2011.04.018.

Chen, L., Liu, T., Tu, Y., Rong, D., and Cao, Y. (2016). Cul1 promotes melanoma cell proliferation by promoting DEPTOR degradation and enhancing cap-dependent translation. *Oncol. Rep.* 35, 1049–1056. doi:10.3892/or.2015.4442.

Chen, P., Chen, J., Huang, K., Ji, W., Wang, T., Li, T., et al. (2012b). Analysis of association between common SNPs in ErbB4 and bipolar affective disorder, major depressive disorder and schizophrenia in the Han Chinese population. *Prog. Neuropsychopharmacol. Biol. Psychiatry* 36, 17–21. doi:10.1016/j.pnpbp.2011.09.011.

Chen, X., Wang, J., Qian, L., Gaughan, S., Xiang, W., Ai, T., et al. (2017). Domestication drive the changes of immune and digestive system of Eurasian perch (Perca fluviatilis). *PLoS One* 12. doi:10.1371/journal.pone.0172903.

Chen, Y., Andres, A. L., Frotscher, M., and Baram, T. Z. (2012c). Tuning synaptic transmission in the hippocampus by stress: the CRH system. *Front Cell Neurosci* 6, 13. doi:10.3389/fncel.2012.00013.

Chen, Y., Zhang, J., Zhang, L., Shen, Y., and Xu, Q. (2012d). Effects of MAOA promoter methylation on susceptibility to paranoid schizophrenia. *Hum. Genet.* 131, 1081–1087. doi:10.1007/s00439-011-1131-5.

Chen, Y.-C., Liu, Y.-L., Tsai, S.-J., Kuo, P.-H., Huang, S.-S., and Lee, Y.-S. (2019). LRRTM4 and PCSK5 Genetic Polymorphisms as Markers for Cognitive Impairment in A Hypotensive Aging Population: A Genome-Wide Association Study in Taiwan. *J Clin Med* 8. doi:10.3390/jcm8081124.

Chen, Z. Y., Denney, R. M., and Breakefield, X. O. (1995). Norrie disease and MAO genes: nearest neighbors. *Hum. Mol. Genet.* 4 Spec No, 1729–1737. doi:10.1093/hmg/4.suppl_1.1729.

Cheng, R., Juo, S. H., Loth, J. E., Nee, J., Iossifov, I., Blumenthal, R., et al. (2006). Genome-wide linkage scan in a large bipolar disorder sample from the National Institute of Mental Health genetics initiative suggests putative loci for bipolar disorder, psychosis, suicide, and panic disorder. *Mol. Psychiatry* 11, 252–260. doi:10.1038/sj.mp.4001778.

Cherlyn, S. Y. T., Woon, P. S., Liu, J. J., Ong, W. Y., Tsai, G. C., and Sim, K. (2010). Genetic association studies of glutamate, GABA and related genes in schizophrenia and bipolar disorder: A decade of advance. *Neuroscience & Biobehavioral Reviews* 34, 958–977. doi:10.1016/j.neubiorev.2010.01.002.

Cheslack-Postava, K., Fallin, M. D., Avramopoulos, D., Connors, S. L., Zimmerman, A. W., Eberhart, C. G., et al. (2007). beta2-Adrenergic receptor gene variants and risk for autism in the AGRE cohort. *Mol. Psychiatry* 12, 283–291. doi:10.1038/sj.mp.4001940.

Ching, M. S. L., Shen, Y., Tan, W.-H., Jeste, S. S., Morrow, E. M., Chen, X., et al. (2010). Deletions of NRXN1 (neurexin-1) predispose to a wide spectrum of developmental disorders. *Am. J. Med. Genet. B Neuropsychiatr. Genet.* 153B, 937–947. doi:10.1002/ajmg.b.31063.

Chitramuthu, B. P., Baranowski, D. C., Cadieux, B., Rousselet, E., Seidah, N. G., and Bennett, H. P. J. (2010). Molecular cloning and embryonic expression of zebrafish PCSK5 co-orthologues:

functional assessment during lateral line development. *Dev. Dyn.* 239, 2933–2946. doi:10.1002/dvdy.22426.

Chobotská, K., Arnold, M., Werner, P., and Pliska, V. (1998). A rapid assay for tyrosine hydroxylase activity, an indicator of chronic stress in laboratory and domestic animals. *Biol. Chem.* 379, 59–63. doi:10.1515/bchm.1998.379.1.59.

Choi, K.-Y., Yoon, H.-K., and Kim, Y.-K. (2010). Association between Serotonin-Related Polymorphisms in 5HT2A, TPH1, TPH2 Genes and Bipolar Disorder in Korean Population. *Psychiatry Investig* 7, 60–67. doi:10.4306/pi.2010.7.1.60.

Chourbaji, S., Vogt, M. A., and Gass, P. (2008). Mice that under- or overexpress glucocorticoid receptors as models for depression or posttraumatic stress disorder. *Prog. Brain Res.* 167, 65–77. doi:10.1016/S0079-6123(07)67005-8.

Chow, M. L., Pramparo, T., Winn, M. E., Barnes, C. C., Li, H.-R., Weiss, L., et al. (2012). Age-Dependent Brain Gene Expression and Copy Number Anomalies in Autism Suggest Distinct Pathological Processes at Young Versus Mature Ages. *PLoS Genet* 8. doi:10.1371/journal.pgen.1002592.

Christophersen, N. S., Grønborg, M., Petersen, T. N., Fjord-Larsen, L., Jørgensen, J. R., Juliusson, B., et al. (2007). Midbrain expression of Delta-like 1 homologue is regulated by GDNF and is associated with dopaminergic differentiation. *Exp. Neurol.* 204, 791–801. doi:10.1016/j.expneurol.2007.01.014.

Cieslak, M., Reissmann, M., Hofreiter, M., and Ludwig, A. (2011). Colours of domestication. *Biological Reviews* 86, 885–899. doi:10.1111/j.1469-185X.2011.00177.x.

Cipriani, A., Saunders, K., Attenburrow, M.-J., Stefaniak, J., Panchal, P., Stockton, S., et al. (2016). A systematic review of calcium channel antagonists in bipolar disorder and some considerations for their future development. *Mol Psychiatry* 21, 1324–1332. doi:10.1038/mp.2016.86.

Clark, A. J., McLoughlin, L., and Grossman, A. (1993). Familial glucocorticoid deficiency associated with point mutation in the adrenocorticotropin receptor. *Lancet* 341, 461–462. doi:10.1016/0140-6736(93)90208-x.

Clements, C. C., Wenger, T. L., Zoltowski, A. R., Bertollo, J. R., Miller, J. S., de Marchena, A. B., et al. (2017). Critical region within 22q11.2 linked to higher rate of autism spectrum disorder. *Mol Autism* 8. doi:10.1186/s13229-017-0171-7.

Coene, K. L. M., Mans, D. A., Boldt, K., Gloeckner, C. J., van Reeuwijk, J., Bolat, E., et al. (2011). The ciliopathy-associated protein homologs RPGRIP1 and RPGRIP1L are linked to cilium integrity through interaction with Nek4 serine/threonine kinase. *Hum. Mol. Genet.* 20, 3592–3605. doi:10.1093/hmg/ddr280.

Collins, F. A., Murphy, D. L., Reiss, A. L., Sims, K. B., Lewis, J. G., Freund, L., et al. (1992). Clinical, biochemical, and neuropsychiatric evaluation of a patient with a contiguous gene syndrome due to a microdeletion Xp11.3 including the Norrie disease locus and monoamine oxidase (MAOA and MAOB) genes. *Am. J. Med. Genet.* 42, 127–134. doi:10.1002/ajmg.1320420126.

Compton, M. T., Celentana, M., Price, B., and Furman, A. C. (2004). A Case of Sotos Syndrome (Cerebral Gigantism) and Psychosis. *PSP* 37, 190–193. doi:10.1159/000079510.

Comunanza, V., Marcantoni, A., Vandael, D. H., Mahapatra, S., Gavello, D., Carabelli, V., et al. (2010). CaV1.3 as pacemaker channels in adrenal chromaffin cells: specific role on exo- and endocytosis? *Channels (Austin)* 4, 440–446. doi:10.4161/chan.4.6.12866.

Connors, S. L., Crowell, D. E., Eberhart, C. G., Copeland, J., Newschaffer, C. J., Spence, S. J., et al. (2005). beta2-adrenergic receptor activation and genetic polymorphisms in autism: data from dizygotic twins. *J. Child Neurol.* 20, 876–884. doi:10.1177/08830738050200110401.

Consoni, F. T., Vital, M. A. B. F., and Andreatini, R. (2006). Dual monoamine modulation for the antidepressant-like effect of lamotrigine in the modified forced swimming test. *European Neuropsychopharmacology* 16, 451–458. doi:10.1016/j.euroneuro.2006.01.003.

Cotter, D., Landau, S., Beasley, C., Stevenson, R., Chana, G., MacMillan, L., et al. (2002). The density and spatial distribution of gabaergic neurons, labelled using calcium binding proteins, in the anterior cingulate cortex in major depressive disorder, bipolar disorder, and schizophrenia. *Biological Psychiatry* 51, 377–386. doi:10.1016/S0006-3223(01)01243-4.

Coulpier, M., Anders, J., and Ibáñez, C. F. (2002). Coordinated activation of autophosphorylation sites in the RET receptor tyrosine kinase: importance of tyrosine 1062 for GDNF mediated neuronal differentiation and survival. *J. Biol. Chem.* 277, 1991–1999. doi:10.1074/jbc.M107992200.

Craddock, N., O'Donovan, M., and Owen, M. (2005). The genetics of schizophrenia and bipolar disorder: dissecting psychosis. *J Med Genet* 42, 193–204. doi:10.1136/jmg.2005.030718.

Crespi, B., and Badcock, C. (2008). Psychosis and autism as diametrical disorders of the social brain. *Behav Brain Sci* 31, 241–261; discussion 261-320. doi:10.1017/S0140525X08004214.

Crespi, B., Stead, P., and Elliot, M. (2010). Comparative genomics of autism and schizophrenia. *Proc Natl Acad Sci U S A* 107, 1736–1741. doi:10.1073/pnas.0906080106.

Cross-Disorder Group of the Psychiatric Genomics Consortium (2013). Identification of risk loci with shared effects on five major psychiatric disorders: a genome-wide analysis. *Lancet* 381, 1371–1379. doi:10.1016/S0140-6736(12)62129-1.

Cubelos, B., Sebastián-Serrano, A., Beccari, L., Calcagnotto, M. E., Cisneros, E., Kim, S., et al. (2010). Cux1 and Cux2 regulate dendritic branching, spine morphology and synapses of the upper layer neurons of the cortex. *Neuron* 66, 523–535. doi:10.1016/j.neuron.2010.04.038.

Cummings, A. C., Jiang, L., Velez Edwards, D. R., McCauley, J. L., Laux, R., McFarland, L. L., et al. (2012). Genome-wide association and linkage study in the Amish detects a novel candidate late-onset Alzheimer disease gene. *Ann Hum Genet* 76, 342–351. doi:10.1111/j.1469-1809.2012.00721.x.

Cunha, P. da S., Pena, H. B., D'Angelo, C. S., Koiffmann, C. P., Rosenfeld, J. A., Shaffer, L. G., et al. (2014). Accurate, Fast and Cost-Effective Diagnostic Test for Monosomy 1p36 Using Real-Time Quantitative PCR. *Dis Markers* 2014. doi:10.1155/2014/836082.

Curran, S., Ahn, J. W., Grayton, H., Collier, D. A., and Ogilvie, C. M. (2013). NRXN1 deletions identified by array comparative genome hybridisation in a clinical case series – further understanding of the relevance of NRXN1 to neurodevelopmental disorders. *J Mol Psychiatry* 1. doi:10.1186/2049-9256-1-4.

Curtis, D., Vine, A., McQuillin, A., Bass, N., Pereira, A., Kandaswamy, R., et al. (2011). Case-case genome wide association analysis reveals markers differentially associated with schizophrenia and bipolar disorder and implicates calcium channel genes. *Psychiatr Genet* 21, 1–4. doi:10.1097/YPG.0b013e3283413382.

Cuscó, I., Corominas, R., Bayés, M., Flores, R., Rivera-Brugués, N., Campuzano, V., et al. (2008). Copy number variation at the 7q11.23 segmental duplications is a susceptibility factor for the Williams-Beuren syndrome deletion. *Genome Res.* 18, 683–694. doi:10.1101/gr.073197.107.

da Silva, E. G., Pfaffenseller, B., Walz, J., Stertz, L., Fries, G., Rosa, A. R., et al. (2017). Peripheral insulin-like growth factor 1 in bipolar disorder. *Psychiatry Res* 250, 30–34. doi:10.1016/j.psychres.2017.01.061.

Dachtler, J., Ivorra, J. L., Rowland, T. E., Lever, C., Rodgers, R. J., and Clapcote, S. J. (2015). Heterozygous Deletion of α-Neurexin I or α-Neurexin II Results in Behaviors Relevant to Autism and Schizophrenia. *Behav Neurosci* 129, 765–776. doi:10.1037/bne0000108.

Dahia, P. L., Toledo, S. P., Mulligan, L. M., Maher, E. R., Grossman, A. B., and Eng, C. (1997). Mutation analysis of glial cell line-derived neurotrophic factor (GDNF), a ligand for the RET/GDNF receptor alpha complex, in sporadic phaeochromocytomas. *Cancer Res.* 57, 310–313.

Daly, C. J., and McGrath, J. C. (2011). Previously unsuspected widespread cellular and tissue distribution of β-adrenoceptors and its relevance to drug action. *Trends Pharmacol Sci*. 32(4), 219-26.

Daly, R. J. (2004). Cortactin signalling and dynamic actin networks. *Biochem. J.* 382, 13–25. doi:10.1042/BJ20040737.

Damaj, L., Lupien-Meilleur, A., Lortie, A., Riou, É., Ospina, L. H., Gagnon, L., et al. (2015). CACNA1A haploinsufficiency causes cognitive impairment, autism and epileptic encephalopathy with mild cerebellar symptoms. *Eur J Hum Genet* 23, 1505–1512. doi:10.1038/ejhg.2015.21.

Dann, J., DeLisi, L. E., Devoto, M., Laval, S., Nancarrow, D. J., Shields, G., et al. (1997). A linkage study of schizophrenia to markers within Xp11 near the MAOB gene. *Psychiatry Res* 70, 131–143. doi:10.1016/s0165-1781(97)03138-7.

Das, M., Bhowmik, A. D., Sinha, S., Chattopadhyay, A., Chaudhuri, K., Singh, M., et al. (2006). MAOA promoter polymorphism and attention deficit hyperactivity disorder (ADHD) in indian children. *Am. J. Med. Genet. B Neuropsychiatr. Genet.* 141B, 637–642. doi:10.1002/ajmg.b.30385.

Davila, S., Furu, L., Gharavi, A. G., Tian, X., Onoe, T., Qian, Q., et al. (2004). Mutations in SEC63 cause autosomal dominant polycystic liver disease. *Nat. Genet.* 36, 575–577. doi:10.1038/ng1357.

Davis, K. L., Stewart, D. G., Friedman, J. I., Buchsbaum, M., Harvey, P. D., Hof, P. R., et al. (2003). White matter changes in schizophrenia: evidence for myelin-related dysfunction. *Arch. Gen. Psychiatry* 60, 443–456. doi:10.1001/archpsyc.60.5.443.

de Ligt, J., Willemsen, M. H., van Bon, B. W. M., Kleefstra, T., Yntema, H. G., Kroes, T., et al. (2012). Diagnostic exome sequencing in persons with severe intellectual disability. *N. Engl. J. Med.* 367, 1921–1929. doi:10.1056/NEJMoa1206524.

De Rubeis, S., He, X., Goldberg, A. P., Poultney, C. S., Samocha, K., Cicek, A. E., et al. (2014). Synaptic, transcriptional and chromatin genes disrupted in autism. *Nature* 515, 209–215. doi:10.1038/nature13772.

Debette, S., Ibrahim Verbaas, C. A., Bressler, J., Schuur, M., Smith, A., Bis, J. C., et al. (2015). Genome-wide Studies of Verbal Declarative Memory in Nondemented Older People: The Cohorts for Heart and Aging Research in Genomic Epidemiology Consortium. *Biol Psychiatry* 77, 749–763. doi:10.1016/j.biopsych.2014.08.027.

Dedic, N., Pöhlmann, M. L., Richter, J. S., Mehta, D., Czamara, D., Metzger, M. W., et al. (2018). Cross-disorder risk gene CACNA1C differentially modulates susceptibility to psychiatric disorders during development and adulthood. *Mol. Psychiatry* 23, 533–543. doi:10.1038/mp.2017.133.

Deitz, S. L. (2013). Molecular Basis and Modification of a Neural Crest Deficit in a Down Syndrome Mouse Model. Available at: https://scholarworks.iupui.edu/handle/1805/3354 [Accessed March 8, 2020].

Delahanty, R. J., Zhang, Y., Bichell, T. J., Shen, W., Verdier, K., Macdonald, R. L., et al. (2016). Beyond Epilepsy and Autism: Disruption of GABRB3 Causes Ocular Hypopigmentation. *Cell Rep* 17, 3115–3124. doi:10.1016/j.celrep.2016.11.067.

Deng, P.-Y., Xiao, Z., Yang, C., Rojanathammanee, L., Grisanti, L., Watt, J., et al. (2009). GABA(B) receptor activation inhibits neuronal excitability and spatial learning in the entorhinal cortex by activating TREK-2 K+ channels. *Neuron* 63, 230–243. doi:10.1016/j.neuron.2009.06.022.

Detera-Wadleigh, S. D., Yoon, S. W., Berrettini, W. H., Goldin, L. R., Turner, G., Yoshikawa, T., et al. (1995). Adrenocorticotropin receptor/melanocortin receptor-2 maps within a reported susceptibility region for bipolar illness on chromosome 18. *Am. J. Med. Genet.* 60, 317–321. doi:10.1002/ajmg.1320600411.

Devlin, R. H., Sakhrani, D., Tymchuk, W. E., Rise, M. L., and Goh, B. (2009). Domestication and growth hormone transgenesis cause similar changes in gene expression in coho salmon (Oncorhynchus kisutch). *Proc Natl Acad Sci U S A* 106, 3047–3052. doi:10.1073/pnas.0809798106.

D'Gama, A. M., Pochareddy, S., Li, M., Jamuar, S. S., Reiff, R. E., Lam, A.-T. N., et al. (2015). Targeted DNA Sequencing from Autism Spectrum Disorder Brains Implicates Multiple Genetic Mechanisms. *Neuron* 88, 910–917. doi:10.1016/j.neuron.2015.11.009.

Dick, D. M., Foroud, T., Flury, L., Bowman, E. S., Miller, M. J., Rau, N. L., et al. (2003). Genomewide linkage analyses of bipolar disorder: a new sample of 250 pedigrees from the National Institute of Mental Health Genetics Initiative. *Am. J. Hum. Genet.* 73, 107–114. doi:10.1086/376562.

Dickey, A. S., Pineda, V. V., Tsunemi, T., Liu, P. P., Miranda, H. C., Gilmore-Hall, S. K., et al. (2016). PPAR-δ is repressed in Huntington's disease, is required for normal neuronal function and can be targeted therapeutically. *Nat. Med.* 22, 37–45. doi:10.1038/nm.4003.

Di-Poï, N., Michalik, L., Desvergne, B., and Wahli, W. (2004). Functions of peroxisome proliferator-activated receptors (PPAR) in skin homeostasis. *Lipids* 39, 1093–1099. doi:10.1007/s11745-004-1335-y.

Dlugos, A. M., Palmer, A. A., and de Wit, H. (2009). Negative emotionality: monoamine oxidase B gene variants modulate personality traits in healthy humans. *J Neural Transm (Vienna)* 116, 1323–1334. doi:10.1007/s00702-009-0281-2.

Dong, Y., Zhang, X., Xie, M., Arefnezhad, B., Wang, Z., Wang, W., et al. (2015). Reference genome of wild goat (capra aegagrus) and sequencing of goat breeds provide insight into genic basis of goat domestication. *BMC Genomics* 16. doi:10.1186/s12864-015-1606-1.

Dow, D. J., Huxley-Jones, J., Hall, J. M., Francks, C., Maycox, P. R., Kew, J. N. C., et al. (2011). ADAMTSL3 as a candidate gene for schizophrenia: Gene sequencing and ultra-high density association analysis by imputation. *Schizophrenia Research* 127, 28–34. doi:10.1016/j.schres.2010.12.009.

Duffy, A., Goodday, S. M., Keown-Stoneman, C., Scotti, M., Maitra, M., Nagy, C., et al. (2019). Epigenetic markers in inflammation-related genes associated with mood disorder: a cross-sectional and longitudinal study in high-risk offspring of bipolar parents. *Int J Bipolar Disord* 7. doi:10.1186/s40345-019-0152-1.

Dygalo, N. N., Bykova, T. S., and Naumenko, E. V. (1988). [Brain tyrosine hydroxylase activity in behavior-selected silver foxes]. *Zh. Evol. Biokhim. Fiziol.* 24, 503–508.

Eastwood, S. L., McDonald, B., Burnet, P. W., Beckwith, J. P., Kerwin, R. W., and Harrison, P. J. (1995). Decreased expression of mRNAs encoding non-NMDA glutamate receptors GluR1 and GluR2 in medial temporal lobe neurons in schizophrenia. *Brain Res. Mol. Brain Res.* 29, 211–223. doi:10.1016/0169-328x(94)00247-c.

Ehrlichman, R. S., Luminais, S. N., White, S. L., Rudnick, N. D., Ma, N., Dow, H. C., et al. (2009). Neuregulin 1 transgenic mice display reduced mismatch negativity, contextual fear conditioning and social interactions. *Brain Res.* 1294, 116–127. doi:10.1016/j.brainres.2009.07.065.

Eo, J., Lee, H.-E., Nam, G.-H., Kwon, Y.-J., Choi, Y., Choi, B.-H., et al. (2016). Association of DNA methylation and monoamine oxidase A gene expression in the brains of different dog breeds. *Gene* 580, 177–182. doi:10.1016/j.gene.2016.01.022.

Eslami Amirabadi, M. R., Rajezi Esfahani, S., Davari-Ashtiani, R., Khademi, M., Emamalizadeh, B., Movafagh, A., et al. (2015). Monoamine Oxidase A Gene Polymorphisms and Bipolar Disorder in Iranian Population. *Iran Red Crescent Med J* 17. doi:10.5812/ircmj.23095.

Essalmani, R., Zaid, A., Marcinkiewicz, J., Chamberland, A., Pasquato, A., Seidah, N. G., et al. (2008). In vivo functions of the proprotein convertase PC5/6 during mouse development: Gdf11 is a likely substrate. *Proc. Natl. Acad. Sci. U.S.A.* 105, 5750–5755. doi:10.1073/pnas.0709428105.

Eusebi, P. G., Sevane, N., Cortés, O., Contreras, E., Cañon, J., and Dunner, S. (2020). Aggressive behavior in cattle is associated with a polymorphism in the MAOA gene promoter. *Animal Genetics* 51, 14–21. doi:10.1111/age.12867.

Fallahsharoudi, A., de Kock, N., Johnsson, M., Ubhayasekera, S. J. K. A., Bergquist, J., Wright, D., et al. (2015). Domestication Effects on Stress Induced Steroid Secretion and Adrenal Gene Expression in Chickens. *Sci Rep* 5. doi:10.1038/srep15345.

Fallin, M. D., Lasseter, V. K., Avramopoulos, D., Nicodemus, K. K., Wolyniec, P. S., McGrath, J. A., et al. (2005). Bipolar I Disorder and Schizophrenia: A 440–Single-Nucleotide Polymorphism Screen of 64 Candidate Genes among Ashkenazi Jewish Case-Parent Trios. *Am J Hum Genet* 77, 918–936.

Fan, M., Liu, B., Jiang, T., Jiang, X., Zhao, H., and Zhang, J. (2010). Meta-analysis of the association between the monoamine oxidase-A gene and mood disorders. *Psychiatr. Genet.* 20, 1–7. doi:10.1097/YPG.0b013e3283351112.

Fan, Y.-C., Zhu, Y.-S., Mei, P.-J., Sun, S.-G., Zhang, H., Chen, H.-F., et al. (2014). Cullin1 regulates proliferation, migration and invasion of glioma cells. *Med. Oncol.* 31, 227. doi:10.1007/s12032-014-0227-x.

Fawcett, J., Busch, K. A., Jacobs, D., Kravitz, H. M., and Fogg, L. (1997). Suicide: a four-pathway clinical-biochemical model. *Ann. N. Y. Acad. Sci.* 836, 288–301. doi:10.1111/j.1749-6632.1997.tb52366.x.

Feder, J., Gurling, H. M., Darby, J., and Cavalli-Sforza, L. L. (1985). DNA restriction fragment analysis of the proopiomelanocortin gene in schizophrenia and bipolar disorders. *Am J Hum Genet* 37, 286–294.

Feng, J., Schroer, R., Yan, J., Song, W., Yang, C., Bockholt, A., et al. (2006). High frequency of neurexin 1beta signal peptide structural variants in patients with autism. *Neurosci. Lett.* 409, 10–13. doi:10.1016/j.neulet.2006.08.017.

Ferensztajn-Rochowiak, E., Kaczmarek, M., Wójcicka, M., Kaufman-Szukalska, E., Dziuda, S., Remlinger-Molenda, A., et al. (2019). Glutamate-Related Antibodies and Peripheral Insulin-Like Growth Factor in Bipolar Disorder and Lithium Prophylaxis. *Neuropsychobiology* 77, 49–56. doi:10.1159/000493740.

Fernandes, B. S., Gama, C. S., Ceresér, K. M., Yatham, L. N., Fries, G. R., Colpo, G., et al. (2011). Brain-derived neurotrophic factor as a state-marker of mood episodes in bipolar disorders: a systematic review and meta-regression analysis. *J Psychiatr Res* 45, 995–1004. doi:10.1016/j.jpsychires.2011.03.002.

Fernandez, S., Fernandez, A. M., Lopez-Lopez, C., and Torres-Aleman, I. (2007). Emerging roles of insulin-like growth factor-I in the adult brain. *Growth Hormone & IGF Research* 17, 89–95. doi:10.1016/j.ghir.2007.01.006.

Ferreira, M. A. R., O'Donovan, M. C., Meng, Y. A., Jones, I. R., Ruderfer, D. M., Jones, L., et al. (2008). Collaborative genome-wide association analysis supports a role for ANK3 and CACNA1C in bipolar disorder. *Nature Genetics* 40, 1056–1058. doi:10.1038/ng.209.

Finci, L., Zhang, Y., Meijers, R., and Wang, J.-H. (2015). Signaling mechanism of the netrin-1 receptor DCC in axon guidance. *Prog. Biophys. Mol. Biol.* 118, 153–160. doi:10.1016/j.pbiomolbio.2015.04.001.

Firth, H. V., Richards, S. M., Bevan, A. P., Clayton, S., Corpas, M., Rajan, D., et al. (2009). DECIPHER: Database of Chromosomal Imbalance and Phenotype in Humans Using Ensembl Resources. *The American Journal of Human Genetics* 84, 524–533. doi:10.1016/j.ajhg.2009.03.010.

Folks, D., and Arnold, E. S. (1983). Pargyline-induced mania in primary affective disorder: case report. *J Clin Psychiatry* 44, 25–26.

Frank, S. M. (1978). Psycholinguistic findings in gilles de la Tourette syndrome. *J Commun Disord* 11, 349–363. doi:10.1016/0021-9924(78)90043-6.

Freedman, A. H., Schweizer, R. M., Vecchyo, D. O.-D., Han, E., Davis, B. W., Gronau, I., et al. (2016). Demographically-Based Evaluation of Genomic Regions under Selection in Domestic Dogs. *PLOS Genetics* 12, e1005851. doi:10.1371/journal.pgen.1005851.

Friedmacher, F., and Puri, P. (2013). Hirschsprung's disease associated with Down syndrome: a meta-analysis of incidence, functional outcomes and mortality. *Pediatr. Surg. Int.* 29, 937–946. doi:10.1007/s00383-013-3361-1.

Friedrich, S. R., Lovell, P. V., Kaser, T. M., and Mello, C. V. (2019). Exploring the molecular basis of neuronal excitability in a vocal learner. *BMC Genomics* 20. doi:10.1186/s12864-019-5871-2.

Fuchs-Telem, D., Stewart, H., Rapaport, D., Nousbeck, J., Gat, A., Gini, M., et al. (2011). CEDNIK syndrome results from loss-of-function mutations in SNAP29. *Br. J. Dermatol.* 164, 610–616. doi:10.1111/j.1365-2133.2010.10133.x.

Fujii, T., Uchiyama, H., Yamamoto, N., Hori, H., Tatsumi, M., Ishikawa, M., et al. (2011). Possible association of the semaphorin 3D gene (SEMA3D) with schizophrenia. *Journal of Psychiatric Research* 45, 47–53. doi:10.1016/j.jpsychires.2010.05.004.

Furlong, R. A., Rubinsztein, J. S., Ho, L., Walsh, C., Coleman, T. A., Muir, W. J., et al. (1999). Analysis and metaanalysis of two polymorphisms within the tyrosine hydroxylase gene in bipolar and unipolar affective disorders. *Am. J. Med. Genet.* 88, 88–94. doi:10.1002/(sici)1096-8628(19990205)88:1<88::aid-ajmg16>3.0.co;2-j.

Gajecka, M., Mackay, K. L., and Shaffer, L. G. (2007). Monosomy 1p36 deletion syndrome. *Am J Med Genet C Semin Med Genet* 145C, 346–356. doi:10.1002/ajmg.c.30154.

Galfalvy, H., Huang, Y.-Y., Oquendo, M. A., Currier, D., and Mann, J. J. (2009). Increased risk of suicide attempt in mood disorders and TPH1 genotype. *J Affect Disord* 115, 331–338. doi:10.1016/j.jad.2008.09.019.

Gamage, T. H., Misceo, D., Fannemel, M., and Frengen, E. (2013). A balanced de novo inv(7)(p14.3q22.3) disrupting PDE1C and ATXN7L1 in a 14-year old developmentally delayed boy. *European Journal of Medical Genetics* 56, 361–364. doi:10.1016/j.ejmg.2013.04.005.

Gao, X., Zheng, R., Ma, X., Gong, Z., Xia, D., and Zhou, Q. (2019). Elevated Level of PKMζ Underlies the Excessive Anxiety in an Autism Model. *Front Mol Neurosci* 12. doi:10.3389/fnmol.2019.00291.

Garai, C., Furuichi, T., Kawamoto, Y., Ryu, H., and Inoue-Murayama, M. (2014). Androgen receptor and monoamine oxidase polymorphism in wild bonobos. *Meta Gene* 2, 831–843. doi:10.1016/j.mgene.2014.10.005.

Gautier, H. O. B., Evans, K. A., Volbracht, K., James, R., Sitnikov, S., Lundgaard, I., et al. (2015). Neuronal activity regulates remyelination via glutamate signalling to oligodendrocyte progenitors. *Nature Communications* 6, 1–15. doi:10.1038/ncomms9518.

Geisheker, M. R., Heymann, G., Wang, T., Coe, B. P., Turner, T. N., Stessman, H. A. F., et al. (2017). Hotspots of missense mutation identify novel neurodevelopmental disorder genes and functional domains. *Nat Neurosci* 20, 1043–1051. doi:10.1038/nn.4589.

George-Abraham, J. K., Zimmerman, S. L., Hinton, R. B., Marino, B. S., Witte, D. P., and Hopkin, R. J. (2012). Tetrasomy 15q25.2 → qter identified with SNP microarray in a patient with multiple anomalies including complex cardiovascular malformation. *American Journal of Medical Genetics Part A* 158A, 1971–1976. doi:10.1002/ajmg.a.35428.

Germer, E. L., Imhoff, S., Vilariño-Güell, C., Kasten, M., Seibler, P., Brüggemann, N., et al. (2019). The Role of Rare Coding Variants in Parkinson's Disease GWAS Loci. *Front Neurol* 10, 1284. doi:10.3389/fneur.2019.01284.

Gershon, E. S., Grennan, K., Busnello, J., Badner, J. A., Ovsiew, F., Memon, S., et al. (2014). A rare mutation of CACNA1C in a patient with Bipolar disorder, and decreased gene expression associated with a Bipolar-associated common SNP of CACNA1C in brain. *Mol Psychiatry* 19, 890–894. doi:10.1038/mp.2013.107.

Giil, L. M., Vedeler, C. A., Kristoffersen, E. K., Nordrehaug, J. E., Heidecke, H., Dechend, R., et al. (2017). Antibodies to Signaling Molecules and Receptors in Alzheimer's Disease are Associated with Psychomotor Slowing, Depression, and Poor Visuospatial Function. *J. Alzheimers Dis.* 59, 929–939. doi:10.3233/JAD-170245.

Gilabert-Juan, J., Sáez, A. R., Lopez-Campos, G., Sebastiá-Ortega, N., González-Martínez, R., Costa, J., et al. (2015). Semaphorin and plexin gene expression is altered in the prefrontal cortex of schizophrenia patients with and without auditory hallucinations. *Psychiatry Research* 229, 850–857. doi:10.1016/j.psychres.2015.07.074.

Gill, K. M., Lodge, D. J., Cook, J. M., Aras, S., and Grace, A. A. (2011). A Novel α5GABAAR-Positive Allosteric Modulator Reverses Hyperactivation of the Dopamine System in the MAM Model of Schizophrenia. *Neuropsychopharmacology* 36, 1903–1911. doi:10.1038/npp.2011.76.

Gimelli, S., Capra, V., Di Rocco, M., Leoni, M., Mirabelli-Badenier, M., Schiaffino, M. C., et al. (2014). Interstitial 7q31.1 copy number variations disrupting IMMP2L gene are associated with a wide spectrum of neurodevelopmental disorders. *Mol Cytogenet* 7, 54. doi:10.1186/s13039-014-0054-y.

Gjørlund, M. D., Nielsen, J., Pankratova, S., Li, S., Korshunova, I., Bock, E., et al. (2012). Neuroligin-1 induces neurite outgrowth through interaction with neurexin-1β and activation of fibroblast growth factor receptor-1. *FASEB J.* 26, 4174–4186. doi:10.1096/fj.11-202242.

Glaser, B., Kirov, G., Green, E., Craddock, N., and Owen, M. J. (2005). Linkage disequilibrium mapping of bipolar affective disorder at 12q23-q24 provides evidence for association at CUX2 and FLJ32356. *American Journal of Medical Genetics Part B: Neuropsychiatric Genetics* 132B, 38–45. doi:10.1002/ajmg.b.30081.

Glatt, S. J., Chandler, S. D., Bousman, C. A., Chana, G., Lucero, G. R., Tatro, E., et al. (2009). Alternatively Spliced Genes as Biomarkers for Schizophrenia, Bipolar Disorder and Psychosis: A Blood-Based Spliceome-Profiling Exploratory Study. *Curr Pharmacogenomics Person Med* 7, 164–188.

Glatt, S. J., Stone, W. S., Nossova, N., Liew, C.-C., Seidman, L. J., and Tsuang, M. T. (2011). Similarities and differences in peripheral blood gene-expression signatures of individuals with schizophrenia and their first-degree biological relatives. *American Journal of Medical Genetics Part B: Neuropsychiatric Genetics* 156, 869–887. doi:10.1002/ajmg.b.31239.

Glessner, J. T., Wang, K., Cai, G., Korvatska, O., Kim, C. E., Wood, S., et al. (2009). Autism genome-wide copy number variation reveals ubiquitin and neuronal genes. *Nature* 459, 569–573. doi:10.1038/nature07953.

Goes, F. S., Rongione, M., Chen, Y.-C., Karchin, R., Elhaik, E., and Potash, J. B. (2011). Exonic DNA Sequencing of ERBB4 in Bipolar Disorder. *PLoS One* 6. doi:10.1371/journal.pone.0020242.

Golding, J. P., Trainor, P., Krumlauf, R., and Gassmann, M. (2000). Defects in pathfinding by cranial neural crest cells in mice lacking the neuregulin receptor ErbB4. *Nat. Cell Biol.* 2, 103–109. doi:10.1038/35000058.

González-Peñas, J., Arrojo, M., Paz, E., Brenlla, J., Páramo, M., and Costas, J. (2015). Cumulative role of rare and common putative functional genetic variants at NPAS3 in schizophrenia susceptibility. *Am. J. Med. Genet. B Neuropsychiatr. Genet.* 168, 528–535. doi:10.1002/ajmg.b.32324.

Gould, P., and Kamnasaran, D. (2011). Immunohistochemical Analyses of NPAS3 Expression in the Developing Human Fetal Brain. *Anatomia, Histologia, Embryologia* 40, 196–203. doi:10.1111/j.1439-0264.2010.01059.x.

Gozzi, M., Nielson, D. M., Lenroot, R. K., Ostuni, J. L., Luckenbaugh, D. A., Thurm, A. E., et al. (2012). A Magnetization Transfer Imaging Study of Corpus Callosum Myelination in Young Children with Autism. *Biological Psychiatry* 72, 215–220. doi:10.1016/j.biopsych.2012.01.026.

Gragnoli, C. (2014). Hypothesis of the neuroendocrine cortisol pathway gene role in the comorbidity of depression, type 2 diabetes, and metabolic syndrome. *Appl Clin Genet* 7, 43–53. doi:10.2147/TACG.S39993.

Grant, A., Fathalli, F., Rouleau, G., Joober, R., and Flores, C. (2012). Association between schizophrenia and genetic variation in DCC: a case-control study. *Schizophr. Res.* 137, 26–31. doi:10.1016/j.schres.2012.02.023.

Grant, A., Hoops, D., Labelle-Dumais, C., Prévost, M., Rajabi, H., Kolb, B., et al. (2007). Netrin-1 receptor-deficient mice show enhanced mesocortical dopamine transmission and blunted behavioural responses to amphetamine. *Eur. J. Neurosci.* 26, 3215–3228. doi:10.1111/j.1460-9568.2007.05888.x.

Gratacòs, M., Costas, J., Cid, R. de, Bayés, M., González, J. R., Baca-García, E., et al. (2009). Identification of new putative susceptibility genes for several psychiatric disorders by association analysis of regulatory and non-synonymous SNPs of 306 genes involved in neurotransmission and neurodevelopment. *American Journal of Medical Genetics Part B: Neuropsychiatric Genetics* 150B, 808–816. doi:10.1002/ajmg.b.30902.

Gray, N. W. (2005). A dynamin-3 spliced variant modulates the actin/cortactin-dependent morphogenesis of dendritic spines. Journal of Cell Science 118, 1279–1290. doi:10.1242/jcs.01711.

Gray, M. M., Sutter, N. B., Ostrander, E. A., and Wayne, R. K. (2010). The IGF1 small dog haplotype is derived from Middle Eastern grey wolves. *BMC Biol.* 8, 16. doi:10.1186/1741-7007-8-16.

Grayton, H. M., Missler, M., Collier, D. A., and Fernandes, C. (2013). Altered Social Behaviours in Neurexin 1α Knockout Mice Resemble Core Symptoms in Neurodevelopmental Disorders. *PLoS One* 8. doi:10.1371/journal.pone.0067114.

Green, E. K., Grozeva, D., Jones, I., Jones, L., Kirov, G., Caesar, S., et al. (2010). The bipolar disorder risk allele at CACNA1C also confers risk of recurrent major depression and of schizophrenia. *Mol Psychiatry* 15, 1016–1022. doi:10.1038/mp.2009.49.

Griswold, A. J., Ma, D., Cukier, H. N., Nations, L. D., Schmidt, M. A., Chung, R.-H., et al. (2012). Evaluation of copy number variations reveals novel candidate genes in autism spectrum disorder-associated pathways. *Hum Mol Genet* 21, 3513–3523. doi:10.1093/hmg/dds164.

Grover, D., Verma, R., Goes, F. S., Mahon, P. L. B., Gershon, E. S., McMahon, F. J., et al. (2009). Family-based association of YWHAH in psychotic bipolar disorder. *Am J Med Genet B Neuropsychiatr Genet* 0, 977–983. doi:10.1002/ajmg.b.30927.

Grozeva, D., Kirov, G., Ivanov, D., Jones, I. R., Jones, L., Green, E. K., et al. (2010). Rare Copy Number Variants. *Arch Gen Psychiatry* 67, 318–327. doi:10.1001/archgenpsychiatry.2010.25.

Gulevich, R. G., Oskina, I. N., Shikhevich, S. G., Fedorova, E. V., and Trut, L. N. (2004). Effect of selection for behavior on pituitary-adrenal axis and proopiomelanocortin gene expression in silver foxes (*Vulpes vulpes*). *Physiol. Behav.* 82, 513–518. doi:10.1016/j.physbeh.2004.04.062.

Guo, H., Wang, T., Wu, H., Long, M., Coe, B. P., Li, H., et al. (2018). Inherited and multiple de novo mutations in autism/developmental delay risk genes suggest a multifactorial model. *Mol Autism* 9. doi:10.1186/s13229-018-0247-z.

Guo, J., Bian, Y., Wang, Y., Chen, L., Yu, A., and Sun, X. (2017). FAM107B is regulated by S100A4 and mediates the effect of S100A4 on the proliferation and migration of MGC803 gastric cancer cells. *Cell Biol. Int.* 41, 1103–1109. doi:10.1002/cbin.10816.

Guo, L., Stormmesand, J., Fang, Z., Zhu, Q., Balesar, R., van Heerikhuize, J., et al. (2019). Quantification of Tyrosine Hydroxylase and ErbB4 in the Locus Coeruleus of Mood Disorder Patients Using a Multispectral Method to Prevent Interference with Immunocytochemical Signals by Neuromelanin. *Neurosci Bull* 35, 205–215. doi:10.1007/s12264-019-00339-y.

Guo, S.-Z., Huang, K., Shi, Y.-Y., Tang, W., Zhou, J., Feng, G.-Y., et al. (2007). A Case-control association study between the GRID1 gene and schizophrenia in the Chinese Northern Han population. *Schizophrenia Research* 93, 385–390. doi:10.1016/j.schres.2007.03.007.

Gupta, A. R., Westphal, A., Yang, D. Y. J., Sullivan, C. A. W., Eilbott, J., Zaidi, S., et al. (2017). Neurogenetic analysis of childhood disintegrative disorder. *Mol Autism* 8, 19. doi:10.1186/s13229-017-0133-0.

Gupta, V., Khan, A. A., Sasi, B. K., and Mahapatra, N. R. (2015). Molecular mechanism of monoamine oxidase A gene regulation under inflammation and ischemia-like conditions: key roles of the transcription factors GATA2, Sp1 and TBP. *Journal of Neurochemistry* 134, 21–38. doi:10.1111/jnc.13099.

Guzmán, Y. F., Ramsey, K., Stolz, J. R., Craig, D. W., Huentelman, M. J., Narayanan, V., et al. (2017). A gain-of-function mutation in the GRIK2 gene causes neurodevelopmental deficits. *Neurol Genet* 3. doi:10.1212/NXG.0000000000000129.

Hall, N. J., and Wynne, C. D. L. (2012). The canid genome: behavioral geneticists' best friend? *Genes, Brain and Behavior* 11, 889–902. doi:10.1111/j.1601-183X.2012.00851.x.

Hare, B., Wobber, V., and Wrangham, R. (2012). The self-domestication hypothesis: evolution of bonobo psychology is due to selection against aggression. *Animal Behaviour* 83, 573–585. doi:10.1016/j.anbehav.2011.12.007.

Haroutunian, V., and Davis, K. L. (2007). Introduction to the Special Section: Myelin and oligodendrocyte abnormalities in schizophrenia. *Int J Neuropsychopharmacol* 10, 499–502. doi:10.1017/S1461145706007449.

Hashizume, C., Masuda, K., Momozawa, Y., Kikusui, T., Takeuchi, Y., and Mori, Y. (2005). Identification of an cysteine-to-arginine substitution caused by a single nucleotide polymorphism in the canine monoamine oxidase B gene. *J. Vet. Med. Sci.* 67, 199–201. doi:10.1292/jvms.67.199.

Hegde, S. S., and Eglen, R. M. (1996). Peripheral 5-HT4 receptors. *FASEB J.* 10, 1398–1407. doi:10.1096/fasebj.10.12.8903510.

Hekman, J. P., Johnson, J. L., Edwards, W., Vladimirova, A. V., Gulevich, R. G., Ford, A. L., et al. (2018). Anterior pituitary transcriptome suggests differences in ACTH release in tame and aggressive foxes. *G3* (*Bethesda*) 8, 859–873. doi:10.1534/g3.117.300508.

Heid, I. M., Jackson, A. U., Randall, J. C., Winkler, T. W., Qi, L., Steinthorsdottir, V., et al. (2010). Meta-analysis identifies 13 new loci associated with waist-hip ratio and reveals sexual dimorphism in the genetic basis of fat distribution. *Nat. Genet.* 42, 949–960. doi:10.1038/ng.685.

Heim, C., Owens, M. J., Plotsky, P. M., and Nemeroff, C. B. (1997). Persistent changes in corticotropin-releasing factor systems due to early life stress: relationship to the pathophysiology of major depression and post-traumatic stress disorder. *Psychopharmacol Bull* 33, 185–192.

Hercig, D., Bau, A., and Reznek, L. (1994). Psychosis associated with lobar holoprosencephaly. *Can J Psychiatry* 39, 449. doi:10.1177/070674379403900716.

Hering, H., and Sheng, M. (2003). Activity-Dependent Redistribution and Essential Role of Cortactin in Dendritic Spine Morphogenesis. *J Neurosci* 23, 11759–11769. doi:10.1523/JNEUROSCI.23-37-11759.2003.

Hernandez, C. C., XiangWei, W., Hu, N., Shen, D., Shen, W., Lagrange, A. H., et al. (2019). Altered inhibitory synapses in de novo GABRA5 and GABRA1 mutations associated with early onset epileptic encephalopathies. *Brain* 142, 1938–1954. doi:10.1093/brain/awz123.

Hibar, D. P., Adams, H. H. H., Jahanshad, N., Chauhan, G., Stein, J. L., Hofer, E., et al. (2017). Novel genetic loci associated with hippocampal volume. *Nat Commun* 8. doi:10.1038/ncomms13624.

Hill, S. Y., Weeks, D. E., Jones, B. L., Zezza, N., and Stiffler, S. (2012). ASTN1 and alcohol dependence: family-based association analysis in multiplex alcohol dependence families. *Am. J. Med. Genet. B Neuropsychiatr. Genet.* 159B, 445–455. doi:10.1002/ajmg.b.32048.

Hitchins, M. P., Bentley, L., Monk, D., Beechey, C., Peters, J., Kelsey, G., et al. (2002). DDC and COBL, flanking the imprinted GRB10 gene on 7p12, are biallelically expressed. *Mamm Genome* 13, 686–691. doi:10.1007/s00335-002-3028-z.

Hofer, N. T., Tuluc, P., Ortner, N. J., Nikonishyna, Y. V., Fernándes-Quintero, M. L., Liedl, K. R., et al. (2020). Biophysical classification of a CACNA1D de novo mutation as a high-risk mutation for a severe neurodevelopmental disorder. *Mol Autism* 11. doi:10.1186/s13229-019-0310-4.

Holmes, D. I. R., Wahab, N. A., and Mason, R. M. (1999). Cloning and Characterization of ZNF236, a Glucose-Regulated Kruppel-like Zinc-Finger Gene Mapping to Human Chromosome 18q22–q23. *Genomics* 60, 105–109. doi:10.1006/geno.1999.5897.

Holt, R., Barnby, G., Maestrini, E., Bacchelli, E., Brocklebank, D., Sousa, I., et al. (2010). Linkage and candidate gene studies of autism spectrum disorders in European populations. *Eur. J. Hum. Genet.* 18, 1013–1019. doi:10.1038/ejhg.2010.69.

Hoogduijn, M. J., Hitchcock, I. S., Smit, N. P. M., Gillbro, J. M., Schallreuter, K. U., and Genever, P. G. (2006). Glutamate receptors on human melanocytes regulate the expression of MiTF. *Pigment Cell Res.* 19, 58–67. doi:10.1111/j.1600-0749.2005.00284.x.

Hooper, A. W. M., Alamilla, J. F., Venier, R. E., Gillespie, D. C., and Igdoura, S. A. (2017). Neuronal pentraxin 1 depletion delays neurodegeneration and extends life in Sandhoff disease mice. *Hum Mol Genet* 26, 661–673. doi:10.1093/hmg/ddw422.

Horiguchi, M., Ohi, K., Hashimoto, R., Hao, Q., Yasuda, Y., Yamamori, H., et al. (2014). Functional polymorphism (C-824T) of the tyrosine hydroxylase gene affects IQ in schizophrenia. *Psychiatry Clin. Neurosci.* 68, 456–462. doi:10.1111/pcn.12157.

Hotamisligil, G. S., Girmen, A. S., Fink, J. S., Tivol, E., Shalish, C., Trofatter, J., et al. (1994). Hereditary variations in monoamine oxidase as a risk factor for Parkinson's disease. *Mov. Disord.* 9, 305–310. doi:10.1002/mds.870090304.

Hou, X.-J., Ni, K.-M., Yang, J.-M., and Li, X.-M. (2014). Neuregulin 1/ErbB4 enhances synchronized oscillations of prefrontal cortex neurons via inhibitory synapses. *Neuroscience* 261, 107–117. doi:10.1016/j.neuroscience.2013.12.040.

Howland, R. H. (1996). Induction of mania with serotonin reuptake inhibitors. *J Clin Psychopharmacol* 16, 425–427. doi:10.1097/00004714-199612000-00003.

Hu, J., Chan, L. F., Souza, R. P., Tampakeras, M., Kennedy, J. L., Zai, C., et al. (2014). The role of tyrosine hydroxylase gene variants in suicide attempt in schizophrenia. *Neurosci. Lett.* 559, 39–43. doi:10.1016/j.neulet.2013.11.025.

Hu, J., Liao, J., Sathanoori, M., Kochmar, S., Sebastian, J., Yatsenko, S. A., et al. (2015). CNTN6 copy number variations in 14 patients: a possible candidate gene for neurodevelopmental and neuropsychiatric disorders. *J Neurodev Disord* 7. doi:10.1186/s11689-015-9122-9.

Hu, V. W., Devlin, C. A., and Debski, J. J. (2019). ASD Phenotype—Genotype Associations in Concordant and Discordant Monozygotic and Dizygotic Twins Stratified by Severity of Autistic Traits. *Int J Mol Sci* 20. doi:10.3390/ijms20153804.

Hu, Z., Ying, X., Huang, L., Zhao, Y., Zhou, D., Liu, J., et al. (2020). Association of human serotonin receptor 4 promoter methylation with autism spectrum disorder. *Medicine (Baltimore)* 99, e18838. doi:10.1097/MD.0000000000018838.

Huang, A. Y., Yu, D., Davis, L. K., Sul, J. H., Tsetsos, F., Ramensky, V., et al. (2017). Rare copy number variants in NRXN1 and CNTN6 increase risk for Tourette syndrome. *Neuron* 94, 1101-1111.e7. doi:10.1016/j.neuron.2017.06.010.

Huang, J., Perlis, R. H., Lee, P. H., Rush, A. J., Fava, M., Sachs, G. S., et al. (2010). Cross-Disorder Genomewide Analysis of Schizophrenia, Bipolar Disorder, and Depression. *Am J Psychiatry* 167. doi:10.1176/appi.ajp.2010.09091335.

Huang, S., Slomianka, L., Farmer, A. J., Kharlamova, A. V., Gulevich, R. G., Herbeck, Y. E., et al. (2015). Selection for tameness, a key behavioral trait of domestication, increases adult hippocampal neurogenesis in foxes. *Hippocampus* 25, 963–975. doi:10.1002/hipo.22420.

Hur, C.-G., Kim, E.-J., Cho, S.-K., Cho, Y.-W., Yoon, S.-Y., Tak, H.-M., et al. (2012). K(+) efflux through two-pore domain K(+) channels is required for mouse embryonic development. *Reproduction* 143, 625–636. doi:10.1530/REP-11-0225.

Hussman, J. P. (2001). Suppressed GABAergic Inhibition as a Common Factor in Suspected Etiologies of Autism. *J Autism Dev Disord* 31, 247–248. doi:10.1023/A:1010715619091.

Imagawa, E., Higashimoto, K., Sakai, Y., Numakura, C., Okamoto, N., Matsunaga, S., et al. (2017). Mutations in genes encoding polycomb repressive complex 2 subunits cause Weaver syndrome. *Human Mutation* 38, 637–648. doi:10.1002/humu.23200.

Inaguma, Y., Matsumoto, A., Noda, M., Tabata, H., Maeda, A., Goto, M., et al. (2016). Role of Class III phosphoinositide 3-kinase in the brain development: possible involvement in specific learning disorders. *Journal of Neurochemistry* 139, 245–255. doi:10.1111/jnc.13832.

Insel, P. A. (2011). β(2)-Adrenergic receptor polymorphisms and signaling: Do variants influence the "memory" of receptor activation? *Sci Signal* 4(185), pe37.

Iossifov, I., O'Roak, B. J., Sanders, S. J., Ronemus, M., Krumm, N., Levy, D., et al. (2014). The contribution of de novo coding mutations to autism spectrum disorder. *Nature* 515, 216–221. doi:10.1038/nature13908.

Iossifov, I., Ronemus, M., Levy, D., Wang, Z., Hakker, I., Rosenbaum, J., et al. (2012). De Novo Gene Disruptions in Children on the Autistic Spectrum. *Neuron* 74, 285–299. doi:10.1016/j.neuron.2012.04.009.

Ishida, M., Cullup, T., Boustred, C., James, C., Docker, J., English, C., et al. (2018). A targeted sequencing panel identifies rare damaging variants in multiple genes in the cranial neural tube defect, anencephaly. *Clin Genet* 93, 870–879. doi:10.1111/cge.13189.

Iurov, I. I., Vorsanova, S. G., Saprina, E. A., and Iurov, I. B. (2010). [Identification of candidate genes of autism on the basis of molecular cytogenetic and in silico studies of the genome organization of chromosomal regions involved in unbalanced rearrangements]. *Genetika* 46, 1348–1351.

Ivanov, H., Stoyanova, V., Ivanov, I., Linev, A., Vazharova, R., Ivanov, S., et al. (2018). Rare Case of a Heterozygous Microdeletion 9q21.11-q21.2: Clinical and Genetic Characteristics. *Balkan J Med Genet* 21, 59–62. doi:10.2478/bjmg-2018-0021.

Iwayama, Y., Hattori, E., Maekawa, M., Yamada, K., Toyota, T., Ohnishi, T., et al. (2010). Association analyses between brain-expressed fatty-acid binding protein (FABP) genes and schizophrenia and bipolar disorder. *Am. J. Med. Genet. B Neuropsychiatr. Genet.* 153B, 484–493. doi:10.1002/ajmg.b.31004.

Jacobsen, N. J., Elvidge, G., Franks, E. K., O'Donovan, M. C., Craddock, N., and Owen, M. J. (2001). CUX2, a potential regulator of NCAM expression: genomic characterization and analysis as a positional candidate susceptibility gene for bipolar disorder. *Am. J. Med. Genet.* 105, 295–300. doi:10.1002/ajmg.1325.

Jahnes, E., Müller, D. J., Schulze, T. G., Windemuth, C., Cichon, S., Ohlraun, S., et al. (2002). Association study between two variants in the DOPA decarboxylase gene in bipolar and unipolar affective disorder. *Am. J. Med. Genet.* 114, 519–522. doi:10.1002/ajmg.10308.

Jamain, S., Betancur, C., Quach, H., Philippe, A., Fellous, M., Giros, B., et al. (2002). Linkage and association of the glutamate receptor 6 gene with autism. *Mol. Psychiatry* 7, 302–310. doi:10.1038/sj.mp.4000979.

Jamra, R. A., Schulze, T. G., Becker, T., Brockschmidt, F. F., Green, E., Alblas, M. A., et al. (2010). A systematic association mapping on chromosome 6q in bipolar affective disorder—evidence for the melanin-concentrating-hormone-receptor-2 gene as a risk factor for bipolar affective disorder. *American Journal of Medical Genetics Part B: Neuropsychiatric Genetics* 153B, 878–884. doi:10.1002/ajmg.b.31051.

Jenkins, A., Apud, J. A., Zhang, F., Decot, H., Weinberger, D. R., and Law, A. J. (2014). Identification of Candidate Single-Nucleotide Polymorphisms in NRXN1 Related to Antipsychotic Treatment Response in Patients with Schizophrenia. *Neuropsychopharmacology* 39, 2170–2178. doi:10.1038/npp.2014.65.

Jenkins, A. K., Paterson, C., Wang, Y., Hyde, T. M., Kleinman, J. E., and Law, A. J. (2016). Neurexin 1 (NRXN1) Splice Isoform Expression During Human Neocortical Development and Aging. *Mol Psychiatry* 21, 701–706. doi:10.1038/mp.2015.107.

Jensen, P. (2015). Adding 'epi-' to behaviour genetics: implications for animal domestication. *Journal of Experimental Biology* 218, 32–40. doi:10.1242/jeb.106799.

Ji, T., Wu, Y., Wang, H., Wang, J., and Jiang, Y. (2010). Diagnosis and fine mapping of a deletion in distal 11q in two Chinese patients with developmental delay. *Journal of Human Genetics* 55, 486–489. doi:10.1038/jhg.2010.51.

Ji, X., Kember, R. L., Brown, C. D., and Bućan, M. (2016). Increased burden of deleterious variants in essential genes in autism spectrum disorder. *Proc Natl Acad Sci U S A* 113, 15054–15059. doi:10.1073/pnas.1613195113.

Jia, B., Huang, L., Chen, Y., Liu, S., Chen, C., Xiong, K., et al. (2017). A novel contiguous deletion involving NDP, MAOB and EFHC2 gene in a patient with familial Norrie disease: bilateral blindness and leucocoria without other deficits. *J. Genet.* 96, 1015–1020. doi:10.1007/s12041-017-0869-5.

Jiang, Q., Arnold, S., Heanue, T., Kilambi, K. P., Doan, B., Kapoor, A., et al. (2015). Functional Loss of Semaphorin 3C and/or Semaphorin 3D and Their Epistatic Interaction with Ret Are Critical to Hirschsprung Disease Liability. *Am J Hum Genet* 96, 581–596. doi:10.1016/j.ajhg.2015.02.014.

Jiang, Y., Yuen, R. K. C., Jin, X., Wang, M., Chen, N., Wu, X., et al. (2013). Detection of clinically relevant genetic variants in autism spectrum disorder by whole-genome sequencing. *Am. J. Hum. Genet.* 93, 249–263. doi:10.1016/j.ajhg.2013.06.012.

Jinnah, H. A., Yitta, S., Drew, T., Kim, B. S., Visser, J. E., and Rothstein, J. D. (1999). Calcium channel activation and self-biting in mice. *Proc Natl Acad Sci U S A* 96, 15228–15232.

Johansen, J. P., Cain, C. K., Ostroff, L. E., and LeDoux, J. E. (2011). Molecular Mechanisms of Fear Learning and Memory. *Cell* 147, 509–524. doi:10.1016/j.cell.2011.10.009.

Johnson, C., Drgon, T., McMahon, F. J., and Uhl, G. R. (2009). Convergent genome wide association results for bipolar disorder and substance dependence. *American Journal of Medical Genetics Part B: Neuropsychiatric Genetics* 150B, 182–190. doi:10.1002/ajmg.b.30900.

Joiner, M. L., Lisé, M. F., Yuen, E. Y., Kam, A. Y., Zhang, M., et al. (2010). Assembly of a beta2-adrenergic receptor--GluR1 signalling complex for localized cAMP signalling. EMBO J. 29(2), 482-95.

Jokinen, J., Boström, A. E., Dadfar, A., Ciuculete, D. M., Chatzittofis, A., Åsberg, M., et al. (2017). Epigenetic Changes in the CRH Gene are Related to Severity of Suicide Attempt and a General Psychiatric Risk Score in Adolescents. *EBioMedicine* 27, 123–133. doi:10.1016/j.ebiom.2017.12.018.

Juan-Perez, C., Farrand, S., and Velakoulis, D. (2018). Schizophrenia and epilepsy as a result of maternally inherited CNTN6 copy number variant. *Schizophr. Res.* 202, 111–112. doi:10.1016/j.schres.2018.06.062.

Kabir, Z., Che, A., Fischer, D., Rice, R., Rizzo, B., Byrne, M., et al. (2017). Rescue of impaired sociability and anxiety-like behavior in adult cacna1c-deficient mice by pharmacologically targeting eIF2α. *Mol Psychiatry* 22, 1096–1109. doi:10.1038/mp.2017.124.

Kabir, Z. D., Lee, A. S., and Rajadhyaksha, A. M. (2016). L-type Ca2+ channels in mood, cognition and addiction: integrating human and rodent studies with a focus on behavioural endophenotypes. *J Physiol* 594, 5823–5837. doi:10.1113/JP270673.

Kähler, A. K., Djurovic, S., Kulle, B., Jönsson, E. G., Agartz, I., Hall, H., et al. (2008). Association analysis of schizophrenia on 18 genes involved in neuronal migration: MDGA1 as a new susceptibility gene. *Am. J. Med. Genet. B Neuropsychiatr. Genet.* 147B, 1089–1100. doi:10.1002/ajmg.b.30726.

Kamm, G. B., Pisciottano, F., Kliger, R., and Franchini, L. F. (2013). The Developmental Brain Gene NPAS3 Contains the Largest Number of Accelerated Regulatory Sequences in the Human Genome. *Mol Biol Evol* 30, 1088–1102. doi:10.1093/molbev/mst023.

Kamnasaran, D., Chen, C.-P., Devriendt, K., Mehta, L., and Cox, D. W. (2005). Defining a holoprosencephaly locus on human chromosome 14q13 and characterization of potential candidate genes. *Genomics* 85, 608–621. doi:10.1016/j.ygeno.2005.01.010.

Kamnasaran, D., Muir, W., Ferguson-Smith, M., and Cox, D. (2003). Disruption of the neuronal PAS3 gene in a family affected with schizophrenia. *J Med Genet* 40, 325–332. doi:10.1136/jmg.40.5.325.

Kandaswamy, R., McQuillin, A., Curtis, D., and Gurling, H. (2012). Tests of linkage and allelic association between markers in the 1p36 PRKCZ (protein kinase C zeta) gene region and bipolar affective disorder. *Am. J. Med. Genet. B Neuropsychiatr. Genet.* 159B, 201–209. doi:10.1002/ajmg.b.32014.

Kang, D., and Kim, D. (2006). TREK-2 (K2P10.1) and TRESK (K2P18.1) are major background K+ channels in dorsal root ganglion neurons. *American Journal of Physiology-Cell Physiology* 291, C138–C146. doi:10.1152/ajpcell.00629.2005.

Kang, D.-S., Yang, Y. R., Lee, C., Park, B., Park, K. I., Seo, J. K., et al. (2018). Netrin-1/DCC-mediated PLCγ1 activation is required for axon guidance and brain structure development. *EMBO Rep.* 19. doi:10.15252/embr.201846250.

Kaphzan, H., Hernandez, P., Jung, J. I., Cowansage, K. K., Deinhardt, K., Chao, M. V., et al. (2012). Reversal of Impaired Hippocampal Long-term Potentiation and Contextual Fear Memory Deficits in Angelman Syndrome Model Mice by ErbB Inhibitors. *Biol Psychiatry* 72, 182–190. doi:10.1016/j.biopsych.2012.01.021.

Kar, S., Chabot, J. G., and Quirion, R. (1993). Quantitative autoradiographic localization of [125I]insulin-like growth factor I, [125I]insulin-like growth factor II, and [125I]insulin receptor binding sites in developing and adult rat brain. *J. Comp. Neurol.* 333, 375–397. doi:10.1002/cne.903330306.

Kashevarova, A. A., Nazarenko, L. P., Schultz-Pedersen, S., Skryabin, N. A., Salyukova, O. A., Chechetkina, N. N., et al. (2014). Single gene microdeletions and microduplication of 3p26.3 in three unrelated families: CNTN6 as a new candidate gene for intellectual disability. *Mol Cytogenet* 7. doi:10.1186/s13039-014-0097-0.

Kasim, S., and Jinnah, H. A. (2003). Self-Biting Induced by Activation of L-Type Calcium Channels in Mice: Dopaminergic Influences. *DNE* 25, 20–25. doi:10.1159/000071464.

Kasnauskiene, J., Ciuladaite, Z., Preiksaitiene, E., Utkus, A., Peciulyte, A., and Kučinskas, V. (2013). A new single gene deletion on 2q34: ERBB4 is associated with intellectual disability. *Am. J. Med. Genet. A* 161A, 1487–1490. doi:10.1002/ajmg.a.35911.

Kato, T. (2007). Molecular genetics of bipolar disorder and depression. *Psychiatry Clin. Neurosci.* 61, 3–19. doi:10.1111/j.1440-1819.2007.01604.x.

Kato, T. M., Kubota-Sakashita, M., Fujimori-Tonou, N., Saitow, F., Fuke, S., Masuda, A., et al. (2018). Ant1 mutant mice bridge the mitochondrial and serotonergic dysfunctions in bipolar disorder. *Mol Psychiatry* 23, 2039–2049. doi:10.1038/s41380-018-0074-9.

Keller, J., Gomez, R., Williams, G., Lembke, A., Lazzeroni, L., Murphy, G. M., et al. (2017). HPA Axis in Major Depression: Cortisol, Clinical Symptomatology, and Genetic Variation Predict Cognition. *Mol Psychiatry* 22, 527–536. doi:10.1038/mp.2016.120.

Kelsoe, J. R., Spence, M. A., Loetscher, E., Foguet, M., Sadovnick, A. D., Remick, R. A., et al. (2001). A genome survey indicates a possible susceptibility locus for bipolar disorder on chromosome 22. *Proc. Natl. Acad. Sci. U.S.A.* 98, 585–590. doi:10.1073/pnas.011358498.

Kempermann, G., Krebs, J., and Fabel, K. (2008). The contribution of failing adult hippocampal neurogenesis to psychiatric disorders. *Curr Opin Psychiatry* 21, 290–295. doi:10.1097/YCO.0b013e3282fad375.

Kennard, L. E., Chumbley, J. R., Ranatunga, K. M., Armstrong, S. J., Veale, E. L., and Mathie, A. (2005). Inhibition of the human two-pore domain potassium channel, TREK-1, by fluoxetine and its metabolite norfluoxetine. *Br. J. Pharmacol.* 144, 821–829. doi:10.1038/sj.bjp.0706068.

Kerner, B., Jasinska, A. J., DeYoung, J., Almonte, M., Choi, O.-W., and Freimer, N. B. (2009). Polymorphisms in the GRIA1 Gene Region in Psychotic Bipolar Disorder. *Am J Med Genet B Neuropsychiatr Genet* 0, 24–32. doi:10.1002/ajmg.b.30780.

Kerner, B., Lambert, C. G., and Muthén, B. O. (2011). Genome-Wide Association Study in Bipolar Patients Stratified by Co-Morbidity. *PLoS One* 6. doi:10.1371/journal.pone.0028477.

Kim, E.-J., Lee, D. K., Hong, S.-G., Han, J., and Kang, D. (2017). Activation of TREK-1, but Not TREK-2, Channel by Mood Stabilizers. *Int J Mol Sci* 18. doi:10.3390/ijms18112460.

Kim, H.-G., Kishikawa, S., Higgins, A. W., Seong, I.-S., Donovan, D. J., Shen, Y., et al. (2008). Disruption of neurexin 1 associated with autism spectrum disorder. *Am. J. Hum. Genet.* 82, 199–207. doi:10.1016/j.ajhg.2007.09.011.

Kim, Y. S., Jang, S.-W., Sung, H. J., Lee, H. J., Kim, I. S., Na, D. S., et al. (2005). Role of 14-3-3 eta as a positive regulator of the glucocorticoid receptor transcriptional activation. *Endocrinology* 146, 3133–3140. doi:10.1210/en.2004-1455.

Kim, Y.-K., Na, K.-S., Hwang, J.-A., Yoon, H.-K., Lee, H.-J., Hahn, S.-W., et al. (2013). High insulin-like growth factor-1 in patients with bipolar I disorder: a trait marker? *J Affect Disord* 151, 738–743. doi:10.1016/j.jad.2013.07.041.

Kircher, T., Krug, A., Markov, V., Whitney, C., Krach, S., Zerres, K., et al. (2009). Genetic variation in the schizophrenia-risk gene neuregulin 1 correlates with brain activation and impaired speech production in a verbal fluency task in healthy individuals. *Human Brain Mapping* 30, 3406–3416. doi:10.1002/hbm.20761.

Kisko, T. M., Braun, M. D., Michels, S., Witt, S. H., Rietschel, M., Culmsee, C., et al. (2018). Cacna1c haploinsufficiency leads to pro-social 50-kHz ultrasonic communication deficits in rats. *Dis Model Mech* 11. doi:10.1242/dmm.034116.

Klitten, L. L., Møller, R. S., Ravn, K., Hjalgrim, H., and Tommerup, N. (2011). Duplication of MAOA, MAOB, and NDP in a patient with mental retardation and epilepsy. *Eur J Hum Genet* 19, 1–2. doi:10.1038/ejhg.2010.149.

Konopaske, G. T., Lange, N., Coyle, J. T., and Benes, F. M. (2014). Prefrontal cortical dendritic spine pathology in schizophrenia and bipolar disorder. *JAMA Psychiatry* 71, 1323–1331. doi:10.1001/jamapsychiatry.2014.1582.

Korenbaum, E., Olski, T. M., and Noegel, A. A. (2001). Genomic organization and expression profile of the parvin family of focal adhesion proteins in mice and humans. *Gene* 279, 69–79. doi:10.1016/s0378-1119(01)00743-0.

Krude, H., Biebermann, H., Luck, W., Horn, R., Brabant, G., and Grüters, A. (1998). Severe early-onset obesity, adrenal insufficiency and red hair pigmentation caused by POMC mutations in humans. *Nat. Genet.* 19, 155–157. doi:10.1038/509.

Krude, H., Biebermann, H., Schnabel, D., Tansek, M. Z., Theunissen, P., Mullis, P. E., et al. (2003). Obesity due to proopiomelanocortin deficiency: three new cases and treatment trials with thyroid hormone and ACTH4-10. *J. Clin. Endocrinol. Metab.* 88, 4633–4640. doi:10.1210/jc.2003-030502.

Krug, A., Nieratschker, V., Markov, V., Krach, S., Jansen, A., Zerres, K., et al. (2010). Effect of CACNA1C rs1006737 on neural correlates of verbal fluency in healthy individuals. *Neuroimage* 49, 1831–1836. doi:10.1016/j.neuroimage.2009.09.028.

Kruger, R. P., Aurandt, J., and Guan, K.-L. (2005). Semaphorins command cells to move. *Nature Reviews Molecular Cell Biology* 6, 789–800. doi:10.1038/nrm1740.

Krumm, N., O'Roak, B. J., Shendure, J., and Eichler, E. E. (2014). A de novo convergence of autism genetics and molecular neuroscience. *Trends Neurosci* 37, 95–105. doi:10.1016/j.tins.2013.11.005.

Kubinyi, E., Vas, J., Hejjas, K., Ronai, Z., Brúder, I., Turcsán, B., et al. (2012). Polymorphism in the tyrosine hydroxylase (TH) gene is associated with activity-impulsivity in German Shepherd Dogs. *PLoS ONE* 7, e30271. doi:10.1371/journal.pone.0030271.

Kubo, Y., Baba, K., Toriyama, M., Minegishi, T., Sugiura, T., Kozawa, S., et al. (2015). Shootin1-cortactin interaction mediates signal-force transduction for axon outgrowth. *J. Cell Biol.* 210, 663–676. doi:10.1083/jcb.201505011.

Kuipers, B., Silfhout, A. V., Marcelis, C., Pfundt, R., Leeuw, N. de, and Vries, B. de (2013). Two patients with intellectual disability, overlapping facial features, and overlapping deletions in 6p25.1p24.3. *Clinical Dysmorphology* 22, 18–21. doi:10.1097/MCD.0b013e32835b6e39.

Kumarasinghe, N., Beveridge, N. J., Gardiner, E., Scott, R. J., Yasawardene, S., Perera, A., et al. (2013). Gene expression profiling in treatment-naive schizophrenia patients identifies abnormalities in biological pathways involving AKT1 that are corrected by antipsychotic medication. *Int J Neuropsychopharmacol* 16, 1483–1503. doi:10.1017/S1461145713000035.

Kundakovic, M., and Jaric, I. (2017). The Epigenetic Link between Prenatal Adverse Environments and Neurodevelopmental Disorders. *Genes (Basel)* 8. doi:10.3390/genes8030104.

Kuo, P. H., Chuang, L. C., Liu, J. R., Liu, C. M., Huang, M. C., Lin, S. K., et al. (2014). Identification of novel loci for bipolar I disorder in a multi-stage genome-wide association study. *Progress in Neuro-Psychopharmacology and Biological Psychiatry* 51, 58–64. doi:10.1016/j.pnpbp.2014.01.003.

Kuroyanagi, T., Yokoyama, M., and Hirano, T. (2009). Postsynaptic glutamate receptor δ family contributes to presynaptic terminal differentiation and establishment of synaptic transmission. *Proc Natl Acad Sci U S A* 106, 4912–4916. doi:10.1073/pnas.0900892106.

Kzhyshkowska, J. (2010). Multifunctional Receptor Stabilin-1 in Homeostasis and Disease. *ScientificWorldJournal* 10, 2039–2053. doi:10.1100/tsw.2010.189.

Lai, T.-J., Wu, C.-Y., Tsai, H.-W., Lin, Y.-M. J., and Sun, H. S. (2005). Polymorphism screening and haplotype analysis of the tryptophan hydroxylase gene (TPH1) and association with bipolar affective disorder in Taiwan. *BMC Med. Genet.* 6, 14. doi:10.1186/1471-2350-6-14.

Lambert, D., Middle, F., Hamshere, M. L., Segurado, R., Raybould, R., Corvin, A., et al. (2005). Stage 2 of the Wellcome Trust UK-Irish bipolar affective disorder sibling-pair genome screen: evidence for linkage on chromosomes 6q16-q21, 4q12-q21, 9p21, 10p14-p12 and 18q22. *Mol. Psychiatry* 10, 831–841. doi:10.1038/sj.mp.4001684.

Lauritsen, M., Børglum, A., Betancur, C., Philippe, A., Kruse, T., Leboyer, M., et al. (2002). Investigation of two variants in the DOPA decarboxylase gene in patients with autism. *Am J Med Genet* 114, 466–470. doi:10.1002/ajmg.10379.

Lavado, A., Jeffery, G., Tovar, V., de la Villa, P., and Montoliu, L. (2006). Ectopic expression of tyrosine hydroxylase in the pigmented epithelium rescues the retinal abnormalities and visual function common in albinos in the absence of melanin. *Journal of Neurochemistry* 96, 1201–1211. doi:10.1111/j.1471-4159.2006.03657.x.

Ledda, F., Paratcha, G., Sandoval-Guzmán, T., and Ibáñez, C. F. (2007). GDNF and GFRalpha1 promote formation of neuronal synapses by ligand-induced cell adhesion. *Nat. Neurosci.* 10, 293–300. doi:10.1038/nn1855.

Lee, A. M., Kanter, B. R., Wang, D., Lim, J. P., Zou, M. E., Qiu, C., et al. (2013a). Prkcz null mice show normal learning and memory. *Nature* 493, 416–419. doi:10.1038/nature11803.

Lee, A. M., Zou, M. E., Lim, J. P., Stecher, J., McMahon, T., and Messing, R. O. (2014). Deletion of Prkcz increases intermittent ethanol consumption in mice. *Alcohol. Clin. Exp. Res.* 38, 170–178. doi:10.1111/acer.12211.

Lee, B.-T., Lee, H.-Y., Lee, B.-C., Pae, C.-U., Yoon, B.-J., Ryu, S.-G., et al. (2009). Impact of the tryptophan hydroxylase 1 gene A218C polymorphism on amygdala activity in response to affective facial stimuli in patients with major depressive disorder. *Genes Brain Behav.* 8, 512–518. doi:10.1111/j.1601-183X.2009.00500.x.

Lee, K. F., Simon, H., Chen, H., Bates, B., Hung, M. C., and Hauser, C. (1995). Requirement for neuregulin receptor erbB2 in neural and cardiac development. *Nature* 378, 394–398. doi:10.1038/378394a0.

Lee, Y. H., Kim, J.-H., and Song, G. G. (2013b). Pathway analysis of a genome-wide association study in schizophrenia. *Gene* 525, 107–115. doi:10.1016/j.gene.2013.04.014.

Le-Niculescu, H., Kurian, S. M., Yehyawi, N., Dike, C., Patel, S. D., Edenberg, H. J., et al. (2009). Identifying blood biomarkers for mood disorders using convergent functional genomics. *Mol. Psychiatry* 14, 156–174. doi:10.1038/mp.2008.11.

Leno-Colorado, J., Hudson, N. J., Reverter, A., and Pérez-Enciso, M. (2017). A Pathway-Centered Analysis of Pig Domestication and Breeding in Eurasia. *G3 (Bethesda)* 7, 2171–2184. doi:10.1534/g3.117.042671.

Levinson, D. F., Holmans, P., Straub, R. E., Owen, M. J., Wildenauer, D. B., Gejman, P. V., et al. (2000). Multicenter Linkage Study of Schizophrenia Candidate Regions on Chromosomes 5q, 6q, 10p, and 13q: Schizophrenia Linkage Collaborative Group III. *Am J Hum Genet* 67, 652–663.

Levy, B., Tegay, D., Papenhausen, P., Tepperberg, J., Nahum, O., Tsuchida, T., et al. (2012). Tetrasomy 15q26: a distinct syndrome or Shprintzen-Goldberg syndrome phenocopy? *Genet Med* 14, 811–818. doi:10.1038/gim.2012.54.

Levy-Shraga, Y., Gothelf, D., Pinchevski-Kadir, S., Katz, U., and Modan-Moses, D. (2018). Endocrine manifestations in children with Williams-Beuren syndrome. *Acta Paediatr.* 107, 678–684. doi:10.1111/apa.14198.

Lewis, C. M., Levinson, D. F., Wise, L. H., DeLisi, L. E., Straub, R. E., Hovatta, I., et al. (2003). Genome scan meta-analysis of schizophrenia and bipolar disorder, part II: Schizophrenia. *Am. J. Hum. Genet.* 73, 34–48. doi:10.1086/376549.

Li, D., and He, L. (2006). Further clarification of the contribution of the tryptophan hydroxylase (TPH) gene to suicidal behavior using systematic allelic and genotypic meta-analyses. *Hum. Genet.* 119, 233–240. doi:10.1007/s00439-005-0113-x.

Li, H., Chou, S.-J., Hamasaki, T., Perez-Garcia, C. G., and O'Leary, D. D. (2012). Neuregulin repellent signaling via ErbB4 restricts GABAergic interneurons to migratory paths from ganglionic eminence to cortical destinations. *Neural Dev* 7, 10. doi:10.1186/1749-8104-7-10.

Li, M., Shen, L., Chen, L., Huai, C., Huang, H., Wu, X., et al. (2020). Novel genetic susceptibility loci identified by family based whole exome sequencing in Han Chinese schizophrenia patients. *Transl Psychiatry* 10, 5. doi:10.1038/s41398-020-0708-y.

Li, Y., vonHoldt, B. M., Reynolds, A., Boyko, A. R., Wayne, R. K., Wu, D.-D., et al. (2013). Artificial Selection on Brain-Expressed Genes during the Domestication of Dog. *Mol Biol Evol* 30, 1867–1876. doi:10.1093/molbev/mst088.

Li, Y., Wang, G.-D., Wang, M.-S., Irwin, D. M., Wu, D.-D., and Zhang, Y.-P. (2014). Domestication of the Dog from the Wolf Was Promoted by Enhanced Excitatory Synaptic Plasticity: A Hypothesis. *Genome Biol Evol* 6, 3115–3121. doi:10.1093/gbe/evu245.

Li, Y., Xue, Y., He, Y., Li, F., Xue, L., Xu, C., et al. (2011). Inhibition of PKMζ in Nucleus Accumbens Core Abolishes Long-Term Drug Reward Memory. *J Neurosci* 31, 5436–5446. doi:10.1523/JNEUROSCI.5884-10.2011.

Liebowitz, M. R., Schneier, F., Gitow, A., and Feerick, J. (1993). Reversible monoamine oxidase-A inhibitors in social phobia. *Clin Neuropharmacol* 16 Suppl 2, S83-88.

Liguori, M., Nuzziello, N., Simone, M., Amoroso, N., Viterbo, R. G., Tangaro, S., et al. (2019). Association between miRNAs expression and cognitive performances of Pediatric Multiple Sclerosis patients: A pilot study. *Brain Behav* 9, e01199. doi:10.1002/brb3.1199.

Lim, L. C. C., Gurling, H., Curtis, D., Brynjolfsson, J., Petursson, H., and Gill, M. (1993). Linkage between tyrosine hydroxylase gene and affective disorder cannot be excluded in two of six pedigrees. *American Journal of Medical Genetics* 48, 223–228. doi:10.1002/ajmg.1320480410.

Lim, L. C., Powell, J., Sham, P., Castle, D., Hunt, N., Murray, R., et al. (1995). Evidence for a genetic association between alleles of monoamine oxidase A gene and bipolar affective disorder. *Am. J. Med. Genet.* 60, 325–331. doi:10.1002/ajmg.1320600413.

Lin, S., Jiang, S., Wu, X., Qian, Y., Wang, D., Tang, G., et al. (2000). Association analysis between mood disorder and monoamine oxidase gene. *Am. J. Med. Genet.* 96, 12–14. doi:10.1002/(sici)1096-8628(20000207)96:1<12::aid-ajmg4>3.0.co;2-s.

Lionel, A. C., Tammimies, K., Vaags, A. K., Rosenfeld, J. A., Ahn, J. W., Merico, D., et al. (2014). Disruption of the ASTN2/TRIM32 locus at 9q33.1 is a risk factor in males for autism spectrum disorders, ADHD and other neurodevelopmental phenotypes. *Hum. Mol. Genet.* 23, 2752–2768. doi:10.1093/hmg/ddt669.

Litonjua, A. A., Gong, L., Duan, Q. L., Shin, J., Moore, M. J., Weiss, S. T., et al. (2010). Very important pharmacogene summary ADRB2. *Pharmacogenet Genomics*. 20(1), 64-9. doi: 10.1097/FPC.0b013e3.

Liu, J., Dietz, K., DeLoyht, J. M., Pedre, X., Kelkar, D., Kaur, J., et al. (2012a). Impaired adult myelination in the prefrontal cortex of socially isolated mice. *Nat Neurosci* 15, 1621–1623. doi:10.1038/nn.3263.

Liu, J., Julnes, P. S., Chen, J., Ehrlich, S., Walton, E., and Calhoun, V. D. (2015). The association of DNA methylation and brain volume in healthy individuals and schizophrenia patients. *Schizophr Res* 169, 447–452. doi:10.1016/j.schres.2015.08.035.

Liu, J., Ulloa, A., Perrone-Bizzozero, N., Yeo, R., Chen, J., and Calhoun, V. D. (2012b). A Pilot Study on Collective Effects of 22q13.31 Deletions on Gray Matter Concentration in Schizophrenia. *PLOS ONE* 7, e52865. doi:10.1371/journal.pone.0052865.

Liu, M., Fitzgibbon, M., Wang, Y., Reilly, J., Qian, X., O'Brien, T., et al. (2018). Ulk4 regulates GABAergic signaling and anxiety-related behavior. *Transl Psychiatry* 8, 43. doi:10.1038/s41398-017-0091-5.

Liu, W., Tang, B., Cao, G., Chen, T., and Li, H. (2004). [TH gene mutation in Chinese patients with autosomal recessive dopa-responsive dystonia]. *Zhonghua Yi Xue Yi Chuan Xue Za Zhi* 21, 452–454.

Liu, X., Zhang, T., He, S., Hong, B., Chen, Z., Peng, D., et al. (2014a). Elevated serum levels of FGF-2, NGF and IGF-1 in patients with manic episode of bipolar disorder. *Psychiatry Res* 218, 54–60. doi:10.1016/j.psychres.2014.03.042.

Liu, Y., Blackwood, D. H., Caesar, S., de Geus, E. J. C., Farmer, A., Ferreira, M. A. R., et al. (2011). Meta-Analysis of Genome-Wide Association Data of Bipolar Disorder and Major Depressive Disorder. *Mol Psychiatry* 16. doi:10.1038/mp.2009.107.

Liu, Y., Harding, M., Pittman, A., Dore, J., Striessnig, J., Rajadhyaksha, A., et al. (2014b). Cav1.2 and Cav1.3 L-type calcium channels regulate dopaminergic firing activity in the mouse ventral tegmental area. *J Neurophysiol* 112, 1119–1130. doi:10.1152/jn.00757.2013.

Liu, Y., Hu, Z., Xun, G., Peng, Y., Lu, L., Xu, X., et al. (2012c). Mutation analysis of the NRXN1 gene in a Chinese autism cohort. *J Psychiatr Res* 46, 630–634. doi:10.1016/j.jpsychires.2011.10.015.

Liu, Y., Wang, Z., and Zhang, B. (2014c). The relationship between monoamine oxidase B (MAOB) A644G polymorphism and Parkinson disease risk: a meta-analysis. *Ann Saudi Med* 34, 12–17. doi:10.5144/0256-4947.2014.12.

López, M. E., Benestan, L., Moore, J., Perrier, C., Gilbey, J., Di Genova, A., et al. (2018). Comparing genomic signatures of domestication in two Atlantic salmon (Salmo salar L.) populations with different geographical origins. *Evol Appl* 12, 137–156. doi:10.1111/eva.12689.

Løtvedt, P., Fallahshahroudi, A., Bektic, L., Altimiras, J., and Jensen, P. (2017). Chicken domestication changes expression of stress-related genes in brain, pituitary and adrenals. *Neurobiol Stress* 7, 113–121. doi:10.1016/j.ynstr.2017.08.002.

Low, D., and Chen, K.-S. (2010). Genome-wide gene expression profiling of the Angelman syndrome mice with Ube3a mutation. *Eur J Hum Genet* 18, 1228–1235. doi:10.1038/ejhg.2010.95.

Ma, H., Xun, G., Zhang, R., Yang, X., and Cao, Y. (2019a). Correlation between GRIK2 rs6922753, rs2227283 polymorphism and aggressive behaviors with Bipolar Mania in the Chinese Han population. *Brain Behav* 9. doi:10.1002/brb3.1449.

Ma, J., Zhang, L.-Q., He, Z.-X., He, X.-X., Wang, Y.-J., Jian, Y.-L., et al. (2019b). Autism candidate gene DIP2A regulates spine morphogenesis via acetylation of cortactin. *PLoS Biol* 17. doi:10.1371/journal.pbio.3000461.

Ma, Y., Li, J., Xu, Y., Wang, Y., Yao, Y., Liu, Q., et al. (2020). Identification of 34 genes conferring genetic and pharmacological risk for the comorbidity of schizophrenia and smoking behaviors. *Aging (Albany NY)* 12. doi:10.18632/aging.102735.

Macintyre, G., Alford, T., Xiong, L., Rouleau, G. A., Tibbo, P. G., and Cox, D. W. (2010). Association of NPAS3 exonic variation with schizophrenia. *Schizophr. Res.* 120, 143–149. doi:10.1016/j.schres.2010.04.002.

Maestrini, E., Pagnamenta, A. T., Lamb, J. A., Bacchelli, E., Sykes, N. H., Sousa, I., et al. (2010). High-density SNP association study and copy number variation analysis of the AUTS1 and AUTS5 loci implicate the IMMP2L-DOCK4 gene region in autism susceptibility. *Mol. Psychiatry* 15, 954–968. doi:10.1038/mp.2009.34.

Magri, C., Sacchetti, E., Traversa, M., Valsecchi, P., Gardella, R., Bonvicini, C., et al. (2010). New Copy Number Variations in Schizophrenia. *PLoS One* 5. doi:10.1371/journal.pone.0013422.

Mahdavi, M., Kheirollahi, M., Riahi, R., Khorvash, F., Khorrami, M., and Mirsafaie, M. (2018). Meta-Analysis of the Association between GABA Receptor Polymorphisms and Autism Spectrum Disorder (ASD). *J Mol Neurosci* 65, 1–9. doi:10.1007/s12031-018-1073-7.

Mallas, E., Carletti, F., Chaddock, C. A., Shergill, S., Woolley, J., Picchioni, M. M., et al. (2017). The impact of CACNA1C gene, and its epistasis with ZNF804A, on white matter microstructure in health, schizophrenia and bipolar disorder1. *Genes Brain Behav.* 16, 479–488. doi:10.1111/gbb.12355.

Maniatis, T. (1999). A ubiquitin ligase complex essential for the NF-κB, Wnt/Wingless, and Hedgehog signaling pathways. *Genes Dev.* 13, 505–510.

Manitt, C., Eng, C., Pokinko, M., Ryan, R. T., Torres-Berrío, A., Lopez, J. P., et al. (2013). dcc orchestrates the development of the prefrontal cortex during adolescence and is altered in psychiatric patients. *Transl Psychiatry* 3, e338. doi:10.1038/tp.2013.105.

Manuck, S. B., Flory, J. D., Ferrell, R. E., Mann, J. J., and Muldoon, M. F. (2000). A regulatory polymorphism of the monoamine oxidase-A gene may be associated with variability in aggression, impulsivity, and central nervous system serotonergic responsivity. *Psychiatry Research* 95, 9–23. doi:10.1016/S0165-1781(00)00162-1.

Marcantoni, A., Vandael, D. H. F., Mahapatra, S., Carabelli, V., Sinnegger-Brauns, M. J., Striessnig, J., et al. (2010). Loss of Cav1.3 Channels Reveals the Critical Role of L-Type and BK Channel Coupling in Pacemaking Mouse Adrenal Chromaffin Cells. *J Neurosci* 30, 491–504. doi:10.1523/JNEUROSCI.4961-09.2010.

Marenco, S., Siuta, M. A., Kippenhan, J. S., Grodofsky, S., Chang, W., Kohn, P., et al. (2007). Genetic contributions to white matter architecture revealed by diffusion tensor imaging in Williams syndrome. *PNAS* 104, 15117–15122. doi:10.1073/pnas.0704311104.

Mark, P. R., Radlinski, B. C., Core, N., Fryer, A., Kirk, E. P., and Haldeman-Englert, C. R. (2013). Narrowing the Critical Region for Congenital Vertical Talus in Patients With Interstitial 18q Deletions. *American Journal of Medical Genetics Part A* 161, 1117–1121. doi:10.1002/ajmg.a.35791.

Marley, A., and von Zastrow, M. (2012). A simple cell-based assay reveals that diverse neuropsychiatric risk genes converge on primary cilia. *PLoS ONE* 7, e46647. doi:10.1371/journal.pone.0046647.

Marschallinger, J., Sah, A., Schmuckermair, C., Unger, M., Rotheneichner, P., Kharitonova, M., et al. (2015). The L-type calcium channel Cav1.3 is required for proper hippocampal neurogenesis and cognitive functions. *Cell Calcium* 58, 606–616. doi:10.1016/j.ceca.2015.09.007.

Marshall, C. R., Howrigan, D. P., Merico, D., Thiruvahindrapuram, B., Wu, W., Greer, D. S., et al. (2017). Contribution of copy number variants to schizophrenia from a genome-wide study of 41,321 subjects. *Nat. Genet.* 49, 27–35. doi:10.1038/ng.3725.

Martin, C.-A., Murray, J. E., Carroll, P., Leitch, A., Mackenzie, K. J., Halachev, M., et al. (2016). Mutations in genes encoding condensin complex proteins cause microcephaly through decatenation failure at mitosis. *Genes Dev* 30, 2158–2172. doi:10.1101/gad.286351.116.

Martin, J., Cooper, M., Hamshere, M. L., Pocklington, A., Scherer, S. W., Kent, L., et al. (2014). Biological Overlap of Attention-Deficit/Hyperactivity Disorder and Autism Spectrum Disorder: Evidence From Copy Number Variants. *J Am Acad Child Adolesc Psychiatry* 53, 761-770.e26. doi:10.1016/j.jaac.2014.03.004.

Martínez-Rivera, A., Hao, J., Tropea, T. F., Giordano, T. P., Kosovsky, M., Rice, R. C., et al. (2017). Enhancing VTA Cav1.3 L-type Ca2+ channel activity promotes cocaine and mood - related behaviors via overlapping AMPA receptor mechanisms in the nucleus accumbens. *Mol Psychiatry* 22, 1735–1745. doi:10.1038/mp.2017.9.

Maruani, A., Huguet, G., Beggiato, A., ElMaleh, M., Toro, R., Leblond, C. S., et al. (2015). 11q24.2-25 micro-rearrangements in autism spectrum disorders: Relation to brain structures. *Am. J. Med. Genet. A* 167A, 3019–3030. doi:10.1002/ajmg.a.37345.

Masana, M., Jukic, M. M., Kretzschmar, A., Wagner, K. V., Westerholz, S., Schmidt, M. V., et al. (2015). Deciphering the spatio-temporal expression and stress regulation of Fam107B, the paralog of the resilience-promoting protein DRR1 in the mouse brain. *Neuroscience* 290, 147–158. doi:10.1016/j.neuroscience.2015.01.026.

Mattheisen, M., Samuels, J. F., Wang, Y., Greenberg, B. D., Fyer, A. J., McCracken, J. T., et al. (2015). Genome-Wide Association Study in Obsessive-Compulsive Disorder: Results from the OCGAS. *Mol Psychiatry* 20, 337–344. doi:10.1038/mp.2014.43.

Mattina, T., Perrotta, C. S., and Grossfeld, P. (2009). Jacobsen syndrome. *Orphanet J Rare Dis* 4, 9. doi:10.1186/1750-1172-4-9.

Maynard, K. R., and Stein, E. (2012). DSCAM Contributes to Dendrite Arborization and Spine Formation in the Developing Cerebral Cortex. *J. Neurosci.* 32, 16637–16650. doi:10.1523/JNEUROSCI.2811-12.2012.

McKinney, B. C., and Murphy, G. G. (2006). The L-Type voltage-gated calcium channel Cav1.3 mediates consolidation, but not extinction, of contextually conditioned fear in mice. *Learn Mem* 13, 584–589. doi:10.1101/lm.279006.

McLean, C. Y., Reno, P. L., Pollen, A. A., Bassan, A. I., Capellini, T. D., Guenther, C., et al. (2011). Human-specific loss of regulatory DNA and the evolution of human-specific traits. *Nature* 471, 216–219. doi:10.1038/nature09774.

Mead, A. N., and Stephens, D. N. (2003). Selective Disruption of Stimulus-Reward Learning in Glutamate Receptor gria1 Knock-Out Mice. *J Neurosci* 23, 1041–1048. doi:10.1523/JNEUROSCI.23-03-01041.2003.

Meador-Woodruff, J. H., Davis, K. L., and Haroutunian, V. (2001). Abnormal kainate receptor expression in prefrontal cortex in schizophrenia. *Neuropsychopharmacology* 24, 545–552. doi:10.1016/S0893-133X(00)00189-5.

Mei, L., and Nave, K.-A. (2014). Neuregulin-ERBB signaling in nervous system development and neuropsychiatric diseases. *Neuron* 83, 27–49. doi:10.1016/j.neuron.2014.06.007.

Mendlewicz, J., and Youdim, M. B. (1983). L-Deprenil, a selective monoamine oxidase type B inhibitor, in the treatment of depression: a double blind evaluation. *Br J Psychiatry* 142, 508–511. doi:10.1192/bjp.142.5.508.

Menold, M. M., Shao, Y., Wolpert, C. M., Donnelly, S. L., Raiford, K. L., Martin, E. R., et al. (2001). Association analysis of chromosome 15 gabaa receptor subunit genes in autistic disorder. *J. Neurogenet.* 15, 245–259. doi:10.3109/01677060109167380.

Mercati, O., Huguet, G., Danckaert, A., André-Leroux, G., Maruani, A., Bellinzoni, M., et al. (2017). CNTN6 mutations are risk factors for abnormal auditory sensory perception in autism spectrum disorders. *Mol. Psychiatry* 22, 625–633. doi:10.1038/mp.2016.61.

Mesbah-Oskui, L., Penna, A., Orser, B. A., and Horner, R. L. (2017). Reduced expression of α5GABAA receptors elicits autism-like alterations in EEG patterns and sleep-wake behavior. *Neurotoxicol Teratol* 61, 115–122. doi:10.1016/j.ntt.2016.10.009.

Michaelson, J. J., Shin, M.-K., Koh, J.-Y., Brueggeman, L., Zhang, A., Katzman, A., et al. (2017). Neuronal PAS Domain Proteins 1 and 3 Are Master Regulators of Neuropsychiatric Risk Genes. *Biol Psychiatry* 82, 213–223. doi:10.1016/j.biopsych.2017.03.021.

Migues, P. V., Hardt, O., Wu, D. C., Gamache, K., Sacktor, T. C., Wang, Y. T., et al. (2010). PKMzeta maintains memories by regulating GluR2-dependent AMPA receptor trafficking. *Nat. Neurosci.* 13, 630–634. doi:10.1038/nn.2531.

Miller, C. L., Murakami, P., Ruczinski, I., Ross, R. G., Sinkus, M., Sullivan, B., et al. (2009). Two Complex Genotypes Relevant to the Kynurenine Pathway and Melanotropin Function show Association with Schizophrenia and Bipolar Disorder. *Schizophr Res* 113, 259–267. doi:10.1016/j.schres.2009.05.014.

Mimica, N., Mück-Seler, D., Pivac, N., Mustapić, M., Dezeljin, M., Stipcević, T., et al. (2008). Platelet serotonin and monoamine oxidase in Alzheimer's disease with psychotic features. *Coll Antropol* 32 Suppl 1, 119–122.

Mio, C., Passon, N., Baldan, F., Bregant, E., Monaco, E., Mancini, L., et al. (2020). CACNA1C haploinsufficiency accounts for the common features of interstitial 12p13.33 deletion carriers. *Eur J Med Genet,* 103843. doi:10.1016/j.ejmg.2020.103843.

Mochida, G. H., Ganesh, V. S., Felie, J. M., Gleason, D., Hill, R. S., Clapham, K. R., et al. (2010). A Homozygous Mutation in the Tight-Junction Protein JAM3 Causes Hemorrhagic Destruction of the Brain, Subependymal Calcification, and Congenital Cataracts. *Am J Hum Genet* 87, 882–889. doi:10.1016/j.ajhg.2010.10.026.

Monfrim, X., Gazal, M., De Leon, P. B., Quevedo, L., Souza, L. D., Jansen, K., et al. (2014). Immune dysfunction in bipolar disorder and suicide risk: is there an association between peripheral corticotropin-releasing hormone and interleukin-1β? *Bipolar Disord* 16, 741–747. doi:10.1111/bdi.12214.

Monk, K. R., Voas, M. G., Franzini-Armstrong, C., Hakkinen, I. S., and Talbot, W. S. (2013). Mutation of sec63 in zebrafish causes defects in myelinated axons and liver pathology. *Dis Model Mech* 6, 135–145. doi:10.1242/dmm.009217.

Montague, M. J., Li, G., Gandolfi, B., Khan, R., Aken, B. L., Searle, S. M. J., et al. (2014). Comparative analysis of the domestic cat genome reveals genetic signatures underlying feline biology and domestication. *Proc Natl Acad Sci U S A* 111, 17230–17235. doi:10.1073/pnas.1410083111.

Montgomery, S. A. (1999). Social phobia: diagnosis, severity and implications for treatment. *Eur Arch Psychiatry Clin Neurosci* 249 Suppl 1, S1-6. doi:10.1007/pl00014161.

Moskvina, V., Craddock, N., Holmans, P., Nikolov, I., Pahwa, J. S., Green, E., et al. (2009). Gene-wide analyses of genome-wide association datasets: evidence for multiple common risk alleles for schizophrenia and bipolar disorder and for overlap in genetic risk. *Mol Psychiatry* 14, 252–260. doi:10.1038/mp.2008.133.

Motazacker, M. M., Rost, B. R., Hucho, T., Garshasbi, M., Kahrizi, K., Ullmann, R., et al. (2007). A Defect in the Ionotropic Glutamate Receptor 6 Gene (GRIK2) Is Associated with Autosomal Recessive Mental Retardation. *Am J Hum Genet* 81, 792–798.

Moullé, V. S. F., Cansell, C., Luquet, S., and Cruciani-Guglielmacci, C. (2012). The multiple roles of fatty acid handling proteins in brain. *Front Physiol* 3. doi:10.3389/fphys.2012.00385.

Mowrey, D. D., Cui, T., Jia, Y., Ma, D., Makhov, A. M., Zhang, P., et al. (2013). Open-channel structures of the human glycine receptor α1 full-length transmembrane domain. *Structure* 21, 1897–1904. doi:10.1016/j.str.2013.07.014.

Moy, S. S., Troy Ghashghaei, H., Nonneman, R. J., Weimer, J. M., Yokota, Y., Lee, D., et al. (2009). Deficient NRG1-ERBB signaling alters social approach: relevance to genetic mouse models of schizophrenia. *J Neurodev Disord* 1, 302–312. doi:10.1007/s11689-009-9017-8.

Muck-Seler, D., Sagud, M., Mustapic, M., Nedic, G., Babic, A., Mihaljevic Peles, A., et al. (2008). The effect of lamotrigine on platelet monoamine oxidase type B activity in patients with bipolar depression. *Prog. Neuropsychopharmacol. Biol. Psychiatry* 32, 1195–1198. doi:10.1016/j.pnpbp.2008.03.004.

Mukherjee, S., Perumal, T. M., Daily, K., Sieberts, S. K., Omberg, L., Preuss, C., et al. (2019). Identifying and ranking potential driver genes of Alzheimer’s disease using multiview evidence aggregation. *Bioinformatics* 35, i568–i576. doi:10.1093/bioinformatics/btz365.

Munafò, M. R., Thiselton, D. L., Clark, T. G., and Flint, J. (2006). Association of the NRG1 gene and schizophrenia: a meta-analysis. *Molecular Psychiatry* 11, 539–546. doi:10.1038/sj.mp.4001817.

Murphy, E. (2015). The brain dynamics of linguistic computation. *Front. Psychol.* 6. doi:10.3389/fpsyg.2015.01515.

Murphy, E., and Benítez-Burraco, A. (2016). Bridging the Gap between Genes and Language Deficits in Schizophrenia: An Oscillopathic Approach. *Front Hum Neurosci* 10. doi:10.3389/fnhum.2016.00422.

Murphy, E., and Benítez-Burraco, A. (2017). Language deficits in schizophrenia and autism as related oscillatory connectomopathies: An evolutionary account. *Neuroscience & Biobehavioral Reviews* 83, 742–764. doi:10.1016/j.neubiorev.2016.07.029.

Murphy, E., and Benítez-Burraco, A. (2018). Paleo-oscillomics: inferring aspects of Neanderthal language abilities from gene regulation of neural oscillations. *J Anthropol Sci* 96, 111–124. doi:10.4436/JASS.96010.

Nagatsu, T. (1995). Tyrosine hydroxylase: human isoforms, structure and regulation in physiology and pathology. *Essays Biochem.* 30, 15–35.

Nagatsu, T., and Ichinose, H. (1991). Comparative studies on the structure of human tyrosine hydroxylase with those of the enzyme of various mammals. *Comp. Biochem. Physiol. C, Comp. Pharmacol. Toxicol.* 98, 203–210.

Nakajima, H., and Koizumi, K. (2014). Family with sequence similarity 107: A family of stress responsive small proteins with diverse functions in cancer and the nervous system (Review). *Biomed Rep* 2, 321–325. doi:10.3892/br.2014.243.

Nakajima, H., Koizumi, K., Tanaka, T., Ishigaki, Y., Yoshitake, Y., Yonekura, H., et al. (2012). Loss of HITS (FAM107B) expression in cancers of multiple organs: tissue microarray analysis. *Int. J. Oncol.* 41, 1347–1357. doi:10.3892/ijo.2012.1550.

Nakamura, K., Sugawara, Y., Sawabe, K., Ohashi, A., Tsurui, H., Xiu, Y., et al. (2006). Late Developmental Stage-Specific Role of Tryptophan Hydroxylase 1 in Brain Serotonin Levels. *J Neurosci* 26, 530–534. doi:10.1523/JNEUROSCI.1835-05.2006.

Nam, K., Mugal, C., Nabholz, B., Schielzeth, H., Wolf, J. B., Backström, N., et al. (2010). Molecular evolution of genes in avian genomes. *Genome Biol* 11, R68. doi:10.1186/gb-2010-11-6-r68.

Napolitano, C., Splawski, I., Timothy, K. W., Bloise, R., and Priori, S. G. (2015). *Timothy Syndrome*. University of Washington, Seattle Available at: https://www.ncbi.nlm.nih.gov/books/NBK1403/ [Accessed July 28, 2019].

Naqvi, S., Cole, T., and Graham, J. M. (2000). Cole-Hughes macrocephaly syndrome and associated autistic manifestations. *Am. J. Med. Genet.* 94, 149–152. doi:10.1002/1096-8628(20000911)94:2<149::aid-ajmg7>3.0.co;2-#.

Need, A. C., Ge, D., Weale, M. E., Maia, J., Feng, S., Heinzen, E. L., et al. (2009). A Genome-Wide Investigation of SNPs and CNVs in Schizophrenia. *PLOS Genetics* 5, e1000373. doi:10.1371/journal.pgen.1000373.

New, A. S., Gelernter, J., Yovell, Y., Trestman, R. L., Nielsen, D. A., Silverman, J., et al. (1998). Tryptophan hydroxylase genotype is associated with impulsive-aggression measures: a preliminary study. *Am. J. Med. Genet.* 81, 13–17. doi:10.1002/(sici)1096-8628(19980207)81:1<13::aid-ajmg3>3.0.co;2-o.

Ng, M., Levinson, D., Faraone, S., Suarez, B., DeLisi, L., Arinami, T., et al. (2009). Meta-analysis of 32 genome-wide linkage studies of schizophrenia. *Mol Psychiatry* 14, 774–785. doi:10.1038/mp.2008.135.

Nguyen, C. L., Possemato, R., Bauerlein, E. L., Xie, A., Scully, R., and Hahn, W. C. (2012). Nek4 Regulates Entry into Replicative Senescence and the Response to DNA Damage in Human Fibroblasts. *Mol Cell Biol* 32, 3963–3977. doi:10.1128/MCB.00436-12.

Niculescu, A. B., Levey, D. F., Phalen, P. L., Le-Niculescu, H., Dainton, H. D., Jain, N., et al. (2015). Understanding and predicting suicidality using a combined genomic and clinical risk assessment approach. *Mol. Psychiatry* 20, 1266–1285. doi:10.1038/mp.2015.112.

Niego, A., and Benítez-Burraco, A. (2019). Williams Syndrome, Human Self-Domestication, and Language Evolution. *Front. Psychol.* 10. doi:10.3389/fpsyg.2019.00521.

Nisar, S., Hashem, S., Bhat, A. A., Syed, N., Yadav, S., Azeem, M. W., et al. (2019). Association of genes with phenotype in autism spectrum disorder. *Aging (Albany NY)* 11, 10742–10770. doi:10.18632/aging.102473.

Nishimura, Y., Martin, C. L., Vazquez-Lopez, A., Spence, S. J., Alvarez-Retuerto, A. I., Sigman, M., et al. (2007). Genome-wide expression profiling of lymphoblastoid cell lines distinguishes different forms of autism and reveals shared pathways. *Hum Mol Genet* 16, 1682–1698. doi:10.1093/hmg/ddm116.

Noor, A., Lionel, A. C., Cohen-Woods, S., Moghimi, N., Rucker, J., Fennell, A., et al. (2014). Copy number variant study of bipolar disorder in Canadian and UK populations implicates synaptic genes. *Am. J. Med. Genet. B Neuropsychiatr. Genet.* 165B, 303–313. doi:10.1002/ajmg.b.32232.

Nucifora, L. G., Wu, Y. C., Lee, B. J., Sha, L., Margolis, R. L., Ross, C. A., et al. (2016). A Mutation in NPAS3 That Segregates with Schizophrenia in a Small Family Leads to Protein Aggregation. *Mol Neuropsychiatry* 2, 133–144. doi:10.1159/000447358.

Nurnberger, J. I., Koller, D. L., Jung, J., Edenberg, H. J., Foroud, T., Guella, I., et al. (2014). Identification of Pathways for Bipolar Disorder A Meta-analysis. *JAMA Psychiatry* 71, 657–664. doi:10.1001/jamapsychiatry.2014.176.

O'Connor, J. A., and Hemby, S. E. (2007). Elevated GRIA1 mRNA expression in Layer II/III and V pyramidal cells of the DLPFC in schizophrenia. *Schizophrenia Research* 97, 277. doi:10.1016/j.schres.2007.09.022.

O'Donnell, L., Soileau, B., Heard, P., Carter, E., Sebold, C., Gelfond, J., et al. (2010). Genetic determinants of autism in individuals with deletions of 18q. *Hum Genet* 128, 155–164. doi:10.1007/s00439-010-0839-y.

O'Dushlaine, C., Kenny, E., Heron, E., Donohoe, G., Gill, M., Morris, D., et al. (2011). Molecular pathways involved in neuronal cell adhesion and membrane scaffolding contribute to schizophrenia and bipolar disorder susceptibility. *Mol. Psychiatry* 16, 286–292. doi:10.1038/mp.2010.7.

Oguro-Ando, A., Zuko, A., Kleijer, K. T. E., and Burbach, J. P. H. (2017). A current view on contactin-4, -5, and -6: Implications in neurodevelopmental disorders. *Mol. Cell. Neurosci.* 81, 72–83. doi:10.1016/j.mcn.2016.12.004.

Ohtsuki, T., Ishiguro, H., Detera-Wadleigh, S. D., Toyota, T., Shimizu, H., Yamada, K., et al. (2002). Association between serotonin 4 receptor gene polymorphisms and bipolar disorder in Japanese case-control samples and the NIMH Genetics Initiative Bipolar Pedigrees. *Molecular Psychiatry* 7, 954–961. doi:10.1038/sj.mp.4001133.

O'Roak, B. J., Vives, L., Girirajan, S., Karakoc, E., Krumm, N., Coe, B. P., et al. (2012). Sporadic autism exomes reveal a highly interconnected protein network of de novo mutations. *Nature* 485, 246–250. doi:10.1038/nature10989.

O'Rourke, T., and Boeckx, C. (2020). Glutamate receptors in domestication and modern human evolution. *Neuroscience & Biobehavioral Reviews* 108, 341–357. doi:10.1016/j.neubiorev.2019.10.004.

Oruc, L., Verheyen, G. R., Furac, I., Ivezić, S., Jakovljević, M., Raeymaekers, P., et al. (1997). Positive association between the GABRA5 gene and unipolar recurrent major depression. *Neuropsychobiology* 36, 62–64. doi:10.1159/000119363.

Otani, K., Ujike, H., Tanaka, Y., Morita, Y., Katsu, T., Nomura, A., et al. (2005). The GABA type A receptor α5 subunit gene is associated with bipolar I disorder. *Neuroscience Letters* 381, 108–113. doi:10.1016/j.neulet.2005.02.010.

Pai, S., Li, P., Killinger, B., Marshall, L., Jia, P., Liao, J., et al. (2019). Differential methylation of enhancer at IGF2 is associated with abnormal dopamine synthesis in major psychosis. *Nat Commun* 10, 2046. doi:10.1038/s41467-019-09786-7.

Palomino, A., González-Pinto, A., Martinez-Cengotitabengoa, M., Ruiz de Azua, S., Alberich, S., Mosquera, F., et al. (2013). Relationship between negative symptoms and plasma levels of insulin-like growth factor 1 in first-episode schizophrenia and bipolar disorder patients. *Prog. Neuropsychopharmacol. Biol. Psychiatry* 44, 29–33. doi:10.1016/j.pnpbp.2013.01.008.

Pan, Y., Wang, K.-S., and Aragam, N. (2011). NTM and NR3C2 polymorphisms influencing intelligence: family-based association studies. *Prog. Neuropsychopharmacol. Biol. Psychiatry* 35, 154–160. doi:10.1016/j.pnpbp.2010.10.016.

Papadimitriou, G., Dikeos, D., Daskalopoulou, E., Karadima, G., Avramopoulos, D., Contis, C., et al. (2001a). Association between GABA-A receptor alpha 5 subunit gene locus and schizophrenia of a later age of onset. *Neuropsychobiology* 43, 141–144. doi:10.1159/000054882.

Papadimitriou, G. N., Dikeos, D. G., Karadima, G., Avramopoulos, D., Daskalopoulou, E. G., and Stefanis, C. N. (2001b). GABA-A receptor beta3 and alpha5 subunit gene cluster on chromosome 15q11-q13 and bipolar disorder: a genetic association study. *Am. J. Med. Genet.* 105, 317–320. doi:10.1002/ajmg.1354.

Papadimitriou, G. N., Dikeos, D. G., Karadima, G., Avramopoulos, D., Daskalopoulou, E. G., Vassilopoulos, D., et al. (1998). Association between the GABA(A) receptor alpha5 subunit gene locus (GABRA5) and bipolar affective disorder. *Am. J. Med. Genet.* 81, 73–80.

Pardiñas, A. F., Holmans, P., Pocklington, A. J., Escott-Price, V., Ripke, S., Carrera, N., et al. (2018). Common schizophrenia alleles are enriched in mutation-intolerant genes and in regions under strong background selection. *Nat Genet* 50, 381–389. doi:10.1038/s41588-018-0059-2.

Parekh, P. K., Becker-Krail, D., Sundaravelu, P., Ishigaki, S., Okado, H., Sobue, G., et al. (2018). Altered GluA1 function and accumbal synaptic plasticity in the ClockΔ19 model of bipolar mania. *Biol Psychiatry* 84, 817–826. doi:10.1016/j.biopsych.2017.06.022.

Park, S., Lim, Y., Lee, D., Cho, B., Bang, Y.-J., Sung, S., et al. (2003). Identification and characterization of a novel cancer/testis antigen gene CAGE-1. *Biochimica et Biophysica Acta (BBA) - Gene Structure and Expression* 1625, 173–182. doi:10.1016/S0167-4781(02)00620-6.

Paucar, M., Waldthaler, J., and Svenningsson, P. (2018). GLRA1 mutation and long-term follow-up of the first hyperekplexia family. *Neurol Genet* 4. doi:10.1212/NXG.0000000000000259.

Payer, D., Williams, B., Mansouri, E., Stevanovski, S., Nakajima, S., Le Foll, B., et al. (2017). Corticotropin-releasing hormone and dopamine release in healthy individuals. *Psychoneuroendocrinology* 76, 192–196. doi:10.1016/j.psyneuen.2016.11.034.

Peche, V. S., Holak, T. A., Burgute, B. D., Kosmas, K., Kale, S. P., Wunderlich, F. T., et al. (2013). Ablation of cyclase-associated protein 2 (CAP2) leads to cardiomyopathy. *Cell. Mol. Life Sci.* 70, 527–543. doi:10.1007/s00018-012-1142-y.

Pendleton, A. L., Shen, F., Taravella, A. M., Emery, S., Veeramah, K. R., Boyko, A. R., et al. (2018). Comparison of village dog and wolf genomes highlights the role of the neural crest in dog domestication. *BMC Biology* 16, 64. doi:10.1186/s12915-018-0535-2.

Penzes, P., Cahill, M. E., Jones, K. A., VanLeeuwen, J.-E., and Woolfrey, K. M. (2011). Dendritic spine pathology in neuropsychiatric disorders. *Nat Neurosci* 14, 285–293. doi:10.1038/nn.2741.

Pereira, A. C. P., McQuillin, A., Puri, V., Anjorin, A., Bass, N., Kandaswamy, R., et al. (2011). Genetic association and sequencing of the insulin-like growth factor 1 gene in bipolar affective disorder. *Am. J. Med. Genet. B Neuropsychiatr. Genet.* 156, 177–187. doi:10.1002/ajmg.b.31153.

Petek, E., Schwarzbraun, T., Noor, A., Patel, M., Nakabayashi, K., Choufani, S., et al. (2007). Molecular and genomic studies of IMMP2L and mutation screening in autism and Tourette syndrome. *Mol. Genet. Genomics* 277, 71–81. doi:10.1007/s00438-006-0173-1.

Petek, E., Windpassinger, C., Vincent, J. B., Cheung, J., Boright, A. P., Scherer, S. W., et al. (2001). Disruption of a novel gene (IMMP2L) by a breakpoint in 7q31 associated with Tourette syndrome. *Am. J. Hum. Genet.* 68, 848–858. doi:10.1086/319523.

Peters, E. J., Slager, S. L., McGrath, P. J., Knowles, J. A., and Hamilton, S. P. (2004). Investigation of serotonin-related genes in antidepressant response. *Mol. Psychiatry* 9, 879–889. doi:10.1038/sj.mp.4001502.

Pickard, B. S., Christoforou, A., Thomson, P. A., Fawkes, A., Evans, K. L., Morris, S. W., et al. (2009). Interacting haplotypes at the NPAS3 locus alter risk of schizophrenia and bipolar disorder. *Molecular Psychiatry* 14, 874–884. doi:10.1038/mp.2008.24.

Pickard, B. S., Malloy, M. P., Porteous, D. J., Blackwood, D. H. R., and Muir, W. J. (2005). Disruption of a brain transcription factor, NPAS3, is associated with schizophrenia and learning disability. *Am. J. Med. Genet. B Neuropsychiatr. Genet.* 136B, 26–32. doi:10.1002/ajmg.b.30204.

Pickard, B. S., Pieper, A. A., Porteous, D. J., Blackwood, D. H., and Muir, W. J. (2006). The NPAS3 gene—emerging evidence for a role in psychiatric illness. *Annals of Medicine* 38, 439–448. doi:10.1080/07853890600946500.

Pickrell, J. K., Coop, G., Novembre, J., Kudaravalli, S., Li, J. Z., Absher, D., et al. (2009). Signals of recent positive selection in a worldwide sample of human populations. *Genome Res.* 19, 826–837. doi:10.1101/gr.087577.108.

Pilorge, M., Fassier, C., Le Corronc, H., Potey, A., Bai, J., De Gois, S., et al. (2016). Genetic and functional analyses demonstrate a role for abnormal glycinergic signaling in autism. *Mol Psychiatry* 21, 936–945. doi:10.1038/mp.2015.139.

Pilot, M., Malewski, T., Moura, A. E., Grzybowski, T., Oleński, K., Kamiński, S., et al. (2016). Diversifying Selection Between Pure-Breed and Free-Breeding Dogs Inferred from Genome-Wide SNP Analysis. *G3 (Bethesda)* 6, 2285–2298. doi:10.1534/g3.116.029678.

Pinggera, A., Lieb, A., Benedetti, B., Lampert, M., Monteleone, S., Liedl, K. R., et al. (2015). CACNA1D De Novo Mutations in Autism Spectrum Disorders Activate Cav1.3 L-Type Calcium Channels. *Biol Psychiatry* 77, 816–822. doi:10.1016/j.biopsych.2014.11.020.

Pinggera, A., Mackenroth, L., Rump, A., Schallner, J., Beleggia, F., Wollnik, B., et al. (2017). New gain-of-function mutation shows CACNA1D as recurrently mutated gene in autism spectrum disorders and epilepsy. *Hum Mol Genet* 26, 2923–2932. doi:10.1093/hmg/ddx175.

Pini, G., Scusa, M. F., Benincasa, A., Bottiglioni, I., Congiu, L., Vadhatpour, C., et al. (2014). Repeated insulin-like growth factor 1 treatment in a patient with rett syndrome: a single case study. *Front Pediatr* 2, 52. doi:10.3389/fped.2014.00052.

Pivac, N., Knezevic, J., Kozaric-Kovacic, D., Dezeljin, M., Mustapic, M., Rak, D., et al. (2007). Monoamine oxidase (MAO) intron 13 polymorphism and platelet MAO-B activity in combat-related posttraumatic stress disorder. *J Affect Disord* 103, 131–138. doi:10.1016/j.jad.2007.01.017.

Ponder, C. A., Kliethermes, C. L., Drew, M. R., Muller, J., Das, K., Risbrough, V. B., et al. (2007). Selection for contextual fear conditioning affects anxiety-like behaviors and gene expression. *Genes Brain Behav*. 6, 736–749. doi:10.1111/j.1601-183X.2007.00306.x.

Popova, N. K., Voitenko, N. N., Kulikov, A. V., and Avgustinovich, D. F. (1991). Evidence for the involvement of central serotonin in mechanism of domestication of silver foxes. *Pharmacol. Biochem. Behav*. 40, 751–756. doi:10.1016/0091-3057(91)90080-l.

Poot, M. (2014). A Candidate Gene Association Study Further Corroborates Involvement of Contactin Genes in Autism. *Mol Syndromol* 5, 229–235. doi:10.1159/000362891.

Powers, J. F., Picard, K. L., and Tischler, A. S. (2009). RET expression and neuron-like differentiation of pheochromocytoma and normal chromaffin cells. *Horm. Metab. Res.* 41, 710–714. doi:10.1055/s-0029-1224136.

Prasad, A., Merico, D., Thiruvahindrapuram, B., Wei, J., Lionel, A. C., Sato, D., et al. (2012). A discovery resource of rare copy number variations in individuals with autism spectrum disorder. *G3 (Bethesda)* 2, 1665–1685. doi:10.1534/g3.112.004689.

Preisig, M., Bellivier, F., Fenton, B. T., Baud, P., Berney, A., Courtet, P., et al. (2000). Association between bipolar disorder and monoamine oxidase A gene polymorphisms: results of a multicenter study. *Am J Psychiatry* 157, 948–955. doi:10.1176/appi.ajp.157.6.948.

Presse, F., Conductier, G., Rovere, C., and Nahon, J.-L. (2014). The melanin-concentrating hormone receptors: neuronal and non-neuronal functions. *Int J Obes Suppl* 4, S31–S36. doi:10.1038/ijosup.2014.9.

Prom-Wormley, E. C., Eaves, L. J., Foley, D. L., Gardner, C. O., Archer, K. J., Wormley, B. K., et al. (2009). Monoamine oxidase A and childhood adversity as risk factors for conduct disorder in females. *Psychol Med* 39, 579–590. doi:10.1017/S0033291708004170.

Psychosis Endophenotypes International Consortium, and Wellcome Trust Case-Control Consortium (2014). A Genome-wide Association Analysis of a Broad Psychosis Phenotype Identifies Three Loci for Further Investigation. *Biol Psychiatry* 75, 386–397. doi:10.1016/j.biopsych.2013.03.033.

PubChem ADRB2 - adrenoceptor beta 2 (human). Available at: https://pubchem.ncbi.nlm.nih.gov/gene/ADRB2/human [Accessed January 18, 2020].

Purves-Tyson, T. D., Owens, S. J., Rothmond, D. A., Halliday, G. M., Double, K. L., Stevens, J., et al. (2017). Putative presynaptic dopamine dysregulation in schizophrenia is supported by molecular evidence from post-mortem human midbrain. *Transl Psychiatry* 7, e1003. doi:10.1038/tp.2016.257.

Qanbari, S., Pausch, H., Jansen, S., Somel, M., Strom, T. M., Fries, R., et al. (2014). Classic Selective Sweeps Revealed by Massive Sequencing in Cattle. *PLOS Genetics* 10, e1004148. doi:10.1371/journal.pgen.1004148.

Quartier, A., Chatrousse, L., Redin, C., Keime, C., Haumesser, N., Maglott-Roth, A., et al. (2018). Genes and Pathways Regulated by Androgens in Human Neural Cells, Potential Candidates for the Male Excess in Autism Spectrum Disorder. *Biol. Psychiatry* 84, 239–252. doi:10.1016/j.biopsych.2018.01.002.

Quilter, C. R., Sargent, C. A., Bauer, J., Bagga, M. R., Reiter, C. P., Hutchinson, E. L., et al. (2012). An association and haplotype analysis of porcine maternal infanticide: A model for human puerperal psychosis? *American Journal of Medical Genetics Part B: Neuropsychiatric Genetics* 159B, 908–927. doi:10.1002/ajmg.b.32097.

Rabaneda, L. G., Robles-Lanuza, E., Nieto-González, J. L., and Scholl, F. G. (2014). Neurexin Dysfunction in Adult Neurons Results in Autistic-like Behavior in Mice. *Cell Reports* 8, 338–346. doi:10.1016/j.celrep.2014.06.022.

Racimo, F. (2016). Testing for Ancient Selection Using Cross-population Allele Frequency Differentiation. *Genetics* 202, 733–750. doi:10.1534/genetics.115.178095.

Raffan, E., Dennis, R. J., O'Donovan, C. J., Becker, J. M., Scott, R. A., Smith, S. P., et al. (2016). A Deletion in the Canine POMC Gene Is Associated with Weight and Appetite in Obesity-Prone Labrador Retriever Dogs. *Cell Metab* 23, 893–900. doi:10.1016/j.cmet.2016.04.012.

Raghanti, M. A., Edler, M. K., Stephenson, A. R., Wilson, L. J., Hopkins, W. D., Ely, J. J., et al. (2016). Human-specific increase of dopaminergic innervation in a striatal region associated with speech and language: a comparative analysis of the primate basal ganglia. *J Comp Neurol* 524, 2117–2129. doi:10.1002/cne.23937.

Rajkumar, A. P., Christensen, J. H., Mattheisen, M., Jacobsen, I., Bache, I., Pallesen, J., et al. (2015). Analysis of t(9;17)(q33.2;q25.3) chromosomal breakpoint regions and genetic association reveals novel candidate genes for bipolar disorder. *Bipolar Disord* 17, 205–211. doi:10.1111/bdi.12239.

Ramachandran, K. V., Hennessey, J. A., Barnett, A. S., Yin, X., Stadt, H. A., Foster, E., et al. (2013). Calcium influx through L-type CaV1.2 Ca2+ channels regulates mandibular development. *J Clin Invest* 123, 1638–1646. doi:10.1172/JCI66903.

Ramey, H. R., Decker, J. E., McKay, S. D., Rolf, M. M., Schnabel, R. D., and Taylor, J. F. (2013). Detection of selective sweeps in cattle using genome-wide SNP data. *BMC Genomics* 14, 382. doi:10.1186/1471-2164-14-382.

Ramos-García, P., González-Moles, M. Á., Ayén, Á., González-Ruiz, L., Ruiz-Ávila, I., and Gil-Montoya, J. A. (2019). Prognostic and clinicopathological significance of CTTN/cortactin alterations in head and neck squamous cell carcinoma: Systematic review and meta-analysis. *Head Neck* 41, 1963–1978. doi:10.1002/hed.25632.

Raphaka, K., Matika, O., Sánchez-Molano, E., Mrode, R., Coffey, M. P., Riggio, V., et al. (2017). Genomic regions underlying susceptibility to bovine tuberculosis in Holstein-Friesian cattle. *BMC Genet* 18. doi:10.1186/s12863-017-0493-7.

Raveendranathan, D., Babu, G. N., Desai, G., and Chandra, P. S. (2012). Bipolar Disorder Co-Occurring With Periodic Paralysis: A Case Report. *JNP* 24, E11–E12. doi:10.1176/appi.neuropsych.11010025.

Redecker, T. M., Kisko, T. M., Schwarting, R. K. W., and Wöhr, M. (2019). Effects of Cacna1c haploinsufficiency on social interaction behavior and 50-kHz ultrasonic vocalizations in adult female rats. *Behavioural Brain Research* 367, 35–52. doi:10.1016/j.bbr.2019.03.032.

Ren, J., Duan, Y., Qiao, R., Yao, F., Zhang, Z., Yang, B., et al. (2011). A Missense Mutation in PPARD Causes a Major QTL Effect on Ear Size in Pigs. *PLoS Genet* 7. doi:10.1371/journal.pgen.1002043.

Ren, Z.-Q., Yan, W.-J., Zhang, X.-Z., Zhang, P.-B., Zhang, C., and Chen, S.-K. (2019). CUL1 Knockdown Attenuates the Adhesion, Invasion, and Migration of Triple-Negative Breast Cancer Cells via Inhibition of Epithelial-Mesenchymal Transition. *Pathol. Oncol. Res.* doi:10.1007/s12253-019-00681-6.

Repnikova, E. A., Lyalin, D. A., McDonald, K., Astbury, C., Hansen-Kiss, E., Cooley, L. D., et al. (2020). CNTN6 copy number variations: Uncertain clinical significance in individuals with neurodevelopmental disorders. *European Journal of Medical Genetics* 63, 103636. doi:10.1016/j.ejmg.2019.02.008.

Reynolds, A. W., Mata-Míguez, J., Miró-Herrans, A., Briggs-Cloud, M., Sylestine, A., Barajas-Olmos, F., et al. (2019). Comparing signals of natural selection between three Indigenous North American populations. *PNAS* 116, 9312–9317. doi:10.1073/pnas.1819467116.

Ribasés, M., Ramos-Quiroga, J. A., Hervás, A., Bosch, R., Bielsa, A., Gastaminza, X., et al. (2009). Exploration of 19 serotoninergic candidate genes in adults and children with attention-deficit/hyperactivity disorder identifies association for 5HT2A, DDC and MAOB. *Mol. Psychiatry* 14, 71–85. doi:10.1038/sj.mp.4002100.

Rigterink, A., and Houpt, K. (2014). Genetics of canine behavior: A review. *World Journal of Medical Genetics* 4, 46–57. doi:10.5496/wjmg.v4.i3.46.

Riikonen, R. (2016). Treatment of autistic spectrum disorder with insulin-like growth factors. *Eur. J. Paediatr. Neurol.* 20, 816–823. doi:10.1016/j.ejpn.2016.08.005.

Ripke, S., Neale, B. M., Corvin, A., Walters, J. T., Farh, K.-H., Holmans, P. A., et al. (2014). Biological Insights From 108 Schizophrenia-Associated Genetic Loci. *Nature* 511, 421–427. doi:10.1038/nature13595.

Rodrigues, D. H., Rocha, N. P., Sousa, L. F. da C., Barbosa, I. G., Kummer, A., and Teixeira, A. L. (2014). Circulating levels of neurotrophic factors in autism spectrum disorders. *Neuro Endocrinol. Lett.* 35, 380–384.

Rodriguez-Revenga, L., Madrigal, I., Alkhalidi, L. S., Armengol, L., González, E., Badenas, C., et al. (2007). Contiguous deletion of the NDP, MAOA, MAOB, and EFHC2 genes in a patient with Norrie disease, severe psychomotor retardation and myoclonic epilepsy. *Am. J. Med. Genet. A* 143A, 916–920. doi:10.1002/ajmg.a.31521.

Rolstad, S., Pålsson, E., Ekman, C. J., Eriksson, E., Sellgren, C., and Landén, M. (2015). Polymorphisms of dopamine pathway genes NRG1 and LMX1A are associated with cognitive performance in bipolar disorder. *Bipolar Disorders* 17, 859–868. doi:10.1111/bdi.12347.

Roohi, J., Montagna, C., Tegay, D. H., Palmer, L. E., DeVincent, C., Pomeroy, J. C., et al. (2009). Disruption of contactin 4 in three subjects with autism spectrum disorder. *J Med Genet* 46, 176–182. doi:10.1136/jmg.2008.057505.

Roper, R. J., VanHorn, J. F., Cain, C. C., and Reeves, R. H. (2009). A neural crest deficit in Down syndrome mice is associated with deficient mitotic response to Sonic hedgehog. *Mech Dev* 126, 212. doi:10.1016/j.mod.2008.11.002.

Rosa, A. R., Frey, B. N., Andreazza, A. C., Ceresér, K. M., Cunha, A. B. M., Quevedo, J., et al. (2006). Increased serum glial cell line-derived neurotrophic factor immunocontent during manic and depressive episodes in individuals with bipolar disorder. *Neuroscience Letters* 407, 146–150. doi:10.1016/j.neulet.2006.08.026.

Ross, J., Gedvilaite, E., Badner, J. A., Erdman, C., Baird, L., Matsunami, N., et al. (2016). A Rare Variant in CACNA1D Segregates with 7 Bipolar I Disorder Cases in a Large Pedigree. *Mol Neuropsychiatry* 2, 145–150. doi:10.1159/000448041.

Roy, K., Murtie, J. C., El-Khodor, B. F., Edgar, N., Sardi, S. P., Hooks, B. M., et al. (2007). Loss of erbB signaling in oligodendrocytes alters myelin and dopaminergic function, a potential mechanism for neuropsychiatric disorders. *Proc Natl Acad Sci U S A* 104, 8131–8136. doi:10.1073/pnas.0702157104.

Royal, P., Andres-Bilbe, A., Ávalos Prado, P., Verkest, C., Wdziekonski, B., Schaub, S., et al. (2019). Migraine-Associated TRESK Mutations Increase Neuronal Excitability through Alternative Translation Initiation and Inhibition of TREK. *Neuron* 101, 232-245.e6. doi:10.1016/j.neuron.2018.11.039.

Ruggieri, V. L., and Arberas, C. L. (2003). [Behavioural phenotypes. Biologically determined neuropsychological patterns]. *Rev Neurol* 37, 239–253.

Ruiz, J. E., Barbosa Neto, J., Schoedl, A. F., and Mello, M. F. (2007). [Psychoneuroendocrinology of posttraumatic stress disorder]. *Braz J Psychiatry* 29 Suppl 1, S7-12. doi:10.1590/s1516-44462007000500003.

Rujescu, D., Giegling, I., Sato, T., Hartmann, A. M., and Möller, H. J. (2003). Genetic variations in tryptophan hydroxylase in suicidal behavior: analysis and meta-analysis. *Biol. Psychiatry* 54, 465–473. doi:10.1016/s0006-3223(02)01748-1.

Rujescu, D., Ingason, A., Cichon, S., Pietiläinen, O. P. H., Barnes, M. R., Toulopoulou, T., et al. (2009). Disruption of the neurexin 1 gene is associated with schizophrenia. *Hum. Mol. Genet.* 18, 988–996. doi:10.1093/hmg/ddn351.

Ruocco, A. C., Rodrigo, A. H., Carcone, D., McMain, S., Jacobs, G., and Kennedy, J. L. (2016). Tryptophan hydroxylase 1 gene polymorphisms alter prefrontal cortex activation during response inhibition. *Neuropsychology* 30, 18–27. doi:10.1037/neu0000237.

Ruzzo, E. K., Pérez-Cano, L., Jung, J.-Y., Wang, L.-K., Kashef-Haghighi, D., Hartl, C., et al. (2019). Inherited and De Novo Genetic Risk for Autism Impacts Shared Networks. *Cell* 178, 850-866.e26. doi:10.1016/j.cell.2019.07.015.

Ryan, M. C. M., Collins, P., and Thakore, J. H. (2003). Impaired fasting glucose tolerance in first-episode, drug-naive patients with schizophrenia. *Am J Psychiatry* 160, 284–289. doi:10.1176/appi.ajp.160.2.284.

Sacco, J., Ruplin, A., Skonieczny, P., and Ohman, M. (2017). Polymorphisms in the canine monoamine oxidase a (MAOA) gene: identification and variation among five broad dog breed groups. *Canine Genet Epidemiol* 4. doi:10.1186/s40575-016-0040-2.

Saetre, P., Lundmark, P., Wang, A., Hansen, T., Rasmussen, H. B., Djurovic, S., et al. (2010). The tryptophan hydroxylase 1 (TPH1) gene, schizophrenia susceptibility, and suicidal behavior: a multi-centre case-control study and meta-analysis. *Am. J. Med. Genet. B Neuropsychiatr. Genet.* 153B, 387–396. doi:10.1002/ajmg.b.30991.

Safari, R., Tunca, Z., Özerdem, A., Ceylan, D., Yalçın, Y., and Sakizli, M. (2017). Glial cell-derived neurotrophic factor gene polymorpisms affect severity and functionality of bipolar disorder. *J. Integr. Neurosci.* 16, 471–481. doi:10.3233/JIN-170031.

Saif, R., Tariq, B., and Naz, N. (2018). Sequence Diversity of MAOA Gene within Wild and Docile Animal Species. *Advancements in Life Sciences*. Available at: http://www.als-journal.com/538-18/ [Accessed January 28, 2020].

Saito, T., Aghalar, M. R., and Lachman, H. M. (2005). Analysis of PIK3C3 promoter variant in African-Americans with schizophrenia. *Schizophr. Res.* 76, 361–362. doi:10.1016/j.schres.2005.01.002.

Saito, T., Guan, F., Papolos, D. F., Rajouria, N., Fann, C. S., and Lachman, H. M. (2001). Polymorphism in SNAP29 gene promoter region associated with schizophrenia. *Mol. Psychiatry* 6, 193–201. doi:10.1038/sj.mp.4000825.

Salmela, E., Renvall, H., Kujala, J., Hakosalo, O., Illman, M., Vihla, M., et al. (2016). Evidence for genetic regulation of the human parieto-occipital 10-Hz rhythmic activity. *Eur. J. Neurosci.* 44, 1963–1971. doi:10.1111/ejn.13300.

Sánchez-Villagra, M. R., Geiger, M., and Schneider, R. A. (2016). The taming of the neural crest: a developmental perspective on the origins of morphological covariation in domesticated mammals. *R Soc Open Sci* 3. doi:10.1098/rsos.160107.

Sanders, S. J., Ercan-Sencicek, A. G., Hus, V., Luo, R., Murtha, M. T., Moreno-De-Luca, D., et al. (2011). Multiple Recurrent De Novo CNVs, Including Duplications of the 7q11.23 Williams Syndrome Region, Are Strongly Associated with Autism. *Neuron* 70, 863–885. doi:10.1016/j.neuron.2011.05.002.

Sandman, C. A., Hetrick, W., Talyor, D., Marion, S., and Chicz-DeMet, A. (2000). Uncoupling of proopiomelanocortin (POMC) fragments is related to self-injury. *Peptides* 21, 785–791. doi:10.1016/s0196-9781(00)00209-6.

Saura, J., Luque, J. M., Cesura, A. M., Da Prada, M., Chan-Palay, V., Huber, G., et al. (1994). Increased monoamine oxidase B activity in plaque-associated astrocytes of Alzheimer brains revealed by quantitative enzyme radioautography. *Neuroscience* 62, 15–30. doi:10.1016/0306-4522(94)90311-5.

Sbacchi, S., Acquadro, F., Calò, I., Calì, F., and Romano, V. (2010). Functional Annotation of Genes Overlapping Copy Number Variants in Autistic Patients: Focus on Axon Pathfinding. *Curr Genomics* 11, 136–145. doi:10.2174/138920210790886880.

Schaaf, C. P., Boone, P. M., Sampath, S., Williams, C., Bader, P. I., Mueller, J. M., et al. (2012). Phenotypic spectrum and genotype-phenotype correlations of NRXN1 exon deletions. *Eur. J. Hum. Genet.* 20, 1240–1247. doi:10.1038/ejhg.2012.95.

Schechter, D. S., Moser, D. A., Paoloni-Giacobino, A., Stenz, L., Gex-Fabry, M., Aue, T., et al. (2015). Methylation of NR3C1 is related to maternal PTSD, parenting stress and maternal medial prefrontal cortical activity in response to child separation among mothers with histories of violence exposure. *Front Psychol* 6. doi:10.3389/fpsyg.2015.00690.

Schiffer, H. H., and Heinemann, S. F. (2007). Association of the human kainate receptor GluR7 gene (GRIK3) with recurrent major depressive disorder. *American Journal of Medical Genetics Part B: Neuropsychiatric Genetics* 144B, 20–26. doi:10.1002/ajmg.b.30374.

Schizophrenia Psychiatric Genome-Wide Association Study (GWAS) Consortium (2011). Genome-wide association study identifies five new schizophrenia loci. *Nat. Genet.* 43, 969–976. doi:10.1038/ng.940.

Schriemer, D., Sribudiani, Y., IJpma, A., Natarajan, D., MacKenzie, K. C., Metzger, M., et al. (2016). Regulators of gene expression in Enteric Neural Crest Cells are putative Hirschsprung disease genes. *Developmental Biology* 416, 255–265. doi:10.1016/j.ydbio.2016.06.004.

Schubert, M., Jónsson, H., Chang, D., Sarkissian, C. D., Ermini, L., Ginolhac, A., et al. (2014). Prehistoric genomes reveal the genetic foundation and cost of horse domestication. *PNAS* 111, E5661–E5669. doi:10.1073/pnas.1416991111.

Scoles, D. R., Meera, P., Schneider, M., Paul, S., Dansithong, W., Figueroa, K. P., et al. (2017). Antisense oligonucleotide therapy for spinocerebellar ataxia type 2. *Nature* 544, 362–366. doi:10.1038/nature22044.

Secolin, R., Banzato, C. E. M., Mella, L. F. B., Santos, M. L., Dalgalarrondo, P., and Lopes-Cendes, I. (2013). Refinement of chromosome 3p22.3 region and identification of a susceptibility gene for bipolar affective disorder. *American Journal of Medical Genetics Part B: Neuropsychiatric Genetics* 162, 163–168. doi:10.1002/ajmg.b.32127.

Segurado, R., Detera-Wadleigh, S. D., Levinson, D. F., Lewis, C. M., Gill, M., Nurnberger, Jr., J. I., et al. (2003). Genome Scan Meta-Analysis of Schizophrenia and Bipolar Disorder, Part III: Bipolar Disorder. *Am J Hum Genet* 73, 49–62.

Sehmbi, M., Rowley, C. D., Minuzzi, L., Kapczinski, F., Steiner, M., Sassi, R. B., et al. (2018). Association of intracortical myelin and cognitive function in bipolar I disorder. *Acta Psychiatr Scand* 138, 62–72. doi:10.1111/acps.12875.

Shah, B., and Tobias, J. D. (2006). Osmotic demyelination and hypertonic dehydration in a 9-year-old girl: changes in cerebrospinal fluid myelin basic protein. *J Intensive Care Med* 21, 372–376. doi:10.1177/0885066606293358.

Shah, K., Mann, I., Reddy, K., and John, G. A Case of Severe Psychosis Due to Cushing's

Syndrome Secondary to Primary Bilateral Macronodular Adrenal Hyperplasia. *Cureus* 11. doi:10.7759/cureus.6162.

Shaltiel, G., Maeng, S., Malkesman, O., Pearson, B., Schloesser, R. J., Tragon, T., et al. (2008). Evidence for the involvement of the kainate receptor subunit GluR6 (GRIK2) in mediating behavioral displays related to behavioral symptoms of mania. *Mol. Psychiatry* 13, 858–872. doi:10.1038/mp.2008.20.

Shao, L., and Vawter, M. P. (2008). Shared gene expression alterations in schizophrenia and bipolar disorder. *Biol. Psychiatry* 64, 89–97. doi:10.1016/j.biopsych.2007.11.010.

Sheth, K., Moss, J., Hyland, S., Stinton, C., Cole, T., and Oliver, C. (2015). The behavioral characteristics of Sotos syndrome. *Am. J. Med. Genet. A* 167A, 2945–2956. doi:10.1002/ajmg.a.37373.

Shi, L., Chen, S.-J., Deng, J.-H., Que, J.-Y., Lin, X., Sun, Y., et al. (2018). ADRB2 gene polymorphism modulates the retention of fear extinction memory. *Neurobiology of Learning and Memory* 156, 96–102. doi:10.1016/j.nlm.2018.11.004.

Shiah, I.-S., and Yatham, L. N. (2000). Serotonin in mania and in the mechanism of action of mood stabilizers: a review of clinical studies. *Bipolar Disorders* 2, 77–92. doi:10.1034/j.1399-5618.2000.020201.x.

Shibata, H., Shibata, A., Ninomiya, H., Tashiro, N., and Fukumaki, Y. (2002). Association study of polymorphisms in the GluR6 kainate receptor gene (GRIK2) with schizophrenia. *Psychiatry Res* 113, 59–67. doi:10.1016/s0165-1781(02)00231-7.

Shih, J. C., and Thompson, R. F. (1999). Monoamine oxidase in neuropsychiatry and behavior. *Am J Hum Genet* 65, 593–598.

Shikhevich, S. G., Os'kina, I. N., and Pliusnina, I. Z. (2002). [Effect of stress and immune stimulus on the pituitary-adrenal axis in gray rats selected for behavior]. *Ross Fiziol Zh Im I M Sechenova* 88, 781–789.

Shuang, M., Liu, J., Jia, M. X., Yang, J. Z., Wu, S. P., Gong, X. H., et al. (2004). Family-based association study between autism and glutamate receptor 6 gene in Chinese Han trios. *Am. J. Med. Genet. B Neuropsychiatr. Genet.* 131B, 48–50. doi:10.1002/ajmg.b.30025.

Sieghart, W., and Sperk, G. (2002). Subunit composition, distribution and function of GABA(A) receptor subtypes. *Curr Top Med Chem* 2, 795–816. doi:10.2174/1568026023393507.

Silberberg, G., Darvasi, A., Pinkas-Kramarski, R., and Navon, R. (2006). The involvement of ErbB4 with schizophrenia: association and expression studies. *Am. J. Med. Genet. B Neuropsychiatr. Genet.* 141B, 142–148. doi:10.1002/ajmg.b.30275.

Simmons, A. B., Bloomsburg, S. J., Sukeena, J. M., Miller, C. J., Ortega-Burgos, Y., Borghuis, B. G., et al. (2017). DSCAM-mediated control of dendritic and axonal arbor outgrowth enforces tiling and inhibits synaptic plasticity. *Proc. Natl. Acad. Sci. U.S.A.* 114, E10224–E10233. doi:10.1073/pnas.1713548114.

Singh, V. K., Warren, R. P., Odell, J. D., Warren, W. L., and Cole, P. (1993). Antibodies to myelin basic protein in children with autistic behavior. *Brain Behav. Immun.* 7, 97–103. doi:10.1006/brbi.1993.1010.

Sklar, P., Ripke, S., Scott, L. J., Andreassen, O. A., Cichon, S., Craddock, N., et al. (2011). Large-scale genome-wide association analysis of bipolar disorder identifies a new susceptibility locus near ODZ4. *Nat Genet* 43, 977–983. doi:10.1038/ng.943.

Smedler, E., Pålsson, E., Hashimoto, K., and Landén, M. (2019). Association of CACNA1C polymorphisms with serum BDNF levels in bipolar disorder. *Br J Psychiatry*, 1–3. doi:10.1192/bjp.2019.173.

Smith, A. K., Dimulescu, I., Falkenberg, V. R., Narasimhan, S., Heim, C., Vernon, S. D., et al. (2008). Genetic evaluation of the serotonergic system in chronic fatigue syndrome. *Psychoneuroendocrinology*. 33(2), 188-197. 10.1016/j.psyneuen.2007.11.001.

Smith, S. E., Mullen, T. E., Graham, D., Sims, K. B., and Rehm, H. L. (2012). Norrie disease: Extraocular clinical manifestations in 56 patients. *American Journal of Medical Genetics Part A* 158A, 1909–1917. doi:10.1002/ajmg.a.35469.

Smyth, C., Kalsi, G., Brynjolfsson, J., O'Neill, J., Curtis, D., Rifkin, L., et al. (1996). Further tests for linkage of bipolar affective disorder to the tyrosine hydroxylase gene locus on chromosome 11p15 in a new series of multiplex British affective disorder pedigrees. *Am J Psychiatry* 153, 271–274. doi:10.1176/ajp.153.2.271.

Soeiro-de-Souza, M. G., Bio, D. S., Dias, V. V., Vieta, E., Machado-Vieira, R., and Moreno, R. A. (2013). The CACNA1C risk allele selectively impacts on executive function in bipolar type I disorder. *Acta Psychiatrica Scandinavica* 128, 362–369. doi:10.1111/acps.12073.

Southam, E., Kirkby, D., Higgins, G. A., and Hagan, R. M. (1998). Lamotrigine inhibits monoamine uptake in vitro and modulates 5-hydroxytryptamine uptake in rats. *European Journal of Pharmacology* 358, 19–24. doi:10.1016/S0014-2999(98)00580-9.

Spiteri, B. S., Stafrace, Y., and Calleja-Agius, J. (2017). Silver-Russell Syndrome: A Review. *Neonatal Netw* 36, 206–212. doi:10.1891/0730-0832.36.4.206.

Spitzer, S., Volbracht, K., Lundgaard, I., and Káradóttir, R. T. (2016). Glutamate signalling: A multifaceted modulator of oligodendrocyte lineage cells in health and disease. *Neuropharmacology* 110, 574–585. doi:10.1016/j.neuropharm.2016.06.014.

Splawski, I., Timothy, K. W., Sharpe, L. M., Decher, N., Kumar, P., Bloise, R., et al. (2004). CaV1.2 Calcium Channel Dysfunction Causes a Multisystem Disorder Including Arrhythmia and Autism. *Cell* 119, 19–31. doi:10.1016/j.cell.2004.09.011.

Squassina, A., Costa, M., Congiu, D., Manchia, M., Angius, A., Deiana, V., et al. (2013). Insulin-like growth factor 1 (IGF-1) expression is up-regulated in lymphoblastoid cell lines of lithium responsive bipolar disorder patients. *Pharmacol. Res.* 73, 1–7. doi:10.1016/j.phrs.2013.04.004.

Srikanth, K., Kim, N.-Y., Park, W., Kim, J.-M., Kim, K.-D., Lee, K.-T., et al. (2019). Comprehensive genome and transcriptome analyses reveal genetic relationship, selection signature, and transcriptome landscape of small-sized Korean native Jeju horse. *Sci Rep* 9. doi:10.1038/s41598-019-53102-8.

Starnawska, A., Demontis, D., Pen, A., Hedemand, A., Nielsen, A. L., Staunstrup, N. H., et al. (2016). CACNA1C hypermethylation is associated with bipolar disorder. *Translational Psychiatry* 6, e831–e831. doi:10.1038/tp.2016.99.

Stassart, R. M., Möbius, W., Nave, K.-A., and Edgar, J. M. (2018). The Axon-Myelin Unit in Development and Degenerative Disease. *Front Neurosci* 12. doi:10.3389/fnins.2018.00467.

Steinman, G., and Mankuta, D. (2019). Molecular biology of autism's etiology – An alternative mechanism. *Medical Hypotheses* 130, 109272. doi:10.1016/j.mehy.2019.109272.

Stelzhammer, V., Alsaif, M., Chan, M. K., Rahmoune, H., Steeb, H., Guest, P. C., et al. (2015). Distinct proteomic profiles in post-mortem pituitary glands from bipolar disorder and major depressive disorder patients. Journal of Psychiatric Research 60, 40–48. doi:10.1016/j.jpsychires.2014.09.022.

Stessman, H. A. F., Xiong, B., Coe, B. P., Wang, T., Hoekzema, K., Fenckova, M., et al. (2017). Targeted sequencing identifies 91 neurodevelopmental disorder risk genes with autism and developmental disability biases. *Nat Genet* 49, 515–526. doi:10.1038/ng.3792.

Stopkova, P., Saito, T., Papolos, D. F., Vevera, J., Paclt, I., Zukov, I., et al. (2004). Identification of PIK3C3 promoter variant associated with bipolar disorder and schizophrenia. *Biol. Psychiatry* 55, 981–988. doi:10.1016/j.biopsych.2004.01.014.

Su, H.-Y., Lin, Z.-Y., Peng, W.-C., Guan, F., Zhu, G.-T., Mao, B.-B., et al. (2018). MiR-448 downregulates CTTN to inhibit cell proliferation and promote apoptosis in glioma. *Eur Rev Med Pharmacol Sci* 22, 3847–3854. doi:10.26355/eurrev_201806_15269.

Su, Q., Mochida, S., Tian, J.-H., Mehta, R., and Sheng, Z.-H. (2001). SNAP-29: A general SNARE protein that inhibits SNARE disassembly and is implicated in synaptic transmission. *Proc Natl Acad Sci U S A* 98, 14038–14043. doi:10.1073/pnas.251532398.

Suárez-Merino, B., Bye, J., McDowall, J., Ross, M., and Craig, I. W. (2001). Sequence analysis and transcript identification within 1.5 MB of DNA deleted together with the NDP and MAO genes in atypical Norrie disease patients presenting with a profound phenotype. *Hum. Mutat.* 17, 523. doi:10.1002/humu.1140.

Sun, J., Kuo, P.-H., Riley, B. P., Kendler, K. S., and Zhao, Z. (2008a). Candidate genes for schizophrenia: a survey of association studies and gene ranking. *Am. J. Med. Genet. B Neuropsychiatr. Genet.* 147B, 1173–1181. doi:10.1002/ajmg.b.30743.

Sun, S., Liu, Y., Wei, J., Liu, S., and Ju, G. (2008b). The PPARD gene may be associated with schizophrenia in a Chinese population. *Psychiatric Genetics* 18, 253–254. doi:10.1097/YPG.0b013e3283053035.

Sutter, N. B., Bustamante, C. D., Chase, K., Gray, M. M., Zhao, K., Zhu, L., et al. (2007). A single IGF1 allele is a major determinant of small size in dogs. *Science* 316, 112–115. doi:10.1126/science.1137045.

Suzuki, T., Iwata, N., Kitamura, Y., Kitajima, T., Yamanouchi, Y., Ikeda, M., et al. (2003). Association of a haplotype in the serotonin 5-HT4 receptor gene (HTR4) with Japanese schizophrenia. *Am. J. Med. Genet. B Neuropsychiatr. Genet.* 121B, 7–13. doi:10.1002/ajmg.b.20060.

Switonski, M., Mankowska, M., and Salamon, S. (2013). Family of melanocortin receptor (MCR) genes in mammals—mutations, polymorphisms and phenotypic effects. *J Appl Genet* 54, 461–472. doi:10.1007/s13353-013-0163-z.

Sykes, L., Clifton, N. E., Hall, J., and Thomas, K. L. (2018). Regulation of the Expression of the Psychiatric Risk Gene Cacna1c during Associative Learning. *MNP* 4, 149–157. doi:10.1159/000493917.

Sykes, L., Haddon, J., Lancaster, T. M., Sykes, A., Azzouni, K., Ihssen, N., et al. (2019). Genetic Variation in the Psychiatric Risk Gene CACNA1C Modulates Reversal Learning Across Species. *Schizophr Bull* 45, 1024–1032. doi:10.1093/schbul/sby146.

Szumska, D., Pieles, G., Essalmani, R., Bilski, M., Mesnard, D., Kaur, K., et al. (2008). VACTERL/caudal regression/Currarino syndrome-like malformations in mice with mutation in the proprotein convertase Pcsk5. *Genes Dev* 22, 1465–1477. doi:10.1101/gad.479408.

Tajdaran, K., Gordon, T., Wood, M. D., Shoichet, M. S., and Borschel, G. H. (2016). A glial cell line-derived neurotrophic factor delivery system enhances nerve regeneration across acellular nerve allografts. *Acta Biomater* 29, 62–70. doi:10.1016/j.actbio.2015.10.001.

Takahashi, K., Ishida, M., and Takahashi, H. (2009). Expression of Sema3D in subsets of neurons in the developing dorsal root ganglia of the rat. *Neurosci. Lett.* 455, 17–21. doi:10.1016/j.neulet.2009.03.050.

Takahashi, N., Sakurai, T., Davis, K. L., and Buxbaum, J. D. (2011). Linking oligodendrocyte and myelin dysfunction to neurocircuitry abnormalities in schizophrenia. *Progress in Neurobiology* 93, 13–24. doi:10.1016/j.pneurobio.2010.09.004.

Takahashi, Y., Fukuda, Y., Yoshimura, J., Toyoda, A., Kurppa, K., Moritoyo, H., et al. (2013). ERBB4 mutations that disrupt the neuregulin-ErbB4 pathway cause amyotrophic lateral sclerosis type 19. *Am. J. Hum. Genet.* 93, 900–905. doi:10.1016/j.ajhg.2013.09.008.

Takata, A., Matsumoto, N., and Kato, T. (2017). Genome-wide identification of splicing QTLs in the human brain and their enrichment among schizophrenia-associated loci. *Nat Commun* 8. doi:10.1038/ncomms14519.

Takeuchi, Y., Hashizume, C., Chon, E. M. H., Momozawa, Y., Masuda, K., Kikusui, T., et al. (2005). Canine tyrosine hydroxylase (TH) gene and dopamine beta -hydroxylase (DBH) gene: their sequences, genetic polymorphisms, and diversities among five different dog breeds. *J. Vet. Med. Sci.* 67, 861–867. doi:10.1292/jvms.67.861.

Talley, E. M., Solórzano, G., Lei, Q., Kim, D., and Bayliss, D. A. (2001). CNS Distribution of Members of the Two-Pore-Domain (KCNK) Potassium Channel Family. *J Neurosci* 21, 7491–7505. doi:10.1523/JNEUROSCI.21-19-07491.2001.

Tang, R., Zhao, X., Fang, C., Tang, W., Huang, K., Wang, L., et al. (2008). Investigation of variants in the promoter region of PIK3C3 in schizophrenia. *Neurosci. Lett.* 437, 42–44. doi:10.1016/j.neulet.2008.03.043.

Tansey, K. E., Rucker, J. J. H., Kavanagh, D. H., Guipponi, M., Perroud, N., Bondolfi, G., et al. (2014). Copy number variants and therapeutic response to antidepressant medication in major depressive disorder. *The Pharmacogenomics Journal* 14, 395–399. doi:10.1038/tpj.2013.51.

Tassano, E., Severino, M., Rosina, S., Papa, R., Tortora, D., Gimelli, G., et al. (2016). Interstitial de novo 18q22.3q23 deletion: clinical, neuroradiological and molecular characterization of a new case and review of the literature. *Mol Cytogenet* 9. doi:10.1186/s13039-016-0285-1.

Tatton-Brown, K., Pilz, D. T., Örstavik, K. H., Patton, M., Barber, J. C. K., Collinson, M. N., et al. (2009). 15q overgrowth syndrome: A newly recognized phenotype associated with overgrowth, learning difficulties, characteristic facial appearance, renal anomalies and increased dosage of distal chromosome 15q. *American Journal of Medical Genetics Part A* 149A, 147–154. doi:10.1002/ajmg.a.32534.

Tejedor-Real, P., Biguet, N. F., Dumas, S., and Mallet, J. (2003). Tyrosine hydroxylase mRNA and protein are down-regulated by chronic clozapine in both the mesocorticolimbic and the nigrostriatal systems. *Journal of Neuroscience Research* 72, 105–115. doi:10.1002/jnr.10555.

Teletchea, F. (2019). Animal Domestication: A Brief Overview. *Animal Domestication*. doi:10.5772/intechopen.86783.

Tesli, M., Skatun, K. C., Ousdal, O. T., Brown, A. A., Thoresen, C., Agartz, I., et al. (2013). CACNA1C Risk Variant and Amygdala Activity in Bipolar Disorder, Schizophrenia and Healthy Controls. *PLoS One* 8. doi:10.1371/journal.pone.0056970.

Theofanopoulou, C., Gastaldon, S., O'Rourke, T., Samuels, B. D., Messner, A., Martins, P. T., et al. (2017). Self-domestication in Homo sapiens: Insights from comparative genomics. *PLOS ONE* 12, e0185306. doi:10.1371/journal.pone.0185306.

Thomas, R. H., Chung, S.-K., Wood, S. E., Cushion, T. D., Drew, C. J. G., Hammond, C. L., et al. (2013). Genotype-phenotype correlations in hyperekplexia: apnoeas, learning difficulties and speech delay. *Brain* 136, 3085–3095. doi:10.1093/brain/awt207.

Thümmler, S., Duprat, F., and Lazdunski, M. (2007). Antipsychotics inhibit TREK but not TRAAK channels. *Biochem. Biophys. Res. Commun.* 354, 284–289. doi:10.1016/j.bbrc.2006.12.199.

Tian, J., Geng, F., Gao, F., Chen, Y.-H., Liu, J.-H., Wu, J.-L., et al. (2017). Down-Regulation of Neuregulin1/ErbB4 Signaling in the Hippocampus Is Critical for Learning and Memory. *Mol. Neurobiol.* 54, 3976–3987. doi:10.1007/s12035-016-9956-5.

Tiihonen, J., Rautiainen, M.-R., Ollila, H., Repo-Tiihonen, E., Virkkunen, M., Palotie, A., et al. (2015). Genetic background of extreme violent behavior. *Mol Psychiatry* 20, 786–792. doi:10.1038/mp.2014.130.

Tilot, A. K., Frazier, T. W., and Eng, C. (2015). Balancing Proliferation and Connectivity in PTEN-associated Autism Spectrum Disorder. *Neurotherapeutics* 12, 609–619. doi:10.1007/s13311-015-0356-8.

Tiwari, V., O'Donnell, C. D., Oh, M.-J., Valyi-Nagy, T., and Shukla, D. (2005). A role for 3-O-sulfotransferase isoform-4 in assisting HSV-1 entry and spread. *Biochem. Biophys. Res. Commun.* 338, 930–937. doi:10.1016/j.bbrc.2005.10.056.

Tkachev, D., Mimmack, M. L., Ryan, M. M., Wayland, M., Freeman, T., Jones, P. B., et al. (2003). Oligodendrocyte dysfunction in schizophrenia and bipolar disorder. *Lancet* 362, 798–805. doi:10.1016/S0140-6736(03)14289-4.

Todarello, G., Feng, N., Kolachana, B. S., Li, C., Vakkalanka, R., Bertolino, A., et al. (2014). Incomplete penetrance of NRXN1 deletions in families with schizophrenia. *Schizophr. Res.* 155, 1–7. doi:10.1016/j.schres.2014.02.023.

Toma, C., Hervás, A., Balmaña, N., Salgado, M., Maristany, M., Vilella, E., et al. (2013). Neurotransmitter systems and neurotrophic factors in autism: association study of 37 genes suggests involvement of DDC. *World J. Biol. Psychiatry* 14, 516–527. doi:10.3109/15622975.2011.602719.

Torres-Berrío, A., Lopez, J. P., Bagot, R. C., Nouel, D., Dal Bo, G., Cuesta, S., et al. (2017). DCC Confers Susceptibility to Depression-like Behaviors in Humans and Mice and Is Regulated by miR-218. *Biol. Psychiatry* 81, 306–315. doi:10.1016/j.biopsych.2016.08.017.

Toth, M. (2019). The other side of the coin: hypersociability. *Genes Brain Behav* 18, e12512. doi:10.1111/gbb.12512.

Trut, L., Oskina, I., and Kharlamova, A. (2009). Animal evolution during domestication: the domesticated fox as a model. *BioEssays* 31, 349–360. doi:10.1002/bies.200800070.

Trut, L. N., Pliusnina, I. Z., and Os'kina, I. N. (2004). [An experiment on fox domestication and debatable issues of evolution of the dog]. *Genetika* 40, 794–807.

Tsilioni, I., Dodman, N., Petra, A. I., Taliou, A., Francis, K., Moon-Fanelli, A., et al. (2014). Elevated serum neurotensin and CRH levels in children with autistic spectrum disorders and tail-chasing Bull Terriers with a phenotype similar to autism. *Transl Psychiatry* 4, e466. doi:10.1038/tp.2014.106.

Tsuda, T., Iwai, N., Deguchi, E., Kimura, O., Ono, S., Furukawa, T., et al. (2011). PCSK5 and GDF11 expression in the hindgut region of mouse embryos with anorectal malformations. *Eur J Pediatr Surg* 21, 238–241. doi:10.1055/s-0031-1273691.

Tunca, Z., Kıvırcık Akdede, B., Özerdem, A., Alkın, T., Polat, S., Ceylan, D., et al. (2015). Diverse glial cell line-derived neurotrophic factor (GDNF) support between mania and schizophrenia: a comparative study in four major psychiatric disorders. *Eur. Psychiatry* 30, 198–204. doi:10.1016/j.eurpsy.2014.11.003.

Turner, T. N., Hormozdiari, F., Duyzend, M. H., McClymont, S. A., Hook, P. W., Iossifov, I., et al. (2016). Genome Sequencing of Autism-Affected Families Reveals Disruption of Putative Noncoding Regulatory DNA. *Am. J. Hum. Genet.* 98, 58–74. doi:10.1016/j.ajhg.2015.11.023.

Tymchuk, W. E., Beckman, B., and Devlin, R. H. (2009). Altered Expression of Growth Hormone/Insulin-Like Growth Factor I Axis Hormones in Domesticated Fish. *Endocrinology* 150, 1809–1816. doi:10.1210/en.2008-0797.

Ueda, S., Negishi, M., and Katoh, H. (2013). Rac GEF Dock4 interacts with cortactin to regulate dendritic spine formation. *Mol Biol Cell* 24, 1602–1613. doi:10.1091/mbc.E12-11-0782.

Uliana, V., Grosso, S., Cioni, M., Ariani, F., Papa, F. T., Tamburello, S., et al. (2010). 3.2 Mb microdeletion in chromosome 7 bands q22.2–q22.3 associated with overgrowth and delayed bone age. *European Journal of Medical Genetics* 53, 168–170. doi:10.1016/j.ejmg.2010.02.003.

van Daalen, E., Kemner, C., Verbeek, N. E., van der Zwaag, B., Dijkhuizen, T., Rump, P., et al. (2011). Social responsiveness scale-aided analysis of the clinical impact of copy number variations in autism. *Neurogenetics* 12, 315–323. doi:10.1007/s10048-011-0297-2.

Vandael, D. H. F., Marcantoni, A., and Carbone, E. (2015). Cav1.3 Channels as Key Regulators of Neuron-Like Firings and Catecholamine Release in Chromaffin Cells. *Curr Mol Pharmacol* 8, 149–161. doi:10.2174/1874467208666150507105443.

Varnäs, K., Halldin, C., Pike, V. W., and Hall, H. (2003). Distribution of 5-HT4 receptors in the postmortem human brain--an autoradiographic study using [125I]SB 207710. *Eur Neuropsychopharmacol* 13, 228–234. doi:10.1016/s0924-977x(03)00009-9.

Venkatasubramanian, G., Chittiprol, S., Neelakantachar, N., Naveen, M. N., Thirthall, J., Gangadhar, B. N., et al. (2007). Insulin and insulin-like growth factor-1 abnormalities in antipsychotic-naive schizophrenia. *Am J Psychiatry* 164, 1557–1560. doi:10.1176/appi.ajp.2007.07020233.

Viikki, M., Kampman, O., Illi, A., Setälä-Soikkeli, E., Anttila, S., Huuhka, M., et al. (2010). TPH1 218A/C polymorphism is associated with major depressive disorder and its treatment response. *Neurosci. Lett.* 468, 80–84. doi:10.1016/j.neulet.2009.10.069.

Viñas-Jornet, M., Esteba-Castillo, S., Gabau, E., Ribas-Vidal, N., Baena, N., San, J., et al. (2014). A common cognitive, psychiatric, and dysmorphic phenotype in carriers of NRXN1 deletion. *Mol Genet Genomic Med* 2, 512–521. doi:10.1002/mgg3.105.

Vincent, J. B., Noor, A., Windpassinger, C., Gianakopoulos, P. J., Schwarzbraun, T., Alfred, S. E., et al. (2009). Characterization of a de novo translocation t(5;18)(q33.1;q12.1) in an autistic boy identifies a breakpoint close to SH3TC2, ADRB2, and HTR4 on 5q, and within the desmocollin gene cluster on 18q. *Am. J. Med. Genet. B Neuropsychiatr. Genet.* 150B, 817–826. doi:10.1002/ajmg.b.30903.

Visser, R., Gijsbers, A., Ruivenkamp, C., Karperien, M., Reeser, H. M., Breuning, M. H., et al. (2010). Genome-Wide SNP Array Analysis in Patients with Features of Sotos Syndrome. *HRP* 73, 265–274. doi:10.1159/000284391.

Vistein, R., and Puthenveedu, M. A. (2014). Src regulates sequence-dependent beta-2 adrenergic receptor recycling via cortactin phosphorylation. *Traffic* 15, 1195–1205. doi:10.1111/tra.12202.

Vitalis, T., Alvarez, C., Chen, K., Shih, J. C., Gaspar, P., and Cases, O. (2003). Developmental expression pattern of monoamine oxidases in sensory organs and neural crest derivatives. *J. Comp. Neurol.* 464, 392–403. doi:10.1002/cne.10804.

Vjugina, U., Zhu, X., Oh, E., Bracero, N. J., and Evans, J. P. (2009). Reduction of Mouse Egg Surface Integrin Alpha9 Subunit (ITGA9) Reduces the Egg's Ability to Support Sperm-Egg Binding and Fusion. *Biol Reprod* 80, 833–841. doi:10.1095/biolreprod.108.075275.

Voineskos, A. N., Lett, T. A. P., Lerch, J. P., Tiwari, A. K., Ameis, S. H., Rajji, T. K., et al. (2011). Neurexin-1 and frontal lobe white matter: an overlapping intermediate phenotype for schizophrenia and autism spectrum disorders. *PLoS ONE* 6, e20982. doi:10.1371/journal.pone.0020982.

Völkening, B., Schönig, K., Kronenberg, G., Bartsch, D., and Weber, T. (2017). Deletion of psychiatric risk gene Cacna1c impairs hippocampal neurogenesis in cell-autonomous fashion. *Glia* 65, 817–827. doi:10.1002/glia.23128.

Voltas, N., Aparicio, E., Arija, V., and Canals, J. (2015). Association study of monoamine oxidase-A gene promoter polymorphism (MAOA-uVNTR) with self-reported anxiety and other psychopathological symptoms in a community sample of early adolescents. *J Anxiety Disord* 31, 65–72. doi:10.1016/j.janxdis.2015.02.004.

vonHoldt, B. M., Shuldiner, E., Koch, I. J., Kartzinel, R. Y., Hogan, A., Brubaker, L., et al. (2017). Structural variants in genes associated with human Williams-Beuren syndrome underlie stereotypical hypersociability in domestic dogs. *Science Advances* 3, e1700398. doi:10.1126/sciadv.1700398.

Vora, A. K., Fisher, A. M., New, A. S., Hazlett, E. A., McNamara, M., Yuan, Q., et al. (2018). Dimensional Traits Of Schizotypy Associated With Glycine receptor GLRA1 Polymorphism: An Exploratory Candidate-Gene Association Study. *J Pers Disord* 32, 421–432. doi:10.1521/pedi_2017_31_303.

Vosberg, D. E., Beaulé, V., Torres-Berrío, A., Cooke, D., Chalupa, A., Jaworska, N., et al. (2019). Neural function in DCC mutation carriers with and without mirror movements. *Ann. Neurol.* 85, 433–442. doi:10.1002/ana.25418.

Vosberg, D. E., Leyton, M., and Flores, C. (2020). The Netrin-1/DCC guidance system: dopamine pathway maturation and psychiatric disorders emerging in adolescence. *Mol. Psychiatry* 25, 297–307. doi:10.1038/s41380-019-0561-7.

Vrijenhoek, T., Buizer-Voskamp, J. E., van der Stelt, I., Strengman, E., Genetic Risk and Outcome in Psychosis (GROUP) Consortium, Sabatti, C., et al. (2008). Recurrent CNVs disrupt three candidate genes in schizophrenia patients. *Am. J. Hum. Genet.* 83, 504–510. doi:10.1016/j.ajhg.2008.09.011.

Walenski, M., Mostofsky, S. H., and Ullman, M. T. (2007). Speeded processing of grammar and tool knowledge in Tourette's syndrome. *Neuropsychologia* 45, 2447–2460. doi:10.1016/j.neuropsychologia.2007.04.001.

Walker, S., and Scherer, S. W. (2013). Identification of candidate intergenic risk loci in autism spectrum disorder. *BMC Genomics* 14, 499. doi:10.1186/1471-2164-14-499.

Walther, D. J., and Bader, M. (2003). A unique central tryptophan hydroxylase isoform. *Biochem. Pharmacol.* 66, 1673–1680. doi:10.1016/s0006-2952(03)00556-2.

Wang, D., Weng, Y., Guo, S., Qin, W., Ni, J., Yu, L., et al. (2019). microRNA-1 Regulates NCC Migration and Differentiation by Targeting sec63. *Int J Biol Sci* 15, 2538–2547. doi:10.7150/ijbs.35357.

Wang, D.-S., Zurek, A. A., Lecker, I., Yu, J., Abramian, A. M., Avramescu, S., et al. (2012). Memory Deficits Induced by Inflammation Are Regulated by α5-Subunit-Containing GABAA Receptors. *Cell Rep* 2, 488–496. doi:10.1016/j.celrep.2012.08.022.

Wang, K.-S., Liu, X.-F., and Aragam, N. (2010). A genome-wide meta-analysis identifies novel loci associated with schizophrenia and bipolar disorder. *Schizophr. Res.* 124, 192–199. doi:10.1016/j.schres.2010.09.002.

Wang, K.-S., Tonarelli, S., Luo, X., Wang, L., Su, B., Zuo, L., et al. (2015a). Polymorphisms within ASTN2 gene are associated with age at onset of Alzheimer's disease. *J Neural Transm (Vienna)* 122, 701–708. doi:10.1007/s00702-014-1306-z.

Wang, L., Budolfson, K., and Wang, F. (2011). Pik3c3 deletion in pyramidal neurons results in loss of synapses, extensive gliosis and progressive neurodegeneration. *Neuroscience* 172, 427–442. doi:10.1016/j.neuroscience.2010.10.035.

Wang, R., Chen, C.-C., Hara, E., Rivas, M. V., Roulhac, P. L., Howard, J. T., et al. (2015b). Convergent differential regulation of SLIT-ROBO axon guidance genes in the brains of vocal learners. *J. Comp. Neurol.* 523, 892–906. doi:10.1002/cne.23719.

Wang, T., Guo, H., Xiong, B., Stessman, H. A. F., Wu, H., Coe, B. P., et al. (2016). De novo genic mutations among a Chinese autism spectrum disorder cohort. *Nat Commun* 7, 13316. doi:10.1038/ncomms13316.

Wang, X., Pipes, L., Trut, L. N., Herbeck, Y., Vladimirova, A. V., Gulevich, R. G., et al. (2018). Genomic responses to selection for tame/aggressive behaviors in the silver fox (Vulpes vulpes). *PNAS* 115, 10398–10403. doi:10.1073/pnas.1800889115.

Ward, J., Strawbridge, R. J., Bailey, M. E. S., Graham, N., Ferguson, A., Lyall, D. M., et al. (2017). Genome-wide analysis in UK Biobank identifies four loci associated with mood instability and genetic correlation with major depressive disorder, anxiety disorder and schizophrenia. *Transl Psychiatry* 7. doi:10.1038/s41398-017-0012-7.

Watanabe, Y., Nunokawa, A., Kaneko, N., and Someya, T. (2007). The tryptophan hydroxylase 1 (TPH1) gene and risk of schizophrenia: a moderate-scale case-control study and meta-analysis. *Neurosci. Res.* 59, 322–326. doi:10.1016/j.neures.2007.08.002.

Watkeys, O. J., Kremerskothen, K., Quidé, Y., Fullerton, J. M., and Green, M. J. (2018). Glucocorticoid receptor gene (NR3C1) DNA methylation in association with trauma, psychopathology, transcript expression, or genotypic variation: A systematic review. *Neurosci Biobehav Rev* 95, 85–122. doi:10.1016/j.neubiorev.2018.08.017.

Watson, S., Gallagher, P., Ritchie, J. C., Ferrier, I. N., and Young, A. H. (2004). Hypothalamic-pituitary-adrenal axis function in patients with bipolar disorder. *The British Journal of Psychiatry* 184, 496–502. doi:10.1192/bjp.184.6.496.

Wayne, R. K., and vonHoldt, B. M. (2012). Evolutionary genomics of dog domestication. *Mamm Genome* 23, 3–18. doi:10.1007/s00335-011-9386-7.

Weaver, A. M. (2008). Cortactin in tumor invasiveness. *Cancer Lett.* 265, 157–166. doi:10.1016/j.canlet.2008.02.066.

Weedon, M. N., Lango, H., Lindgren, C. M., Wallace, C., Evans, D. M., Mangino, M., et al. (2008). Genome-wide association analysis identifies 20 loci that influence adult height. *Nat Genet* 40, 575–583. doi:10.1038/ng.121.

Whibley, A., Urquhart, J., Dore, J., Willatt, L., Parkin, G., Gaunt, L., et al. (2010). Deletion of MAOA and MAOB in a male patient causes severe developmental delay, intermittent hypotonia and stereotypical hand movements. *Eur. J. Hum. Genet.* 18, 1095–1099. doi:10.1038/ejhg.2010.41.

Wilkins, A. S., Wrangham, R. W., and Fitch, W. T. (2014). The "Domestication Syndrome" in Mammals: A Unified Explanation Based on Neural Crest Cell Behavior and Genetics. *Genetics* 197, 795–808. doi:10.1534/genetics.114.165423.

Wilkinson, S., Lu, Z. H., Megens, H.-J., Archibald, A. L., Haley, C., Jackson, I. J., et al. (2013). Signatures of Diversifying Selection in European Pig Breeds. *PLoS Genet* 9. doi:10.1371/journal.pgen.1003453.

Wilson, P. M., Fryer, R. H., Fang, Y., and Hatten, M. E. (2010). Astn2, A Novel Member of the Astrotactin Gene Family, Regulates the Trafficking of ASTN1 during Glial-Guided Neuronal Migration. *J Neurosci* 30, 8529–8540. doi:10.1523/JNEUROSCI.0032-10.2010.

Wilson, S. T., Stanley, B., Brent, D. A., Oquendo, M. A., Huang, Y., Haghighi, F., et al. (2012). Interaction between tryptophan hydroxylase I polymorphisms and childhood abuse is associated with increased risk for borderline personality disorder in adulthood. *Psychiatr. Genet.* 22, 15–24. doi:10.1097/YPG.0b013e32834c0c4c.

Witt, S. H., Juraeva, D., Sticht, C., Strohmaier, J., Meier, S., Treutlein, J., et al. (2014). Investigation of manic and euthymic episodes identifies state- and trait-specific gene expression and STAB1 as a new candidate gene for bipolar disorder. *Transl Psychiatry* 4, e426. doi:10.1038/tp.2014.71.

Wolf, C., Mohr, H., Schneider-Axmann, T., Reif, A., Wobrock, T., Scherk, H., et al. (2014). CACNA1C genotype explains interindividual differences in amygdala volume among patients with schizophrenia. *Eur Arch Psychiatry Clin Neurosci* 264, 93–102. doi:10.1007/s00406-013-0427-y.

Wonodi, I., Hong, L. E., Avila, M. T., Buchanan, R. W., Carpenter, W. T., Stine, O. C., et al. (2005). Association between polymorphism of the SNAP29 gene promoter region and schizophrenia. *Schizophr. Res.* 78, 339–341. doi:10.1016/j.schres.2005.03.023.

Woodward, E. R., Eng, C., McMahon, R., Voutilainen, R., Affara, N. A., Ponder, B. A., et al. (1997). Genetic predisposition to phaeochromocytoma: analysis of candidate genes GDNF, RET and VHL. *Hum. Mol. Genet.* 6, 1051–1056. doi:10.1093/hmg/6.7.1051.

Wrzal, P. K., Goupil, E., Laporte, S. A., Hébert, T. E., and Zingg, H. H. (2012). Functional interactions between the oxytocin receptor and the β2-adrenergic receptor: implications for ERK1/2 activation in human myometrial cells. *Cell. Signal.* 24, 333–341. doi:10.1016/j.cellsig.2011.09.019.

Xu, H., Xiao, B., Ji, X., Hu, Q., Chen, Y., and Qiu, W. (2014). Nonmosaic tetrasomy 15q25.2 → qter identified with SNP microarray in a patient with characteristic facial appearance and review of the literature. *European Journal of Medical Genetics* 57, 329–333. doi:10.1016/j.ejmg.2014.04.011.

Yadav, R., Gupta, S. C., Hillman, B. G., Bhatt, J. M., Stairs, D. J., and Dravid, S. M. (2012). Deletion of Glutamate Delta-1 Receptor in Mouse Leads to Aberrant Emotional and Social Behaviors. *PLoS One* 7. doi:10.1371/journal.pone.0032969.

Yamada, M., Clark, J., McClelland, C., Capaldo, E., Ray, A., and Iulianella, A. (2015). Cux2 activity defines a subpopulation of perinatal neurogenic progenitors in the hippocampus. *Hippocampus* 25, 253–267. doi:10.1002/hipo.22370.

Yamakawa, K., Huot, Y. K., Haendelt, M. A., Hubert, R., Chen, X. N., Lyons, G. E., et al. (1998). DSCAM: a novel member of the immunoglobulin superfamily maps in a Down syndrome region and is involved in the development of the nervous system. *Hum. Mol. Genet.* 7, 227–237. doi:10.1093/hmg/7.2.227.

Yan, J., Noltner, K., Feng, J., Li, W., Schroer, R., Skinner, C., et al. (2008). Neurexin 1alpha structural variants associated with autism. *Neurosci. Lett.* 438, 368–370. doi:10.1016/j.neulet.2008.04.074.

Yang, H., Kim, J., Kim, Y., Jang, S.-W., Sestan, N., and Shim, S. (2020a). Cux2 expression regulated by Lhx2 in the upper layer neurons of the developing cortex. *Biochemical and Biophysical Research Communications* 521, 874–879. doi:10.1016/j.bbrc.2019.11.004.

Yang, J. Z., Si, T. M., Ruan, Y., Ling, Y. S., Han, Y. H., Wang, X. L., et al. (2003). Association study of neuregulin 1 gene with schizophrenia. *Mol. Psychiatry* 8, 706–709. doi:10.1038/sj.mp.4001377.

Yang, S., Guo, X., Dong, X., Han, Y., Gao, L., Su, Y., et al. (2017). GABA A receptor subunit gene polymorphisms predict symptom-based and developmental deficits in Chinese Han children and adolescents with autistic spectrum disorders. *Scientific Reports* 7, 1–9. doi:10.1038/s41598-017-03666-0.

Yang, Z., Zhou, D., Li, H., Cai, X., Liu, W., Wang, L., et al. (2020b). The genome-wide risk alleles for psychiatric disorders at 3p21.1 show convergent effects on mRNA expression, cognitive function, and mushroom dendritic spine. *Molecular Psychiatry* 25, 48–66. doi:10.1038/s41380-019-0592-0.

Yehuda, R., Halligan, S. L., Grossman, R., Golier, J. A., and Wong, C. (2002). The cortisol and glucocorticoid receptor response to low dose dexamethasone administration in aging combat veterans and holocaust survivors with and without posttraumatic stress disorder. *Biol. Psychiatry* 52, 393–403. doi:10.1016/s0006-3223(02)01357-4.

Yin, D.-M., Chen, Y.-J., Lu, Y.-S., Bean, J. C., Sathyamurthy, A., Shen, C., et al. (2013). Reversal of behavioral deficits and synaptic dysfunction in mice overexpressing neuregulin 1. *Neuron* 78, 644–657. doi:10.1016/j.neuron.2013.03.028.

Young, E. A., Watson, S. J., Kotun, J., Haskett, R. F., Grunhaus, L., Murphy-Weinberg, V., et al. (1990). Beta-lipotropin-beta-endorphin response to low-dose ovine corticotropin releasing factor in endogenous depression. Preliminary studies. *Arch. Gen. Psychiatry* 47, 449–457. doi:10.1001/archpsyc.1990.01810170049008.

Yu, H., Bi, W., Liu, C., Zhao, Y., Zhang, D., and Yue, W. (2014a). A hypothesis-driven pathway analysis reveals myelin-related pathways that contribute to the risk of schizophrenia and bipolar disorder. *Progress in Neuro-Psychopharmacology and Biological Psychiatry* 51, 140–145. doi:10.1016/j.pnpbp.2014.01.006.

Yu, H.-S., Shen, Y.-H., Yuan, G.-X., Hu, Y.-G., Xu, H.-E., Xiang, Z.-H., et al. (2011). Evidence of Selection at Melanin Synthesis Pathway Loci during Silkworm Domestication. *Mol Biol Evol* 28, 1785–1799. doi:10.1093/molbev/msr002.

Yu, L., Arbez, N., Nucifora, L. G., Sell, G. L., Delisi, L. E., Ross, C. A., et al. (2014b). A mutation in NPAS3 segregates with mental illness in a small family. *Mol. Psychiatry* 19, 7–8. doi:10.1038/mp.2012.192.

Zadeh, N., and John M Graham, J. (2017). *KCNK9 Imprinting Syndrome*. University of Washington, Seattle Available at: https://www.ncbi.nlm.nih.gov/books/NBK425128/ [Accessed February 10, 2020].

Zandi, P. P., Belmonte, P. L., Willour, V. L., Goes, F. S., Badner, J. A., Simpson, S. G., et al. (2008a). Association Study of Wnt Signaling Pathway Genes in Bipolar Disorder. *Arch Gen Psychiatry* 65, 785–793. doi:10.1001/archpsyc.65.7.785.

Zapata, I., Serpell, J. A., and Alvarez, C. E. (2016). Genetic mapping of canine fear and aggression. *BMC Genomics* 17, 572. doi:10.1186/s12864-016-2936-3.

Zavala, J., Ramirez, M., Medina, R., Heard, P., Carter, E., Crandall, A., et al. (2010). Psychiatric syndromes in individuals with chromosome 18 abnormalities. *Am. J. Med. Genet. B Neuropsychiatr. Genet.* 153B, 837–845. doi:10.1002/ajmg.b.31047.

Zaveri, H. P., Beck, T. F., Hernández-García, A., Shelly, K. E., Montgomery, T., Haeringen, A. van, et al. (2014). Identification of Critical Regions and Candidate Genes for Cardiovascular Malformations and Cardiomyopathy Associated with Deletions of Chromosome 1p36. *PLOS ONE* 9, e85600. doi:10.1371/journal.pone.0085600.

Zeidán-Chuliá, F., de Oliveira, B.-H. N., Casanova, M. F., Casanova, E. L., Noda, M., Salmina, A. B., et al. (2016). Up-Regulation of Oligodendrocyte Lineage Markers in the Cerebellum of Autistic Patients: Evidence from Network Analysis of Gene Expression. *Mol Neurobiol* 53, 4019–4025. doi:10.1007/s12035-015-9351-7.

Zhan, C. (2018). POMC Neurons: Feeding, Energy Metabolism, and Beyond. *Adv. Exp. Med. Biol.* 1090, 17–29. doi:10.1007/978-981-13-1286-1_2.

Zhang, B., Jia, Y., Yuan, Y., Yu, X., Xu, Q., Shen, Y., et al. (2004). No association between polymorphisms in the DDC gene and paranoid schizophrenia in a northern Chinese population. *Psychiatr. Genet.* 14, 161–163. doi:10.1097/00041444-200409000-00008.

Zhang, P., Li, G., Li, H., Tan, X., and Cheng, H.-Y. M. (2017a). Environmental perturbation of the circadian clock during pregnancy leads to transgenerational mood disorder-like behaviors in mice. *Scientific Reports* 7, 1–14. doi:10.1038/s41598-017-13067-y.

Zhang, S., Wang, W., Li, J., Cheng, K., Zhou, J., Zhu, D., et al. (2016). Behavioral characterization of CD36 knockout mice with SHIRPA primary screen. *Behav. Brain Res.* 299, 90–96. doi:10.1016/j.bbr.2015.11.027.

Zhang, T., Hou, L., Chen, D. T., McMahon, F. J., Wang, J.-C., and Rice, J. P. (2018a). Exome sequencing of a large family identifies potential candidate genes contributing risk to bipolar disorder. *Gene* 645, 119–123. doi:10.1016/j.gene.2017.12.025.

Zhang, X., Beaulieu, J.-M., Gainetdinov, R. R., and Caron, M. G. (2006). Functional polymorphisms of the brain serotonin synthesizing enzyme tryptophan hydroxylase-2. *Cell Mol Life Sci* 63, 6–11. doi:10.1007/s00018-005-5417-4.

Zhang, X., Zhang, C., Wu, Z., Wang, Z., Peng, D., Chen, J., et al. (2013). Association of genetic variation in CACNA1C with bipolar disorder in Han Chinese. *J Affect Disord* 150, 261–265. doi:10.1016/j.jad.2013.04.004.

Zhang, Y., Liu, Y., Zarrei, M., Tong, W., Dong, R., Wang, Y., et al. (2018b). Association of IMMP2L deletions with autism spectrum disorder: A trio family study and meta-analysis. *Am. J. Med. Genet. B Neuropsychiatr. Genet.* 177, 93–100. doi:10.1002/ajmg.b.32608.

Zhang, Z., Duan, Y., Wu, Z., Zhang, H., Ren, J., and Huang, L. (2017b). PPARD is an Inhibitor of Cartilage Growth in External Ears. *Int J Biol Sci* 13, 669–681. doi:10.7150/ijbs.19714.

Zhang, Z., Jia, Y., Almeida, P., Mank, J. E., van Tuinen, M., Wang, Q., et al. (2018c). Whole-genome resequencing reveals signatures of selection and timing of duck domestication. *Gigascience* 7. doi:10.1093/gigascience/giy027.

Zhou, Q., Eldakhakhny, S., Conforti, F., Crosbie, E. J., Melino, G., and Sayan, B. S. (2018). Pir2/Rnf144b is a potential endometrial cancer biomarker that promotes cell proliferation. *Cell Death Dis* 9. doi:10.1038/s41419-018-0521-1.

Zhou, W.-Z., Zhang, J., Li, Z., Lin, X., Li, J., Wang, S., et al. (2019). Targeted resequencing of 358 candidate genes for autism spectrum disorder in a Chinese cohort reveals diagnostic potential and genotype-phenotype correlations. *Hum. Mutat.* 40, 801–815. doi:10.1002/humu.23724.

Zhou, X., Wang, L., Hasegawa, H., Amin, P., Han, B.-X., Kaneko, S., et al. (2010). Deletion of PIK3C3/Vps34 in sensory neurons causes rapid neurodegeneration by disrupting the endosomal but not the autophagic pathway. *Proc Natl Acad Sci U S A* 107, 9424–9429. doi:10.1073/pnas.0914725107.

Ziegler, C., and Domschke, K. (2018). Epigenetic signature of MAOA and MAOB genes in mental disorders. *J Neural Transm (Vienna)* 125, 1581–1588. doi:10.1007/s00702-018-1929-6.

Zill, P., Büttner, A., Eisenmenger, W., Möller, H.-J., Ackenheil, M., and Bondy, B. (2007). Analysis of tryptophan hydroxylase I and II mRNA expression in the human brain: a post-mortem study. *J Psychiatr Res* 41, 168–173. doi:10.1016/j.jpsychires.2005.05.004.

Zurek, A. A., Kemp, S. W. P., Aga, Z., Walker, S., Milenkovic, M., Ramsey, A. J., et al. (2016). α5GABAA receptor deficiency causes autism-like behaviors. *Ann Clin Transl Neurol* 3, 392–398. doi:10.1002/acn3.303.

Zweier, C., de Jong, E. K., Zweier, M., Orrico, A., Ousager, L. B., Collins, A. L., et al. (2009). CNTNAP2 and NRXN1 Are Mutated in Autosomal-Recessive Pitt-Hopkins-like Mental Retardation and Determine the Level of a Common Synaptic Protein in Drosophila. *Am J Hum Genet* 85, 655–666. doi:10.1016/j.ajhg.2009.10.004.
